# PBX1 and PBX3 transcription factors regulate *SHH* expression in the Frontonasal Ectodermal Zone through complementary mechanisms

**DOI:** 10.1101/2024.06.04.597450

**Authors:** Chan Hee Mok, Diane Hu, Marta Losa, Maurizio Risolino, Licia Selleri, Ralph S. Marcucio

## Abstract

Sonic hedgehog (SHH) signaling from the frontonasal ectodermal zone (FEZ) is a key regulator of craniofacial morphogenesis. Along with SHH, pre-B-cell leukemia homeobox (PBX) transcription factors regulate midfacial development. PBXs act in the epithelium during fusion of facial primordia, but their specific interactions with *SHH* have not been fully investigated. We hypothesized that PBX1/3 regulate *SHH* expression in the FEZ by activating or repressing transcription. The hypothesis was tested by manipulating *PBX1/3* expression in chick embryos and profiling epigenomic landscapes at early developmental stages. *PBX1/3* expression was perturbed in the chick face beginning at stage 10 (HH10) using RCAS viruses, and the resulting *SHH* expression was assessed at HH22. Overexpressing *PBX1* expanded *SHH* expression, while overexpressing *PBX3* decreased *SHH* expression. Conversely, reducing *PBX1* expression decreased *SHH* expression, but reducing *PBX3* induced ectopic *SHH* expression. We performed ATAC-seq and mapped binding of PBX1 and PBX3 with ChIP-seq on the FEZ at HH22 to assess direct interactions of PBX1/3 with the *SHH* locus. These multi-omics approaches uncovered a 400 bp PBX1-enriched element within intron 1 of *SHH* (chr2:8,173,222-8,173,621). Enhancer activity of this element was demonstrated by electroporation of reporter constructs *in ovo* and luciferase reporter assays *in vitro*. When bound by PBX1, this element upregulates transcription, while it downregulates transcription when bound by PBX3. The present study identifies a *cis-*regulatory element, named SFE1, that interacts with PBX1/3 to modulate *SHH* expression in the FEZ and establishes that PBX1 and PBX3 play complementary roles in *SHH* regulation during embryonic development.

## Introduction

During facial development, signaling among the forebrain, the surface cephalic ectoderm, and the neural crest controls morphogenesis of the upper jaw [1–7]. In the chick, signals from the brain [8] and the neural crest cells [9] act sequentially to induce expression of Sonic hedgehog (*SHH*) in the surface cephalic ectoderm covering the Frontonasal Process (FNP), where it forms a boundary with a domain of ectoderm expressing Fibroblast growth factor 8 (*FGF8*). This ectoderm comprises a signaling center that we named the Frontonasal Ectodermal Zone (FEZ) [10]. This signaling center is highly conserved among amniotes and regulates patterned development of the upper jaw [10]. Differences in the shape of the *SHH* domain in the FEZ contribute to different patterns of morphogenesis among embryos [11]. Hence, understanding how the spatial domain of *SHH* expression is established is important for understanding morphogenesis of the embryonic upper jaw and thus, more broadly, of the head. However, molecular regulation of the pattern of *SHH* expression in the FEZ is not known.

Control of *SHH* expression in the FEZ appears to be a two-step process^2,3^. Initially, either *SHH* or a *SHH* dependent signal from the brain to the cephalic ectoderm is required prior to Hamburger-Hamilton stage [12] 17 (HH17). Then as neural crest cells colonize the FNP, *SHH* expression is induced in the FEZ. *SHH* expression is confined to the ventral ectoderm of the mouth cavity by HH22 and becomes restricted to the tip of the upper jaw by HH26. Additional *SHH* expression domains also appear, including its expansion into the ectoderm of the maxillary process and the globular process of the FNP by HH25^15^.

To date, little is known about the regulatory landscape controlling *SHH* expression in the FEZ. However, in other tissues *Shh* expression is regulated by enhancers located within the gene locus itself or via multiple long-range enhancers located up to 1 Mb upstream of *Shh*. For example, in the neural tube of the mouse embryo three distinct but adjacent regulatory motifs are present within the *Shh* locus or just upstream of the promoter region: *Shh* floor plate enhancers (SFPE1/2) and *Shh* brain enhancer 1 (SBE1) [13]. In the mouse embryonic brain, a series of SBEs (SBE2/3/4/6/7) regulate *Shh* expression. These enhancers are located in the intergenic region between *Shh* and *Lmbr1* and in an intron of *Lmbr1,* which is about 1Mb away from *Shh* [14–18]. During murine oral and dental development, MRCS1 and MFCS4 regulate *Shh* expression [19, 20]. In addition, a regulatory sequence for *Shh* expression in endodermal organs (SLGE) is located 100 kb upstream of *Shh* [21]. Lastly, In the mouse embryonic limb bud, the zone of polarizing activity regulatory sequence (ZRS) is positioned in intron 5 of *Lmbr1*, which is 1 Mb upstream of *Shh* [22]. It is thought that chromatin folding of the *Shh* Topologically associating domain (TAD) brings these distant regulatory elements in proximity of the *Shh* promoter to initiate transcription [23].

Interestingly, pre-B-cell leukemia homeobox (PBX) transcription factors control *Shh* expression during mouse limb patterning [24, 25]. PBX homeoproteins control spatial distribution of *Hox* genes and, in turn, *Shh* expression in the posterior limb mesenchyme during mouse limb development^8^. *Shh* is not expressed in the hindlimb buds of *Pbx1/2* mutant embryos from E9.5 to E13.5, and levels of *Hoxa/Hoxd* gene transcripts were significantly decreased or absent in future *Shh*-positive domains before the onset of *Shh* expression. Thus, we hypothesized that PBX transcription factors could similarly participate in the regulation of *SHH/Shh* expression in the avian and mammalian face. In addition, *Pbx* genes control the morphogenesis of the mouse primary palate [26]. Indeed, conditional inactivation of murine *Pbx1* in the cephalic epithelium or mesenchyme, respectively, on a *Pbx2-* or *Pbx3-*deficient background, demonstrated that PBX transcription factors are essential in the epithelium and dispensable in the mesenchyme for upper lip/primary palate fusion [26]. Accordingly, loss-of-function of PBX proteins in the murine cephalic epithelium, but not in the mesenchyme, results in cleft lip/palate (CL/P). However, despite the critical functions of PBX transcription factors in the craniofacial epithelium, potential direct roles of PBX homeoproteins in regulating *SHH* expression in the FEZ have not been investigated so far.

Given the known functions of PBX transcription factors in regulating *Shh* expression in the murine limb bud and their roles in primary palate morphogenesis, we set out to evaluate potential PBX-dependent mechanisms that regulate expression of *SHH* in the avian FEZ. As birds do not have a gene encoding PBX2, we hypothesized that *PBX1* and *PBX3* participate in regulating *SHH* expression in the FEZ by activating or repressing *SHH* transcription. We tested our working hypothesis by manipulating *PBX1* and *PBX3* expression in the FEZ and assessing the chromatin landscape at the *SHH* locus in developing avian embryos.

## Results

### *PBX1* and *PBX3* have complementary expression patterns that delineate *SHH* expression in the FEZ

To begin, we analyzed expression of *SHH* relative to *PBX1* and *PBX3* by *in situ* hybridization in chick embryonic faces at HH22 (n=6). At this developmental stage, the FEZ is active, and the FNP is undergoing patterned outgrowth to form the upper jaw. *SHH* is strongly expressed in the roof of the developing mouth (Fig 1A). At this same time, *PBX1* expression overlaps the *SHH* expression domain (Fig 1B). In contrast, *PBX3* is expressed in the globular process and the dorsal surface of the FNP tip just outside of the *SHH* expression domain, in a manner that appears to define the dorsal boundary of *SHH* expression in the FEZ (Fig 1C). This spatial pattern prompted us to assess whether *PBX1* and *PBX3* play complementary roles in regulating *SHH* expression in the FEZ and help delineate the boundaries of *SHH* expression in this region.

### PBX1 and PBX3 exert opposite effects on *SHH* expression in the FEZ

To assess the extent to which PBX1 and PBX3 may regulate the pattern of *SHH* expression in the FEZ, we used both gain- and loss-of-function approaches. First, we over-expressed each gene using a replication competent retroviral vector (RCAS) encoding PBX1 or PBX3 (RCAS-*PBX1* and RCAS-*PBX3*; Fig 2A, 2B, 2F, and 2G). Second, we knocked down each gene using an RCAS vector that encodes a microRNA targeting either *PBX1* or *PBX3* transcripts (RCAS-mir*PBX1* and RCAS-mir*PBX3*; Fig 2C, 2D, 2H, and 2I). In both experiments RCAS-AP (expressing alkaline phosphatase) was used as a control for retroviral infection and spread (Fig 2E and 2J). Embryos (n=12) were infected at HH10 by injecting viral supernatant (titer, 10^8^) into the mesenchyme adjacent to the anterior neural tube. This time point was chosen as it precedes the onset of *SHH* expression and leads to widespread infection of the face. Embryos were allowed to develop for additional 72 hours (~HH22) and then collected for whole mount *in situ* hybridization and RT-qPCR. Overexpression of *PBX1* led to upregulation of *SHH* expression and expansion of the *SHH* expression domain (Fig 2A), while knock-down of *PBX1* reduced *SHH* expression in the FEZ (Fig 2C). In contrast, overexpression of *PBX3* led to downregulation of *SHH* in the FEZ as well as facial malformations including a smaller head (Fig 2B), and knock-down of *PBX3* led to an expansion of *SHH* expression and premature/ectopic expression of *SHH* in the globular process (Fig 2D; red arrow). At the same time, both overexpression and knock-down of *PBX1* or *PBX3* did not affect *SHH* expression in the brain (Fig 2F – 2I). Expression of *SHH* in the FEZ from each treatment group was quantified by RT-qPCR, and the results were concordant with the *in situ* hybridization data (Fig 2K). Both RCAS-AP and RCAS-mirLacZ controls did not affect *SHH* expression compared to the normal control (Fig 2K). We confirmed with RT-qPCR that RCAS-mir*PBX1* decreased expression of *PBX1* and RCAS-mir*PBX3* decreased expression of *PBX3* (Fig 2L and 2M). To ensure that the gene expression changes did not result from apoptosis after RCAS-miRNA virus infection, we used terminal deoxynucleotidyl transferase dUTP nick end labeling (TUNEL) assays in chick embryonic faces and determined that no cell death was apparent 24 hours after infection (~HH14/15, n=6; S1 Fig).

Together, these data suggest that *PBX1* is a transcriptional activator and *PBX3* a transcriptional repressor of *SHH* expression in the FEZ. However, whether this is direct via transcription factor binding to PBX consensus sites in the *SHH* locus was unexplored and became the focus of the rest of the study.

### Profiling epigenomic landscapes at the *SHH* locus by high-throughput sequencing

We assessed whether PBX1 and PBX3 directly participate in transcriptional regulation of *SHH* expression in cells comprising the FEZ (Fig 3A). First, we performed genome-wide assays for transposase-accessible chromatin using sequencing (ATAC-Seq, n=2) to profile open chromatin regions near the *SHH* locus. Second, we performed chromatin immunoprecipitation followed by sequencing (ChIP-seq, n=2) to identify PBX1 and PBX3 binding to DNA genome-wide and specifically to the *SHH* locus. We then intersected these datasets to identify regions of open chromatin that were bound by PBX1 and/or PBX3.

First, ATAC-seq profiles were used to evaluate open chromatin. For these data sets, the percentage of reads mapped onto the reference genome (mapping rate) were greater than 95%, and the number of unique reads (sequencing depth) ranged from 585,502,859 – 707,110,287 reads (Table S1). Open chromatin peak profiles from two biological replicates resulted in 65,556 peaks that have an Irreproducibility Discovery Rate (IDR) [27] less than 0.05 (Table S1). These open chromatin regions were used for subsequent intersections with ChIP-seq data sets.

Next, we performed ChIP-seq using specific antibodies directed against PBX1 and PBX3. For these data, mapping rates ranged from 88.56 – 98.18%, and the sequencing depth ranged from 25,146,384 – 90,585,798 reads (Table S1). PBX1 peak profiles from two biological replicates resulted in 13,430 overlapping peaks (IDR<0.05), and PBX3 peak profiles from two biological replicates resulted in overlapping 36,122 peaks (IDR<0.05; Table S1). PBX1 and PBX3 peaks (IDR<0.05) were then analyzed to identify differentially bound sites by DiffBind [28, 29]. Differentially bound peak profiles of PBX1 and PBX3 underwent the KEGG pathway enrichment analysis [30, 31], and each PBX protein’s binding profiles resulted in its own exclusive overrepresented pathways (Fig 3B). From their top 12 overrepresented biological processes, PBX1 and PBX3 binding profiles shared seven biological processes: regulation of actin cytoskeleton, salmonella infection, focal adhesion, Toll-like receptor signaling pathway, FoxO signaling pathway, TGF-beta signaling pathway, and melanogenesis. While the other five pathways were exclusive to either PBX1- or PBX3-specific binding signatures: cell cycle, citrate cycle (TCA cycle), mitophagy, apoptosis, and cellular senescence for PBX1 profiles; and cytokine-cytokine receptor interaction, neuroactive ligand-receptor interaction, C-type lectin receptor signaling pathway, MAPK signaling pathway, and Wnt signaling pathway for PBX3 profiles.

Sequentially, we conducted motif analysis on ATAC-seq peaks with IDR<0.05 by Hypergeometric Optimization of Motif EnRichment (HOMER [32], version 4.11) to identify known motifs. The top three motifs were AYAGTGCCMYCTRGTGGCCA (CTCF, CCCTC-Binding Factor), CNNBRGCGCCCCCTGSTGGC (BORIS, CCCTC-Binding Factor Like), and GKVTCADRTTWC (SIX1, SIX homeobox 1; Fig 2C). PBX3 motif (SCTGTCAMTCAN) ranked 17^th^, and PBX1 motif (GSCTGTCACTCA) ranked 39^th^ (Table S2). Also, the motif analysis was conducted on PBX1 and PBX3 peaks with IDR<0.05 to discover both known and *de novo* motifs (Fig 2D – 2G). The top three known motifs from PBX1 peak profiles were VGCTGWCAVB (MEIS1, Myeloid Ecotropic Integration Site 1), YTGWCADY (TGIF1; TGFB Induced Factor Homeobox 1), and SCTGTCAMTCAN (PBX3). The top three known motifs from PBX3 peak profiles were YTGWCADY (TGIF1), VGCTGWCAVB (MEIS1), and TGTCANYT (TGIF2). The top three *de novo* motifs discovered from PBX1 peak profiles were NRNCTGWCAG, TGATTGRCNG, and RDRGGMGGDR. The top three *de novo* motifs discovered from PBX3 peak profiles were NCTGWCAGNH, TGATTGRCGG, and NAGGAATGYG. Notably, PREP1/PKNOX1 (PBX regulating protein 1, PREP1; also called PBX/Knotted Homeobox 1, PKNOX1) motif is the second best match for *de novo* motif discovery from both PBX1 and PBX3 ChIP-seq datasets. The full lists of motif discovery are shown in supplemental tables: ATAC-seq data (known motif discovery, Table S2), ChIP-seq data targeting PBX1 (known motif discovery, Table S3; *de novo* motif discovery, Table S4), and ChIP-seq data targeting PBX3 (known motif discovery, Table S5; *de novo* motif discovery, Table S6).

Finally, we intersected the replicated peaks from the ATAC-seq and ChIP-seq datasets. All PBX1 and PBX3 binding sequences on the *SHH* locus are located within intronic regions and span ATAC-seq peaks (Fig 3H). In intron 1, PBX1 and PBX3 are bound to the same region, suggesting a potential competitive binding within that region of chromatin. Among PBX1 and PBX3 differentially bound sites, one 400 bp long sequence was identified as a PBX1-enriched region within the first intron of *SHH* (chr2:8,173,222-621; Fig 3H, yellow rectangle). Within this 400 bp PBX1-enriched sequence, the TGIF2 motif (the third known motif from the PBX3 binding profile) was found once (Fig 3J). The third *de novo* motif from the PBX1 binding profiles (RDRGGMGGDR, best match POLII – GC box) was found four times within the 400 bp PBX1-enriched sequence (Fig 3J). This third *de novo* motif from the PBX1 peak profile and the second *de novo* motif from the PBX3 peak profile (TGATTGRCGG) contain GGMGG and GRCGG sequences which match putative MEIS-PREP/PBX1-PREP binding sites [33, 34] (CCCGCCC and GGCGG), and these sequences appear at three regions within the 400 bp PBX1-enriched sequence identified within intron 1 of *SHH* (Fig 3I and 3J). Altogether, these genome-wide high-throughput sequencing data suggest potential interactions between PBX proteins and MEIS-PREP complexes at this 400 bp sequence within intron 1 of *SHH* in the chick FEZ at HH22.

### Activity of the regulatory sequence within the *SHH* locus

With the series of sequencing data analyses reported above, the 400 bp PBX1-enriched sequence (chr2:8,173,222-8,173,621) containing multiple PBX motifs became our region of interest. To test whether this 400 bp genomic sequence has enhancer activity, we cloned a 1.8 kb (chr2: 8,172,590-8,174,390) fragment that included our region of interest (Fig 4A) into the TKp-TK vector. This vector has a minimal promoter with no basal reporter (LacZ) activity and can be used to assess enhancer activity via electroporation into chick embryos [3]. This construct was electroporated into the ectoderm covering the FNP at HH20 (n=20). After 24 hours (HH24), this fragment exhibited transcriptional activity in the *SHH* expression domain of the FEZ suggesting that this element has cis-regulatory functions (Fig 4B).

To identify the minimal region with enhancer activity, we divided the 1.8kb segment (chr2:8,172,590-8,174,390) into three 600 bp fragments (the 3’ end, 3’ of SFE1; the middle, SFE1; the 5’ end, 5’ of SFE1; Fig 5A). Each of these fragments were cloned into a luciferase expression vector and transfected into DF1 cells [35] along with RCAS-*PBX1* (3’ of SFE1, n=7; SFE1, n=6; 5’ of SFE1, n=4; triplicate technical replicates). Luciferase intensity of each experimental group was normalized to the RCAS-AP controls, of which the luciferase expression was normalized to 1 and shown as a red dashed line in Fig 5B – 5D. Luciferase activity was only detected when the fragment containing the 400 bp peak (SFE1; chr2:8,173,096-688) was inserted (P<0.001; Fig 5B). In this region, three PBX consensus binding sites are present (chr2:8,173,380-386; chr2:8,173,425-429; and chr2:8,173,497-503; Fig 3I). Accordingly, we made one construct completely deleting all three PBX consensus sites within SFE1 (referred as “Deleted SFE1”) and a second construct where we mutated all three PBX consensus sites (“Mutated SFE1”; n=3/each). Both deletion and mutation of the putative binding sites significantly decreased luciferase activity compared to the intact 400 bp element (P<0.001) and resulted in no significant luciferase activity compared to the RCAS-AP controls (P>0.05; Fig 5C). The results indicate that PBX1 physically interacts with the three PBX consensus sequences. We named this enhancer *SHH* FEZ Enhancer 1 (SFE1).

In order to confirm the repressive role of *PBX3* on SFE1, we transfected DF1 cells with SFE1 cloned in a luciferase reporter construct with either *PBX3* (n=9) or mir*PBX3* (n=5) expression vectors. Repression of *PBX3* using RCAS-mir*PBX3* in cell culture increased luciferase activity almost two-fold (P<0.0001; Fig 5D). These results suggest that PBX3 plays a repressive role when it interacts with SFE1. We then cloned the SFE1 fragment into the HSP68-lacZ vector that has strong basal expression and transfected it into DF1 cells along with RCAS-*PBX3* (n=4; triplicate technical replicates). A fluorometric β-galactosidase assay demonstrated that SFE1 significantly reduced β-galactosidase fluorescent intensity in the presence of PBX3 compared to the controls (HSP68-lacZ with no insert, shown as a red dashed line; P<0.0001; Fig 5E). Together, these data confirm that PBX3 acts as a repressor of *SHH* when it binds to SFE1, while PBX1 activates transcription from this element.

## Discussion

### SHH as a key molecular regulator in the FEZ and PBX1/3 as potential regulators of *SHH* expression

The FEZ was first identified in the chicken embryo based on the observation of a transient boundary between *SHH* and *FGF8* expression domains in the ectoderm covering the FNP [10]. In the chick FEZ, *SHH* expression begins around HH20 [1, 2] at which time it forms a boundary with a previously established domain of *FGF8* expression. At this time, *FGF8* expression becomes down-regulated across the midline of the FNP and restricted to the nasal pits. In contrast, *SHH* expression is maintained in the FEZ for a long period of time and is required for the FEZ to function. This has been observed in other amniote embryos as well [7]. Interestingly, our data revealed that *PBX1* was co-expressed with *SHH* in the FEZ, while *PBX3* was expressed at the boundary of the *SHH* expression domain. By using gain- and loss-of-function approaches in the chick, we demonstrated that PBX1 induces and PBX3 represses *SHH* transcription. Further, these opposite effects on *SHH* transcription appear to be mediated by a regulatory element in an intronic region of *SHH* that acts as a switch. The same element behaves as an enhancer when bound by PBX1, and as a repressor when bound by PBX3. Hence, PBX1 and PBX3 play complementary roles in the regulation of *SHH* expression in the FEZ during early stages of its function.

### *PBX* genes in development

The *PBX* gene family comprises four transcription factors that have essential roles in embryonic development. *PBX1*, the gene encoding the Pre-B-cell leukemia homeobox transcription factor 1 [36], is the mammalian homologue of the extradenticle (*exd*) gene in *Drosophila melanogaster* and shares over 70% identity to *exd* [37, 38]. *PBX1* encodes a homeodomain transcription factor of the three amino acid loop extension (TALE) family. The PBX1 protein and its related family members PBX2–4 dimerize with other TALE class homeodomain proteins from the MEIS and PREP families through a PBC domain to form nuclear complexes that enhance the binding specificity of HOX proteins to DNA, as demonstrated in some embryonic contexts (reviewed by Moens and Selleri [39]). In the mouse, PBX-PREP or PBX-MEIS complexes regulate target genes that control segment identity and organ patterning during embryogenesis [39–42]. Indeed, the discovery from the present motif analysis supports that PBX1 and PBX3 binding patterns within the *SHH* locus in the FEZ at HH22 are substantially associated with binding of PREP and MEIS complexes. Also, these transcription factors can act upstream of *Hox* genes, or even independently of HOX proteins in collaboration with other cofactors, in various developmental processes [39]. In addition to forming heterodimers with HOX proteins and with TALE partners MEIS/PREP, PBX proteins can form multimeric complexes with other transcription factors, such as MYOD, EN, PDX1 (reviewed by Selleri et al. [43]) and HAND2 [44]. Lastly, PBX proteins can regulate transcription by interacting with basic transcription regulators, such as histone acetyltransferases (HATs) and CBP coactivators, and with histone deacetylases (HDACs) and the corepressor N-CoR/SMRT (reviewed by Selleri et al. [43]). Thus, PBX transcription factors can be part of activating or repressing transcriptional complexes.

In mice *Pbx1* is widely transcribed throughout the developing embryo, and *Pbx1*-deficient embryos that are homozygous for a null allele (*Pbx1*^−/−^) develop pleiotropic developmental defects. These abnormalities include perturbed morphogenesis, severe hypoplasia, or aplasia, of multiple organ systems, including the craniofacial skeleton, ear pinnae, branchial arch-derived structures, limbs, heart, hematopoietic system, lungs, diaphragm, liver, stomach, gut, pancreas, spleen, kidneys, and gonads [25, 40, 45–48]. Heterozygous mice are viable and fertile but are smaller in size than wild type mice [40]. Overall, our previous studies have established that transcription factors of the PBX family are critical developmental regulators. Accordingly, *Pbx* genes play critical roles during morphogenesis of the murine midface, especially in the FNP epithelium, which expresses *Pbx1/2/3* transcripts. Interestingly, PBX transcription factors are essential in the cephalic epithelium but dispensable in the mesenchyme for the proper fusion of the upper lip and the palate [26]. The *Pbx2* gene is absent from the avian genome suggesting that there may be differences in the roles of members of this gene family among organisms. Evolutionary-developmental biology approaches will be needed to elucidate potential unique roles of these transcription factors in different organisms.

### Epigenomic landscapes in the FEZ and a regulatory element within the *SHH* locus

Regions of high sequence conservation across species and with elevated DNaseI hypersensitivity [49, 50] usually correspond to accessible regions of the genome that are functionally related to transcriptional activity, since this remodeled state is necessary for the binding of proteins such as transcription factors. Accessible regions have also been shown to comprise many types of cis-regulatory elements including promoters, enhancers, silencers, and locus control regions. In the present study, we assessed open chromatin sites from the FEZ of chick embryos at HH22 using ATAC-seq. We combined the obtained data sets with results from ChIP-seq assays that profiled genome-wide binding of PBX1 and PBX3 to DNA. These combined high-throughput analyses enabled us to: 1) broadly demarcate the chromatin landscape during FEZ morphogenesis in the chick embryo; and 2) locate a candidate regulatory element within the *SHH* locus that is bound by PBX proteins and controls facial development by regulating *SHH* expression.

In the epigenomic landscapes, all PBX1- and PBX3-bound sites within the *SHH* locus were located at the open, accessible chromatin regions uncovered by ATAC-seq peaks (Fig 3H). Both PBX1 and PBX3 bound to the same region in intron 1 of the *SHH* locus. However, the differentially bound peak analysis determined that PBX1 enrichment at the 400 bp element is greater than PBX3 binding (Fig 3H), suggesting this 400 bp sequence is a candidate regulatory region that is bound preferentially by PBX1. Indeed, upregulation of transcription was observed when PBX1 is bound to the DNA element spanning the PBX1 enriched peak both *in vivo* by electroporation experiments (Fig 4B) and *in vitro* by luciferase reporter assays (Fig 5B). In addition, the reporter assay data also suggest preferential binding of PBX1 to this sequence. Overexpression of PBX3 in cultured cells transfected with the PBX1-bound element cloned upstream of the luciferase reporter neither increased nor decreased luciferase intensity (Fig 5D). This result may indicate that 1) the luciferase assay was simply not sensitive enough to detect reduced expression and/or 2) expression did not change by the additional PBX3 over expression in the presence of endogenous PBX1 proteins. In other words, increased levels of PBX3 proteins did not overcome the binding competition by endogenous PBX1 factors. However, downregulation of PBX3 in this cell culture system increased the luciferase intensity (Fig 5D), indicating that expression was enhanced when competition between PBX1 and PBX3 was altered by decrease of PBX3 protein levels. Reduced PBX3 expression appeared to allow greater enrichment of PBX1 binding to this regulatory sequence, increasing in turn the expression of luciferase. The repressive role of PBX3 was further confirmed by the fluorometric β-galactosidase assay (Fig 5E).

### *PBX* mutations are associated with craniofacial birth defects in humans

There is significant clinical relevance in understanding the roles of PBX transcription factors in regulating facial morphogenesis. Indeed, we reported in a previous study eight patients with craniofacial dysmorphology who had *de novo*, deleterious sequence variants in the *PBX1* gene [51]. The cases exhibited varying expressivity and severity of facial dysmorphology, as well as other affected organ systems. The sequence variants in these cases included missense substitutions adjacent to the PBX1 homeodomain or within the homeodomain, and mutations yielding truncated PBX1 proteins. Functional studies on five *PBX1* sequence variants revealed perturbation of intrinsic, PBX-dependent transactivation ability and altered nuclear translocation, suggesting abnormal interactions between mutant PBX1 proteins and wild-type TALE or HOX cofactors. These mutations may directly affect transcription of PBX1 target genes to impact development [51]. Also, genomic analyses revealed evidence of gene-gene interactions between the human genes *PBX1*/*2* and SNPs in the *ARHGAP29* locus, a candidate for CL/P [52].

In the present study, using the chick embryo, we explored the specific roles of *PBX1* and *PBX3* in regulating FEZ morphogenesis by controlling *SHH* expression. In mammals, *Pbx2* may compensate for some aspects of *Pbx1* loss, thus confounding the potential effect of *Pbx1* loss on *Shh* expression [26, 53, 54]. As discussed above, birds do not appear to have the *PBX2* gene, making the chick embryo an ideal model system to address the functions of *PBX1* and *PBX3*. With the complementary expression patterns of *PBX1* and *PBX3* around the *SHH* domain in normal chick embryos at HH22, we examined how altered *PBX1/3* expression prior to the onset of *SHH* expression would affect the spatial *SHH* domain in the FEZ by HH22. By gain-of-function and loss-of-function experiments in the chick, we showed that over-expressing *PBX1* resulted in increased and expanded *SHH* expression, while repressing *PBX1* decreased *SHH* expression in the FEZ, suggesting that PBX1 is a positive regulator of *SHH* transcription in the FEZ. We can envision that PBX1 binding to the 400 bp *SHH* regulatory element identified in this study could play: 1) an ‘instructive’ role in target gene transcriptional activation; or 2) a ‘permissive’ (pioneer factor) role [55, 56] as a recruiter of cofactors that modify the chromatin, marking control regions for activation by other transcription factors to target enhancer elements of craniofacial developmental genes. In contrast, repressing *PBX3* increased expression of *SHH* in its original expression domains on the ventral surface of the FNP around the mouth cavity and induced premature ectopic expression of *SHH* in the FNP globular process at HH22, where *SHH* expression is observed only by HH25 [57]. Overexpression of *PBX3* resulted in decreased *SHH* expression in the FEZ at HH22 and malformations including a smaller head, which is associated with reduced *SHH*-signaling [1, 2, 8]. Together, our results demonstrate complementary functions of *PBX1/3* in the regulation of *SHH* expression in the developing FNP. Future research will aim at identifying co-factors that partner with PBX homeoproteins to activate or repress *SHH* expression in the FEZ of avian embryos.

## Conclusion

Collectively, our findings demonstrate complementary regulatory mechanisms involving PBX1 and PBX3 transcription factors in the control of *SHH* expression in the chick FEZ. The high-throughput sequencing data sets we generated establish that PBX1 and PBX3 bind to open chromatin regions and specifically interact with a regulatory sequence in a non-coding region within the *SHH* locus in the chicken FEZ. Specifically, PBX1 acts as an activator and PBX3 acts as a repressor of *SHH* expression. Further investigations on the relationships between histone mark binding and PBX binding at the *SHH* locus will provide additional information on PBX/SHH-dependent regulatory mechanisms that drive craniofacial morphogenesis.

## Materials and Methods

### Experimental design

The aim of the present study was to assess regulatory mechanisms involving PBX1 and PBX3 that induce and maintain *SHH* expression in the FEZ during early craniofacial development. The study is divided into two major parts: 1) demonstrating expression patterns of *PBX1*, *PBX3*, and *SHH* in normal chick embryos at HH22 and evaluating *SHH* expression patterns after manipulating *PBX1* and *PBX3* expression and 2) profiling open chromatin configurations and PBX binding sites in the *SHH* locus of the normal chick FEZ at HH22 and testing activity of a potential regulatory sequence.

First, we demonstrated normal expression patterns of *SHH*, *PBX1*, and *PBX3* in the FNP of normal HH22 chick embryos (n=6) by whole-mount *in situ* hybridization. Next, we demonstrated how overexpression and repression of *PBX1* and *PBX3* affect *SHH* in the FEZ by standard gain- and loss-of-function experiments with RCAS virus infection (n=12).

Assessment of accessible chromatin signatures and the binding patterns of PBX1 and PBX3 to regulatory regions of the chick *SHH* locus was done in order to determine the regulatory mechanisms of PBX1 and PBX3 on *SHH* expression in the FEZ. Open chromatin configurations were evaluated by utilizing ATAC-seq (n=2), and PBX binding sites were evaluated by utilizing ChIP-seq (n=2). Then, a potential regulatory sequence analyzed from the high-throughput sequencing data was tested both *in vivo* and *in vitro* by electroporation (n=20) and luciferase assays (n= 3 – 7), respectively.

### Chick embryos

The experimental procedures involving chick embryos met the institutional and national regulatory standards applied for vertebrate embryos. Fertile White Leghorn chicken (*Gallus gallus*) eggs (Petaluma Farms, Petaluma, CA) or SPAFAS Flock-C17 RCAS free eggs from Charles River (Wilmington, MA) were incubated at 38°C until either sample collection or experiments. The embryos were staged using a strategy that relies on external morphological characters and that is independent of body size and incubation time [58, 59]. Specifically, we applied the Hamburger and Hamilton staging system, which was originally developed for chick [12, 60].

### Whole-mount *in situ* hybridization

*In situ* hybridization was performed on whole embryos as described [61]. Briefly, subclones of chick *SHH*, *PBX1* and *PBX3* were linearized to transcribe digoxygenin-labeled antisense riboprobes. The chick embryos underwent hybridization with 0.5 – 1.0 μg/ml digoxygenin-labeled cRNA probes, and, after washing, the embryos were incubated with an alkaline phosphatase-conjugated anti-digoxygenin antibody (Boehringer Ingelheim, Ingelheim, Germany). Nitro blue tetrazolium/5-bromo-4-chloro-3-indolyl phosphate substrate (NBT/BCIP substrate; Roche, Basel, Switzerland) was used for color detection. Stained embryos were observed and imaged by the Leica MZ FLIII Stereomicroscope.

### Gain-of-function and loss-of-function experiments by RCAS virus infection

Chick embryos were incubated until HH10. Then, 1.0 ml of albumin was removed from the egg, and a small window was cut out on the top of the shell to expose the embryo. At HH10, overexpression or knock-down of *PBX1* and *PBX3* in the embryo face was achieved by RCAS virus infection into the mesenchyme adjacent to the anterior neural tube. For overexpression, we engineered RCAS virus to encode each gene (RCAS-*PBX1*, RCAS-*PBX3*, and RCAS-AP as a control). For knock-down experiments, we modified the BLOCK-iT Pol-II RNAi expression vector (Cat #, K493500; Invitrogen, Waltham, MA) to clone into RCAS virus. We designed multiple miRNAs targeting the open reading frame and the 3’UTR of *PBX1* and *PBX3*, as well as scrambled controls using Invitrogen’s BLOCK-iT^TM^ RNAi Designer. In our experience, these constructs significantly reduced toxicity of knock-down by employing an artificial miRNA that is processed as endogenous miRNAs rather than as shRNA. After RCAS infection, embryos were returned to the incubator, and the whole heads were collected at HH22 for whole-mount *in situ* hybridization to detect *SHH* expression (conducted as described above). Also, *SHH* expression from the FEZ at HH22 was quantified by RT-qPCR as described below.

### Terminal deoxynucleotidyl transferase dUTP nick end labeling (TUNEL) assay on embryos infected with miRNA-RCAS virus

To assess apoptotic cells in embryos that were infected with miRNA-RCAS virus, the embryo heads were fixed in 4% paraformaldehyde (PFA) solution, paraffin-embedded, and sectioned at 8 μm. On the sections, DNA fragmentation was visualized by using an *in situ* cell death detection kit (Fluorescein; Cat #, 11684795910; Roche, Basel, Switzerland) following the manufacturer’s instructions.

### Total RNA preparation and RT-qPCR

The FEZ was isolated as described below. RNA was then extracted from the FEZ using the RNeasy Kit (Cat #, 74104; Qiagen, Hilden, Germany). cDNA was synthesized by the Invitrogen Superscript III kit following the manufacturer’s instructions. RT-qPCR was performed using a Bio-Rad CFX 96 real-time PCR machine. The qPCR primers for *SHH*, *PBX1*, and *PBX3* were as below:

*SHH* – forward 5’ GCTGACAGACTGATGACTCA 3’ and reverse 5’ TCGTAGTGCAGCGATTCCTC 3’;

*PBX1* – forward 5’ GGCTACGGAAATCCTGAATGAG 3’ and reverse 5’ ACCAGTTTGATACCTGTGAGAC 3’;

*PBX3* – forward 5’ GCATCGATATGGACGAGCAGTCC 3’ and reverse 5’ GCATCGATTTAGTTAGAGGTATCA 3’.

Relative gene expression was calculated based on the ΔΔCt method. ΔCt was calculated between each target gene and *GAPDH* as an endogenous control (forward 5’ CTGGTATGACAATGAGTTTGG 3’; reverse 5’ ATCAGTTTCTATCAGCCTCTC 3’), and non-infected, normal chick FEZ was used as a control sample for calculating ΔΔCt. The relative quantity (RQ) was then calculated by the equation: RQ=2^-ΔΔCt^. At least three biological replicates were prepared, and one-way analysis of variance (ANOVA) test was used to assess the statistical significance. (Prism 10, GraphPad).

### Frontonasal ectodermal zone isolation

The isolation of the chick FEZ was conducted as previously described [62]. Chick embryo heads at collection points were removed and placed into serum free DMEM, and the FNP was dissected (Fig 3A) so that only the region covered by the FEZ remains. Then, the tissue was digested in dispase (2.4U/ml) in DMEM on ice for 20 minutes. Digestion was quenched by transferring the tissue to DMEM with 1% BSA. Subsequently, using a sharpened tungsten needle, the surface ectoderm was gently lifted off of the FNP. Fresh tissues were used immediately for ATAC-seq, since fixation of cells has been found to reduce transposition frequency and is not recommended [63]. In contrast, tissues were cross-linked and snap-frozen to be stored at −80°C for later use in ChIP-seq assays [64–66].

### ATAC-Sequencing

The FEZ tissues from 10-15 normal chick embryos at HH22 were pooled for each replicate for ATAC-seq analysis (n=2). ATAC-seq was performed on the FEZ cells employing an established two-step protocol that can use a low number of cells (as low as 50,000 cells) [63]. DNA samples were incubated with the Tn5 transposome, which performs both adaptor ligation and fragmentation of open chromatin regions. Paired-end reads (2×150 nt) were generated by HiSeq 4000 (Illumina) at the UCSF Center for Advanced Technology (CAT) Facility. The quality of sequencing was assessed by FastQC (version 0.11.9) [67]. The raw reads were trimmed and mapped onto the latest NCBI chicken reference genome (galGal6) by the local alignment of Bowtie2 (version 2.4.5) [68]. After read alignment, duplicates and unmapped reads were removed by Sambamba (version 0.6.8) [69], and mitochondrial reads were also removed by Samtool (version 1.11) [70]. The filtered reads were then processed by MAC2S [71] to call narrow peaks. For peak calling, the effective genome size of 1.03E+9 was applied (for 150 bp k-mer, calculated with the NCBI chicken genome size), and the statistical threshold was set at q-value (minimum false discovery rate) <0.05. Simple overlaps of peaks from both biological replicate were determined by the intersect function from BEDTools (version 2.31.1) [72]. Irreproducibility discovery rate (IDR) for every peaks between two replicates were calculated, and only peaks with IDR<0.05 were merged as overlapping peaks [27]. The analyzed peaks and overall peak profiles were then visualized on the Integrative Genomics Viewer (IGV viewer, version 2.15.4) [73]. Lastly, the motif analysis on ATAC-seq peaks (IDR<0.05) was conducted by the findMotifsGenome.pl function in the HOMER [32] package to identify known motifs from the open chromatin profiles. The region size setting used for motif discovery was given peak size.

### ChIP-Sequencing targeting PBX1 and PBX3

Since ChIP-seq requires a substantially greater number of cells compared to ATAC-seq, the FEZ tissues from ~80-85 normal chick embryos were pooled for each replicate (n=2). For immunoprecipitation, cells from the FEZ were crosslinked with 1% PFA for 10 min at room temperature and sonicated by 30 cycles (1 min/cycle; reset every 5 min). The cells were then incubated with Pbx1 antibody (Cat #, 4342; Cell Signaling Technology, Danvers, MA) or PBX3 antibody (Cat #, 12571-1-AP; Proteintech, Rosemont, IL). Input controls, which were crosslinked and sonicated but immunoprecipitated with any antibody, were also generated for each biological replicate. DNA libraries were constructed using the TruSeq ChIP Sample Preparation Kit (Cat #, IP-202-1012; Illumina, San Diego, CA), and single-end reads (50 nt) were generated by HiSeq 4000 (Illumina) at the UCSF CAT Facility. The same sequencing data analysis pipeline (including quality check, read trimming, alignment, peak calling, and merging peaks between biological replicates) that was used for ATAC-seq was also applied for the ChIP-seq data sets with a few modifications described below. The effective genome size of 1.02E+9 (for 50 bp k-mer, calculated with the same NCBI chicken genome size) was used for peak calling by MAC2S [71] to call narrow peaks (q<0.05). Peaks from each sample were normalized by an input control during peak calling. After merging IDR<0.05 peaks from each experimental group, PBX1- and PBX3-enriched peaks were further analyzed by an R package, DiffBind [28, 29], to identify differential binding affinity. The pathway analysis was conducted based on the KEGG database [30, 31]. Lastly, we conducted the motif analysis on PBX1 and PBX3 peaks with IDR<0.05 by the findMotifsGenome.pl function in the HOMER [32] package to identify both *de novo* and known motifs in our data sets. The region size setting used for motif discovery was 200 bp as recommended for transcription factors by the developers to identify both primary and co-enriched motifs.

Both ATAC-seq and ChIP-seq data are openly accessible through National Center for Biotechnology Information Sequence Read Archive (NCBI SRA; accession number: PRJNA1111924) and FaceBase (https://doi.org/10.25550/5G-0TNJ).

### Electroporation

A reporter vector comprised of a minimal Thymidine kinase promoter, β-galactosidase, and a polyadenylation sequence (TK-p-β-gal) was used for the analyses as the previously established approach [3]. The 1.8kb sequence (chr2: 8,172,590-8,174,390) spanning the 400 bp-long PBX1-enriched region on intron 1 of *SHH* was cloned by Long-Range PCR and then ligated the cloned genomic fragment into TK-p-β-gal. This construct (TK-p-β-gal-1.8kb PBX1 bound intron 1) was electroporated into the developing FEZ of embryos at HH20 (n=20). As a negative control, we electroporated the TK-p-β-gal vector without the cloned intron 1 insert, and as a positive control to assess the extent of transfection, we electroporated a constitutively activated vector comprised of heat shock protein promoter that encodes β-galactosidase (HSP-β-gal). Embryos were then allowed to develop for 24 hours (~HH24), fixed, and assessed for β-galactosidase activity by a standard X-gal reaction as previously described [57].

### Luciferase assay

The 1.8kb sequence (chr2: 8,172,590-8,174,390) spanning the 400 bp PBX1-enriched locus was divided into three 600 bp segments. Each of these segments were transfected into DF1 cells along with a luciferase reporter construct and either of an expression vector (RCAS-*PBX1*, RCAS-*PBX3*, RCAS-mir*PBX1*, RCAS-mir*PBX3*, and RCAS-AP as control). Luciferase intensity was measured by the Luciferase Assay System kit (Cat #, E1500; Promega, Madison, WI) on Promega GloMax Explorer (Cat #, GM3500) and normalized with the RCAS-AP control. One assay includes three technical replicates, and total 3 – 9 inter-assays were conducted for each combination. Outliers were determined by Grubbs’ test (extreme studentized deviate method) and then removed for further data analyses. Then, unpaired t-test was conducted, and the significance was determined when P<0.05 (Prism 10, GraphPad).

### Fluorometric β-galactosidase assay

The middle 600 bp fragment (SFE1; chr2:8,173,096-688) was cloned into the HSP68-LacZ vector with ApaI/HindIII cloning sites. This construct was transfected into DF1 cell cultures either with 300 ng of RCAS-*PBX3* and 50 ng of HSP68-LacZ or 300 ng RCAS-*PBX3* and 50 ng of HSP68-LacZ-SFE1. The transfected cells were cultured for 48 hours. For quantitation of β-galactosidase, we used the sensolyte-mug-beta-galactosidase-assay-kit-fluorimetric (Cat #, AS-72132; Eurogentec, Seraing, Belgium). Fluorescence signal was recorded on Promega GloMax Explorer (Cat #, GM3500) at EX/EM=365/445 (four inter-assays with triplicate technical replicates). The relative fluorescent intensity underwent unpaired t-test, and the statistical significance was determined when P<0.05 (Prism 10, GraphPad).

## Acknowledgment

This research was funded by the National Institutes of Health/National Institute of Dental and Craniofacial Research (NIH/NIDCR; R01 DE028324 to R.S.M. and L.S.). We would like to thank the Center for Advanced Technology Facility (the Facility Director, Dr. Eric Chow) at the University of California, San Francisco, for their sequencing service.

---

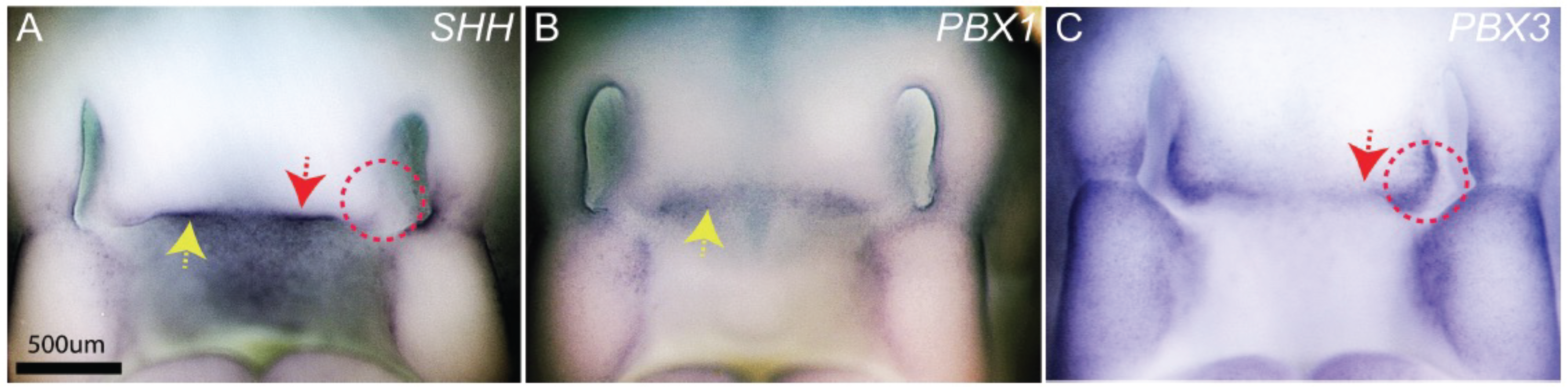

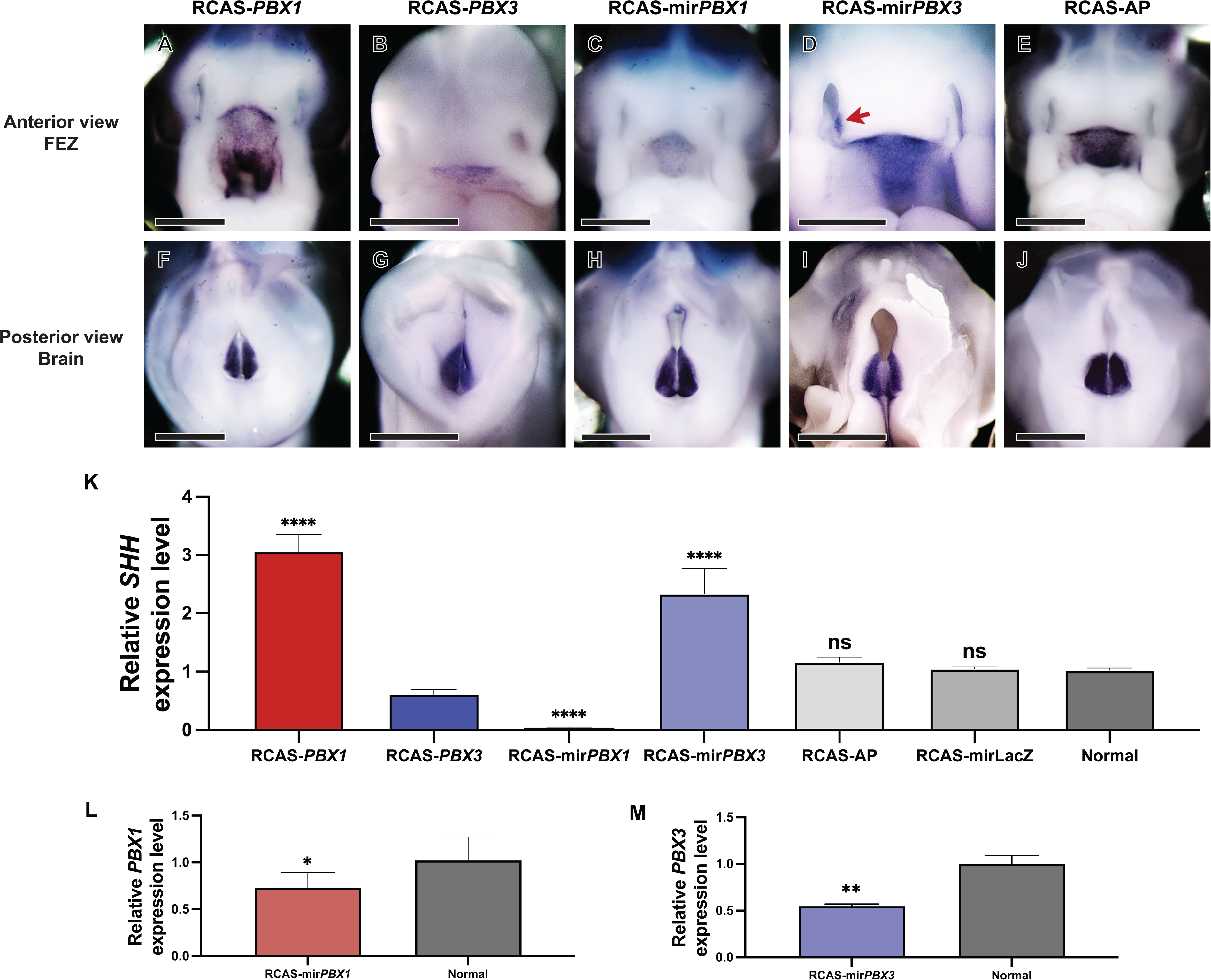

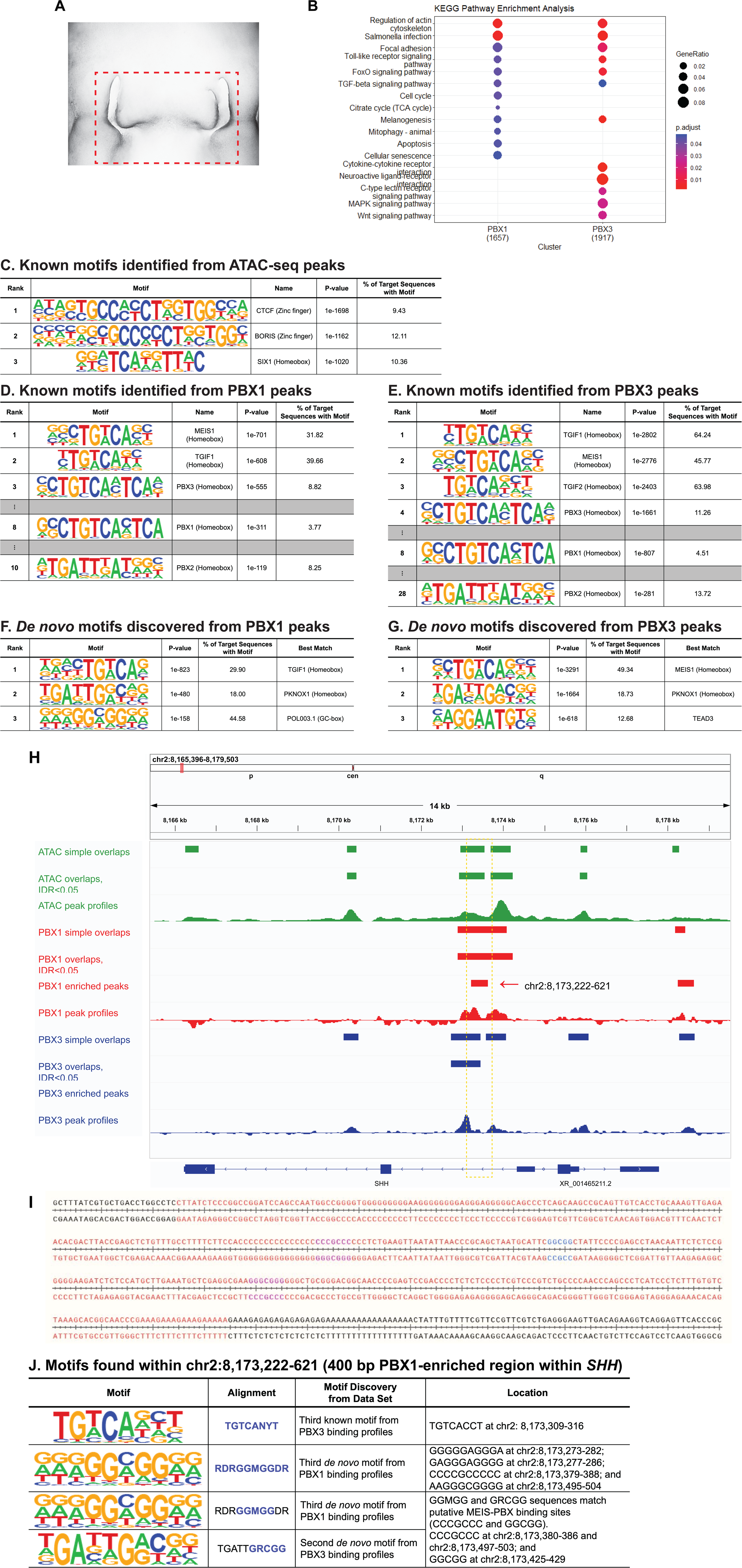

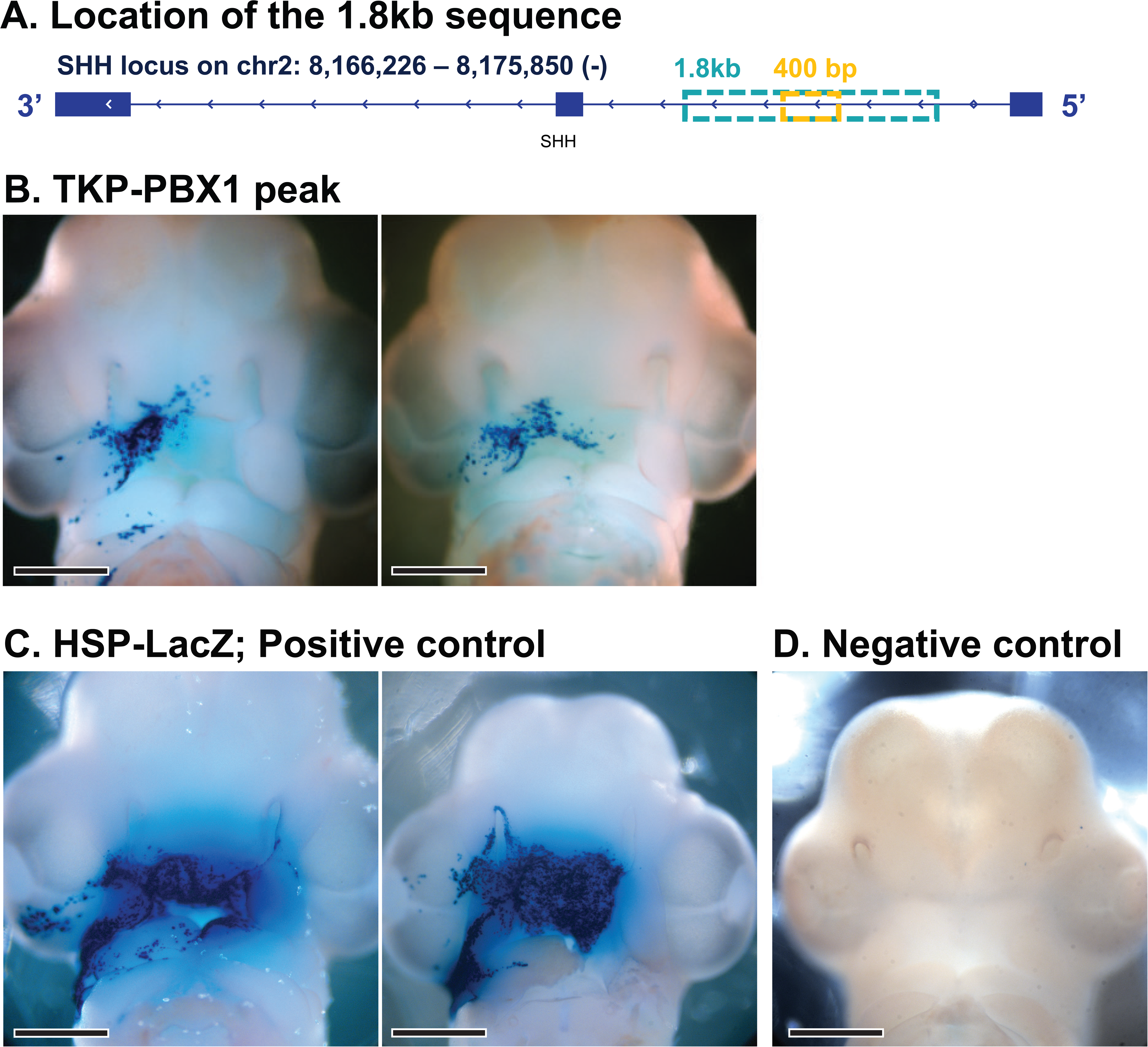

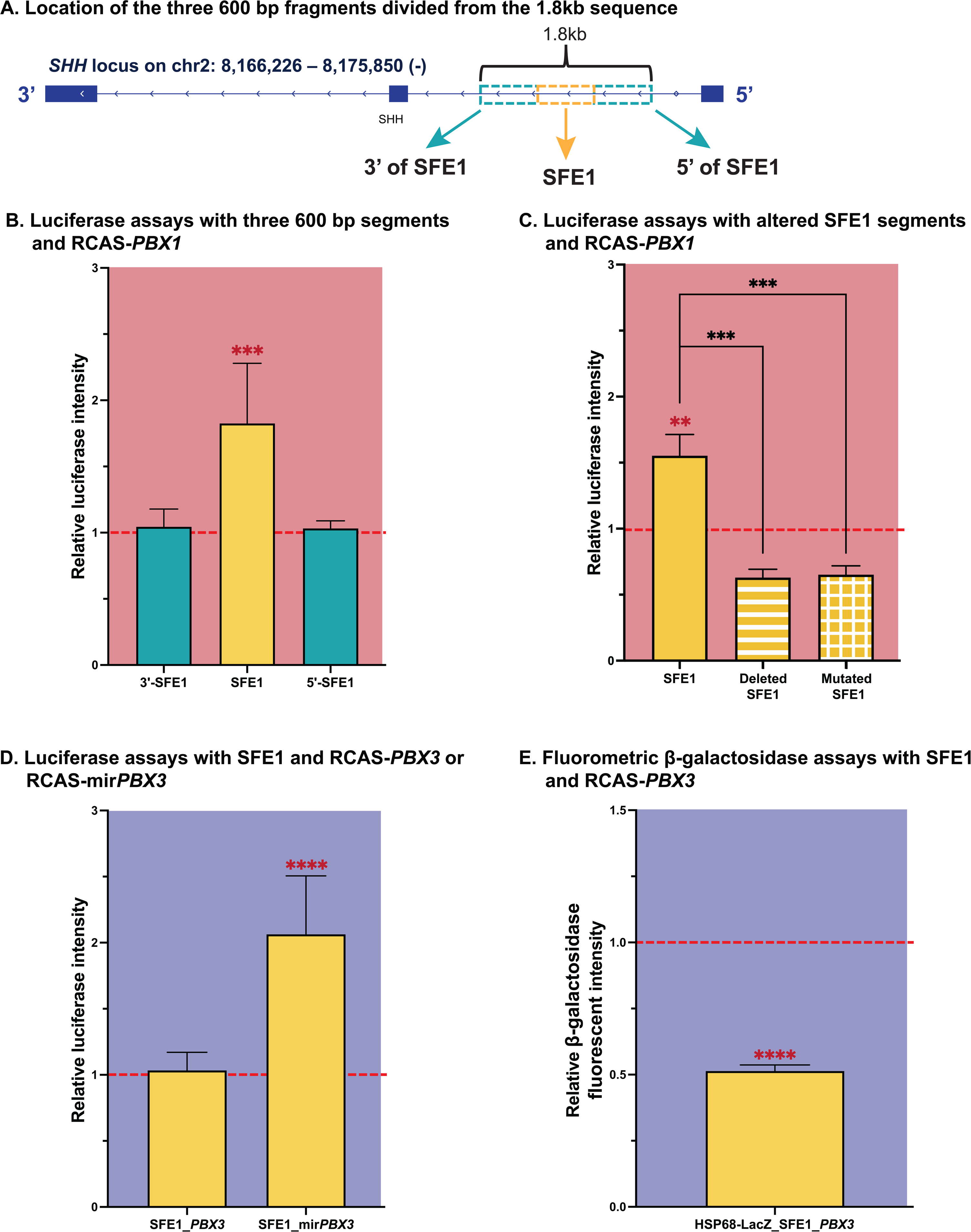

## Supporting information

**S1 Fig. TUNEL (terminal deoxynucleotidyl transferase dUTP nick end labeling) staining, 24 hours after RCAS-miRNA virus infection.** The TUNEL assay demonstrated that RCAS-miRNA treated embryos did not experience significant increase in apoptosis in the FEZ compared to normal embryos.

**S1 Table. Sequencing results of ATAC-seq and ChIP-seq data.**

**S2 Table. Full list of known motif discovery from ATAC-seq data.**

**S3 Table. Full list of known motif discovery from ChIP-seq data targeting PBX1.**

**S4 Table. Full list of *de novo* motif discovery from ChIP-seq data targeting PBX1.**

**S5 Table. Full list of known motif discovery from ChIP-seq data targeting PBX3.**

**S6 Table. Full list of *de novo* motif discovery from ChIP-seq data targeting PBX3.**

## Notes

### Competing Interest Statement

The authors have declared no competing interest.

