## Supporting information for "PBX1 and PBX3 transcription factors regulate *SHH* expression in the Frontonasal Ectodermal Zone through complementary mechanisms"

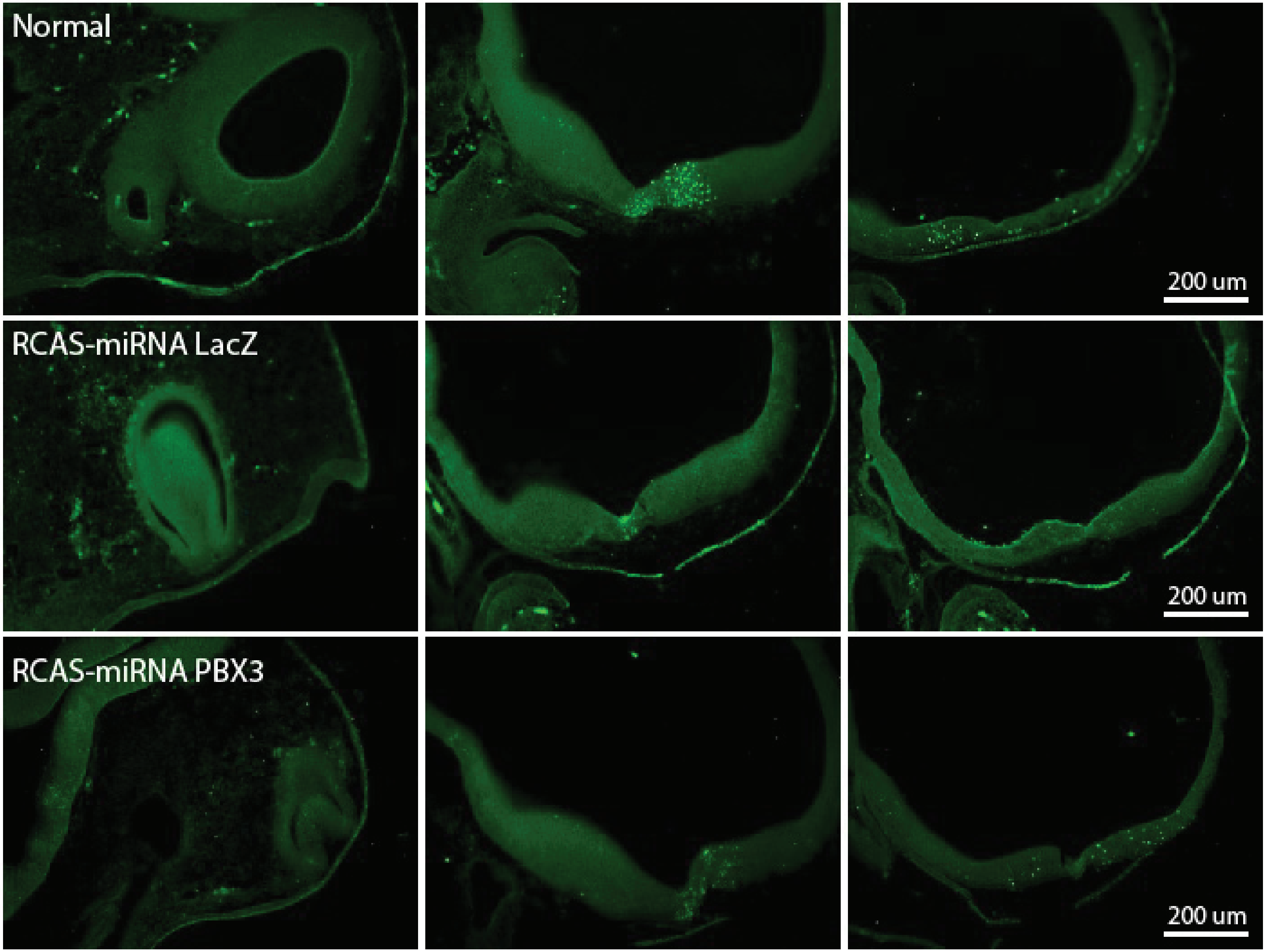

**S1 Fig. TUNEL (terminal deoxynucleotidyl transferase dUTP nick end labeling) staining, 24 hours after RCAS-miRNA virus infection.** The TUNEL assay demonstrated that RCAS-miRNA treated embryos did not experience significant increase in apoptosis in the FEZ compared to normal embryos.

**S1 Table. Sequencing results of ATAC-seq and ChIP-seq data**

| Sequencing type | Sample | Mapping rate, % | Sequencing depth, reads | Peaks of reads | Simple overlapping peaks | Overlapping peaks (IDR<0.05) |
|---|---|---|---|---|---|---|
| ATAC-seq | Replicate 1 | 95.50 | 585,502,859 | 246,299 | 188,247 | 65,556 |
| | Replicate 2 | 96.19 | 707,110,287 | 286,526 | | |
| ChIP-seq for PBX1 | Replicate 1 | 88.56 | 30,851,196 | 44,892 | 32,849 | 13,430 |
| | Input of Replicate 1 | 94.20 | 29,674,869 | | | |
| | Replicate 2 | 91.13 | 36,440,529 | 37,647 | | |
| | Input of Replicate 2 | 93.25 | 40,986,372 | | | |
| ChIP-seq for PBX3 | Replicate 1 | 93.42 | 80,040,505 | 138,442 | 86,512 | 36,122 |
| | Input of Replicate 1 | 97.75 | 90,585,798 | | | |
| | Replicate 2 | 95.29 | 25,146,384 | 99,061 | | |
| | Input of Replicate 2 | 98.18 | 46,876,410 | | | |

Samples were generated from the chick FEZ at HH22 (n=2).

Sequencing was conducted by HiSeq 4000 (Illumina).

Paired-end reads (2×150 nt) were sequenced for ATAC-seq data, and single-end reads (50 nt) were sequenced for ChIP-seq data.

S2 Table. Full list of known motif discovery from ATAC-seq data.

### Homer Known Motif Enrichment Results (/wynton/group/marcucio/2022CHM/Data2022Mar/Motif/HomerATACgiven)

Homer *de novo* Motif Results
Gene Ontology Enrichment Results
Known Motif Enrichment Results (txt file)
Total Target Sequences = 65546, Total Background Sequences = 65439

| Rank | Motif | Name | P-value | log P-pvalue | q-value (Benjamini) | # Target Sequences with Motif | % of Targets Sequenc with Motif |
|---|---|---|---|---|---|---|---|
| 1 | 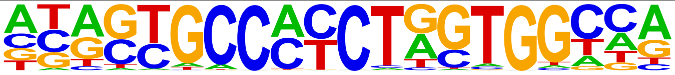 | CTCF(Zf)/CD4+-CTCF-ChIP-Seq(Barski_et_al.)/Homer | 1e-1698 | -3.910e+03 | 0.0000 | 6181.0 | 9.43% |
| 2 | 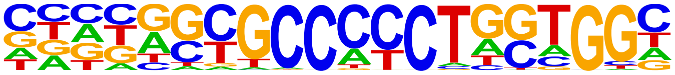 | BORIS(Zf)/K562-CTCFL-ChIP-Seq(GSE32465)/Homer | 1e-1162 | -2.676e+03 | 0.0000 | 7937.0 | 12.11% |
| 3 | 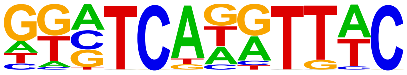 | Six1(Homeobox)/Myoblast-Six1-ChIP-Chip(GSE20150)/Homer | 1e-1020 | -2.349e+03 | 0.0000 | 6792.0 | 10.36% |
| 4 | 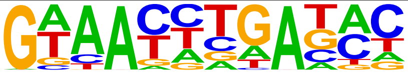 | Six2(Homeobox)/NephronProgenitor-Six2-ChIP-Seq(GSE39837)/Homer | 1e-926 | -2.132e+03 | 0.0000 | 18107.0 | 27.62% |
| 5 | 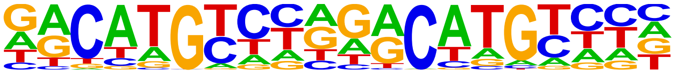 | p53(p53)/Saos-p53-ChIP-Seq(GSE15780)/Homer | 1e-681 | -1.568e+03 | 0.0000 | 3107.0 | 4.74% |
| 6 | 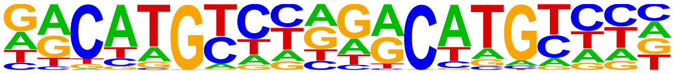 | p53(p53)/Saos-p53-ChIP-Seq/Homer | 1e-681 | -1.568e+03 | 0.0000 | 3107.0 | 4.74% |
| 7 | 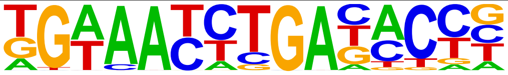 | Six4(Homeobox)/MCF7-SIX4-ChIP-Seq(Encode)/Homer | 1e-649 | -1.497e+03 | 0.0000 | 2040.0 | 3.11% |
| 8 | 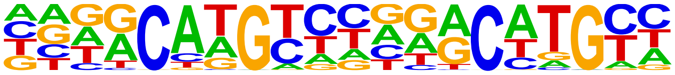 | p63(p53)/Keratinocyte-p63-ChIP-Seq(GSE17611)/Homer | 1e-647 | -1.491e+03 | 0.0000 | 8171.0 | 12.46% |
| 9 | 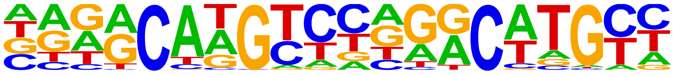 | p73(p53)/Trachea-p73-ChIP-Seq(PRJNA310161)/Homer | 1e-572 | -1.319e+03 | 0.0000 | 2037.0 | 3.11% |
| 10 | 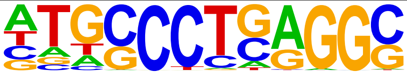 | AP-2alpha(AP2)/Hela-AP2alpha-ChIP-Seq(GSE31477)/Homer | 1e-520 | -1.198e+03 | 0.0000 | 16736.0 | 25.53% |
| 11 | 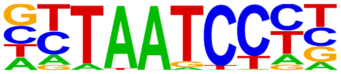 | Otx2(Homeobox)/EpiLC-Otx2-ChIP-Seq(GSE56098)/Homer | 1e-450 | -1.037e+03 | 0.0000 | 13393.0 | 20.43% |
| 12 | 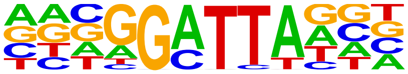 | bcd(Homeobox)/Embryo-Bcd-ChIP-Seq(GSE86966)/Homer | 1e-426 | -9.817e+02 | 0.0000 | 18093.0 | 27.60% |
| 13 | 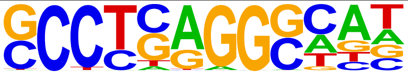 | AP-2gamma(AP2)/MCF7-TFAP2C-ChIP-Seq(GSE21234)/Homer | 1e-425 | -9.805e+02 | 0.0000 | 20164.0 | 30.76% |
| 14 | 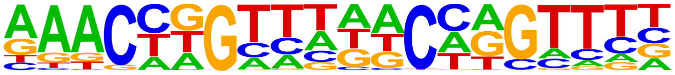 | GRHL2(CP2)/HBE-GRHL2-ChIP-Seq(GSE46194)/Homer | 1e-400 | -9.223e+02 | 0.0000 | 7540.0 | 11.50% |
| 15 | 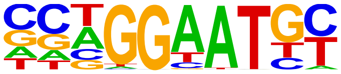 | TEAD4(TEA)/Tropoblast-Tead4-ChIP-Seq(GSE37350)/Homer | 1e-360 | -8.300e+02 | 0.0000 | 13905.0 | 21.21% |
| 16 | 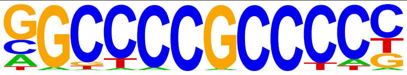 | Sp1(Zf)/Promoter/Homer | 1e-359 | -8.281e+02 | 0.0000 | 6284.0 | 9.59% |
| 17 | 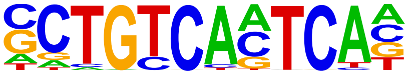 | Pbx3(Homeobox)/GM12878-PBX3-ChIP-Seq(GSE32465)/Homer | 1e-343 | -7.901e+02 | 0.0000 | 5100.0 | 7.78% |
| 18 | 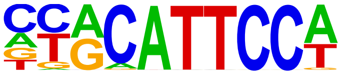 | TEAD1(TEAD)/HepG2-TEAD1-ChIP-Seq(Encode)/Homer | 1e-331 | -7.631e+02 | 0.0000 | 14993.0 | 22.87% |
| 19 | 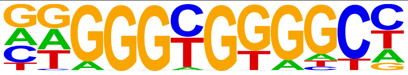 | KLF1(Zf)/HUDEP2-KLF1-CutnRun(GSE136251)/Homer | 1e-325 | -7.486e+02 | 0.0000 | 11796.0 | 17.99% |
| 20 | 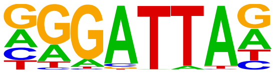 | GSC(Homeobox)/FrogEmbryos-GSC-ChIP-Seq(DRA000576)/Homer | 1e-319 | -7.357e+02 | 0.0000 | 17430.0 | 26.59% |

| | | | | | | | |
|---|---|---|---|---|---|---|---|
| 21 | 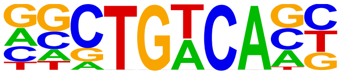 | Meis1(Homeobox)/MastCells-Meis1-ChIP-Seq(GSE48085)/Homer | 1e-309 | -7.116e+02 | 0.0000 | 28924.0 | 44.12% |
| 22 | 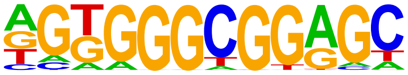 | Sp5(Zf)/mES-Sp5.Flag-ChIP-Seq(GSE72989)/Homer | 1e-307 | -7.071e+02 | 0.0000 | 14866.0 | 22.68% |
| 23 | 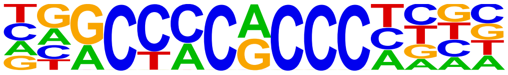 | KLF3(Zf)/MEF-Klf3-ChIP-Seq(GSE44748)/Homer | 1e-296 | -6.822e+02 | 0.0000 | 6815.0 | 10.40% |
| 24 | 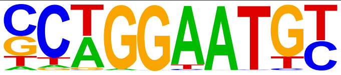 | TEAD2(TEA)/Py2T-Tead2-ChIP-Seq(GSE55709)/Homer | 1e-293 | -6.769e+02 | 0.0000 | 8930.0 | 13.62% |
| 25 | 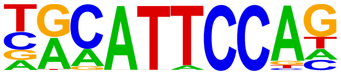 | TEAD3(TEA)/HepG2-TEAD3-ChIP-Seq(Encode)/Homer | 1e-280 | -6.465e+02 | 0.0000 | 17088.0 | 26.07% |
| 26 | 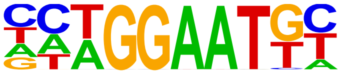 | TEAD(TEA)/Fibroblast-PU.1-ChIP-Seq(Unpublished)/Homer | 1e-266 | -6.147e+02 | 0.0000 | 10858.0 | 16.56% |
| 27 | 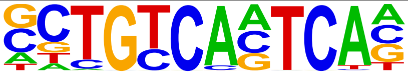 | Pknox1(Homeobox)/ES-Prep1-ChIP-Seq(GSE63282)/Homer | 1e-260 | -6.002e+02 | 0.0000 | 4840.0 | 7.38% |
| 28 | 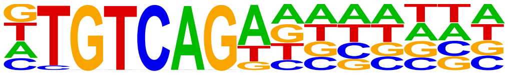 | AtGRF6(GRF)/col-AtGRF6-DAP-Seq(GSE60143)/Homer | 1e-256 | -5.902e+02 | 0.0000 | 21327.0 | 32.53% |
| 29 | 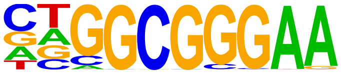 | E2F1(E2F)/Hela-E2F1-ChIP-Seq(GSE22478)/Homer | 1e-251 | -5.788e+02 | 0.0000 | 4978.0 | 7.59% |
| 30 | 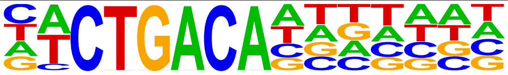 | GRF9(GRF)/colamp-GRF9-DAP-Seq(GSE60143)/Homer | 1e-248 | -5.731e+02 | 0.0000 | 18346.0 | 27.99% |
| 31 | 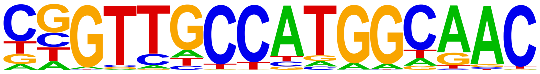 | RFX(HTH)/K562-RFX3-ChIP-Seq(SRA012198)/Homer | 1e-244 | -5.620e+02 | 0.0000 | 2160.0 | 3.29% |
| 32 | 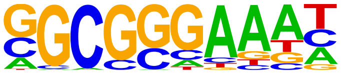 | E2F4(E2F)/K562-E2F4-ChIP-Seq(GSE31477)/Homer | 1e-228 | -5.269e+02 | 0.0000 | 7658.0 | 11.68% |
| 33 | 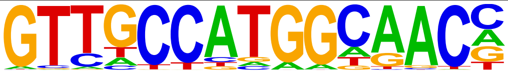 | Rfx2(HTH)/LoVo-RFX2-ChIP-Seq(GSE49402)/Homer | 1e-214 | -4.939e+02 | 0.0000 | 2291.0 | 3.49% |
| 34 | 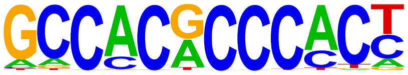 | Klf9(Zf)/GBM-Klf9-ChIP-Seq(GSE62211)/Homer | 1e-202 | -4.661e+02 | 0.0000 | 4982.0 | 7.60% |
| 35 | 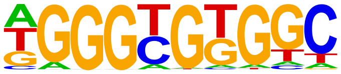 | KLF5(Zf)/LoVo-KLF5-ChIP-Seq(GSE49402)/Homer | 1e-200 | -4.615e+02 | 0.0000 | 16131.0 | 24.61% |
| 36 | 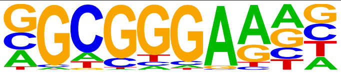 | E2F6(E2F)/Hela-E2F6-ChIP-Seq(GSE31477)/Homer | 1e-196 | -4.515e+02 | 0.0000 | 9873.0 | 15.06% |
| 37 | 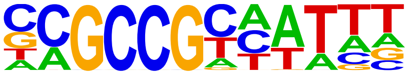 | RRTF1(AP2EREBP)/colamp-RRTF1-DAP-Seq(GSE60143)/Homer | 1e-188 | -4.351e+02 | 0.0000 | 5225.0 | 7.97% |
| 38 | 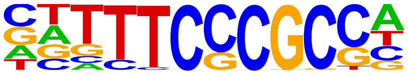 | E2F7(E2F)/Hela-E2F7-ChIP-Seq(GSE32673)/Homer | 1e-187 | -4.315e+02 | 0.0000 | 2379.0 | 3.63% |
| 39 | 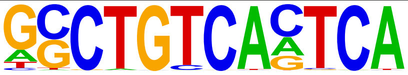 | PBX1(Homeobox)/MCF7-PBX1-ChIP-Seq(GSE28007)/Homer | 1e-185 | -4.269e+02 | 0.0000 | 1800.0 | 2.75% |
| 40 | 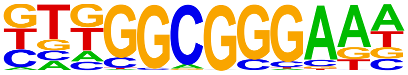 | E2F3(E2F)/MEF-E2F3-ChIP-Seq(GSE71376)/Homer | 1e-179 | -4.130e+02 | 0.0000 | 10719.0 | 16.35% |
| 41 | 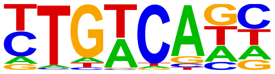 | Tgif1(Homeobox)/mES-Tgif1-ChIP-Seq(GSE55404)/Homer | 1e-160 | -3.687e+02 | 0.0000 | 40421.0 | 61.66% |
| 42 | 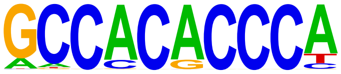 | Klf4(Zf)/mES-Klf4-ChIP-Seq(GSE11431)/Homer | 1e-152 | -3.515e+02 | 0.0000 | 3886.0 | 5.93% |
| 43 | 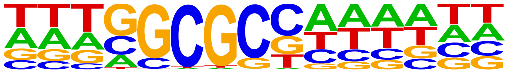 | E2FA(E2FDP)/colamp-E2FA-DAP-Seq(GSE60143)/Homer | 1e-152 | -3.502e+02 | 0.0000 | 4833.0 | 7.37% |
| 44 | 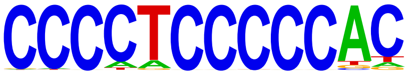 | Zfp281(Zf)/ES-Zfp281-ChIP-Seq(GSE81042)/Homer | 1e-146 | -3.381e+02 | 0.0000 | 3370.0 | 5.14% |
| 45 | 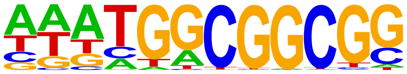 | ERF9(AP2EREBP)/colamp-ERF9-DAP-Seq(GSE60143)/Homer | 1e-137 | -3.174e+02 | 0.0000 | 7959.0 | 12.14% |

| | | | | | | | |
|---|---|---|---|---|---|---|---|
| 46 | 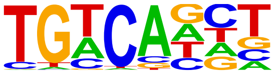 | Tgif2(Homeobox)/mES-Tgif2-ChIP-Seq(GSE55404)/Homer | 1e-134 | -3.106e+02 | 0.0000 | 42535.0 | 64.88% |
| 47 | 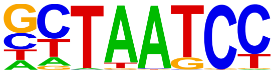 | CRX(Homeobox)/Retina-Crx-ChIP-Seq(GSE20012)/Homer | 1e-132 | -3.043e+02 | 0.0000 | 29266.0 | 44.64% |
| 48 | 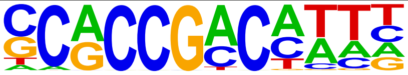 | At5g65130(AP2EREBP)/colamp-At5g65130-DAP-Seq(GSE60143)/Homer | 1e-127 | -2.939e+02 | 0.0000 | 5130.0 | 7.83% |
| 49 | 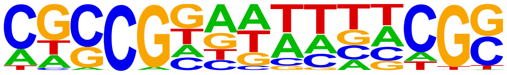 | LOB(LOBAS2)/col-LOB-DAP-Seq(GSE60143)/Homer | 1e-127 | -2.930e+02 | 0.0000 | 5221.0 | 7.96% |
| 50 | 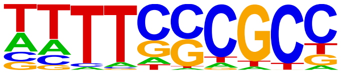 | DEL2(E2FDP)/col-DEL2-DAP-Seq(GSE60143)/Homer | 1e-118 | -2.732e+02 | 0.0000 | 5603.0 | 8.55% |
| 51 | 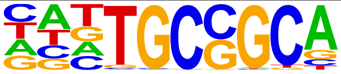 | Zfp57(Zf)/H1-ZFP57.HA-ChIP-Seq(GSE115387)/Homer | 1e-111 | -2.558e+02 | 0.0000 | 9810.0 | 14.96% |
| 52 | 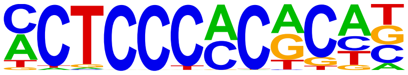 | WT1(Zf)/Kidney-WT1-ChIP-Seq(GSE90016)/Homer | 1e-109 | -2.525e+02 | 0.0000 | 10217.0 | 15.59% |
| 53 | 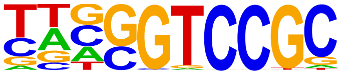 | HINFP(Zf)/K562-HINFP.eGFP-ChIP-Seq(Encode)/Homer | 1e-108 | -2.490e+02 | 0.0000 | 5830.0 | 8.89% |
| 54 | 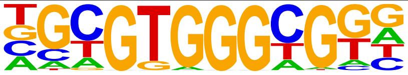 | Egr2(Zf)/Thymocytes-Egr2-ChIP-Seq(GSE34254)/Homer | 1e-107 | -2.476e+02 | 0.0000 | 3312.0 | 5.05% |
| 55 | 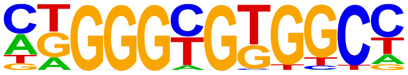 | KLF6(Zf)/PDAC-KLF6-ChIP-Seq(GSE64557)/Homer | 1e-97 | -2.253e+02 | 0.0000 | 14737.0 | 22.48% |
| 56 | 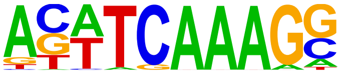 | Tcf3(HMG)/mES-Tcf3-ChIP-Seq(GSE11724)/Homer | 1e-97 | -2.247e+02 | 0.0000 | 4561.0 | 6.96% |
| 57 | 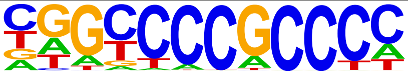 | Sp2(Zf)/HEK293-Sp2.eGFP-ChIP-Seq(Encode)/Homer | 1e-96 | -2.218e+02 | 0.0000 | 19513.0 | 29.77% |
| 58 | 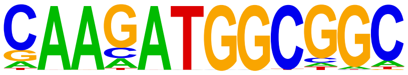 | YY1(Zf)/Promoter/Homer | 1e-96 | -2.216e+02 | 0.0000 | 1814.0 | 2.77% |
| 59 | 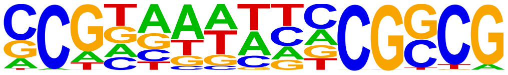 | AS2(LOBAS2)/col-AS2-DAP-Seq(GSE60143)/Homer | 1e-95 | -2.188e+02 | 0.0000 | 2387.0 | 3.64% |
| 60 | 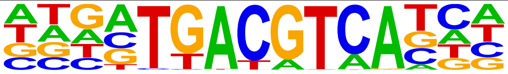 | TGA5(bZIP)/col-TGA5-DAP-Seq(GSE60143)/Homer | 1e-91 | -2.116e+02 | 0.0000 | 1631.0 | 2.49% |
| 61 | 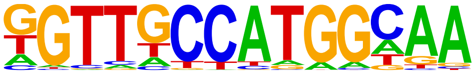 | X-box(HTH)/NPC-H3K4me1-ChIP-Seq(GSE16256)/Homer | 1e-91 | -2.116e+02 | 0.0000 | 2083.0 | 3.18% |
| 62 | 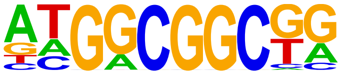 | AT4G18450(AP2EREBP)/col-AT4G18450-DAP-Seq(GSE60143)/Homer | 1e-88 | -2.037e+02 | 0.0000 | 9805.0 | 14.96% |
| 63 | 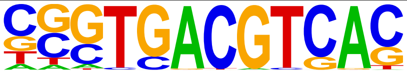 | CRE(bZIP)/Promoter/Homer | 1e-87 | -2.017e+02 | 0.0000 | 4204.0 | 6.41% |
| 64 | 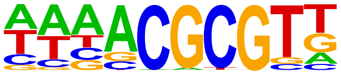 | CAMTA1(CAMTA)/col-CAMTA1-DAP-Seq(GSE60143)/Homer | 1e-85 | -1.958e+02 | 0.0000 | 8099.0 | 12.35% |
| 65 | 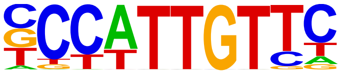 | Sox2(HMG)/mES-Sox2-ChIP-Seq(GSE11431)/Homer | 1e-83 | -1.918e+02 | 0.0000 | 13016.0 | 19.85% |
| 66 | 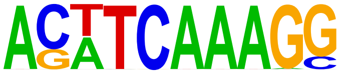 | TCFL2(HMG)/K562-TCF7L2-ChIP-Seq(GSE29196)/Homer | 1e-81 | -1.878e+02 | 0.0000 | 1673.0 | 2.55% |
| 67 | 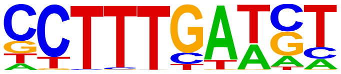 | LEF1(HMG)/H1-LEF1-ChIP-Seq(GSE64758)/Homer | 1e-80 | -1.862e+02 | 0.0000 | 10647.0 | 16.24% |
| 68 | 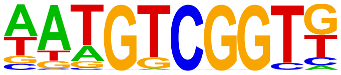 | DREB19(AP2EREBP)/colamp-DREB19-DAP-Seq(GSE60143)/Homer | 1e-80 | -1.849e+02 | 0.0000 | 7275.0 | 11.10% |
| 69 | 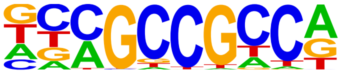 | ERF5(AP2EREBP)/colamp-ERF5-DAP-Seq(GSE60143)/Homer | 1e-80 | -1.845e+02 | 0.0000 | 10184.0 | 15.53% |
| 70 | 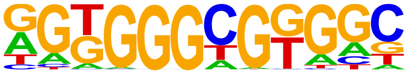 | KLF14(Zf)/HEK293-KLF14.GFP-ChIP-Seq(GSE58341)/Homer | 1e-79 | -1.841e+02 | 0.0000 | 21932.0 | 33.46% |

| | | | | | | | |
|---|---|---|---|---|---|---|---|
| 71 | 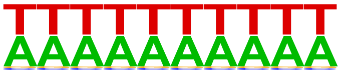 | SeqBias: A/T bias | 1e-79 | -1.829e+02 | 0.0000 | 57774.0 | 88.13% |
| 72 | 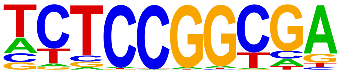 | AT3G58630(Trihelix)/col-AT3G58630-DAP-Seq(GSE60143)/Homer | 1e-79 | -1.820e+02 | 0.0000 | 2755.0 | 4.20% |
| 73 | 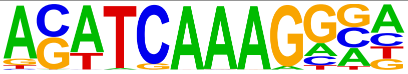 | Tcf4(HMG)/Hct116-Tcf4-ChIP-Seq(SRA012054)/Homer | 1e-78 | -1.802e+02 | 0.0000 | 7523.0 | 11.48% |
| 74 | 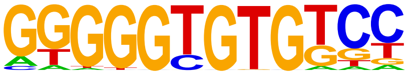 | KLF10(Zf)/HEK293-KLF10.GFP-ChIP-Seq(GSE58341)/Homer | 1e-77 | -1.783e+02 | 0.0000 | 6890.0 | 10.51% |
| 75 | 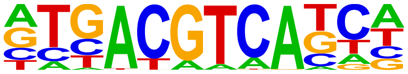 | TGA4(bZIP)/colamp-TGA4-DAP-Seq(GSE60143)/Homer | 1e-74 | -1.707e+02 | 0.0000 | 4433.0 | 6.76% |
| 76 | 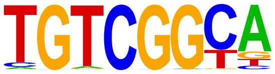 | AT1G12630(AP2EREBP)/colamp-AT1G12630-DAP-Seq(GSE60143)/Homer | 1e-73 | -1.693e+02 | 0.0000 | 6241.0 | 9.52% |
| 77 | 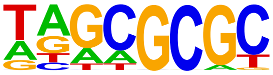 | DPL-1(E2F)/cElegans-Adult-ChIP-Seq(modEncode)/Homer | 1e-73 | -1.687e+02 | 0.0000 | 11143.0 | 17.00% |
| 78 | 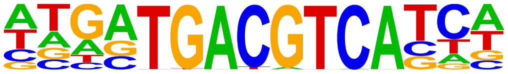 | TGA3(bZIP)/colamp-TGA3-DAP-Seq(GSE60143)/Homer | 1e-73 | -1.684e+02 | 0.0000 | 1050.0 | 1.60% |
| 79 | 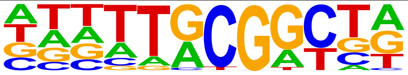 | At5g08750(C3H)/col-At5g08750-DAP-Seq(GSE60143)/Homer | 1e-73 | -1.681e+02 | 0.0000 | 6796.0 | 10.37% |
| 80 | 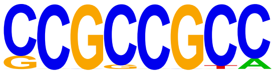 | SHN3(AP2EREBP)/col-SHN3-DAP-Seq(GSE60143)/Homer | 1e-71 | -1.648e+02 | 0.0000 | 9682.0 | 14.77% |
| 81 | 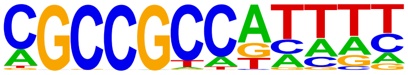 | RAP212(AP2EREBP)/col-RAP212-DAP-Seq(GSE60143)/Homer | 1e-69 | -1.607e+02 | 0.0000 | 12162.0 | 18.55% |
| 82 | 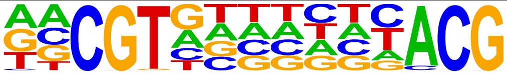 | ANAC094(NAC)/col-ANAC094-DAP-Seq(GSE60143)/Homer | 1e-67 | -1.553e+02 | 0.0000 | 6768.0 | 10.32% |
| 83 | 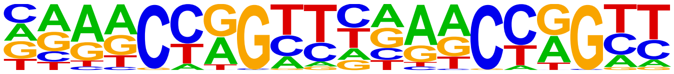 | Tcfcp2l1(CP2)/mES-Tcfcp2l1-ChIP-Seq(GSE11431)/Homer | 1e-66 | -1.521e+02 | 0.0000 | 2238.0 | 3.41% |
| 84 | 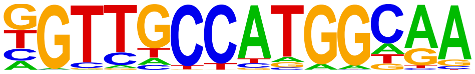 | Rfx1(HTH)/NPC-H3K4me1-ChIP-Seq(GSE16256)/Homer | 1e-65 | -1.513e+02 | 0.0000 | 3518.0 | 5.37% |
| 85 | 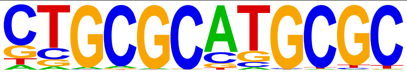 | NRF1(NRF)/MCF7-NRF1-ChIP-Seq(Unpublished)/Homer | 1e-65 | -1.505e+02 | 0.0000 | 2523.0 | 3.85% |
| 86 | 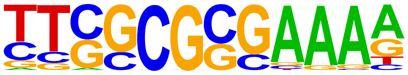 | E2F(E2F)/Hela-CellCycle-Expression/Homer | 1e-63 | -1.471e+02 | 0.0000 | 823.0 | 1.26% |
| 87 | 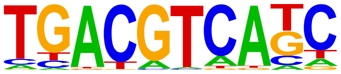 | TGA1(bZIP)/colamp-TGA1-DAP-Seq(GSE60143)/Homer | 1e-62 | -1.435e+02 | 0.0000 | 5846.0 | 8.92% |
| 88 | 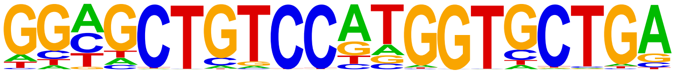 | REST-NRSF(Zf)/Jurkat-NRSF-ChIP-Seq/Homer | 1e-60 | -1.400e+02 | 0.0000 | 299.0 | 0.46% |
| 89 | 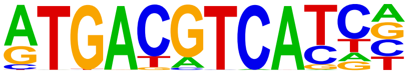 | JunD(bZIP)/K562-JunD-ChIP-Seq/Homer | 1e-60 | -1.385e+02 | 0.0000 | 1478.0 | 2.25% |
| 90 | 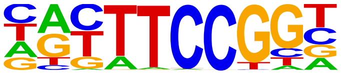 | Elk4(ETS)/Hela-Elk4-ChIP-Seq(GSE31477)/Homer | 1e-60 | -1.382e+02 | 0.0000 | 6826.0 | 10.41% |
| 91 | 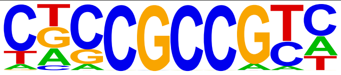 | LEP(AP2EREBP)/col-LEP-DAP-Seq(GSE60143)/Homer | 1e-59 | -1.371e+02 | 0.0000 | 9299.0 | 14.18% |
| 92 | 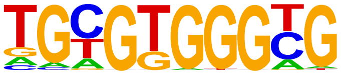 | Egr1(Zf)/K562-Egr1-ChIP-Seq(GSE32465)/Homer | 1e-59 | -1.369e+02 | 0.0000 | 10300.0 | 15.71% |
| 93 | 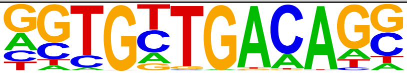 | Tbx20(T-box)/Heart-Tbx20-ChIP-Seq(GSE29636)/Homer | 1e-57 | -1.332e+02 | 0.0000 | 5048.0 | 7.70% |
| 94 | 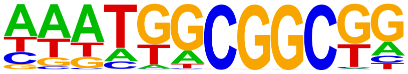 | ABR1(AP2EREBP)/colamp-ABR1-DAP-Seq(GSE60143)/Homer | 1e-57 | -1.325e+02 | 0.0000 | 14302.0 | 21.82% |
| 95 | 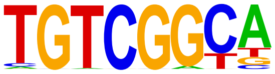 | Rap210(AP2EREBP)/col-Rap210-DAP-Seq(GSE60143)/Homer | 1e-56 | -1.302e+02 | 0.0000 | 6646.0 | 10.14% |

| | | | | | | | |
|---|---|---|---|---|---|---|---|
| 96 | 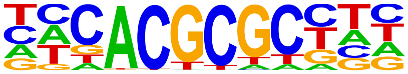 | FHY3(FAR1)/Arabidopsis-FHY3-ChIP-Seq(GSE30711)/Homer | 1e-56 | -1.294e+02 | 0.0000 | 5633.0 | 8.59% |
| 97 | 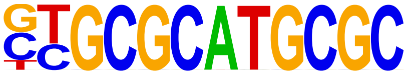 | NRF(NRF)/Promoter/Homer | 1e-55 | -1.281e+02 | 0.0000 | 2826.0 | 4.31% |
| 98 | 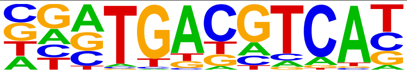 | Atf2(bZIP)/3T3L1-Atf2-ChIP-Seq(GSE56872)/Homer | 1e-55 | -1.272e+02 | 0.0000 | 4635.0 | 7.07% |
| 99 | 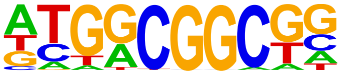 | ERF3(AP2EREBP)/colamp-ERF3-DAP-Seq(GSE60143)/Homer | 1e-55 | -1.267e+02 | 0.0000 | 11992.0 | 18.29% |
| 100 | 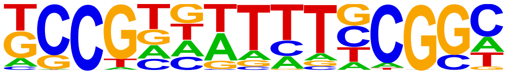 | LBD13(LOBAS2)/colamp-LBD13-DAP-Seq(GSE60143)/Homer | 1e-54 | -1.260e+02 | 0.0000 | 12439.0 | 18.97% |
| 101 | 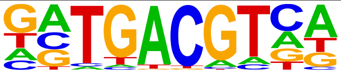 | bZIP50(bZIP)/colamp-bZIP50-DAP-Seq(GSE60143)/Homer | 1e-54 | -1.256e+02 | 0.0000 | 13339.0 | 20.35% |
| 102 | 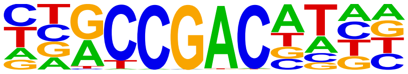 | CBF3(AP2EREBP)/colamp-CBF3-DAP-Seq(GSE60143)/Homer | 1e-54 | -1.246e+02 | 0.0000 | 4860.0 | 7.41% |
| 103 | 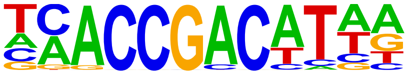 | DREB2(AP2EREBP)/col-DREB2-DAP-Seq(GSE60143)/Homer | 1e-54 | -1.244e+02 | 0.0000 | 5507.0 | 8.40% |
| 104 | 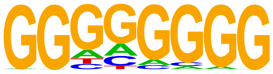 | Maz(Zf)/HepG2-Maz-ChIP-Seq(GSE31477)/Homer | 1e-53 | -1.239e+02 | 0.0000 | 19125.0 | 29.17% |
| 105 | 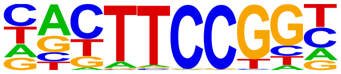 | Elk1(ETS)/Hela-Elk1-ChIP-Seq(GSE31477)/Homer | 1e-52 | -1.219e+02 | 0.0000 | 6697.0 | 10.22% |
| 106 | 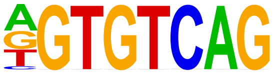 | CELF2(RRM)/JSL1-CELF2-CLIP-Seq(GSE71264)/Homer | 1e-52 | -1.210e+02 | 0.0000 | 6494.0 | 9.91% |
| 107 | 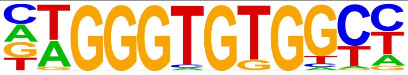 | EKLF(Zf)/Erythrocyte-Klf1-ChIP-Seq(GSE20478)/Homer | 1e-52 | -1.207e+02 | 0.0000 | 1634.0 | 2.49% |
| 108 | 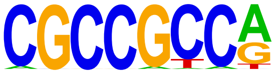 | CRF4(AP2EREBP)/colamp-CRF4-DAP-Seq(GSE60143)/Homer | 1e-51 | -1.178e+02 | 0.0000 | 11038.0 | 16.84% |
| 109 | 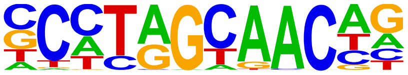 | Rfx5(HTH)/GM12878-Rfx5-ChIP-Seq(GSE31477)/Homer | 1e-49 | -1.144e+02 | 0.0000 | 4860.0 | 7.41% |
| 110 | 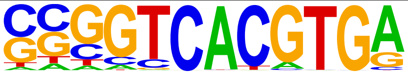 | E-box(bHLH)/Promoter/Homer | 1e-49 | -1.131e+02 | 0.0000 | 1314.0 | 2.00% |
| 111 | 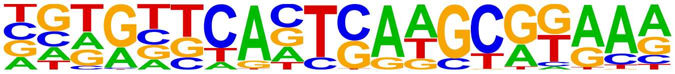 | PAX6(Paired,Homeobox)/Forebrain-Pax6-ChIP-Seq(GSE66961)/Homer | 1e-47 | -1.096e+02 | 0.0000 | 1731.0 | 2.64% |
| 112 | 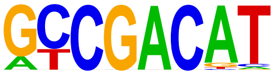 | DDF1(AP2EREBP)/col-DDF1-DAP-Seq(GSE60143)/Homer | 1e-47 | -1.089e+02 | 0.0000 | 4199.0 | 6.41% |
| 113 | 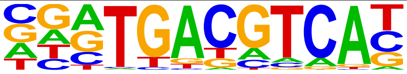 | Atf7(bZIP)/3T3L1-Atf7-ChIP-Seq(GSE56872)/Homer | 1e-47 | -1.087e+02 | 0.0000 | 6674.0 | 10.18% |
| 114 | 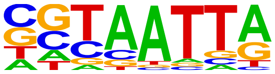 | DLX5(Homeobox)/BasalGanglia-Dlx5-ChIP-seq(GSE124936)/Homer | 1e-44 | -1.033e+02 | 0.0000 | 13172.0 | 20.09% |
| 115 | 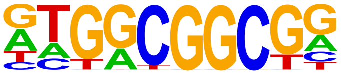 | ERF10(AP2EREBP)/col-ERF10-DAP-Seq(GSE60143)/Homer | 1e-44 | -1.032e+02 | 0.0000 | 11969.0 | 18.26% |
| 116 | 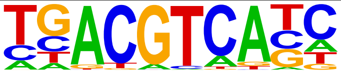 | TGA6(bZIP)/colamp-TGA6-DAP-Seq(GSE60143)/Homer | 1e-44 | -1.018e+02 | 0.0000 | 8335.0 | 12.71% |
| 117 | 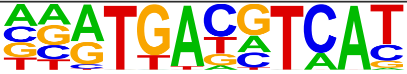 | CREB5(bZIP)/LNCaP-CREB5.V5-ChIP-Seq(GSE137775)/Homer | 1e-44 | -1.015e+02 | 0.0000 | 5333.0 | 8.14% |
| 118 | 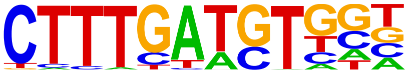 | Tcf7(HMG)/GM12878-TCF7-ChIP-Seq(Encode)/Homer | 1e-42 | -9.812e+01 | 0.0000 | 5568.0 | 8.49% |
| 119 | 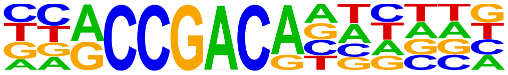 | AT3G60490(AP2EREBP)/colamp-AT3G60490-DAP-Seq(GSE60143)/Homer | 1e-42 | -9.770e+01 | 0.0000 | 3581.0 | 5.46% |
| 120 | | ERF1(AP2EREBP)/colamp-ERF1-DAP-Seq(GSE60143)/Homer | 1e-42 | -9.675e+01 | 0.0000 | 11945.0 | 18.22% |

| | | | | | | | |
|---|---|---|---|---|---|---|---|
| | 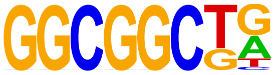 | | | | | | |
| 121 | 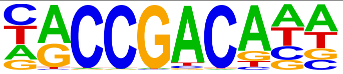 | At4g16750(AP2EREBP)/col-At4g16750-DAP-Seq(GSE60143)/Homer | 1e-41 | -9.449e+01 | 0.0000 | 7962.0 | 12.15% |
| 122 | 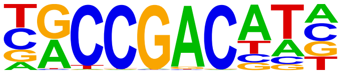 | CBF1(AP2EREBP)/colamp-CBF1-DAP-Seq(GSE60143)/Homer | 1e-40 | -9.317e+01 | 0.0000 | 7058.0 | 10.77% |
| 123 | 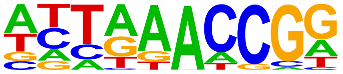 | Unknown3/Arabidopsis-Promoters/Homer | 1e-40 | -9.257e+01 | 0.0000 | 2139.0 | 3.26% |
| 124 | 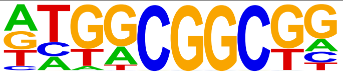 | ERF11(AP2EREBP)/col-ERF11-DAP-Seq(GSE60143)/Homer | 1e-39 | -9.163e+01 | 0.0000 | 13574.0 | 20.71% |
| 125 | 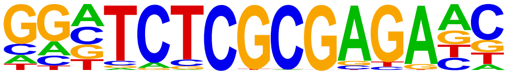 | ZBTB33(Zf)/GM12878-ZBTB33-ChIP-Seq(GSE32465)/Homer | 1e-39 | -8.993e+01 | 0.0000 | 767.0 | 1.17% |
| 126 | 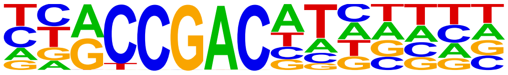 | CBF2(AP2EREBP)/colamp-CBF2-DAP-Seq(GSE60143)/Homer | 1e-38 | -8.969e+01 | 0.0000 | 4822.0 | 7.36% |
| 127 | 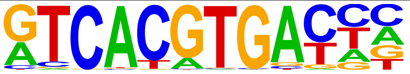 | TFE3(bHLH)/MEF-TFE3-ChIP-Seq(GSE75757)/Homer | 1e-38 | -8.954e+01 | 0.0000 | 1018.0 | 1.55% |
| 128 | 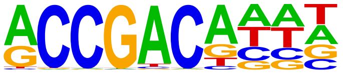 | AT1G77200(AP2EREBP)/colamp-AT1G77200-DAP-Seq(GSE60143)/Homer | 1e-38 | -8.932e+01 | 0.0000 | 7957.0 | 12.14% |
| 129 | 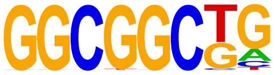 | ERF2(AP2EREBP)/colamp-ERF2-DAP-Seq(GSE60143)/Homer | 1e-38 | -8.773e+01 | 0.0000 | 12660.0 | 19.31% |
| 130 | 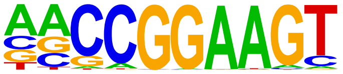 | ETS(ETS)/Promoter/Homer | 1e-37 | -8.710e+01 | 0.0000 | 3569.0 | 5.44% |
| 131 | 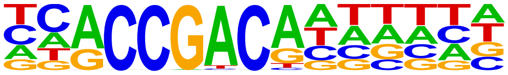 | AT3G16280(AP2EREBP)/colamp-AT3G16280-DAP-Seq(GSE60143)/Homer | 1e-37 | -8.578e+01 | 0.0000 | 4029.0 | 6.15% |
| 132 | 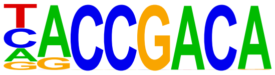 | bHLH10(bHLH)/colamp-bHLH10-DAP-Seq(GSE60143)/Homer | 1e-36 | -8.323e+01 | 0.0000 | 3571.0 | 5.45% |
| 133 | 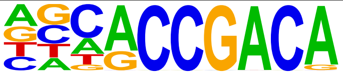 | TINY(AP2EREBP)/col-TINY-DAP-Seq(GSE60143)/Homer | 1e-35 | -8.116e+01 | 0.0000 | 3326.0 | 5.07% |
| 134 | 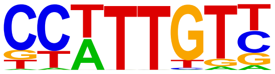 | Sox3(HMG)/NPC-Sox3-ChIP-Seq(GSE33059)/Homer | 1e-34 | -8.029e+01 | 0.0000 | 23264.0 | 35.49% |
| 135 | 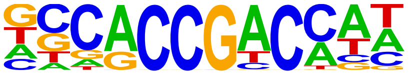 | At4g28140(AP2EREBP)/colamp-At4g28140-DAP-Seq(GSE60143)/Homer | 1e-34 | -7.967e+01 | 0.0000 | 4459.0 | 6.80% |
| 136 | 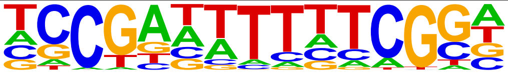 | LBD2(LOBAS2)/colamp-LBD2-DAP-Seq(GSE60143)/Homer | 1e-34 | -7.937e+01 | 0.0000 | 3435.0 | 5.24% |
| 137 | 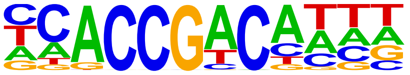 | DEAR2(AP2EREBP)/colamp-DEAR2-DAP-Seq(GSE60143)/Homer | 1e-34 | -7.928e+01 | 0.0000 | 12190.0 | 18.59% |
| 138 | 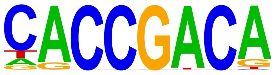 | ERF38(AP2EREBP)/col-ERF38-DAP-Seq(GSE60143)/Homer | 1e-34 | -7.925e+01 | 0.0000 | 5828.0 | 8.89% |
| 139 | 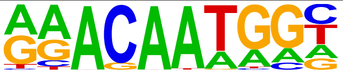 | Sox15(HMG)/CPA-Sox15-ChIP-Seq(GSE62909)/Homer | 1e-34 | -7.902e+01 | 0.0000 | 14121.0 | 21.54% |
| 140 | 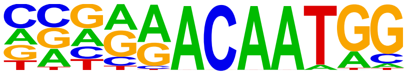 | Sox7(HMG)/ESC-Sox7-ChIP-Seq(GSE133899)/Homer | 1e-33 | -7.737e+01 | 0.0000 | 4009.0 | 6.12% |
| 141 | 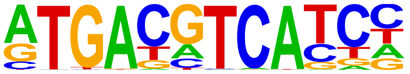 | c-Jun-CRE(bZIP)/K562-cJun-ChIP-Seq(GSE31477)/Homer | 1e-33 | -7.729e+01 | 0.0000 | 4250.0 | 6.48% |
| 142 | 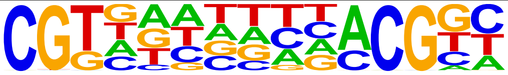 | ANAC042(NAC)/col-ANAC042-DAP-Seq(GSE60143)/Homer | 1e-33 | -7.723e+01 | 0.0000 | 11711.0 | 17.86% |
| 143 | 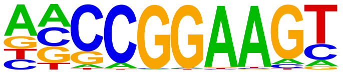 | ELF1(ETS)/Jurkat-ELF1-ChIP-Seq(SRA014231)/Homer | 1e-33 | -7.630e+01 | 0.0000 | 5784.0 | 8.82% |
| 144 | 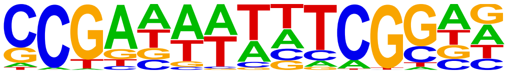 | CDM1(C3H)/colamp-CDM1-DAP-Seq(GSE60143)/Homer | 1e-33 | -7.617e+01 | 0.0000 | 1112.0 | 1.70% |

| | | | | | | | |
|---|---|---|---|---|---|---|---|
| 145 | 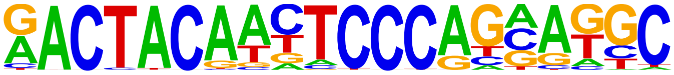 | Ronin(THAP)/ES-Thap11-ChIP-Seq(GSE51522)/Homer | 1e-32 | -7.564e+01 | 0.0000 | 569.0 | 0.87% |
| 146 | 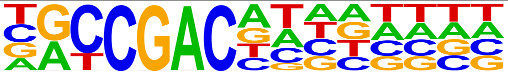 | CBF4(AP2EREBP)/colamp-CBF4-DAP-Seq(GSE60143)/Homer | 1e-32 | -7.511e+01 | 0.0000 | 6800.0 | 10.37% |
| 147 | 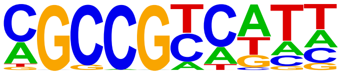 | ERF8(AP2EREBP)/colamp-ERF8-DAP-Seq(GSE60143)/Homer | 1e-32 | -7.460e+01 | 0.0000 | 14675.0 | 22.39% |
| 148 | 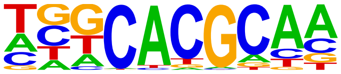 | Arnt:Ahr(bHLH)/MCF7-Arnt-ChIP-Seq(Lo_et_al.)/Homer | 1e-32 | -7.425e+01 | 0.0000 | 12813.0 | 19.55% |
| 149 | 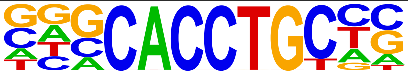 | Slug(Zf)/Mesoderm-Snai2-ChIP-Seq(GSE61475)/Homer | 1e-31 | -7.302e+01 | 0.0000 | 9711.0 | 14.81% |
| 150 | 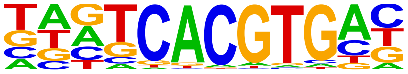 | BIM3(bHLH)/col-BIM3-DAP-Seq(GSE60143)/Homer | 1e-31 | -7.209e+01 | 0.0000 | 2038.0 | 3.11% |
| 151 | 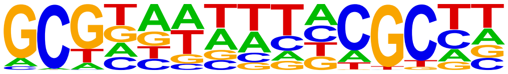 | LBD23(LOBAS2)/colamp-LBD23-DAP-Seq(GSE60143)/Homer | 1e-31 | -7.179e+01 | 0.0000 | 15795.0 | 24.09% |
| 152 | 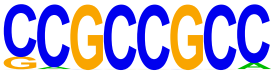 | ERF73(AP2EREBP)/col-ERF73-DAP-Seq(GSE60143)/Homer | 1e-30 | -7.135e+01 | 0.0000 | 12933.0 | 19.73% |
| 153 | 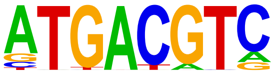 | TGA10(bZIP)/colamp-TGA10-DAP-Seq(GSE60143)/Homer | 1e-30 | -7.032e+01 | 0.0000 | 7958.0 | 12.14% |
| 154 | 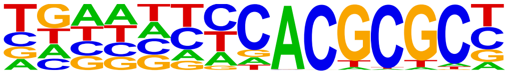 | FAR1(FAR1)/col-FAR1-DAP-Seq(GSE60143)/Homer | 1e-30 | -6.982e+01 | 0.0000 | 2323.0 | 3.54% |
| 155 | 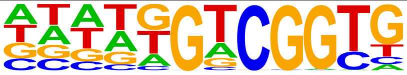 | At1g36060(AP2EREBP)/colamp-At1g36060-DAP-Seq(GSE60143)/Homer | 1e-29 | -6.741e+01 | 0.0000 | 8136.0 | 12.41% |
| 156 | 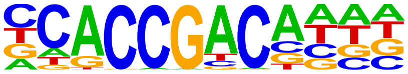 | At1g22810(AP2EREBP)/colamp-At1g22810-DAP-Seq(GSE60143)/Homer | 1e-28 | -6.646e+01 | 0.0000 | 5627.0 | 8.58% |
| 157 | 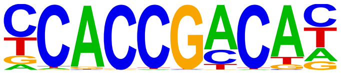 | DREB26(AP2EREBP)/col-DREB26-DAP-Seq(GSE60143)/Homer | 1e-28 | -6.639e+01 | 0.0000 | 4115.0 | 6.28% |
| 158 | 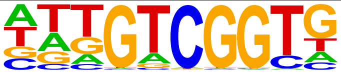 | CEJ1(AP2EREBP)/col-CEJ1-DAP-Seq(GSE60143)/Homer | 1e-28 | -6.629e+01 | 0.0000 | 10751.0 | 16.40% |
| 159 | 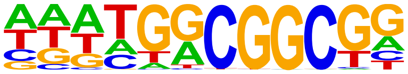 | ERF4(AP2EREBP)/colamp-ERF4-DAP-Seq(GSE60143)/Homer | 1e-27 | -6.431e+01 | 0.0000 | 14765.0 | 22.52% |
| 160 | 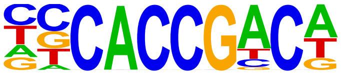 | AT1G44830(AP2EREBP)/col-AT1G44830-DAP-Seq(GSE60143)/Homer | 1e-27 | -6.356e+01 | 0.0000 | 5258.0 | 8.02% |
| 161 | 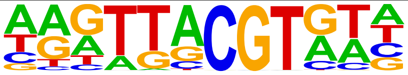 | NAP(NAC)/col-NAP-DAP-Seq(GSE60143)/Homer | 1e-27 | -6.338e+01 | 0.0000 | 11852.0 | 18.08% |
| 162 | 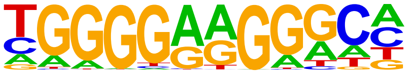 | ZNF467(Zf)/HEK293-ZNF467.GFP-ChIP-Seq(GSE58341)/Homer | 1e-27 | -6.270e+01 | 0.0000 | 14572.0 | 22.23% |
| 163 | 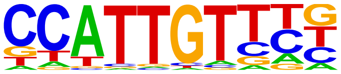 | Sox17(HMG)/Endoderm-Sox17-ChIP-Seq(GSE61475)/Homer | 1e-27 | -6.229e+01 | 0.0000 | 9345.0 | 14.25% |
| 164 | 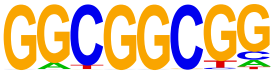 | ESE1(AP2EREBP)/col-ESE1-DAP-Seq(GSE60143)/Homer | 1e-26 | -6.141e+01 | 0.0000 | 13116.0 | 20.01% |
| 165 | 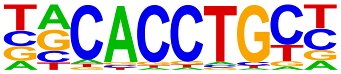 | Snail1(Zf)/LS174T-SNAIL1.HA-ChIP-Seq(GSE127183)/Homer | 1e-26 | -6.075e+01 | 0.0000 | 14168.0 | 21.61% |
| 166 | 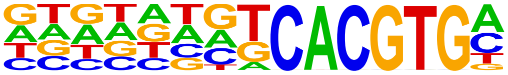 | BIM1(bHLH)/colamp-BIM1-DAP-Seq(GSE60143)/Homer | 1e-25 | -5.970e+01 | 0.0000 | 1895.0 | 2.89% |
| 167 | 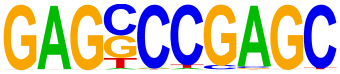 | ZNF519(Zf)/HEK293-ZNF519.GFP-ChIP-Seq(GSE58341)/Homer | 1e-25 | -5.896e+01 | 0.0000 | 4176.0 | 6.37% |
| 168 | 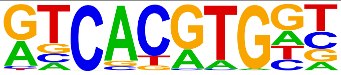 | Usf2(bHLH)/C2C12-Usf2-ChIP-Seq(GSE36030)/Homer | 1e-25 | -5.846e+01 | 0.0000 | 4277.0 | 6.52% |
| 169 | 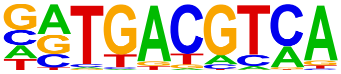 | Atf1(bZIP)/K562-ATF1-ChIP-Seq(GSE31477)/Homer | 1e-25 | -5.846e+01 | 0.0000 | 9145.0 | 13.95% |

| | | | | | | | |
|---|---|---|---|---|---|---|---|
| 170 | 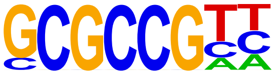 | PUCHI(AP2EREBP)/colamp-PUCHI-DAP-Seq(GSE60143)/Homer | 1e-25 | -5.815e+01 | 0.0000 | 12642.0 | 19.28% |
| 171 | 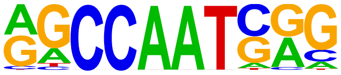 | NFY(CCAAT)/Promoter/Homer | 1e-24 | -5.665e+01 | 0.0000 | 9423.0 | 14.37% |
| 172 | 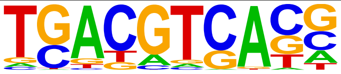 | FEA4(bZIP)/Corn-FEA4-ChIP-Seq(GSE61954)/Homer | 1e-24 | -5.533e+01 | 0.0000 | 15186.0 | 23.16% |
| 173 | 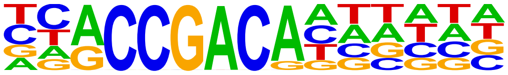 | At2g44940(AP2EREBP)/colamp-At2g44940-DAP-Seq(GSE60143)/Homer | 1e-24 | -5.531e+01 | 0.0000 | 2500.0 | 3.81% |
| 174 | 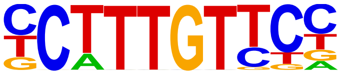 | Sox4(HMG)/proB-Sox4-ChIP-Seq(GSE50066)/Homer | 1e-23 | -5.476e+01 | 0.0000 | 12684.0 | 19.35% |
| 175 | 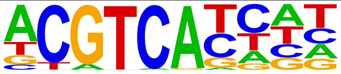 | TGA2(bZIP)/colamp-TGA2-DAP-Seq(GSE60143)/Homer | 1e-23 | -5.410e+01 | 0.0000 | 7610.0 | 11.61% |
| 176 | 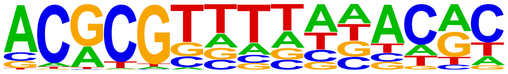 | CAMTA5(CAMTA)/col-CAMTA5-DAP-Seq(GSE60143)/Homer | 1e-23 | -5.396e+01 | 0.0000 | 4414.0 | 6.73% |
| 177 | 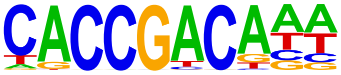 | At4g31060(AP2EREBP)/colamp-At4g31060-DAP-Seq(GSE60143)/Homer | 1e-23 | -5.338e+01 | 0.0000 | 4783.0 | 7.30% |
| 178 | 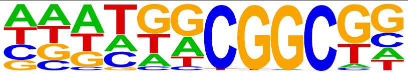 | RAP26(AP2EREBP)/colamp-RAP26-DAP-Seq(GSE60143)/Homer | 1e-23 | -5.317e+01 | 0.0000 | 16001.0 | 24.41% |
| 179 | 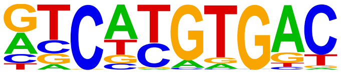 | MITF(bHLH)/MastCells-MITF-ChIP-Seq(GSE48085)/Homer | 1e-22 | -5.194e+01 | 0.0000 | 11075.0 | 16.89% |
| 180 | 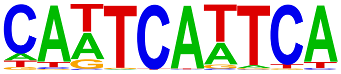 | WUS1(Homeobox)/colamp-WUS1-DAP-Seq(GSE60143)/Homer | 1e-21 | -4.895e+01 | 0.0000 | 5556.0 | 8.48% |
| 181 | 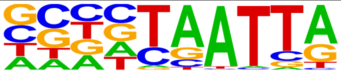 | DLX1(Homeobox)/BasalGanglia-Dlx1-ChIP-seq(GSE124936)/Homer | 1e-21 | -4.860e+01 | 0.0000 | 19445.0 | 29.66% |
| 182 | 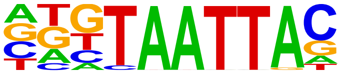 | Dlx3(Homeobox)/Kerainocytes-Dlx3-ChIP-Seq(GSE89884)/Homer | 1e-20 | -4.774e+01 | 0.0000 | 11342.0 | 17.30% |
| 183 | 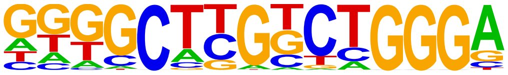 | Zfp809(Zf)/ES-Zfp809-ChIP-Seq(GSE70799)/Homer | 1e-20 | -4.724e+01 | 0.0000 | 4059.0 | 6.19% |
| 184 | 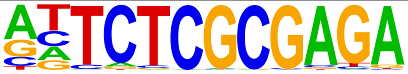 | GFX(?)/Promoter/Homer | 1e-19 | -4.565e+01 | 0.0000 | 263.0 | 0.40% |
| 185 | 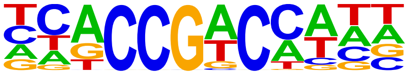 | At1g19210(AP2EREBP)/colamp-At1g19210-DAP-Seq(GSE60143)/Homer | 1e-19 | -4.456e+01 | 0.0000 | 14017.0 | 21.38% |
| 186 | 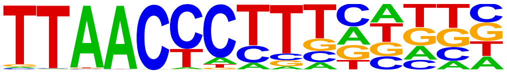 | ZNF652/HepG2-ZNF652.Flag-ChIP-Seq(Encode)/Homer | 1e-18 | -4.365e+01 | 0.0000 | 3438.0 | 5.24% |
| 187 | 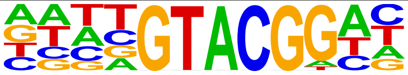 | bHLH28(bHLH)/col-bHLH28-DAP-Seq(GSE60143)/Homer | 1e-18 | -4.158e+01 | 0.0000 | 4565.0 | 6.96% |
| 188 | 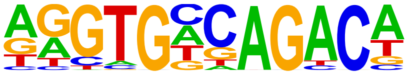 | Tbox:Smad(T-box,MAD)/ESCd5-Smad2_3-ChIP-Seq(GSE29422)/Homer | 1e-17 | -4.112e+01 | 0.0000 | 2489.0 | 3.80% |
| 189 | 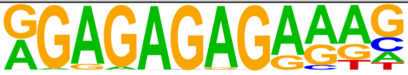 | FRS9(ND)/col-FRS9-DAP-Seq(GSE60143)/Homer | 1e-17 | -4.088e+01 | 0.0000 | 3382.0 | 5.16% |
| 190 | 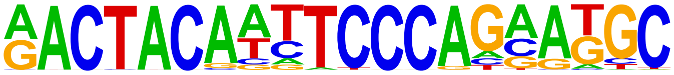 | GFY-Staf(?,Zf)/Promoter/Homer | 1e-17 | -4.066e+01 | 0.0000 | 766.0 | 1.17% |
| 191 | 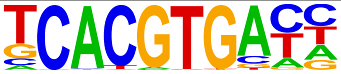 | Cbf1(bHLH)/Yeast-Cbf1-ChIP-Seq(GSE29506)/Homer | 1e-17 | -4.009e+01 | 0.0000 | 4254.0 | 6.49% |
| 192 | 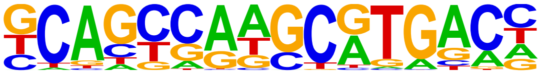 | PAX5(Paired,Homeobox)/GM12878-PAX5-ChIP-Seq(GSE32465)/Homer | 1e-17 | -3.950e+01 | 0.0000 | 6786.0 | 10.35% |
| 193 | 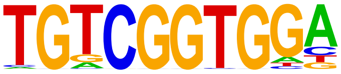 | At1g77640(AP2EREBP)/col-At1g77640-DAP-Seq(GSE60143)/Homer | 1e-16 | -3.861e+01 | 0.0000 | 2799.0 | 4.27% |
| 194 | 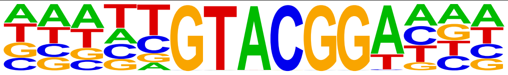 | SPL14(SBP)/col-SPL14-DAP-Seq(GSE60143)/Homer | 1e-16 | -3.810e+01 | 0.0000 | 2731.0 | 4.17% |

| | | | | | | | |
|---|---|---|---|---|---|---|---|
| 195 | 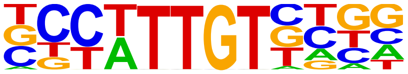 | Sox21(HMG)/ESC-SOX21-ChIP-Seq(GSE110505)/Homer | 1e-16 | -3.746e+01 | 0.0000 | 24065.0 | 36.71% |
| 196 | 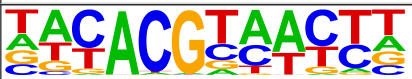 | ANAC047(NAC)/colamp-ANAC047-DAP-Seq(GSE60143)/Homer | 1e-15 | -3.642e+01 | 0.0000 | 9314.0 | 14.21% |
| 197 | 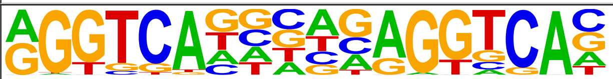 | RAR:RXR(NR),DR5/ES-RAR-ChIP-Seq(GSE56893)/Homer | 1e-15 | -3.462e+01 | 0.0000 | 399.0 | 0.61% |
| 198 | 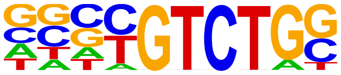 | Smad4(MAD)/ESC-SMAD4-ChIP-Seq(GSE29422)/Homer | 1e-14 | -3.443e+01 | 0.0000 | 21865.0 | 33.35% |
| 199 | 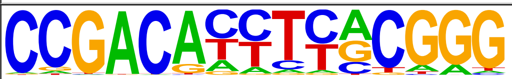 | GEI-11(Myb?)/cElegans-L4-GEI11-ChIP-Seq(modEncode)/Homer | 1e-14 | -3.422e+01 | 0.0000 | 1000.0 | 1.53% |
| 200 | 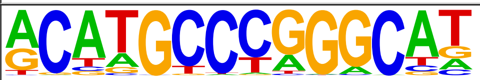 | p53(p53)/mES-cMyc-ChIP-Seq(GSE11431)/Homer | 1e-14 | -3.385e+01 | 0.0000 | 275.0 | 0.42% |
| 201 | 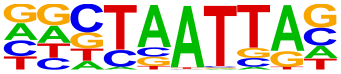 | En1(Homeobox)/SUM149-EN1-ChIP-Seq(GSE120957)/Homer | 1e-14 | -3.355e+01 | 0.0000 | 24552.0 | 37.45% |
| 202 | 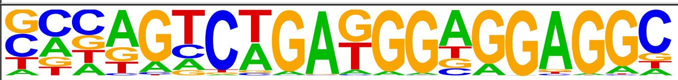 | ZSCAN22(Zf)/HEK293-ZSCAN22.GFP-ChIP-Seq(GSE58341)/Homer | 1e-14 | -3.281e+01 | 0.0000 | 1056.0 | 1.61% |
| 203 | 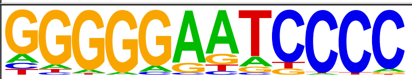 | NFkB-p50,p52(RHD)/Monocyte-p50-ChIP-Chip(Schreiber_et_al.)/Homer | 1e-13 | -3.149e+01 | 0.0000 | 2345.0 | 3.58% |
| 204 | 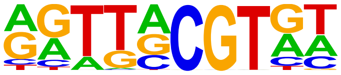 | NAM(NAC)/col-NAM-DAP-Seq(GSE60143)/Homer | 1e-13 | -3.126e+01 | 0.0000 | 15540.0 | 23.70% |
| 205 | 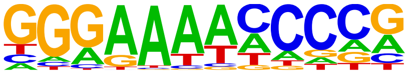 | Dorsal(RHD)/Embryo-dl-ChIP-Seq(GSE65441)/Homer | 1e-13 | -3.036e+01 | 0.0000 | 2860.0 | 4.36% |
| 206 | 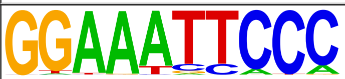 | NFkB-p65-Rel(RHD)/ThioMac-LPS-Expression(GSE23622)/Homer | 1e-11 | -2.745e+01 | 0.0000 | 867.0 | 1.32% |
| 207 | 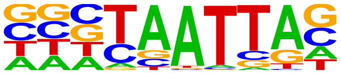 | DLX2(Homeobox)/BasalGanglia-Dlx2-ChIP-seq(GSE124936)/Homer | 1e-11 | -2.719e+01 | 0.0000 | 20933.0 | 31.93% |
| 208 | 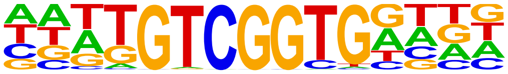 | DEAR5(AP2EREBP)/col-DEAR5-DAP-Seq(GSE60143)/Homer | 1e-11 | -2.617e+01 | 0.0000 | 2095.0 | 3.20% |
| 209 | 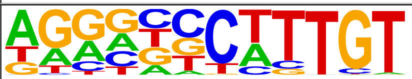 | Sox9(HMG)/Limb-SOX9-ChIP-Seq(GSE73225)/Homer | 1e-11 | -2.595e+01 | 0.0000 | 11873.0 | 18.11% |
| 210 | 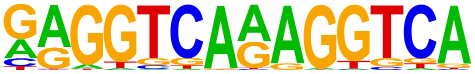 | TR4(NR),DR1/Hela-TR4-ChIP-Seq(GSE24685)/Homer | 1e-11 | -2.560e+01 | 0.0000 | 971.0 | 1.48% |
| 211 | 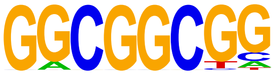 | ERF104(AP2EREBP)/col-ERF104-DAP-Seq(GSE60143)/Homer | 1e-10 | -2.442e+01 | 0.0000 | 14638.0 | 22.33% |
| 212 | 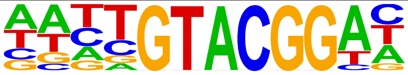 | SPL13(SBP)/col-SPL13-DAP-Seq(GSE60143)/Homer | 1e-10 | -2.392e+01 | 0.0000 | 1545.0 | 2.36% |
| 213 | 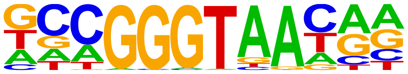 | REB1/SacCer-Promoters/Homer | 1e-10 | -2.365e+01 | 0.0000 | 2351.0 | 3.59% |
| 214 | 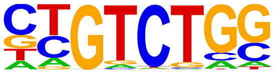 | Smad2(MAD)/ES-SMAD2-ChIP-Seq(GSE29422)/Homer | 1e-9 | -2.281e+01 | 0.0000 | 21520.0 | 32.83% |
| 215 | 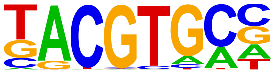 | HIF-1a(bHLH)/MCF7-HIF1a-ChIP-Seq(GSE28352)/Homer | 1e-9 | -2.237e+01 | 0.0000 | 5840.0 | 8.91% |
| 216 | 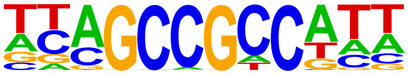 | ERF13(AP2EREBP)/colamp-ERF13-DAP-Seq(GSE60143)/Homer | 1e-9 | -2.209e+01 | 0.0000 | 17044.0 | 26.00% |
| 217 | 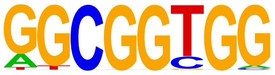 | AT3G57600(AP2EREBP)/col-AT3G57600-DAP-Seq(GSE60143)/Homer | 1e-9 | -2.138e+01 | 0.0000 | 13820.0 | 21.08% |
| 218 | 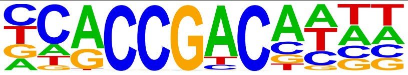 | RAP21(AP2EREBP)/colamp-RAP21-DAP-Seq(GSE60143)/Homer | 1e-9 | -2.088e+01 | 0.0000 | 2064.0 | 3.15% |
| 219 | 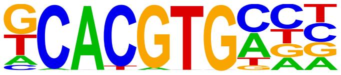 | bHLHE40(bHLH)/HepG2-BHLHE40-ChIP-Seq(GSE31477)/Homer | 1e-9 | -2.084e+01 | 0.0000 | 5538.0 | 8.45% |

| | | | | | | | |
|---|---|---|---|---|---|---|---|
| 220 | 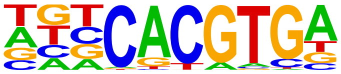 | SPCH(bHLH)/Seedling-SPCH-ChIP-Seq(GSE57497)/Homer | 1e-8 | -2.061e+01 | 0.0000 | 15134.0 | 23.09% |
| 221 | 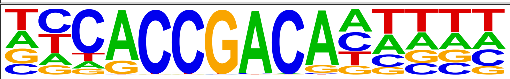 | At4g32800(AP2EREBP)/colamp-At4g32800-DAP-Seq(GSE60143)/Homer | 1e-8 | -2.008e+01 | 0.0000 | 1339.0 | 2.04% |
| 222 | 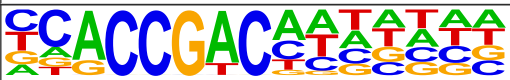 | DEAR3(AP2EREBP)/colamp-DEAR3-DAP-Seq(GSE60143)/Homer | 1e-8 | -2.004e+01 | 0.0000 | 3476.0 | 5.30% |
| 223 | 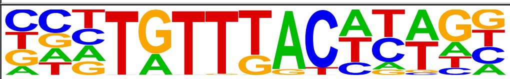 | Foxa3(Forkhead)/Liver-Foxa3-ChIP-Seq(GSE77670)/Homer | 1e-8 | -1.995e+01 | 0.0000 | 4315.0 | 6.58% |
| 224 | 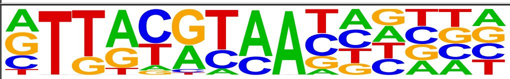 | NFIL3(bZIP)/HepG2-NFIL3-ChIP-Seq(Encode)/Homer | 1e-8 | -1.945e+01 | 0.0000 | 8842.0 | 13.49% |
| 225 | 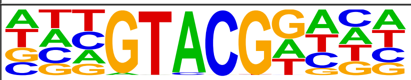 | SPL5(SBP)/colamp-SPL5-DAP-Seq(GSE60143)/Homer | 1e-8 | -1.885e+01 | 0.0000 | 7148.0 | 10.90% |
| 226 | 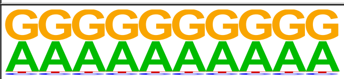 | SeqBias: G/A bias | 1e-8 | -1.855e+01 | 0.0000 | 65524.0 | 99.95% |
| 227 | 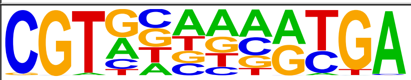 | ABF1/SacCer-Promoters/Homer | 1e-8 | -1.848e+01 | 0.0000 | 3266.0 | 4.98% |
| 228 | 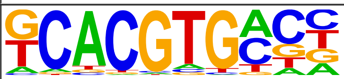 | bHLHE41(bHLH)/proB-Bhlhe41-ChIP-Seq(GSE93764)/Homer | 1e-8 | -1.847e+01 | 0.0000 | 17735.0 | 27.05% |
| 229 | 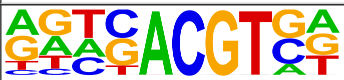 | HY5(bZIP)/colamp-HY5-DAP-Seq(GSE60143)/Homer | 1e-7 | -1.656e+01 | 0.0000 | 10980.0 | 16.75% |
| 230 | 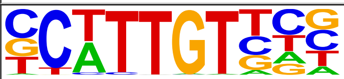 | Sox10(HMG)/SciaticNerve-Sox3-ChIP-Seq(GSE35132)/Homer | 1e-7 | -1.651e+01 | 0.0000 | 22613.0 | 34.49% |
| 231 | 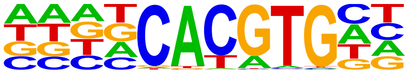 | BIM2(bHLH)/col-BIM2-DAP-Seq(GSE60143)/Homer | 1e-6 | -1.521e+01 | 0.0000 | 12508.0 | 19.08% |
| 232 | 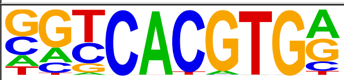 | USF1(bHLH)/GM12878-Usf1-ChIP-Seq(GSE32465)/Homer | 1e-6 | -1.511e+01 | 0.0000 | 6887.0 | 10.51% |
| 233 | 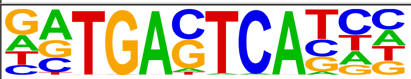 | Jun-AP1(bZIP)/K562-cJun-ChIP-Seq(GSE31477)/Homer | 1e-6 | -1.468e+01 | 0.0000 | 2605.0 | 3.97% |
| 234 | 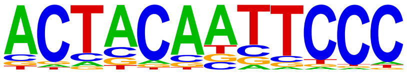 | GFY(?)/Promoter/Homer | 1e-6 | -1.448e+01 | 0.0000 | 953.0 | 1.45% |
| 235 | 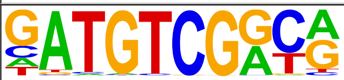 | DDF2(AP2EREBP)/col-DDF2-DAP-Seq(GSE60143)/Homer | 1e-6 | -1.426e+01 | 0.0000 | 426.0 | 0.65% |
| 236 | 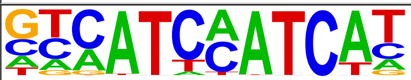 | HOXA2(Homeobox)/mES-Hoxa2-ChIP-Seq(Donaldson_et_al.)/Homer | 1e-6 | -1.419e+01 | 0.0000 | 1383.0 | 2.11% |
| 237 | 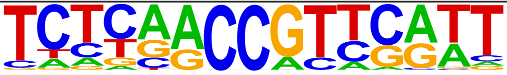 | AT5G59990(C2C2COlike)/colamp-AT5G59990-DAP-Seq(GSE60143)/Homer | 1e-6 | -1.400e+01 | 0.0000 | 251.0 | 0.38% |
| 238 | 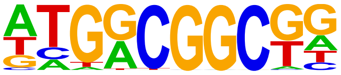 | ERF7(AP2EREBP)/col-ERF7-DAP-Seq(GSE60143)/Homer | 1e-5 | -1.295e+01 | 0.0000 | 16565.0 | 25.27% |
| 239 | 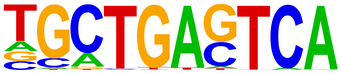 | Bach2(bZIP)/OCILy7-Bach2-ChIP-Seq(GSE44420)/Homer | 1e-5 | -1.291e+01 | 0.0000 | 2380.0 | 3.63% |
| 240 | 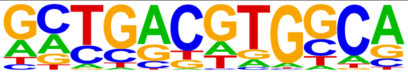 | O2(bZIP)/Corn-O2-ChIP-Seq(GSE63991)/Homer | 1e-5 | -1.284e+01 | 0.0000 | 2330.0 | 3.55% |
| 241 | 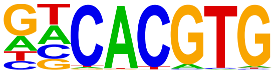 | CLOCK(bHLH)/Liver-Clock-ChIP-Seq(GSE39860)/Homer | 1e-5 | -1.280e+01 | 0.0000 | 8301.0 | 12.66% |
| 242 | 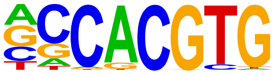 | c-Myc(bHLH)/LNCAP-cMyc-ChIP-Seq(Unpublished)/Homer | 1e-5 | -1.264e+01 | 0.0000 | 9046.0 | 13.80% |
| 243 | 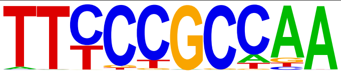 | DEL1(E2FDP)/colamp-DEL1-DAP-Seq(GSE60143)/Homer | 1e-5 | -1.250e+01 | 0.0000 | 48.0 | 0.07% |
| 244 | | Pax8(Paired,Homeobox)/Thyroid-Pax8-ChIP-Seq(GSE26938)/Homer | 1e-5 | -1.198e+01 | 0.0000 | 4890.0 | 7.46% |

| | | | | | | | |
|---|---|---|---|---|---|---|---|
| | 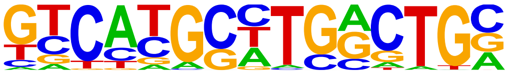 | | | | | | |
| 245 | 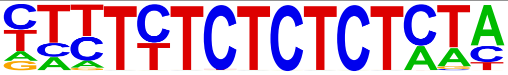 | BPC6(BBRBPC)/col-BPC6-DAP-Seq(GSE60143)/Homer | 1e-5 | -1.193e+01 | 0.0000 | 450.0 | 0.69% |
| 246 | 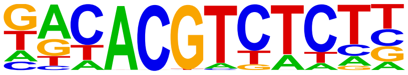 | bHLH157(bHLH)/col-bHLH157-DAP-Seq(GSE60143)/Homer | 1e-5 | -1.164e+01 | 0.0000 | 3131.0 | 4.78% |
| 247 | 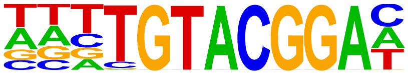 | SPL3(SBP)/colamp-SPL3-DAP-Seq(GSE60143)/Homer | 1e-4 | -1.142e+01 | 0.0000 | 647.0 | 0.99% |
| 248 | 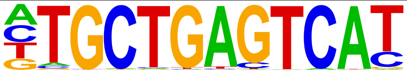 | Nrf2(bZIP)/Lymphoblast-Nrf2-ChIP-Seq(GSE37589)/Homer | 1e-4 | -1.098e+01 | 0.0001 | 662.0 | 1.01% |
| 249 | 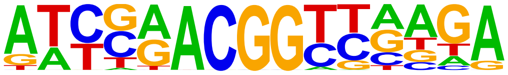 | ZIM(C2C2gata)/col-ZIM-DAP-Seq(GSE60143)/Homer | 1e-4 | -1.080e+01 | 0.0001 | 105.0 | 0.16% |
| 250 | 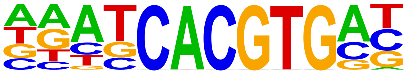 | bHLH74(bHLH)/col-bHLH74-DAP-Seq(GSE60143)/Homer | 1e-4 | -1.075e+01 | 0.0001 | 4271.0 | 6.52% |
| 251 | 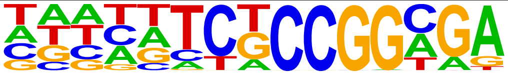 | AT5G05550(Trihelix)/col-AT5G05550-DAP-Seq(GSE60143)/Homer | 1e-4 | -1.048e+01 | 0.0001 | 18080.0 | 27.58% |
| 252 | 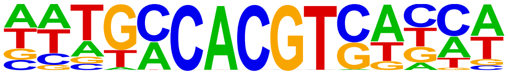 | GBF6(bZIP)/colamp-GBF6-DAP-Seq(GSE60143)/Homer | 1e-4 | -1.044e+01 | 0.0001 | 2917.0 | 4.45% |
| 253 | 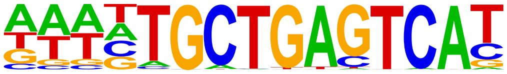 | Bach1(bZIP)/K562-Bach1-ChIP-Seq(GSE31477)/Homer | 1e-4 | -1.042e+01 | 0.0001 | 754.0 | 1.15% |
| 254 | 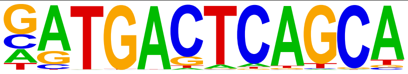 | NF-E2(bZIP)/K562-NFE2-ChIP-Seq(GSE31477)/Homer | 1e-4 | -1.035e+01 | 0.0001 | 773.0 | 1.18% |
| 255 | 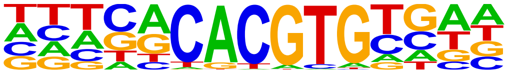 | BAM8(BES1)/col-BAM8-DAP-Seq(GSE60143)/Homer | 1e-4 | -1.028e+01 | 0.0001 | 2343.0 | 3.57% |
| 256 | 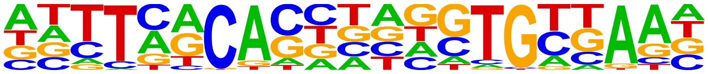 | Brachyury(T-box)/Mesoendoderm-Brachyury-ChIP-exo(GSE54963)/Homer | 1e-4 | -1.015e+01 | 0.0002 | 4496.0 | 6.86% |
| 257 | 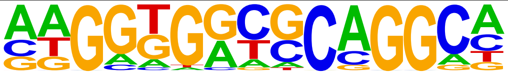 | ZNF165(Zf)/WHIM12-ZNF165-ChIP-Seq(GSE65937)/Homer | 1e-4 | -9.888e+00 | 0.0002 | 1990.0 | 3.04% |
| 258 | 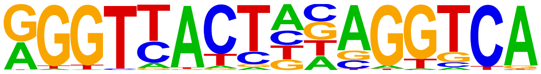 | LXRE(NR),DR4/RAW-LXRb.biotin-ChIP-Seq(GSE21512)/Homer | 1e-4 | -9.849e+00 | 0.0002 | 563.0 | 0.86% |
| 259 | 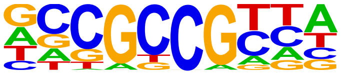 | CRF10(AP2EREBP)/col100-CRF10-DAP-Seq(GSE60143)/Homer | 1e-4 | -9.641e+00 | 0.0003 | 15923.0 | 24.29% |
| 260 | 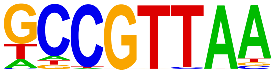 | AT3G10030(Trihelix)/colamp-AT3G10030-DAP-Seq(GSE60143)/Homer | 1e-4 | -9.419e+00 | 0.0003 | 9312.0 | 14.20% |
| 261 | 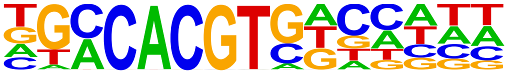 | ABF2(bZIP)/col-ABF2-DAP-Seq(GSE60143)/Homer | 1e-4 | -9.410e+00 | 0.0003 | 2991.0 | 4.56% |
| 262 | 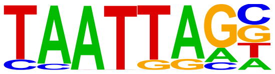 | Lhx2(Homeobox)/HFSC-Lhx2-ChIP-Seq(GSE48068)/Homer | 1e-4 | -9.354e+00 | 0.0003 | 15009.0 | 22.89% |
| 263 | 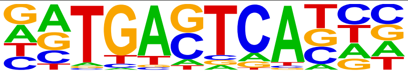 | Fosl2(bZIP)/3T3L1-Fosl2-ChIP-Seq(GSE56872)/Homer | 1e-3 | -8.392e+00 | 0.0009 | 3564.0 | 5.44% |
| 264 | 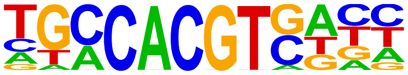 | bZIP28(bZIP)/col-bZIP28-DAP-Seq(GSE60143)/Homer | 1e-3 | -8.342e+00 | 0.0009 | 3730.0 | 5.69% |
| 265 | 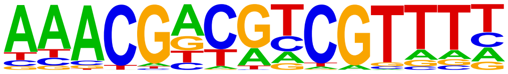 | Unknown2/Arabidopsis-Promoters/Homer | 1e-3 | -7.319e+00 | 0.0025 | 99.0 | 0.15% |
| 266 | 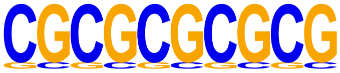 | SeqBias: CG-repeat | 1e-3 | -7.234e+00 | 0.0027 | 17509.0 | 26.71% |
| 267 | 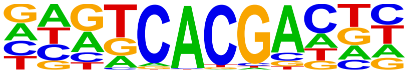 | Npas4(bHLH)/Neuron-Npas4-ChIP-Seq(GSE127793)/Homer | 1e-3 | -7.097e+00 | 0.0031 | 13728.0 | 20.94% |
| 268 | 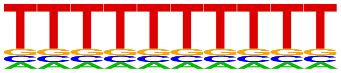 | VRN1(ABI3VP1)/col-VRN1-DAP-Seq(GSE60143)/Homer | 1e-3 | -6.938e+00 | 0.0036 | 3793.0 | 5.79% |

| | | | | | | | |
|---|---|---|---|---|---|---|---|
| 269 | 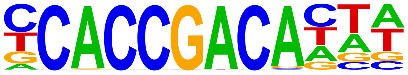 | AT1G01250(AP2EREBP)/col-AT1G01250-DAP-Seq(GSE60143)/Homer | 1e-2 | -6.872e+00 | 0.0039 | 1174.0 | 1.79% |
| 270 | 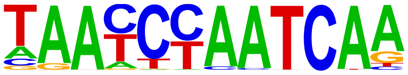 | Duxbl(Homeobox)/NIH3T3-Duxbl.HA-ChIP-Seq(GSE119782)/Homer | 1e-2 | -6.783e+00 | 0.0042 | 702.0 | 1.07% |
| 271 | 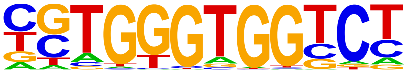 | Gli2(Zf)/GM2-Gli2-ChIP-Chip(GSE112702)/Homer | 1e-2 | -6.683e+00 | 0.0046 | 2788.0 | 4.25% |
| 272 | 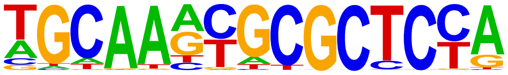 | EFL-1(E2F)/cElegans-L1-EFL1-ChIP-Seq(modEncode)/Homer | 1e-2 | -6.567e+00 | 0.0052 | 341.0 | 0.52% |
| 273 | 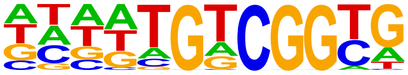 | AT1G71450(AP2EREBP)/col-AT1G71450-DAP-Seq(GSE60143)/Homer | 1e-2 | -6.143e+00 | 0.0079 | 17262.0 | 26.33% |
| 274 | 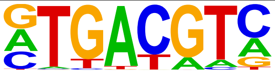 | TGA9(bZIP)/colamp-TGA9-DAP-Seq(GSE60143)/Homer | 1e-2 | -6.065e+00 | 0.0085 | 16267.0 | 24.81% |
| 275 | 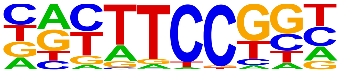 | Fli1(ETS)/CD8-FLI-ChIP-Seq(GSE20898)/Homer | 1e-2 | -5.843e+00 | 0.0106 | 13404.0 | 20.45% |
| 276 | 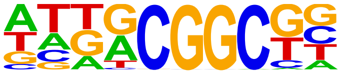 | ERF115(AP2EREBP)/colamp-ERF115-DAP-Seq(GSE60143)/Homer | 1e-2 | -5.671e+00 | 0.0126 | 18487.0 | 28.20% |
| 277 | 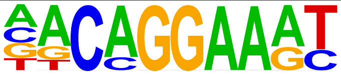 | EWS:FLI1-fusion(ETS)/SK_N_MC-EWS:FLI1-ChIP-Seq(SRA014231)/Homer | 1e-2 | -5.632e+00 | 0.0130 | 6765.0 | 10.32% |
| 278 | 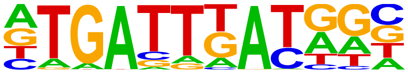 | PBX2(Homeobox)/K562-PBX2-ChIP-Seq(Encode)/Homer | 1e-2 | -5.609e+00 | 0.0133 | 10928.0 | 16.67% |
| 279 | 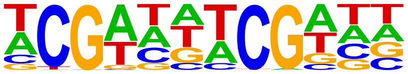 | AT5G22990(C2H2)/col-AT5G22990-DAP-Seq(GSE60143)/Homer | 1e-2 | -5.398e+00 | 0.0163 | 956.0 | 1.46% |
| 280 | 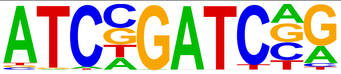 | GATA19(C2C2gata)/colamp-GATA19-DAP-Seq(GSE60143)/Homer | 1e-2 | -5.223e+00 | 0.0194 | 974.0 | 1.49% |
| 281 | 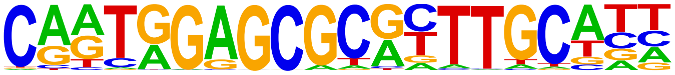 | LIN-15B(Zf)/cElegans-L3-LIN15B-ChIP-Seq(modEncode)/Homer | 1e-2 | -5.098e+00 | 0.0219 | 89.0 | 0.14% |
| 282 | 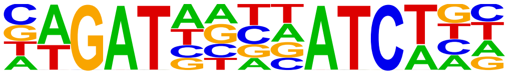 | GATA(Zf),IR4/iTreg-Gata3-ChIP-Seq(GSE20898)/Homer | 1e-2 | -4.970e+00 | 0.0248 | 1171.0 | 1.79% |
| 283 | 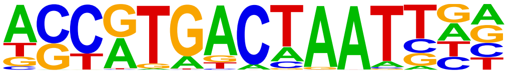 | PAX3:FKHR-fusion(Paired,Homeobox)/Rh4-PAX3:FKHR-ChIP-Seq(GSE19063)/Homer | 1e-2 | -4.911e+00 | 0.0262 | 2794.0 | 4.26% |
| 284 | 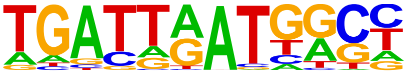 | Hoxb4(Homeobox)/ES-Hoxb4-ChIP-Seq(GSE34014)/Homer | 1e-2 | -4.813e+00 | 0.0288 | 2616.0 | 3.99% |
| 285 | 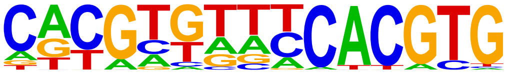 | bHLH18(bHLH)/col-bHLH18-DAP-Seq(GSE60143)/Homer | 1e-2 | -4.644e+00 | 0.0340 | 265.0 | 0.40% |

S3 Table. Full list of known motif discovery from ChIP-seq data targeting PBX1.

### Homer Known Motif Enrichment Results (/wynton/group/marcucio/2022CHM/Data2022Mar/Motif/HomerPBX1IDR)

[Homer *de novo* Motif Results]
[Gene Ontology Enrichment Results]
[Known Motif Enrichment Results (txt file)]

Total Target Sequences = 13432, Total Background Sequences = 36359

| Rank | Motif | Name | P-value | log P-pvalue | q-value (Benjamini) | # Target Sequenc with Motif |
|---|---|---|---|---|---|---|
| 1 | 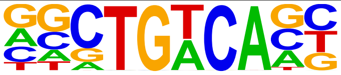 | Meis1(Homeobox)/MastCells-Meis1-ChIP-Seq(GSE48085)/Homer | 1e-701 | -1.614e+03 | 0.0000 | 4274.0 |
| 2 | 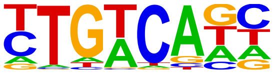 | Tgif1(Homeobox)/mES-Tgif1-ChIP-Seq(GSE55404)/Homer | 1e-608 | -1.402e+03 | 0.0000 | 5327.0 |
| 3 | 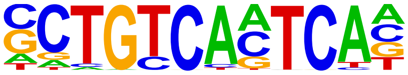 | Pbx3(Homeobox)/GM12878-PBX3-ChIP-Seq(GSE32465)/Homer | 1e-555 | -1.279e+03 | 0.0000 | 1184.0 |
| 4 | 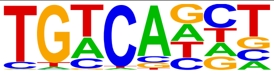 | Tgif2(Homeobox)/mES-Tgif2-ChIP-Seq(GSE55404)/Homer | 1e-487 | -1.122e+03 | 0.0000 | 5396.0 |
| 5 | 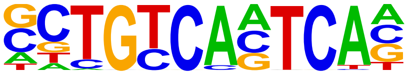 | Pknox1(Homeobox)/ES-Prep1-ChIP-Seq(GSE63282)/Homer | 1e-404 | -9.305e+02 | 0.0000 | 1120.0 |
| 6 | 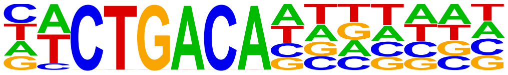 | GRF9(GRF)/colamp-GRF9-DAP-Seq(GSE60143)/Homer | 1e-395 | -9.103e+02 | 0.0000 | 2189.0 |
| 7 | 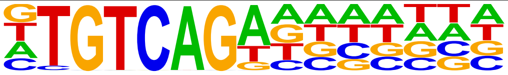 | AtGRF6(GRF)/col-AtGRF6-DAP-Seq(GSE60143)/Homer | 1e-356 | -8.200e+02 | 0.0000 | 2646.0 |
| 8 | 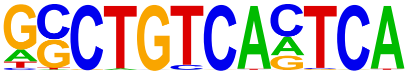 | PBX1(Homeobox)/MCF7-PBX1-ChIP-Seq(GSE28007)/Homer | 1e-311 | -7.181e+02 | 0.0000 | 506.0 |
| 9 | 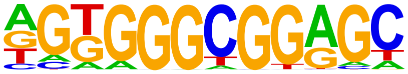 | Sp5(Zf)/mES-Sp5.Flag-ChIP-Seq(GSE72989)/Homer | 1e-123 | -2.833e+02 | 0.0000 | 4245.0 |
| 10 | 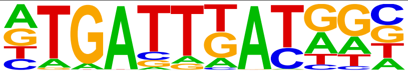 | PBX2(Homeobox)/K562-PBX2-ChIP-Seq(Encode)/Homer | 1e-119 | -2.749e+02 | 0.0000 | 1108.0 |
| 11 | 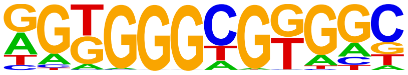 | KLF14(Zf)/HEK293-KLF14.GFP-ChIP-Seq(GSE58341)/Homer | 1e-115 | -2.666e+02 | 0.0000 | 6303.0 |
| 12 | 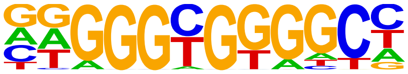 | KLF1(Zf)/HUDEP2-KLF1-CutnRun(GSE136251)/Homer | 1e-115 | -2.662e+02 | 0.0000 | 3253.0 |
| 13 | 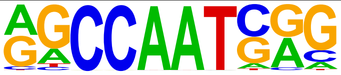 | NFY(CCAAT)/Promoter/Homer | 1e-108 | -2.487e+02 | 0.0000 | 1512.0 |
| 14 | 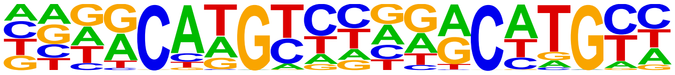 | p63(p53)/Keratinocyte-p63-ChIP-Seq(GSE17611)/Homer | 1e-106 | -2.443e+02 | 0.0000 | 653.0 |
| 15 | 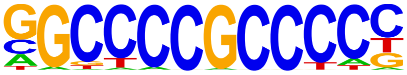 | Sp1(Zf)/Promoter/Homer | 1e-95 | -2.209e+02 | 0.0000 | 2341.0 |
| 16 | 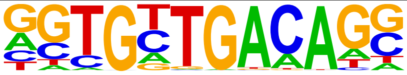 | Tbx20(T-box)/Heart-Tbx20-ChIP-Seq(GSE29636)/Homer | 1e-94 | -2.168e+02 | 0.0000 | 738.0 |
| 17 | 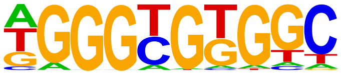 | KLF5(Zf)/LoVo-KLF5-ChIP-Seq(GSE49402)/Homer | 1e-93 | -2.142e+02 | 0.0000 | 4106.0 |
| 18 | 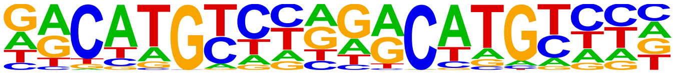 | p53(p53)/Saos-p53-ChIP-Seq(GSE15780)/Homer | 1e-92 | -2.124e+02 | 0.0000 | 261.0 |
| 19 | 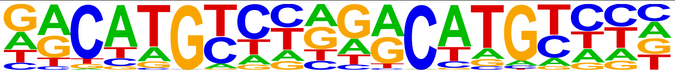 | p53(p53)/Saos-p53-ChIP-Seq/Homer | 1e-92 | -2.124e+02 | 0.0000 | 261.0 |
| 20 | 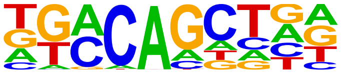 | bZIP18(bZIP)/colamp-bZIP18-DAP-Seq(GSE60143)/Homer | 1e-91 | -2.097e+02 | 0.0000 | 7253.0 |

| | | | | | | |
|---|---|---|---|---|---|---|
| 21 | 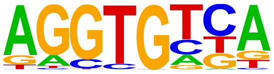 | Tbx5(T-box)/HL1-Tbx5.biotin-ChIP-Seq(GSE21529)/Homer | 1e-87 | -2.025e+02 | 0.0000 | 4208.0 |
| 22 | 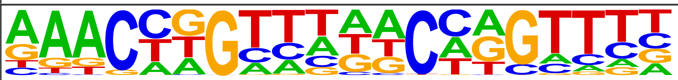 | GRHL2(CP2)/HBE-GRHL2-ChIP-Seq(GSE46194)/Homer | 1e-87 | -2.022e+02 | 0.0000 | 520.0 |
| 23 | 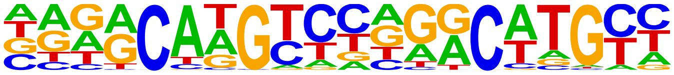 | p73(p53)/Trachea-p73-ChIP-Seq(PRJNA310161)/Homer | 1e-85 | -1.978e+02 | 0.0000 | 170.0 |
| 24 | 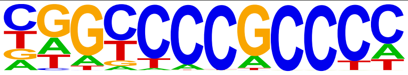 | Sp2(Zf)/HEK293-Sp2.eGFP-ChIP-Seq(Encode)/Homer | 1e-85 | -1.975e+02 | 0.0000 | 5298.0 |
| 25 | 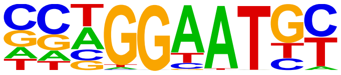 | TEAD4(TEA)/Tropoblast-Tead4-ChIP-Seq(GSE37350)/Homer | 1e-85 | -1.973e+02 | 0.0000 | 1080.0 |
| 26 | 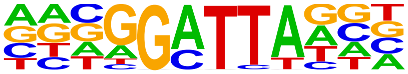 | bcd(Homeobox)/Embryo-Bcd-ChIP-Seq(GSE86966)/Homer | 1e-84 | -1.940e+02 | 0.0000 | 1262.0 |
| 27 | 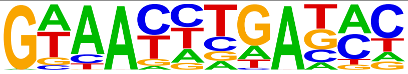 | Six2(Homeobox)/NephronProgenitor-Six2-ChIP-Seq(GSE39837)/Homer | 1e-82 | -1.889e+02 | 0.0000 | 1102.0 |
| 28 | 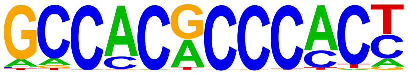 | Klf9(Zf)/GBM-Klf9-ChIP-Seq(GSE62211)/Homer | 1e-81 | -1.883e+02 | 0.0000 | 1362.0 |
| 29 | 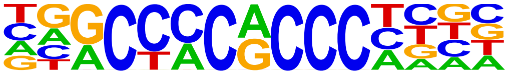 | KLF3(Zf)/MEF-Klf3-ChIP-Seq(GSE44748)/Homer | 1e-81 | -1.865e+02 | 0.0000 | 1901.0 |
| 30 | 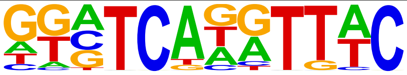 | Six1(Homeobox)/Myoblast-Six1-ChIP-Chip(GSE20150)/Homer | 1e-80 | -1.845e+02 | 0.0000 | 388.0 |
| 31 | 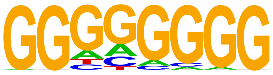 | Maz(Zf)/HepG2-Maz-ChIP-Seq(GSE31477)/Homer | 1e-78 | -1.817e+02 | 0.0000 | 5912.0 |
| 32 | 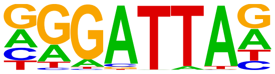 | GSC(Homeobox)/FrogEmbryos-GSC-ChIP-Seq(DRA000576)/Homer | 1e-74 | -1.710e+02 | 0.0000 | 1189.0 |
| 33 | 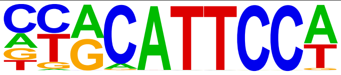 | TEAD1(TEAD)/HepG2-TEAD1-ChIP-Seq(Encode)/Homer | 1e-73 | -1.691e+02 | 0.0000 | 1106.0 |
| 34 | 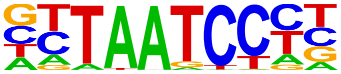 | Otx2(Homeobox)/EpiLC-Otx2-ChIP-Seq(GSE56098)/Homer | 1e-73 | -1.690e+02 | 0.0000 | 890.0 |
| 35 | 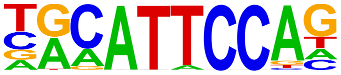 | TEAD3(TEA)/HepG2-TEAD3-ChIP-Seq(Encode)/Homer | 1e-72 | -1.677e+02 | 0.0000 | 1226.0 |
| 36 | 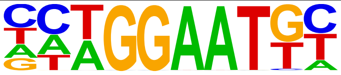 | TEAD(TEA)/Fibroblast-PU.1-ChIP-Seq(Unpublished)/Homer | 1e-68 | -1.584e+02 | 0.0000 | 774.0 |
| 37 | 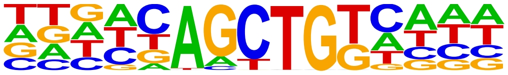 | bZIP52(bZIP)/colamp-bZIP52-DAP-Seq(GSE60143)/Homer | 1e-65 | -1.518e+02 | 0.0000 | 1884.0 |
| 38 | 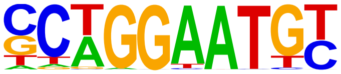 | TEAD2(TEA)/Py2T-Tead2-ChIP-Seq(GSE55709)/Homer | 1e-64 | -1.494e+02 | 0.0000 | 684.0 |
| 39 | 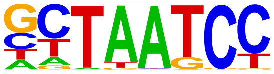 | CRX(Homeobox)/Retina-Crx-ChIP-Seq(GSE20012)/Homer | 1e-58 | -1.356e+02 | 0.0000 | 2170.0 |
| 40 | 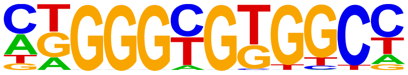 | KLF6(Zf)/PDAC-KLF6-ChIP-Seq(GSE64557)/Homer | 1e-57 | -1.327e+02 | 0.0000 | 3653.0 |
| 41 | 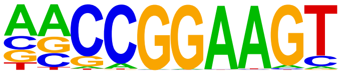 | ETS(ETS)/Promoter/Homer | 1e-53 | -1.237e+02 | 0.0000 | 801.0 |
| 42 | 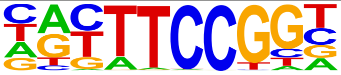 | Elk4(ETS)/Hela-Elk4-ChIP-Seq(GSE31477)/Homer | 1e-53 | -1.235e+02 | 0.0000 | 1381.0 |
| 43 | 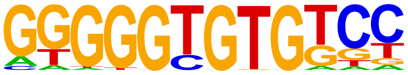 | KLF10(Zf)/HEK293-KLF10.GFP-ChIP-Seq(GSE58341)/Homer | 1e-52 | -1.209e+02 | 0.0000 | 1075.0 |
| 44 | 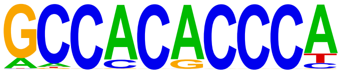 | Klf4(Zf)/mES-Klf4-ChIP-Seq(GSE11431)/Homer | 1e-51 | -1.188e+02 | 0.0000 | 783.0 |
| 45 | 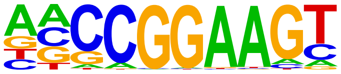 | ELF1(ETS)/Jurkat-ELF1-ChIP-Seq(SRA014231)/Homer | 1e-49 | -1.132e+02 | 0.0000 | 1196.0 |

| | | | | | | |
|---|---|---|---|---|---|---|
| 46 | 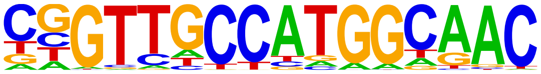 | RFX(HTH)/K562-RFX3-ChIP-Seq(SRA012198)/Homer | 1e-49 | -1.129e+02 | 0.0000 | 341.0 |
| 47 | 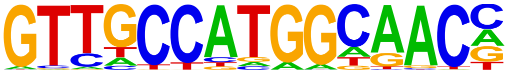 | Rfx2(HTH)/LoVo-RFX2-ChIP-Seq(GSE49402)/Homer | 1e-48 | -1.120e+02 | 0.0000 | 346.0 |
| 48 | 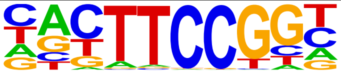 | Elk1(ETS)/Hela-Elk1-ChIP-Seq(GSE31477)/Homer | 1e-45 | -1.054e+02 | 0.0000 | 1399.0 |
| 49 | 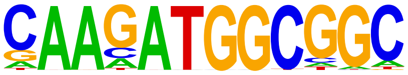 | YY1(Zf)/Promoter/Homer | 1e-44 | -1.021e+02 | 0.0000 | 525.0 |
| 50 | 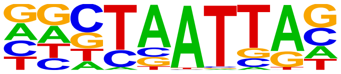 | En1(Homeobox)/SUM149-EN1-ChIP-Seq(GSE120957)/Homer | 1e-43 | -1.003e+02 | 0.0000 | 1858.0 |
| 51 | 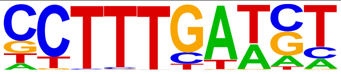 | LEF1(HMG)/H1-LEF1-ChIP-Seq(GSE64758)/Homer | 1e-42 | -9.707e+01 | 0.0000 | 774.0 |
| 52 | 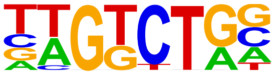 | Smad3(MAD)/NPC-Smad3-ChIP-Seq(GSE36673)/Homer | 1e-40 | -9.394e+01 | 0.0000 | 2779.0 |
| 53 | 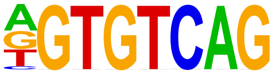 | CELF2(RRM)/JSL1-CELF2-CLIP-Seq(GSE71264)/Homer | 1e-40 | -9.290e+01 | 0.0000 | 559.0 |
| 54 | 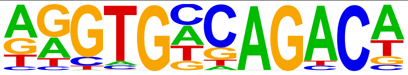 | Tbox:Smad(T-box,MAD)/ESCd5-Smad2_3-ChIP-Seq(GSE29422)/Homer | 1e-39 | -8.981e+01 | 0.0000 | 276.0 |
| 55 | 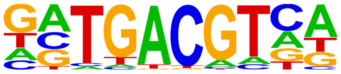 | bZIP50(bZIP)/colamp-bZIP50-DAP-Seq(GSE60143)/Homer | 1e-38 | -8.798e+01 | 0.0000 | 1495.0 |
| 56 | 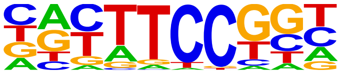 | Fli1(ETS)/CD8-FLI-ChIP-Seq(GSE20898)/Homer | 1e-37 | -8.620e+01 | 0.0000 | 1980.0 |
| 57 | 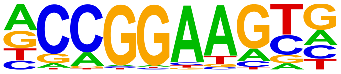 | ETV4(ETS)/HepG2-ETV4-ChIP-Seq(ENCODE)/Homer | 1e-37 | -8.610e+01 | 0.0000 | 2156.0 |
| 58 | 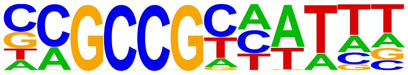 | RRTF1(AP2EREBP)/colamp-RRTF1-DAP-Seq(GSE60143)/Homer | 1e-37 | -8.596e+01 | 0.0000 | 1317.0 |
| 59 | 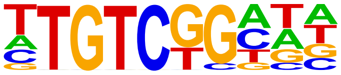 | ARF2(ARF)/col-ARF2-DAP-Seq(GSE60143)/Homer | 1e-37 | -8.583e+01 | 0.0000 | 3585.0 |
| 60 | 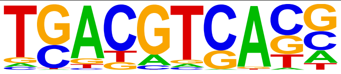 | FEA4(bZIP)/Corn-FEA4-ChIP-Seq(GSE61954)/Homer | 1e-37 | -8.538e+01 | 0.0000 | 1759.0 |
| 61 | 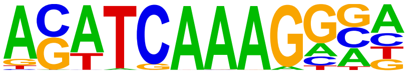 | Tcf4(HMG)/Hct116-Tcf4-ChIP-Seq(SRA012054)/Homer | 1e-36 | -8.445e+01 | 0.0000 | 539.0 |
| 62 | 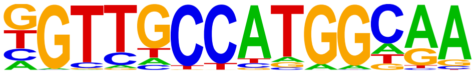 | Rfx1(HTH)/NPC-H3K4me1-ChIP-Seq(GSE16256)/Homer | 1e-36 | -8.419e+01 | 0.0000 | 449.0 |
| 63 | 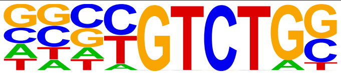 | Smad4(MAD)/ESC-SMAD4-ChIP-Seq(GSE29422)/Homer | 1e-36 | -8.304e+01 | 0.0000 | 2074.0 |
| 64 | 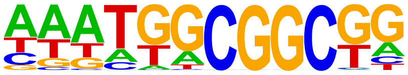 | ABR1(AP2EREBP)/colamp-ABR1-DAP-Seq(GSE60143)/Homer | 1e-35 | -8.227e+01 | 0.0000 | 5148.0 |
| 65 | 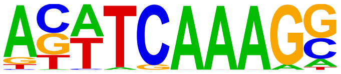 | Tcf3(HMG)/mES-Tcf3-ChIP-Seq(GSE11724)/Homer | 1e-34 | -7.976e+01 | 0.0000 | 331.0 |
| 66 | 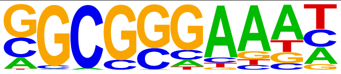 | E2F4(E2F)/K562-E2F4-ChIP-Seq(GSE31477)/Homer | 1e-34 | -7.944e+01 | 0.0000 | 1991.0 |
| 67 | 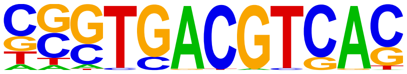 | CRE(bZIP)/Promoter/Homer | 1e-34 | -7.941e+01 | 0.0000 | 763.0 |
| 68 | 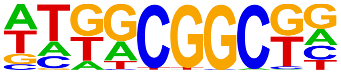 | At2g33710(AP2EREBP)/colamp-At2g33710-DAP-Seq(GSE60143)/Homer | 1e-34 | -7.852e+01 | 0.0000 | 7310.0 |
| 69 | 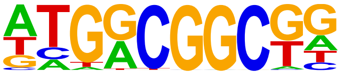 | ERF7(AP2EREBP)/col-ERF7-DAP-Seq(GSE60143)/Homer | 1e-32 | -7.447e+01 | 0.0000 | 6148.0 |
| 70 | 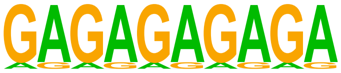 | SeqBias: GA-repeat | 1e-32 | -7.379e+01 | 0.0000 | 8836.0 |

| | | | | | | |
|---|---|---|---|---|---|---|
| 71 | 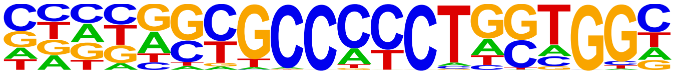 | BORIS(Zf)/K562-CTCFL-ChIP-Seq(GSE32465)/Homer | 1e-32 | -7.375e+01 | 0.0000 | 1084.0 |
| 72 | 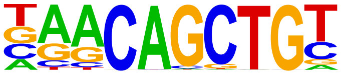 | Myf5(bHLH)/GM-Myf5-ChIP-Seq(GSE24852)/Homer | 1e-31 | -7.300e+01 | 0.0000 | 1373.0 |
| 73 | 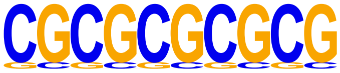 | SeqBias: CG-repeat | 1e-31 | -7.287e+01 | 0.0000 | 8741.0 |
| 74 | 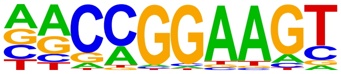 | GABPA(ETS)/Jurkat-GABPa-ChIP-Seq(GSE17954)/Homer | 1e-31 | -7.185e+01 | 0.0000 | 1416.0 |
| 75 | 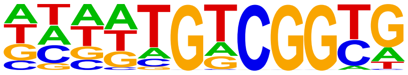 | AT1G71450(AP2EREBP)/col-AT1G71450-DAP-Seq(GSE60143)/Homer | 1e-31 | -7.156e+01 | 0.0000 | 5605.0 |
| 76 | 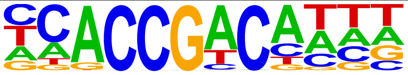 | DEAR2(AP2EREBP)/colamp-DEAR2-DAP-Seq(GSE60143)/Homer | 1e-30 | -7.120e+01 | 0.0000 | 2788.0 |
| 77 | 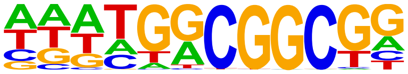 | ERF4(AP2EREBP)/colamp-ERF4-DAP-Seq(GSE60143)/Homer | 1e-30 | -7.102e+01 | 0.0000 | 5732.0 |
| 78 | 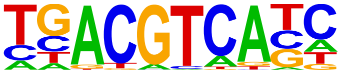 | TGA6(bZIP)/colamp-TGA6-DAP-Seq(GSE60143)/Homer | 1e-30 | -7.017e+01 | 0.0000 | 983.0 |
| 79 | 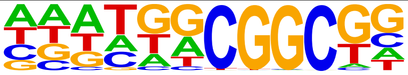 | RAP26(AP2EREBP)/colamp-RAP26-DAP-Seq(GSE60143)/Homer | 1e-30 | -7.010e+01 | 0.0000 | 6295.0 |
| 80 | 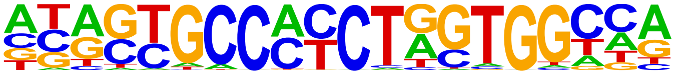 | CTCF(Zf)/CD4+-CTCF-ChIP-Seq(Barski_et_al.)/Homer | 1e-30 | -6.995e+01 | 0.0000 | 491.0 |
| 81 | 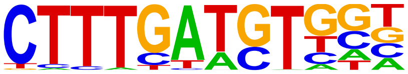 | Tcf7(HMG)/GM12878-TCF7-ChIP-Seq(Encode)/Homer | 1e-30 | -6.957e+01 | 0.0000 | 417.0 |
| 82 | 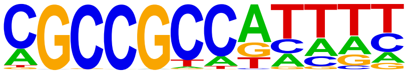 | RAP212(AP2EREBP)/col-RAP212-DAP-Seq(GSE60143)/Homer | 1e-29 | -6.876e+01 | 0.0000 | 4669.0 |
| 83 | 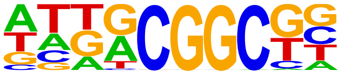 | ERF115(AP2EREBP)/colamp-ERF115-DAP-Seq(GSE60143)/Homer | 1e-29 | -6.800e+01 | 0.0000 | 7330.0 |
| 84 | 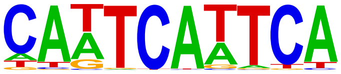 | WUS1(Homeobox)/colamp-WUS1-DAP-Seq(GSE60143)/Homer | 1e-29 | -6.731e+01 | 0.0000 | 434.0 |
| 85 | 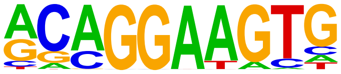 | ETS1(ETS)/Jurkat-ETS1-ChIP-Seq(GSE17954)/Homer | 1e-28 | -6.640e+01 | 0.0000 | 1611.0 |
| 86 | 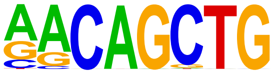 | MyoG(bHLH)/C2C12-MyoG-ChIP-Seq(GSE36024)/Homer | 1e-28 | -6.589e+01 | 0.0000 | 1963.0 |
| 87 | 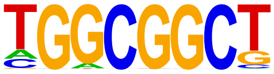 | ERF105(AP2EREBP)/colamp-ERF105-DAP-Seq(GSE60143)/Homer | 1e-28 | -6.492e+01 | 0.0000 | 7098.0 |
| 88 | 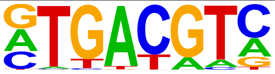 | TGA9(bZIP)/colamp-TGA9-DAP-Seq(GSE60143)/Homer | 1e-28 | -6.485e+01 | 0.0000 | 1533.0 |
| 89 | 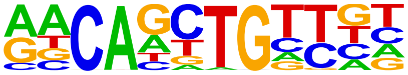 | HLH-1(bHLH)/cElegans-Embryo-HLH1-ChIP-Seq(modEncode)/Homer | 1e-28 | -6.469e+01 | 0.0000 | 1023.0 |
| 90 | 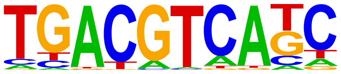 | TGA1(bZIP)/colamp-TGA1-DAP-Seq(GSE60143)/Homer | 1e-27 | -6.410e+01 | 0.0000 | 677.0 |
| 91 | 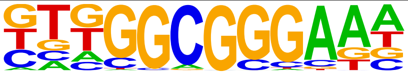 | E2F3(E2F)/MEF-E2F3-ChIP-Seq(GSE71376)/Homer | 1e-27 | -6.391e+01 | 0.0000 | 2825.0 |
| 92 | 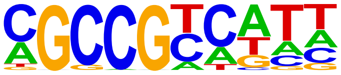 | ERF8(AP2EREBP)/colamp-ERF8-DAP-Seq(GSE60143)/Homer | 1e-27 | -6.366e+01 | 0.0000 | 5619.0 |
| 93 | 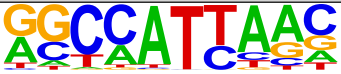 | Nanog(Homeobox)/mES-Nanog-ChIP-Seq(GSE11724)/Homer | 1e-27 | -6.350e+01 | 0.0000 | 4240.0 |
| 94 | 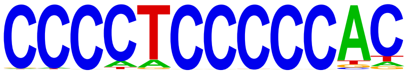 | Zfp281(Zf)/ES-Zfp281-ChIP-Seq(GSE81042)/Homer | 1e-27 | -6.300e+01 | 0.0000 | 688.0 |
| 95 | 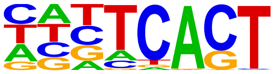 | STZ(C2H2)/colamp-STZ-DAP-Seq(GSE60143)/Homer | 1e-26 | -6.145e+01 | 0.0000 | 5336.0 |

| | | | | | | |
|---|---|---|---|---|---|---|
| 96 | 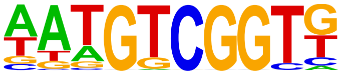 | DREB19(AP2EREBP)/colamp-DREB19-DAP-Seq(GSE60143)/Homer | 1e-26 | -6.115e+01 | 0.0000 | 1480.0 |
| 97 | 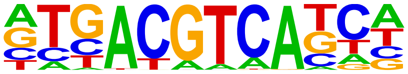 | TGA4(bZIP)/colamp-TGA4-DAP-Seq(GSE60143)/Homer | 1e-26 | -6.045e+01 | 0.0000 | 534.0 |
| 98 | 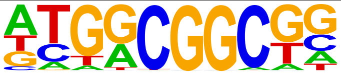 | ERF3(AP2EREBP)/colamp-ERF3-DAP-Seq(GSE60143)/Homer | 1e-26 | -6.012e+01 | 0.0000 | 4988.0 |
| 99 | 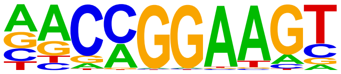 | ETV1(ETS)/GIST48-ETV1-ChIP-Seq(GSE22441)/Homer | 1e-26 | -6.006e+01 | 0.0000 | 1922.0 |
| 100 | 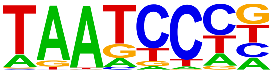 | Pitx1(Homeobox)/Chicken-Pitx1-ChIP-Seq(GSE38910)/Homer | 1e-26 | -5.990e+01 | 0.0000 | 3718.0 |
| 101 | 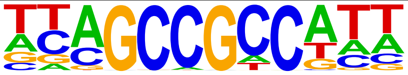 | ERF13(AP2EREBP)/colamp-ERF13-DAP-Seq(GSE60143)/Homer | 1e-25 | -5.945e+01 | 0.0000 | 5520.0 |
| 102 | 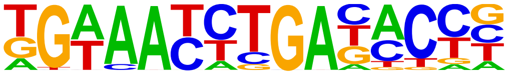 | Six4(Homeobox)/MCF7-SIX4-ChIP-Seq(Encode)/Homer | 1e-25 | -5.803e+01 | 0.0000 | 105.0 |
| 103 | 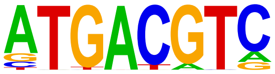 | TGA10(bZIP)/colamp-TGA10-DAP-Seq(GSE60143)/Homer | 1e-24 | -5.694e+01 | 0.0000 | 878.0 |
| 104 | 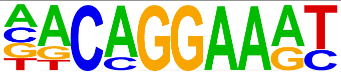 | EWS:FLI1-fusion(ETS)/SK_N_MC-EWS:FLI1-ChIP-Seq(SRA014231)/Homer | 1e-24 | -5.595e+01 | 0.0000 | 875.0 |
| 105 | 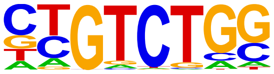 | Smad2(MAD)/ES-SMAD2-ChIP-Seq(GSE29422)/Homer | 1e-24 | -5.578e+01 | 0.0000 | 2023.0 |
| 106 | 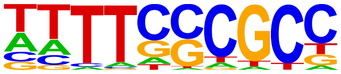 | DEL2(E2FDP)/col-DEL2-DAP-Seq(GSE60143)/Homer | 1e-24 | -5.570e+01 | 0.0000 | 947.0 |
| 107 | 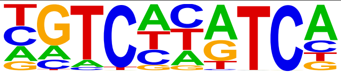 | Replumless(BLH)/Arabidopsis-RPL.GFP-ChIP-Seq(GSE78727)/Homer | 1e-23 | -5.366e+01 | 0.0000 | 1220.0 |
| 108 | 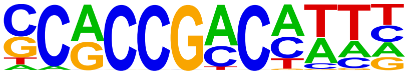 | At5g65130(AP2EREBP)/colamp-At5g65130-DAP-Seq(GSE60143)/Homer | 1e-23 | -5.312e+01 | 0.0000 | 1392.0 |
| 109 | 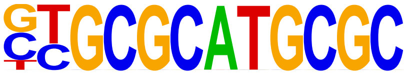 | NRF(NRF)/Promoter/Homer | 1e-23 | -5.305e+01 | 0.0000 | 777.0 |
| 110 | 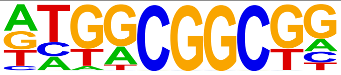 | ERF11(AP2EREBP)/col-ERF11-DAP-Seq(GSE60143)/Homer | 1e-23 | -5.298e+01 | 0.0000 | 5499.0 |
| 111 | 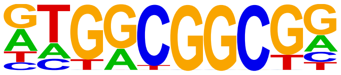 | ERF10(AP2EREBP)/col-ERF10-DAP-Seq(GSE60143)/Homer | 1e-22 | -5.295e+01 | 0.0000 | 4957.0 |
| 112 | 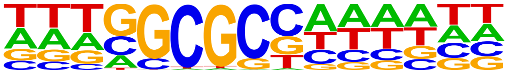 | E2FA(E2FDP)/colamp-E2FA-DAP-Seq(GSE60143)/Homer | 1e-22 | -5.269e+01 | 0.0000 | 757.0 |
| 113 | 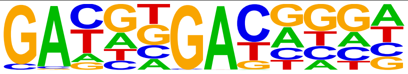 | Knotted(Homeobox)/Corn-KN1-ChIP-Seq(GSE39161)/Homer | 1e-22 | -5.249e+01 | 0.0000 | 2474.0 |
| 114 | 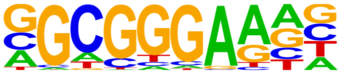 | E2F6(E2F)/Hela-E2F6-ChIP-Seq(GSE31477)/Homer | 1e-22 | -5.127e+01 | 0.0000 | 2621.0 |
| 115 | 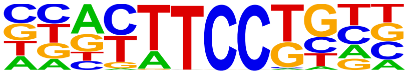 | Etv2(ETS)/ES-ER71-ChIP-Seq(GSE59402)/Homer | 1e-22 | -5.124e+01 | 0.0000 | 1233.0 |
| 116 | 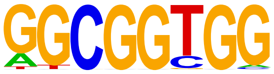 | AT3G57600(AP2EREBP)/col-AT3G57600-DAP-Seq(GSE60143)/Homer | 1e-22 | -5.121e+01 | 0.0000 | 5864.0 |
| 117 | 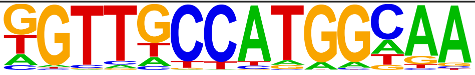 | X-box(HTH)/NPC-H3K4me1-ChIP-Seq(GSE16256)/Homer | 1e-21 | -5.055e+01 | 0.0000 | 219.0 |
| 118 | 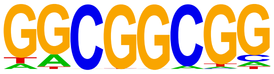 | AT1G28160(AP2EREBP)/colamp-AT1G28160-DAP-Seq(GSE60143)/Homer | 1e-21 | -5.031e+01 | 0.0000 | 8242.0 |
| 119 | 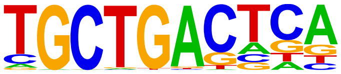 | MafA(bZIP)/Islet-MafA-ChIP-Seq(GSE30298)/Homer | 1e-21 | -4.994e+01 | 0.0000 | 1344.0 |
| 120 | | E2F7(E2F)/Hela-E2F7-ChIP-Seq(GSE32673)/Homer | 1e-21 | -4.978e+01 | 0.0000 | 535.0 |

| | | | | | | |
|---|---|---|---|---|---|---|
| | 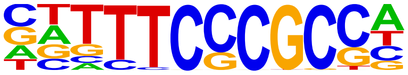 | | | | | |
| 121 | 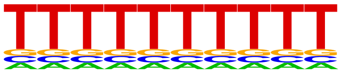 | VRN1(ABI3VP1)/col-VRN1-DAP-Seq(GSE60143)/Homer | 1e-20 | -4.793e+01 | 0.0000 | 299.0 |
| 122 | 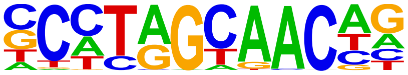 | Rfx5(HTH)/GM12878-Rfx5-ChIP-Seq(GSE31477)/Homer | 1e-20 | -4.740e+01 | 0.0000 | 450.0 |
| 123 | 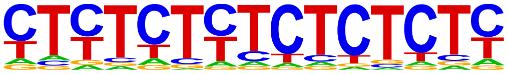 | GAGA-repeat/SacCer-Promoters/Homer | 1e-20 | -4.717e+01 | 0.0000 | 4209.0 |
| 124 | 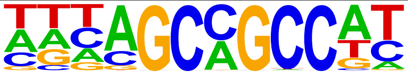 | ERF15(AP2EREBP)/colamp-ERF15-DAP-Seq(GSE60143)/Homer | 1e-20 | -4.688e+01 | 0.0000 | 7653.0 |
| 125 | 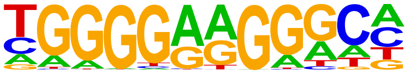 | ZNF467(Zf)/HEK293-ZNF467.GFP-ChIP-Seq(GSE58341)/Homer | 1e-20 | -4.681e+01 | 0.0000 | 2628.0 |
| 126 | 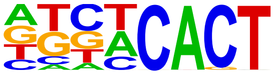 | AZF1(C2H2)/colamp-AZF1-DAP-Seq(GSE60143)/Homer | 1e-20 | -4.670e+01 | 0.0000 | 4879.0 |
| 127 | 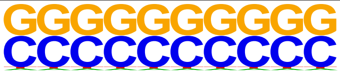 | SeqBias: CG bias | 1e-20 | -4.664e+01 | 0.0000 | 11976.0 |
| 128 | 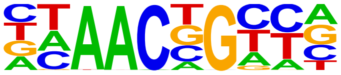 | BMYB(HTH)/Hela-BMYB-ChIP-Seq(GSE27030)/Homer | 1e-19 | -4.522e+01 | 0.0000 | 2102.0 |
| 129 | 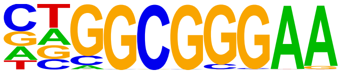 | E2F1(E2F)/Hela-E2F1-ChIP-Seq(GSE22478)/Homer | 1e-19 | -4.468e+01 | 0.0000 | 1279.0 |
| 130 | 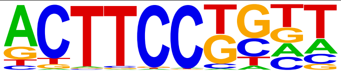 | Elf4(ETS)/BMDM-Elf4-ChIP-Seq(GSE88699)/Homer | 1e-18 | -4.336e+01 | 0.0000 | 1236.0 |
| 131 | 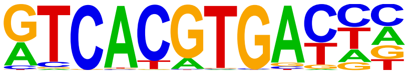 | TFE3(bHLH)/MEF-TFE3-ChIP-Seq(GSE75757)/Homer | 1e-18 | -4.233e+01 | 0.0000 | 189.0 |
| 132 | 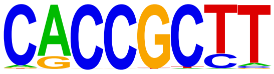 | At5g18450(AP2EREBP)/col-At5g18450-DAP-Seq(GSE60143)/Homer | 1e-18 | -4.204e+01 | 0.0000 | 6977.0 |
| 133 | 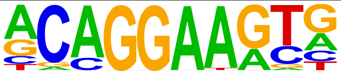 | ERG(ETS)/VCaP-ERG-ChIP-Seq(GSE14097)/Homer | 1e-18 | -4.184e+01 | 0.0000 | 1965.0 |
| 134 | 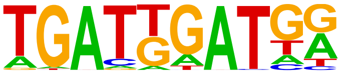 | HOXA1(Homeobox)/mES-Hoxa1-ChIP-Seq(SRP084292)/Homer | 1e-17 | -4.126e+01 | 0.0000 | 303.0 |
| 135 | 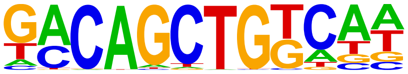 | bZIP69(bZIP)/col-bZIP69-DAP-Seq(GSE60143)/Homer | 1e-17 | -4.056e+01 | 0.0000 | 215.0 |
| 136 | 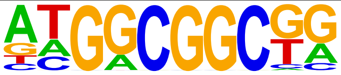 | AT4G18450(AP2EREBP)/col-AT4G18450-DAP-Seq(GSE60143)/Homer | 1e-17 | -4.033e+01 | 0.0000 | 4016.0 |
| 137 | 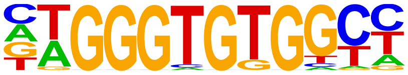 | EKLF(Zf)/Erythrocyte-Klf1-ChIP-Seq(GSE20478)/Homer | 1e-17 | -3.997e+01 | 0.0000 | 182.0 |
| 138 | 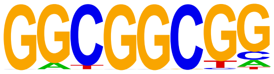 | ESE1(AP2EREBP)/col-ESE1-DAP-Seq(GSE60143)/Homer | 1e-17 | -3.984e+01 | 0.0000 | 6283.0 |
| 139 | 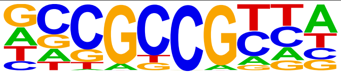 | CRF10(AP2EREBP)/col100-CRF10-DAP-Seq(GSE60143)/Homer | 1e-17 | -3.958e+01 | 0.0000 | 7424.0 |
| 140 | 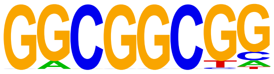 | ERF104(AP2EREBP)/col-ERF104-DAP-Seq(GSE60143)/Homer | 1e-17 | -3.953e+01 | 0.0000 | 6848.0 |
| 141 | 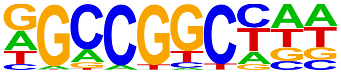 | RAP211(AP2EREBP)/colamp-RAP211-DAP-Seq(GSE60143)/Homer | 1e-17 | -3.950e+01 | 0.0000 | 6697.0 |
| 142 | 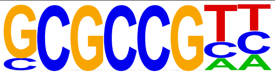 | PUCHI(AP2EREBP)/colamp-PUCHI-DAP-Seq(GSE60143)/Homer | 1e-17 | -3.944e+01 | 0.0000 | 6249.0 |
| 143 | 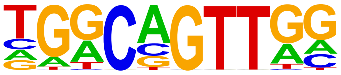 | AMYB(HTH)/Testes-AMYB-ChIP-Seq(GSE44588)/Homer | 1e-17 | -3.940e+01 | 0.0000 | 2128.0 |
| 144 | 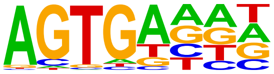 | At5g04390(C2H2)/col200-At5g04390-DAP-Seq(GSE60143)/Homer | 1e-17 | -3.925e+01 | 0.0000 | 4922.0 |

| | | | | | | |
|---|---|---|---|---|---|---|
| 145 | 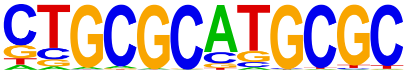 | NRF1(NRF)/MCF7-NRF1-ChIP-Seq(Unpublished)/Homer | 1e-16 | -3.896e+01 | 0.0000 | 814.0 |
| 146 | 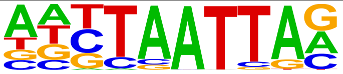 | Lhx3(Homeobox)/Neuron-Lhx3-ChIP-Seq(GSE31456)/Homer | 1e-16 | -3.866e+01 | 0.0000 | 1406.0 |
| 147 | 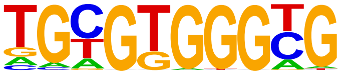 | Egr1(Zf)/K562-Egr1-ChIP-Seq(GSE32465)/Homer | 1e-16 | -3.830e+01 | 0.0000 | 2170.0 |
| 148 | 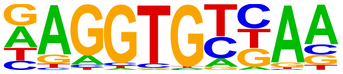 | Tbx6(T-box)/ESC-Tbx6-ChIP-Seq(GSE93524)/Homer | 1e-16 | -3.821e+01 | 0.0000 | 1133.0 |
| 149 | 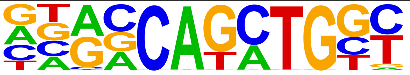 | Atoh1(bHLH)/Cerebellum-Atoh1-ChIP-Seq(GSE22111)/Homer | 1e-16 | -3.798e+01 | 0.0000 | 1558.0 |
| 150 | 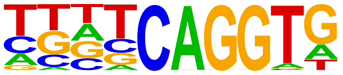 | NGA4(ABI3VP1)/col-NGA4-DAP-Seq(GSE60143)/Homer | 1e-16 | -3.750e+01 | 0.0000 | 3258.0 |
| 151 | 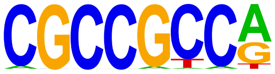 | CRF4(AP2EREBP)/colamp-CRF4-DAP-Seq(GSE60143)/Homer | 1e-15 | -3.669e+01 | 0.0000 | 4833.0 |
| 152 | 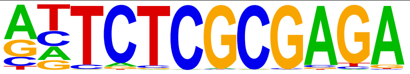 | GFX(?)/Promoter/Homer | 1e-15 | -3.654e+01 | 0.0000 | 102.0 |
| 153 | 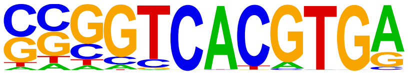 | E-box(bHLH)/Promoter/Homer | 1e-15 | -3.638e+01 | 0.0000 | 263.0 |
| 154 | 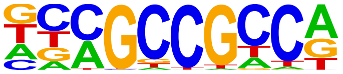 | ERF5(AP2EREBP)/colamp-ERF5-DAP-Seq(GSE60143)/Homer | 1e-15 | -3.586e+01 | 0.0000 | 4296.0 |
| 155 | 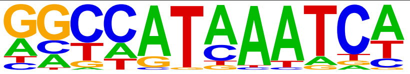 | HOXA9(Homeobox)/HSC-Hoxa9-ChIP-Seq(GSE33509)/Homer | 1e-15 | -3.568e+01 | 0.0000 | 598.0 |
| 156 | 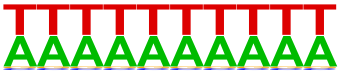 | SeqBias: A/T bias | 1e-15 | -3.558e+01 | 0.0000 | 5914.0 |
| 157 | 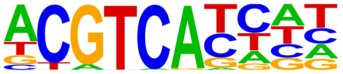 | TGA2(bZIP)/colamp-TGA2-DAP-Seq(GSE60143)/Homer | 1e-15 | -3.554e+01 | 0.0000 | 985.0 |
| 158 | 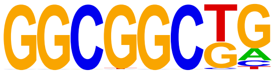 | ERF2(AP2EREBP)/colamp-ERF2-DAP-Seq(GSE60143)/Homer | 1e-15 | -3.502e+01 | 0.0000 | 5993.0 |
| 159 | 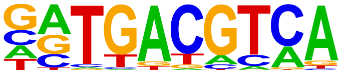 | Atf1(bZIP)/K562-ATF1-ChIP-Seq(GSE31477)/Homer | 1e-14 | -3.449e+01 | 0.0000 | 849.0 |
| 160 | 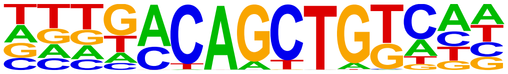 | VIP1(bZIP)/col-VIP1-DAP-Seq(GSE60143)/Homer | 1e-14 | -3.442e+01 | 0.0000 | 286.0 |
| 161 | 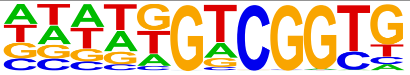 | At1g36060(AP2EREBP)/colamp-At1g36060-DAP-Seq(GSE60143)/Homer | 1e-14 | -3.411e+01 | 0.0000 | 1600.0 |
| 162 | 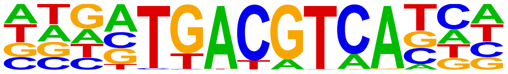 | TGA5(bZIP)/col-TGA5-DAP-Seq(GSE60143)/Homer | 1e-14 | -3.407e+01 | 0.0000 | 253.0 |
| 163 | 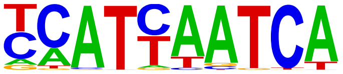 | Pdx1(Homeobox)/Islet-Pdx1-ChIP-Seq(SRA008281)/Homer | 1e-14 | -3.390e+01 | 0.0000 | 817.0 |
| 164 | 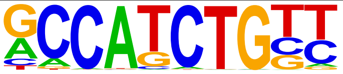 | NeuroD1(bHLH)/Islet-NeuroD1-ChIP-Seq(GSE30298)/Homer | 1e-14 | -3.367e+01 | 0.0000 | 1007.0 |
| 165 | 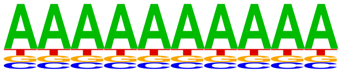 | SeqBias: polyA-repeat | 1e-14 | -3.342e+01 | 0.0000 | 10468.0 |
| 166 | 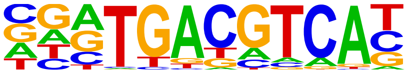 | Atf7(bZIP)/3T3L1-Atf7-ChIP-Seq(GSE56872)/Homer | 1e-14 | -3.342e+01 | 0.0000 | 670.0 |
| 167 | 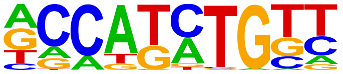 | Olig2(bHLH)/Neuron-Olig2-ChIP-Seq(GSE30882)/Homer | 1e-14 | -3.324e+01 | 0.0000 | 2102.0 |
| 168 | 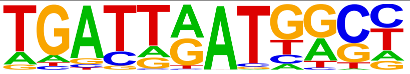 | Hoxb4(Homeobox)/ES-Hoxb4-ChIP-Seq(GSE34014)/Homer | 1e-14 | -3.290e+01 | 0.0000 | 220.0 |
| 169 | 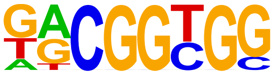 | ESE3(AP2EREBP)/col-ESE3-DAP-Seq(GSE60143)/Homer | 1e-14 | -3.286e+01 | 0.0000 | 7119.0 |

| | | | | | | |
|---|---|---|---|---|---|---|
| 170 | 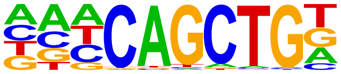 | Ap4(bHLH)/AML-Tfap4-ChIP-Seq(GSE45738)/Homer | 1e-14 | -3.240e+01 | 0.0000 | 2047.0 |
| 171 | 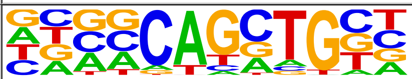 | Ascl1(bHLH)/NeuralTubes-Ascl1-ChIP-Seq(GSE55840)/Homer | 1e-14 | -3.229e+01 | 0.0000 | 2475.0 |
| 172 | 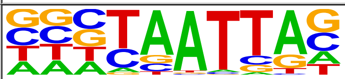 | DLX2(Homeobox)/BasalGanglia-Dlx2-ChIP-seq(GSE124936)/Homer | 1e-13 | -3.222e+01 | 0.0000 | 1239.0 |
| 173 | 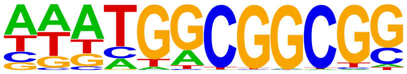 | ERF9(AP2EREBP)/colamp-ERF9-DAP-Seq(GSE60143)/Homer | 1e-13 | -3.208e+01 | 0.0000 | 2994.0 |
| 174 | 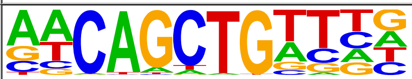 | E-box/Drosophila-Promoters/Homer | 1e-13 | -3.177e+01 | 0.0000 | 395.0 |
| 175 | 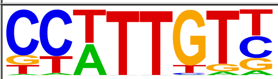 | Sox3(HMG)/NPC-Sox3-ChIP-Seq(GSE33059)/Homer | 1e-13 | -3.174e+01 | 0.0000 | 1787.0 |
| 176 | 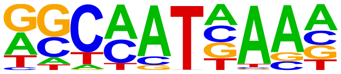 | Hoxa9(Homeobox)/ChickenMSG-Hoxa9.Flag-ChIP-Seq(GSE86088)/Homer | 1e-13 | -3.146e+01 | 0.0000 | 2410.0 |
| 177 | 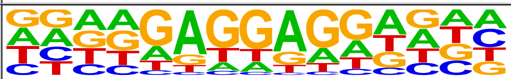 | TF3A(C2H2)/col-TF3A-DAP-Seq(GSE60143)/Homer | 1e-13 | -3.120e+01 | 0.0000 | 2294.0 |
| 178 | 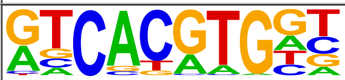 | Usf2(bHLH)/C2C12-Usf2-ChIP-Seq(GSE36030)/Homer | 1e-13 | -3.111e+01 | 0.0000 | 469.0 |
| 179 | 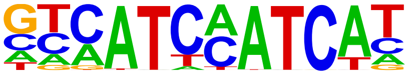 | HOXA2(Homeobox)/mES-Hoxa2-ChIP-Seq(Donaldson_et_al.)/Homer | 1e-13 | -3.043e+01 | 0.0000 | 134.0 |
| 180 | 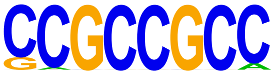 | ERF73(AP2EREBP)/col-ERF73-DAP-Seq(GSE60143)/Homer | 1e-13 | -3.010e+01 | 0.0000 | 5840.0 |
| 181 | 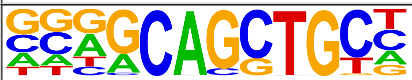 | Ascl2(bHLH)/ESC-Ascl2-ChIP-Seq(GSE97712)/Homer | 1e-12 | -2.950e+01 | 0.0000 | 2089.0 |
| 182 | 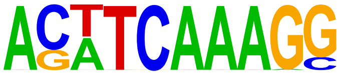 | TCFL2(HMG)/K562-TCF7L2-ChIP-Seq(GSE29196)/Homer | 1e-12 | -2.943e+01 | 0.0000 | 124.0 |
| 183 | 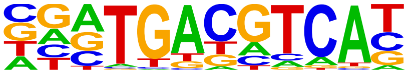 | Atf2(bZIP)/3T3L1-Atf2-ChIP-Seq(GSE56872)/Homer | 1e-12 | -2.925e+01 | 0.0000 | 507.0 |
| 184 | 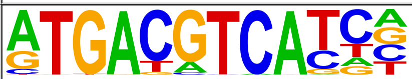 | JunD(bZIP)/K562-JunD-ChIP-Seq/Homer | 1e-12 | -2.884e+01 | 0.0000 | 247.0 |
| 185 | 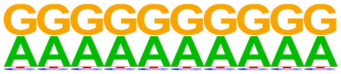 | SeqBias: G/A bias | 1e-12 | -2.867e+01 | 0.0000 | 13429.0 |
| 186 | 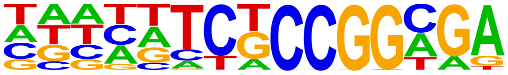 | AT5G05550(Trihelix)/col-AT5G05550-DAP-Seq(GSE60143)/Homer | 1e-12 | -2.855e+01 | 0.0000 | 3546.0 |
| 187 | 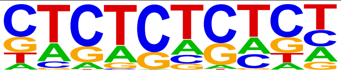 | GAGA-repeat/Arabidopsis-Promoters/Homer | 1e-12 | -2.839e+01 | 0.0000 | 1246.0 |
| 188 | 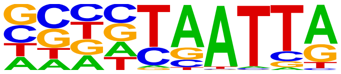 | DLX1(Homeobox)/BasalGanglia-Dlx1-ChIP-seq(GSE124936)/Homer | 1e-12 | -2.826e+01 | 0.0000 | 1141.0 |
| 189 | 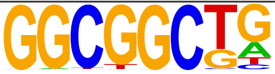 | AT5G23930(mTERF)/col-AT5G23930-DAP-Seq(GSE60143)/Homer | 1e-12 | -2.795e+01 | 0.0000 | 7628.0 |
| 190 | 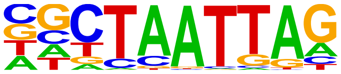 | LHX9(Homeobox)/Hct116-LHX9.V5-ChIP-Seq(GSE116822)/Homer | 1e-12 | -2.789e+01 | 0.0000 | 1202.0 |
| 191 | 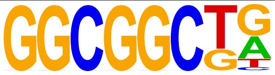 | ERF1(AP2EREBP)/colamp-ERF1-DAP-Seq(GSE60143)/Homer | 1e-12 | -2.775e+01 | 0.0000 | 5439.0 |
| 192 | 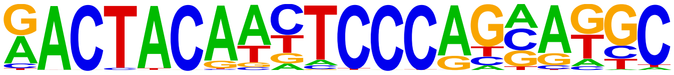 | Ronin(THAP)/ES-Thap11-ChIP-Seq(GSE51522)/Homer | 1e-11 | -2.756e+01 | 0.0000 | 158.0 |
| 193 | 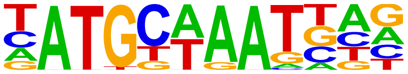 | Brn1(POU,Homeobox)/NPC-Brn1-ChIP-Seq(GSE35496)/Homer | 1e-11 | -2.728e+01 | 0.0000 | 233.0 |
| 194 | 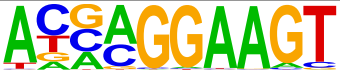 | ELF5(ETS)/T47D-ELF5-ChIP-Seq(GSE30407)/Homer | 1e-11 | -2.713e+01 | 0.0000 | 772.0 |

| | | | | | | |
|---|---|---|---|---|---|---|
| 195 | 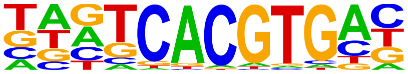 | BIM3(bHLH)/col-BIM3-DAP-Seq(GSE60143)/Homer | 1e-11 | -2.694e+01 | 0.0000 | 247.0 |
| 196 | 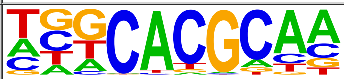 | Arnt:Ahr(bHLH)/MCF7-Arnt-ChIP-Seq(Lo_et_al.)/Homer | 1e-11 | -2.692e+01 | 0.0000 | 1451.0 |
| 197 | 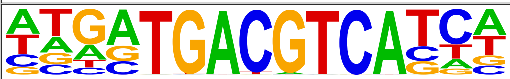 | TGA3(bZIP)/colamp-TGA3-DAP-Seq(GSE60143)/Homer | 1e-11 | -2.679e+01 | 0.0000 | 195.0 |
| 198 | 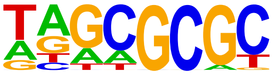 | DPL-1(E2F)/cElegans-Adult-ChIP-Seq(modEncode)/Homer | 1e-11 | -2.637e+01 | 0.0000 | 3146.0 |
| 199 | 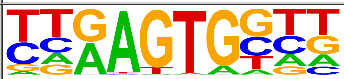 | Bapx1(Homeobox)/VertebralCol-Bapx1-ChIP-Seq(GSE36672)/Homer | 1e-11 | -2.620e+01 | 0.0000 | 1894.0 |
| 200 | 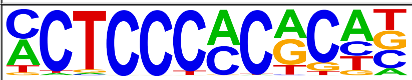 | WT1(Zf)/Kidney-WT1-ChIP-Seq(GSE90016)/Homer | 1e-11 | -2.618e+01 | 0.0000 | 1716.0 |
| 201 | 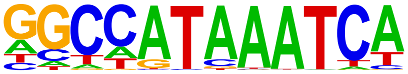 | Hoxc9(Homeobox)/Ainv15-Hoxc9-ChIP-Seq(GSE21812)/Homer | 1e-11 | -2.602e+01 | 0.0000 | 445.0 |
| 202 | 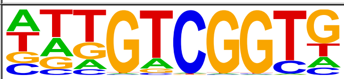 | CEJ1(AP2EREBP)/col-CEJ1-DAP-Seq(GSE60143)/Homer | 1e-11 | -2.555e+01 | 0.0000 | 2031.0 |
| 203 | 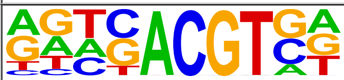 | HY5(bZIP)/colamp-HY5-DAP-Seq(GSE60143)/Homer | 1e-10 | -2.523e+01 | 0.0000 | 1258.0 |
| 204 | 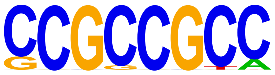 | SHN3(AP2EREBP)/col-SHN3-DAP-Seq(GSE60143)/Homer | 1e-10 | -2.485e+01 | 0.0000 | 4558.0 |
| 205 | 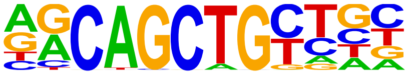 | MyoD(bHLH)/Myotube-MyoD-ChIP-Seq(GSE21614)/Homer | 1e-10 | -2.474e+01 | 0.0000 | 1765.0 |
| 206 | 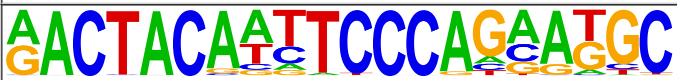 | GFY-Staf(?,Zf)/Promoter/Homer | 1e-10 | -2.466e+01 | 0.0000 | 178.0 |
| 207 | 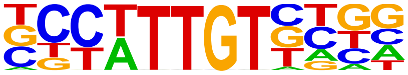 | Sox21(HMG)/ESC-SOX21-ChIP-Seq(GSE110505)/Homer | 1e-10 | -2.458e+01 | 0.0000 | 1876.0 |
| 208 | 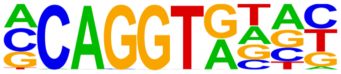 | ZEB1(Zf)/PDAC-ZEB1-ChIP-Seq(GSE64557)/Homer | 1e-10 | -2.451e+01 | 0.0000 | 2443.0 |
| 209 | 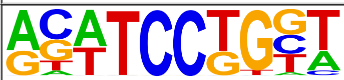 | SPDEF(ETS)/VCaP-SPDEF-ChIP-Seq(SRA014231)/Homer | 1e-10 | -2.447e+01 | 0.0000 | 1073.0 |
| 210 | 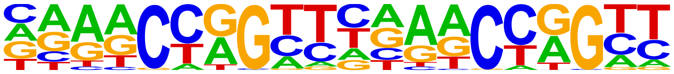 | Tcfcp2l1(CP2)/mES-Tcfcp2l1-ChIP-Seq(GSE11431)/Homer | 1e-10 | -2.381e+01 | 0.0000 | 175.0 |
| 211 | 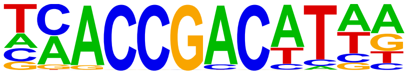 | DREB2(AP2EREBP)/col-DREB2-DAP-Seq(GSE60143)/Homer | 1e-10 | -2.367e+01 | 0.0000 | 924.0 |
| 212 | 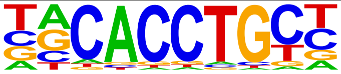 | Snail1(Zf)/LS174T-SNAIL1.HA-ChIP-Seq(GSE127183)/Homer | 1e-10 | -2.360e+01 | 0.0000 | 1365.0 |
| 213 | 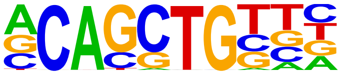 | Ptf1a(bHLH)/Panc1-Ptf1a-ChIP-Seq(GSE47459)/Homer | 1e-10 | -2.354e+01 | 0.0000 | 3871.0 |
| 214 | 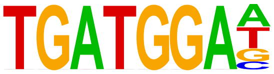 | HAP3(CCAATHAP3)/col-HAP3-DAP-Seq(GSE60143)/Homer | 1e-10 | -2.330e+01 | 0.0000 | 394.0 |
| 215 | 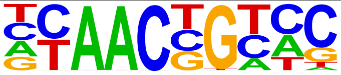 | MYB77(MYB)/col-MYB77-DAP-Seq(GSE60143)/Homer | 1e-10 | -2.303e+01 | 0.0000 | 1752.0 |
| 216 | 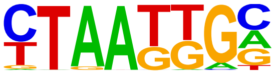 | Isl1(Homeobox)/Neuron-Isl1-ChIP-Seq(GSE31456)/Homer | 1e-9 | -2.297e+01 | 0.0000 | 1604.0 |
| 217 | 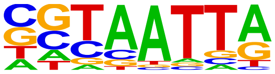 | DLX5(Homeobox)/BasalGanglia-Dlx5-ChIP-seq(GSE124936)/Homer | 1e-9 | -2.235e+01 | 0.0000 | 697.0 |
| 218 | 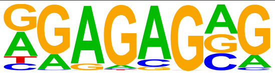 | Trl(Zf)/S2-GAGAfactor-ChIP-Seq(GSE40646)/Homer | 1e-9 | -2.212e+01 | 0.0000 | 5249.0 |
| 219 | 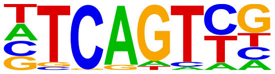 | Initiator/Drosophila-Promoters/Homer | 1e-9 | -2.211e+01 | 0.0000 | 1972.0 |

| | | | | | | |
|---|---|---|---|---|---|---|
| 220 | 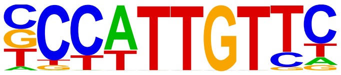 | Sox2(HMG)/mES-Sox2-ChIP-Seq(GSE11431)/Homer | 1e-9 | -2.205e+01 | 0.0000 | 945.0 |
| 221 | 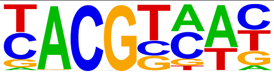 | ATAF1(NAC)/col-ATAF1-DAP-Seq(GSE60143)/Homer | 1e-9 | -2.197e+01 | 0.0000 | 4034.0 |
| 222 | 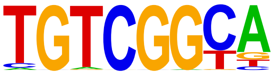 | AT1G12630(AP2EREBP)/colamp-AT1G12630-DAP-Seq(GSE60143)/Homer | 1e-9 | -2.190e+01 | 0.0000 | 1111.0 |
| 223 | 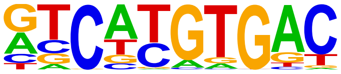 | MITF(bHLH)/MastCells-MITF-ChIP-Seq(GSE48085)/Homer | 1e-9 | -2.180e+01 | 0.0000 | 1012.0 |
| 224 | 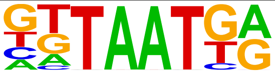 | Nkx6.1(Homeobox)/Islet-Nkx6.1-ChIP-Seq(GSE40975)/Homer | 1e-9 | -2.106e+01 | 0.0000 | 2025.0 |
| 225 | 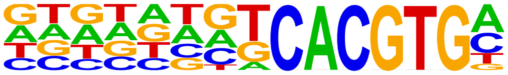 | BIM1(bHLH)/colamp-BIM1-DAP-Seq(GSE60143)/Homer | 1e-9 | -2.079e+01 | 0.0000 | 219.0 |
| 226 | 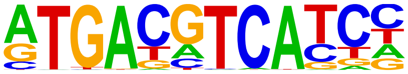 | c-Jun-CRE(bZIP)/K562-cJun-ChIP-Seq(GSE31477)/Homer | 1e-9 | -2.076e+01 | 0.0000 | 440.0 |
| 227 | 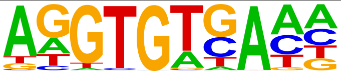 | Tbet(T-box)/CD8-Tbet-ChIP-Seq(GSE33802)/Homer | 1e-8 | -2.072e+01 | 0.0000 | 869.0 |
| 228 | 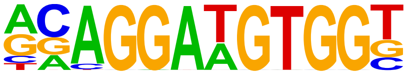 | ETS:RUNX(ETS,Runt)/Jurkat-RUNX1-ChIP-Seq(GSE17954)/Homer | 1e-8 | -2.050e+01 | 0.0000 | 150.0 |
| 229 | 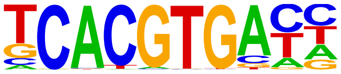 | Cbf1(bHLH)/Yeast-Cbf1-ChIP-Seq(GSE29506)/Homer | 1e-8 | -2.016e+01 | 0.0000 | 456.0 |
| 230 | 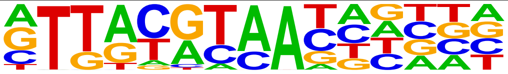 | NFIL3(bZIP)/HepG2-NFIL3-ChIP-Seq(Encode)/Homer | 1e-8 | -2.007e+01 | 0.0000 | 620.0 |
| 231 | 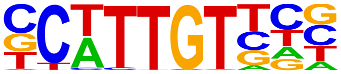 | Sox10(HMG)/SciaticNerve-Sox3-ChIP-Seq(GSE35132)/Homer | 1e-8 | -2.000e+01 | 0.0000 | 1739.0 |
| 232 | 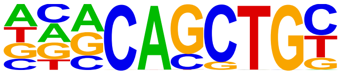 | E2A(bHLH)/proBcell-E2A-ChIP-Seq(GSE21978)/Homer | 1e-8 | -1.989e+01 | 0.0000 | 2393.0 |
| 233 | 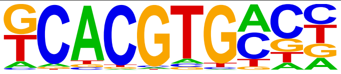 | bHLHE41(bHLH)/proB-Bhlhe41-ChIP-Seq(GSE93764)/Homer | 1e-8 | -1.986e+01 | 0.0000 | 2380.0 |
| 234 | 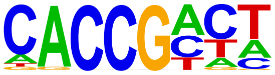 | At1g75490(AP2EREBP)/colamp-At1g75490-DAP-Seq(GSE60143)/Homer | 1e-8 | -1.956e+01 | 0.0000 | 5183.0 |
| 235 | 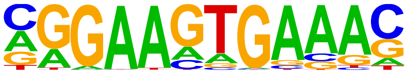 | PU.1-IRF(ETS:IRF)/Bcell-PU.1-ChIP-Seq(GSE21512)/Homer | 1e-8 | -1.944e+01 | 0.0000 | 1342.0 |
| 236 | 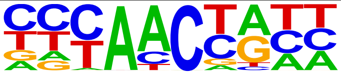 | ATY13(MYB)/col-ATY13-DAP-Seq(GSE60143)/Homer | 1e-8 | -1.934e+01 | 0.0000 | 3166.0 |
| 237 | 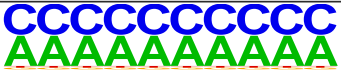 | SeqBias: C/A-bias | 1e-8 | -1.905e+01 | 0.0000 | 13419.0 |
| 238 | 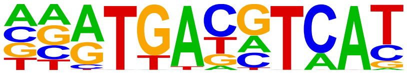 | CREB5(bZIP)/LNCaP-CREB5.V5-ChIP-Seq(GSE137775)/Homer | 1e-8 | -1.879e+01 | 0.0000 | 448.0 |
| 239 | 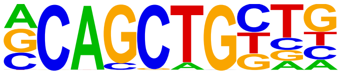 | Tcf12(bHLH)/GM12878-Tcf12-ChIP-Seq(GSE32465)/Homer | 1e-8 | -1.875e+01 | 0.0000 | 1774.0 |
| 240 | 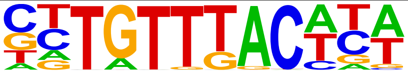 | Foxa2(Forkhead)/Liver-Foxa2-ChIP-Seq(GSE25694)/Homer | 1e-8 | -1.872e+01 | 0.0000 | 623.0 |
| 241 | 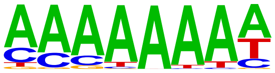 | REM19(REM)/colamp-REM19-DAP-Seq(GSE60143)/Homer | 1e-7 | -1.837e+01 | 0.0000 | 629.0 |
| 242 | 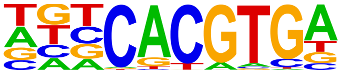 | SPCH(bHLH)/Seedling-SPCH-ChIP-Seq(GSE57497)/Homer | 1e-7 | -1.831e+01 | 0.0000 | 1518.0 |
| 243 | 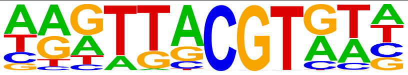 | NAP(NAC)/col-NAP-DAP-Seq(GSE60143)/Homer | 1e-7 | -1.816e+01 | 0.0000 | 962.0 |
| 244 | | At3g60580(C2H2)/col-At3g60580-DAP-Seq(GSE60143)/Homer | 1e-7 | -1.732e+01 | 0.0000 | 4075.0 |

| | | | | | | |
|---|---|---|---|---|---|---|
| | 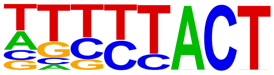 | | | | | |
| 245 | 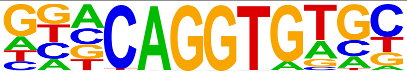 | ZEB2(Zf)/SNU398-ZEB2-ChIP-Seq(GSE103048)/Homer | 1e-7 | -1.714e+01 | 0.0000 | 1360.0 |
| 246 | 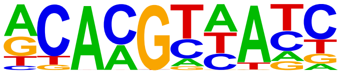 | ANAC038(NAC)/col-ANAC038-DAP-Seq(GSE60143)/Homer | 1e-7 | -1.709e+01 | 0.0000 | 1878.0 |
| 247 | 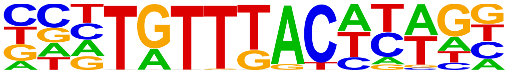 | Foxa3(Forkhead)/Liver-Foxa3-ChIP-Seq(GSE77670)/Homer | 1e-7 | -1.705e+01 | 0.0000 | 247.0 |
| 248 | 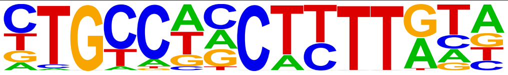 | ZNF7(Zf)/HepG2-ZNF7.Flag-ChIP-Seq(Encode)/Homer | 1e-7 | -1.692e+01 | 0.0000 | 457.0 |
| 249 | 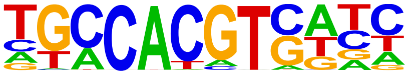 | GBF3(bZIP)/Arabidopsis-GBF3-ChIP-Seq(GSE80564)/Homer | 1e-7 | -1.691e+01 | 0.0000 | 480.0 |
| 250 | 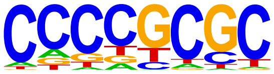 | SUT1?/SacCer-Promoters/Homer | 1e-7 | -1.682e+01 | 0.0000 | 11234.0 |
| 251 | 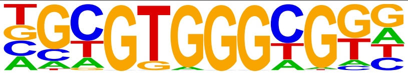 | Egr2(Zf)/Thymocytes-Egr2-ChIP-Seq(GSE34254)/Homer | 1e-7 | -1.652e+01 | 0.0000 | 744.0 |
| 252 | 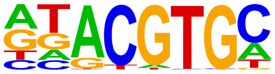 | HIF-1b(HLH)/T47D-HIF1b-ChIP-Seq(GSE59937)/Homer | 1e-7 | -1.646e+01 | 0.0000 | 2313.0 |
| 253 | 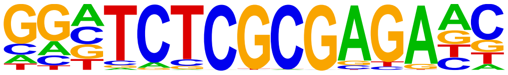 | ZBTB33(Zf)/GM12878-ZBTB33-ChIP-Seq(GSE32465)/Homer | 1e-7 | -1.640e+01 | 0.0000 | 215.0 |
| 254 | 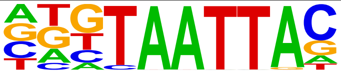 | Dlx3(Homeobox)/Kerainocytes-Dlx3-ChIP-Seq(GSE89884)/Homer | 1e-7 | -1.628e+01 | 0.0000 | 582.0 |
| 255 | 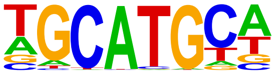 | RBFox2(?)/Heart-RBFox2-CLIP-Seq(GSE57926)/Homer | 1e-7 | -1.624e+01 | 0.0000 | 2181.0 |
| 256 | 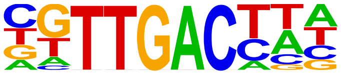 | WRKY28(WRKY)/col-WRKY28-DAP-Seq(GSE60143)/Homer | 1e-7 | -1.622e+01 | 0.0000 | 715.0 |
| 257 | 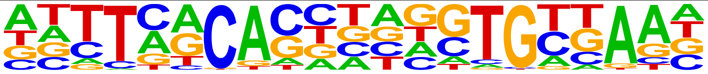 | Brachyury(T-box)/Mesoendoderm-Brachyury-ChIP-exo(GSE54963)/Homer | 1e-6 | -1.609e+01 | 0.0000 | 269.0 |
| 258 | 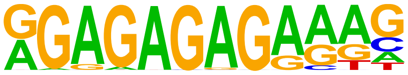 | FRS9(ND)/col-FRS9-DAP-Seq(GSE60143)/Homer | 1e-6 | -1.573e+01 | 0.0000 | 354.0 |
| 259 | 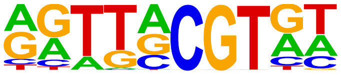 | NAM(NAC)/col-NAM-DAP-Seq(GSE60143)/Homer | 1e-6 | -1.571e+01 | 0.0000 | 1598.0 |
| 260 | 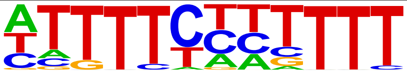 | RLR1?/SacCer-Promoters/Homer | 1e-6 | -1.548e+01 | 0.0000 | 516.0 |
| 261 | 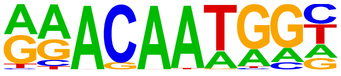 | Sox15(HMG)/CPA-Sox15-ChIP-Seq(GSE62909)/Homer | 1e-6 | -1.504e+01 | 0.0000 | 951.0 |
| 262 | 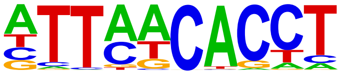 | Eomes(T-box)/H9-Eomes-ChIP-Seq(GSE26097)/Homer | 1e-6 | -1.497e+01 | 0.0000 | 1748.0 |
| 263 | 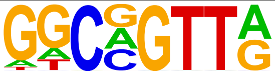 | MYB(HTH)/ERMYB-Myb-ChIPSeq(GSE22095)/Homer | 1e-6 | -1.470e+01 | 0.0000 | 2619.0 |
| 264 | 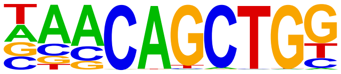 | Tcf21(bHLH)/ArterySmoothMuscle-Tcf21-ChIP-Seq(GSE61369)/Homer | 1e-6 | -1.450e+01 | 0.0000 | 1499.0 |
| 265 | 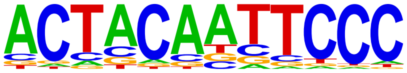 | GFY(?)/Promoter/Homer | 1e-6 | -1.436e+01 | 0.0000 | 154.0 |
| 266 | 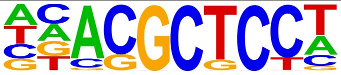 | MYB88(MYB)/col-MYB88-DAP-Seq(GSE60143)/Homer | 1e-6 | -1.412e+01 | 0.0000 | 6533.0 |
| 267 | 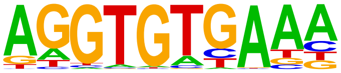 | Tbx21(T-box)/GM12878-TBX21-ChIP-Seq(Encode)/Homer | 1e-6 | -1.410e+01 | 0.0000 | 792.0 |
| 268 | 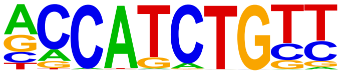 | NeuroG2(bHLH)/Fibroblast-NeuroG2-ChIP-Seq(GSE75910)/Homer | 1e-6 | -1.407e+01 | 0.0000 | 1771.0 |

| | | | | | | |
|---|---|---|---|---|---|---|
| 269 | 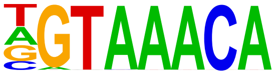 | Foxo3(Forkhead)/U2OS-Foxo3-ChIP-Seq(E-MTAB-2701)/Homer | 1e-6 | -1.390e+01 | 0.0000 | 554.0 |
| 270 | 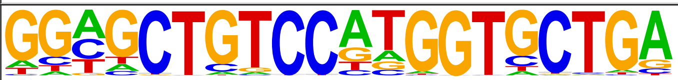 | REST-NRSF(Zf)/Jurkat-NRSF-ChIP-Seq/Homer | 1e-5 | -1.364e+01 | 0.0000 | 47.0 |
| 271 | 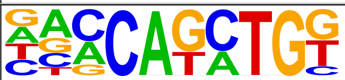 | BHLHA15(bHLH)/NIH3T3-BHLHB8.HA-ChIP-Seq(GSE119782)/Homer | 1e-5 | -1.362e+01 | 0.0000 | 1730.0 |
| 272 | 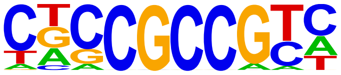 | LEP(AP2EREBP)/col-LEP-DAP-Seq(GSE60143)/Homer | 1e-5 | -1.338e+01 | 0.0000 | 3849.0 |
| 273 | 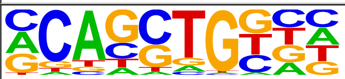 | HEB(bHLH)/mES-Heb-ChIP-Seq(GSE53233)/Homer | 1e-5 | -1.312e+01 | 0.0000 | 2902.0 |
| 274 | 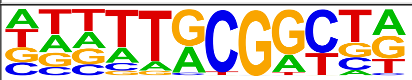 | At5g08750(C3H)/col-At5g08750-DAP-Seq(GSE60143)/Homer | 1e-5 | -1.299e+01 | 0.0000 | 845.0 |
| 275 | 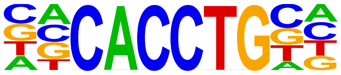 | E2A(bHLH),near_PU.1/Bcell-PU.1-ChIP-Seq(GSE21512)/Homer | 1e-5 | -1.296e+01 | 0.0000 | 1994.0 |
| 276 | 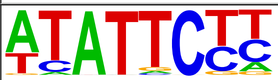 | KAN2(G2like)/colamp-KAN2-DAP-Seq(GSE60143)/Homer | 1e-5 | -1.246e+01 | 0.0000 | 882.0 |
| 277 | 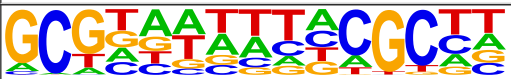 | LBD23(LOBAS2)/colamp-LBD23-DAP-Seq(GSE60143)/Homer | 1e-5 | -1.193e+01 | 0.0000 | 2887.0 |
| 278 | 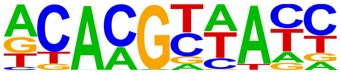 | ANAC046(NAC)/colamp-ANAC046-DAP-Seq(GSE60143)/Homer | 1e-5 | -1.188e+01 | 0.0000 | 1680.0 |
| 279 | 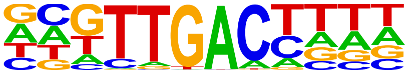 | WRKY27(WRKY)/colamp-WRKY27-DAP-Seq(GSE60143)/Homer | 1e-5 | -1.169e+01 | 0.0000 | 475.0 |
| 280 | 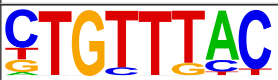 | Foxo1(Forkhead)/RAW-Foxo1-ChIP-Seq(Fan_et_al.)/Homer | 1e-5 | -1.167e+01 | 0.0000 | 1669.0 |
| 281 | 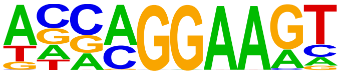 | EHF(ETS)/LoVo-EHF-ChIP-Seq(GSE49402)/Homer | 1e-5 | -1.160e+01 | 0.0000 | 1287.0 |
| 282 | 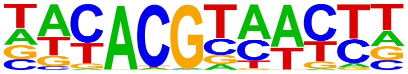 | ANAC047(NAC)/colamp-ANAC047-DAP-Seq(GSE60143)/Homer | 1e-4 | -1.150e+01 | 0.0000 | 683.0 |
| 283 | 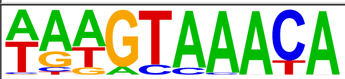 | FOXA1(Forkhead)/LNCAP-FOXA1-ChIP-Seq(GSE27824)/Homer | 1e-4 | -1.134e+01 | 0.0000 | 853.0 |
| 284 | 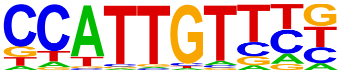 | Sox17(HMG)/Endoderm-Sox17-ChIP-Seq(GSE61475)/Homer | 1e-4 | -1.111e+01 | 0.0001 | 616.0 |
| 285 | 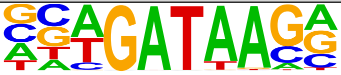 | Gata4(Zf)/Heart-Gata4-ChIP-Seq(GSE35151)/Homer | 1e-4 | -1.102e+01 | 0.0001 | 703.0 |
| 286 | 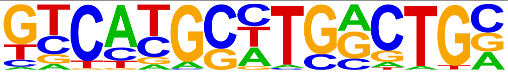 | Pax8(Paired,Homeobox)/Thyroid-Pax8-ChIP-Seq(GSE26938)/Homer | 1e-4 | -1.091e+01 | 0.0001 | 420.0 |
| 287 | 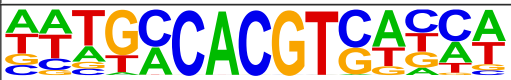 | GBF6(bZIP)/colamp-GBF6-DAP-Seq(GSE60143)/Homer | 1e-4 | -1.083e+01 | 0.0001 | 255.0 |
| 288 | 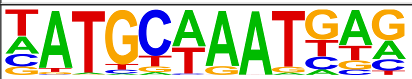 | Oct6(POU,Homeobox)/NPC-Pou3f1-ChIP-Seq(GSE35496)/Homer | 1e-4 | -1.076e+01 | 0.0001 | 280.0 |
| 289 | 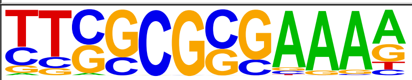 | E2F(E2F)/Hela-CellCycle-Expression/Homer | 1e-4 | -1.073e+01 | 0.0001 | 154.0 |
| 290 | 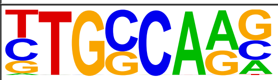 | NF1-halfsite(CTF)/LNCaP-NF1-ChIP-Seq(Unpublished)/Homer | 1e-4 | -1.067e+01 | 0.0001 | 2003.0 |
| 291 | 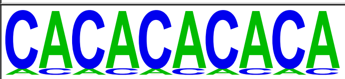 | SeqBias: CA-repeat | 1e-4 | -1.051e+01 | 0.0001 | 5835.0 |
| 292 | 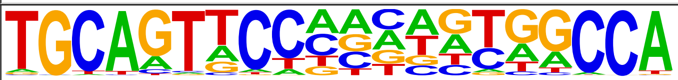 | CTCF-SatelliteElement(Zf?)/CD4+-CTCF-ChIP-Seq(Barski_et_al.)/Homer | 1e-4 | -1.023e+01 | 0.0001 | 24.0 |
| 293 | 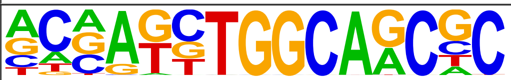 | Unknown3/Drosophila-Promoters/Homer | 1e-4 | -9.783e+00 | 0.0002 | 215.0 |

| | | | | | | |
|---|---|---|---|---|---|---|
| 294 | 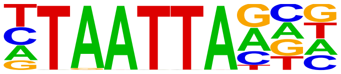 | ATHB23(ZFHD)/col-ATHB23-DAP-Seq(GSE60143)/Homer | 1e-4 | -9.693e+00 | 0.0002 | 632.0 |
| 295 | 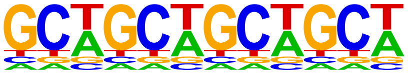 | SeqBias: GCW-triplet | 1e-4 | -9.607e+00 | 0.0002 | 13430.0 |
| 296 | 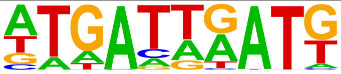 | LIN-39(Homeobox)/cElegans.L3-LIN39-ChIP-Seq(modEncode)/Homer | 1e-4 | -9.579e+00 | 0.0002 | 827.0 |
| 297 | 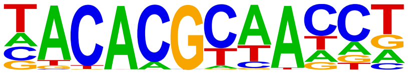 | ANAC079(NAC)/colamp-ANAC079-DAP-Seq(GSE60143)/Homer | 1e-4 | -9.567e+00 | 0.0002 | 321.0 |
| 298 | 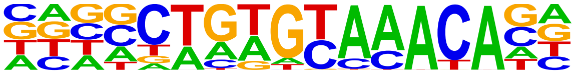 | Fox:Ebox(Forkhead,bHLH)/Panc1-Foxa2-ChIP-Seq(GSE47459)/Homer | 1e-4 | -9.547e+00 | 0.0002 | 789.0 |
| 299 | 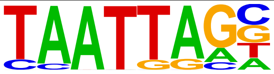 | Lhx2(Homeobox)/HFSC-Lhx2-ChIP-Seq(GSE48068)/Homer | 1e-4 | -9.496e+00 | 0.0003 | 817.0 |
| 300 | 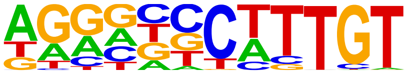 | Sox9(HMG)/Limb-SOX9-ChIP-Seq(GSE73225)/Homer | 1e-4 | -9.451e+00 | 0.0003 | 730.0 |
| 301 | 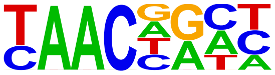 | MYB101(MYB)/colamp-MYB101-DAP-Seq(GSE60143)/Homer | 1e-3 | -9.208e+00 | 0.0003 | 2087.0 |
| 302 | 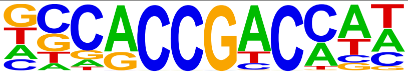 | At4g28140(AP2EREBP)/colamp-At4g28140-DAP-Seq(GSE60143)/Homer | 1e-3 | -9.130e+00 | 0.0004 | 898.0 |
| 303 | 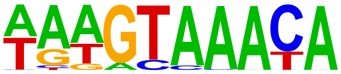 | FOXA1(Forkhead)/MCF7-FOXA1-ChIP-Seq(GSE26831)/Homer | 1e-3 | -9.021e+00 | 0.0004 | 683.0 |
| 304 | 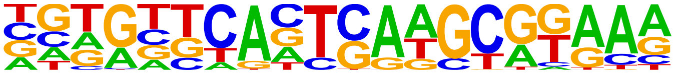 | PAX6(Paired,Homeobox)/Forebrain-Pax6-ChIP-Seq(GSE66961)/Homer | 1e-3 | -9.017e+00 | 0.0004 | 97.0 |
| 305 | 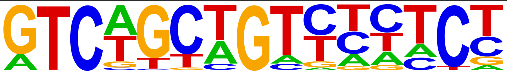 | ZNF317(Zf)/HEK293-ZNF317.GFP-ChIP-Seq(GSE58341)/Homer | 1e-3 | -8.928e+00 | 0.0004 | 104.0 |
| 306 | 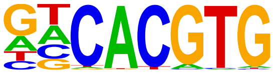 | CLOCK(bHLH)/Liver-Clock-ChIP-Seq(GSE39860)/Homer | 1e-3 | -8.896e+00 | 0.0005 | 807.0 |
| 307 | 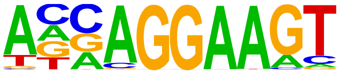 | ELF3(ETS)/PDAC-ELF3-ChIP-Seq(GSE64557)/Homer | 1e-3 | -8.799e+00 | 0.0005 | 627.0 |
| 308 | 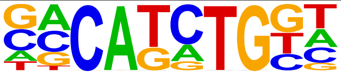 | TCF4(bHLH)/SHSY5Y-TCF4-ChIP-Seq(GSE96915)/Homer | 1e-3 | -8.655e+00 | 0.0006 | 1710.0 |
| 309 | 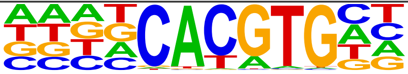 | BIM2(bHLH)/col-BIM2-DAP-Seq(GSE60143)/Homer | 1e-3 | -8.401e+00 | 0.0007 | 990.0 |
| 310 | 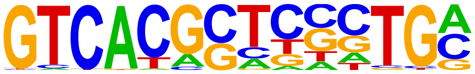 | PAX5(Paired,Homeobox),condensed/GM12878-PAX5-ChIP-Seq(GSE32465)/Homer | 1e-3 | -8.368e+00 | 0.0008 | 175.0 |
| 311 | 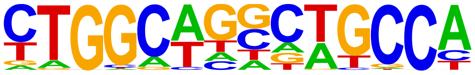 | Tlx?(NR)/NPC-H3K4me1-ChIP-Seq(GSE16256)/Homer | 1e-3 | -8.344e+00 | 0.0008 | 398.0 |
| 312 | 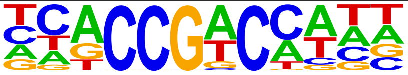 | At1g19210(AP2EREBP)/colamp-At1g19210-DAP-Seq(GSE60143)/Homer | 1e-3 | -8.343e+00 | 0.0008 | 2691.0 |
| 313 | 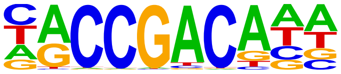 | At4g16750(AP2EREBP)/col-At4g16750-DAP-Seq(GSE60143)/Homer | 1e-3 | -8.103e+00 | 0.0010 | 1342.0 |
| 314 | 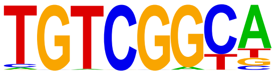 | Rap210(AP2EREBP)/col-Rap210-DAP-Seq(GSE60143)/Homer | 1e-3 | -8.090e+00 | 0.0010 | 1207.0 |
| 315 | 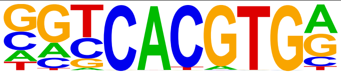 | USF1(bHLH)/GM12878-Usf1-ChIP-Seq(GSE32465)/Homer | 1e-3 | -8.074e+00 | 0.0010 | 711.0 |
| 316 | 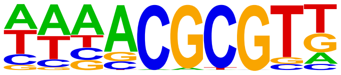 | CAMTA1(CAMTA)/col-CAMTA1-DAP-Seq(GSE60143)/Homer | 1e-3 | -8.061e+00 | 0.0010 | 1275.0 |
| 317 | 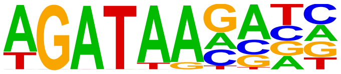 | TRPS1(Zf)/MCF7-TRPS1-ChIP-Seq(GSE107013)/Homer | 1e-3 | -7.945e+00 | 0.0011 | 1267.0 |
| 318 | 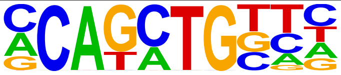 | Twist2(bHLH)/Myoblast-Twist2.Ty1-ChIP-Seq(GSE127998)/Homer | 1e-3 | -7.904e+00 | 0.0012 | 2159.0 |

| | | | | | | |
|---|---|---|---|---|---|---|
| 319 | 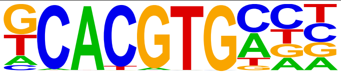 | bHLHE40(bHLH)/HepG2-BHLHE40-ChIP-Seq(GSE31477)/Homer | 1e-3 | -7.806e+00 | 0.0013 | 580.0 |
| 320 | 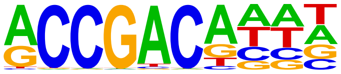 | AT1G77200(AP2EREBP)/colamp-AT1G77200-DAP-Seq(GSE60143)/Homer | 1e-3 | -7.713e+00 | 0.0014 | 1356.0 |
| 321 | 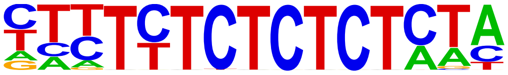 | BPC6(BBRBPC)/col-BPC6-DAP-Seq(GSE60143)/Homer | 1e-3 | -7.578e+00 | 0.0016 | 47.0 |
| 322 | 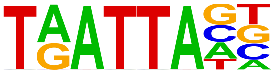 | ATHB25(ZFHD)/colamp-ATHB25-DAP-Seq(GSE60143)/Homer | 1e-3 | -7.556e+00 | 0.0016 | 759.0 |
| 323 | 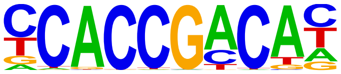 | DREB26(AP2EREBP)/col-DREB26-DAP-Seq(GSE60143)/Homer | 1e-3 | -7.556e+00 | 0.0016 | 759.0 |
| 324 | 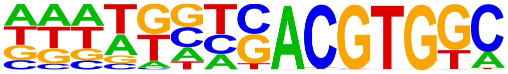 | bZIP48(bZIP)/colamp-bZIP48-DAP-Seq(GSE60143)/Homer | 1e-3 | -7.533e+00 | 0.0017 | 362.0 |
| 325 | 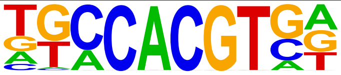 | bZIP16(bZIP)/colamp-bZIP16-DAP-Seq(GSE60143)/Homer | 1e-3 | -7.468e+00 | 0.0018 | 391.0 |
| 326 | 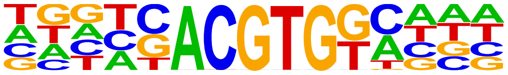 | bZIP53(bZIP)/colamp-bZIP53-DAP-Seq(GSE60143)/Homer | 1e-3 | -7.462e+00 | 0.0018 | 411.0 |
| 327 | 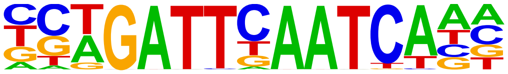 | DUX(Homeobox)/C2C12-Dux-ChIP-Seq(GSE87279)/Homer | 1e-3 | -7.456e+00 | 0.0018 | 4.0 |
| 328 | 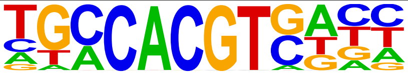 | bZIP28(bZIP)/col-bZIP28-DAP-Seq(GSE60143)/Homer | 1e-3 | -7.330e+00 | 0.0020 | 373.0 |
| 329 | 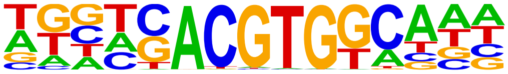 | GBF5(bZIP)/colamp-GBF5-DAP-Seq(GSE60143)/Homer | 1e-3 | -7.289e+00 | 0.0021 | 341.0 |
| 330 | 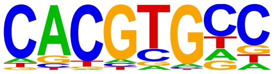 | IBL1(bHLH)/Seedling-IBL1-ChIP-Seq(GSE51120)/Homer | 1e-3 | -7.254e+00 | 0.0022 | 4060.0 |
| 331 | 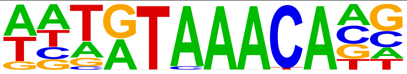 | FoxL2(Forkhead)/Ovary-FoxL2-ChIP-Seq(GSE60858)/Homer | 1e-3 | -7.204e+00 | 0.0023 | 554.0 |
| 332 | 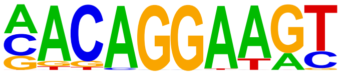 | Ets1-distal(ETS)/CD4+-PolII-ChIP-Seq(Barski_et_al.)/Homer | 1e-3 | -7.153e+00 | 0.0024 | 226.0 |
| 333 | 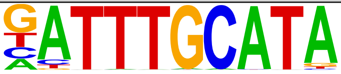 | Oct11(POU,Homeobox)/NCIH1048-POU2F3-ChIP-seq(GSE115123)/Homer | 1e-3 | -7.043e+00 | 0.0026 | 214.0 |
| 334 | 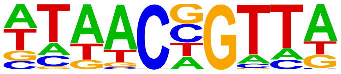 | MYB70(MYB)/col-MYB70-DAP-Seq(GSE60143)/Homer | 1e-3 | -6.926e+00 | 0.0030 | 1137.0 |
| 335 | 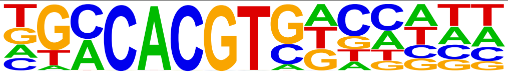 | ABF2(bZIP)/col-ABF2-DAP-Seq(GSE60143)/Homer | 1e-2 | -6.905e+00 | 0.0030 | 291.0 |
| 336 | 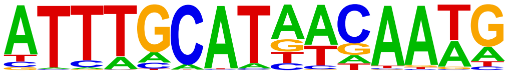 | OCT4-SOX2-TCF-NANOG(POU,Homeobox,HMG)/mES-Oct4-ChIP-Seq(GSE11431)/Homer | 1e-2 | -6.898e+00 | 0.0030 | 110.0 |
| 337 | 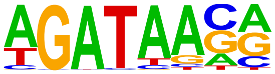 | GATA3(Zf)/iTreg-Gata3-ChIP-Seq(GSE20898)/Homer | 1e-2 | -6.765e+00 | 0.0034 | 950.0 |
| 338 | 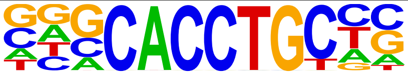 | Slug(Zf)/Mesoderm-Snai2-ChIP-Seq(GSE61475)/Homer | 1e-2 | -6.661e+00 | 0.0038 | 869.0 |
| 339 | 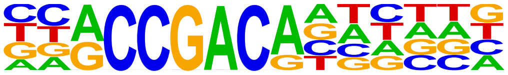 | AT3G60490(AP2EREBP)/colamp-AT3G60490-DAP-Seq(GSE60143)/Homer | 1e-2 | -6.619e+00 | 0.0040 | 587.0 |
| 340 | 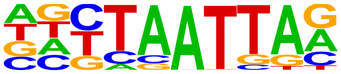 | Lhx1(Homeobox)/EmbryoCarcinoma-Lhx1-ChIP-Seq(GSE70957)/Homer | 1e-2 | -6.607e+00 | 0.0040 | 848.0 |
| 341 | 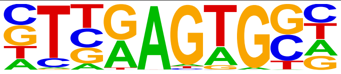 | Nkx2.2(Homeobox)/NPC-Nkx2.2-ChIP-Seq(GSE61673)/Homer | 1e-2 | -6.574e+00 | 0.0041 | 1712.0 |
| 342 | 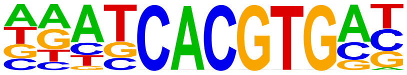 | bHLH74(bHLH)/col-bHLH74-DAP-Seq(GSE60143)/Homer | 1e-2 | -6.564e+00 | 0.0041 | 493.0 |
| 343 | | bZIP44(bZIP)/colamp-bZIP44-DAP-Seq(GSE60143)/Homer | 1e-2 | -6.554e+00 | 0.0042 | 34.0 |

| | | | | | | |
|---|---|---|---|---|---|---|
| | 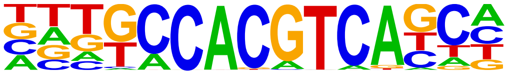 | | | | | |
| 344 | 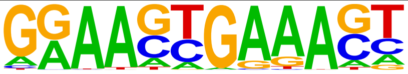 | IRF8(IRF)/BMDM-IRF8-ChIP-Seq(GSE77884)/Homer | 1e-2 | -6.542e+00 | 0.0042 | 288.0 |
| 345 | 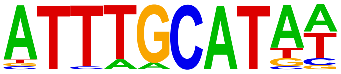 | Oct4(POU,Homeobox)/mES-Oct4-ChIP-Seq(GSE11431)/Homer | 1e-2 | -6.476e+00 | 0.0045 | 305.0 |
| 346 | 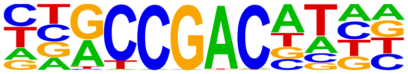 | CBF3(AP2EREBP)/colamp-CBF3-DAP-Seq(GSE60143)/Homer | 1e-2 | -6.419e+00 | 0.0047 | 777.0 |
| 347 | 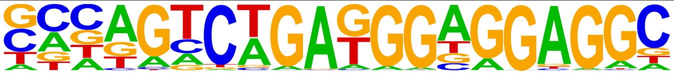 | ZSCAN22(Zf)/HEK293-ZSCAN22.GFP-ChIP-Seq(GSE58341)/Homer | 1e-2 | -6.372e+00 | 0.0050 | 139.0 |
| 348 | 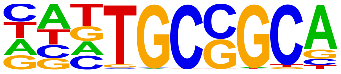 | Zfp57(Zf)/H1-ZFP57.HA-ChIP-Seq(GSE115387)/Homer | 1e-2 | -6.325e+00 | 0.0052 | 2135.0 |
| 349 | 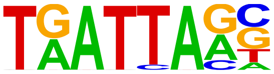 | ATHB34(ZFHD)/colamp-ATHB34-DAP-Seq(GSE60143)/Homer | 1e-2 | -6.116e+00 | 0.0064 | 545.0 |
| 350 | 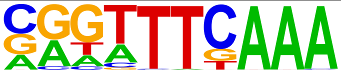 | CHR(?)/Hela-CellCycle-Expression/Homer | 1e-2 | -6.082e+00 | 0.0066 | 418.0 |
| 351 | 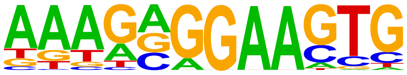 | SpiB(ETS)/OCILY3-SPIB-ChIP-Seq(GSE56857)/Homer | 1e-2 | -6.077e+00 | 0.0066 | 227.0 |
| 352 | 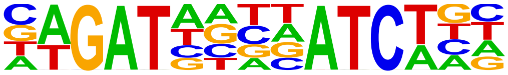 | GATA(Zf),IR4/iTreg-Gata3-ChIP-Seq(GSE20898)/Homer | 1e-2 | -6.072e+00 | 0.0066 | 67.0 |
| 353 | 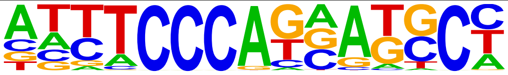 | ZNF143\|STAF(Zf)/CUTLL-ZNF143-ChIP-Seq(GSE29600)/Homer | 1e-2 | -5.982e+00 | 0.0072 | 322.0 |
| 354 | 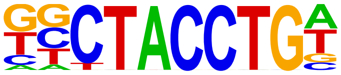 | Zelda(Zf)/Embryo-zld-ChIP-Seq(GSE65441)/Homer | 1e-2 | -5.965e+00 | 0.0073 | 540.0 |
| 355 | 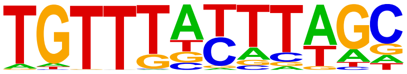 | FoxD3(forkhead)/ZebrafishEmbryo-Foxd3.biotin-ChIP-seq(GSE106676)/Homer | 1e-2 | -5.958e+00 | 0.0073 | 622.0 |
| 356 | 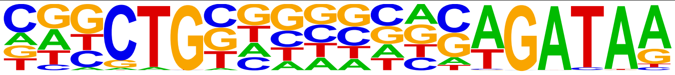 | GATA:SCL(Zf,bHLH)/Ter119-SCL-ChIP-Seq(GSE18720)/Homer | 1e-2 | -5.872e+00 | 0.0080 | 106.0 |
| 357 | 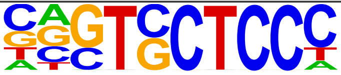 | Znf263(Zf)/K562-Znf263-ChIP-Seq(GSE31477)/Homer | 1e-2 | -5.836e+00 | 0.0082 | 3646.0 |
| 358 | 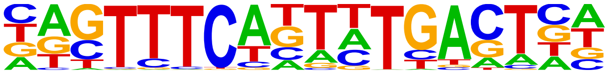 | bZIP:IRF(bZIP,IRF)/Th17-BatF-ChIP-Seq(GSE39756)/Homer | 1e-2 | -5.797e+00 | 0.0085 | 254.0 |
| 359 | 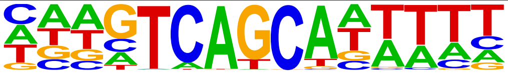 | MafF(bZIP)/HepG2-MafF-ChIP-Seq(GSE31477)/Homer | 1e-2 | -5.796e+00 | 0.0085 | 220.0 |
| 360 | 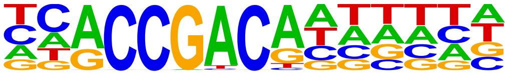 | AT3G16280(AP2EREBP)/colamp-AT3G16280-DAP-Seq(GSE60143)/Homer | 1e-2 | -5.750e+00 | 0.0089 | 597.0 |
| 361 | 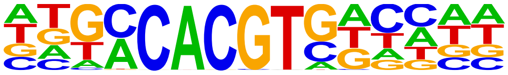 | AREB3(bZIP)/col-AREB3-DAP-Seq(GSE60143)/Homer | 1e-2 | -5.682e+00 | 0.0095 | 387.0 |
| 362 | 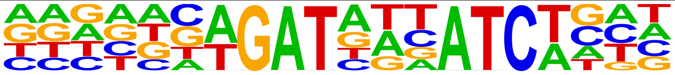 | GATA(Zf),IR3/iTreg-Gata3-ChIP-Seq(GSE20898)/Homer | 1e-2 | -5.655e+00 | 0.0097 | 107.0 |
| 363 | 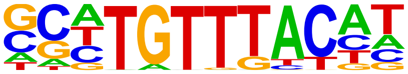 | FOXK2(Forkhead)/U2OS-FOXK2-ChIP-Seq(E-MTAB-2204)/Homer | 1e-2 | -5.635e+00 | 0.0099 | 460.0 |
| 364 | 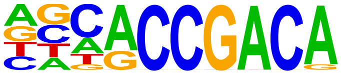 | TINY(AP2EREBP)/col-TINY-DAP-Seq(GSE60143)/Homer | 1e-2 | -5.618e+00 | 0.0100 | 548.0 |
| 365 | 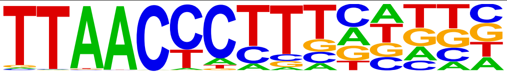 | ZNF652/HepG2-ZNF652.Flag-ChIP-Seq(Encode)/Homer | 1e-2 | -5.585e+00 | 0.0103 | 184.0 |
| 366 | 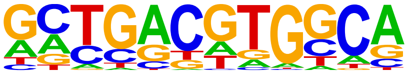 | O2(bZIP)/Corn-O2-ChIP-Seq(GSE63991)/Homer | 1e-2 | -5.514e+00 | 0.0111 | 263.0 |
| 367 | | IRF4(IRF)/GM12878-IRF4-ChIP- | 1e-2 | -5.476e+00 | 0.0115 | 337.0 |

| | | | | | | |
|---|---|---|---|---|---|---|
| | 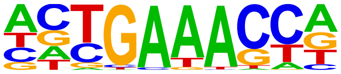 | Seq(GSE32465)/Homer | | | | |
| 368 | 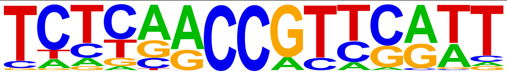 | AT5G59990(C2C2COlike)/colamp-AT5G59990-DAP-Seq(GSE60143)/Homer | 1e-2 | -5.425e+00 | 0.0120 | 34.0 |
| 369 | 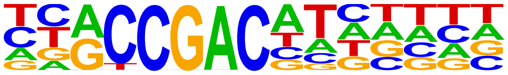 | CBF2(AP2EREBP)/colamp-CBF2-DAP-Seq(GSE60143)/Homer | 1e-2 | -5.420e+00 | 0.0121 | 722.0 |
| 370 | 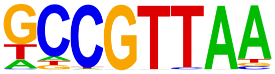 | AT3G10030(Trihelix)/colamp-AT3G10030-DAP-Seq(GSE60143)/Homer | 1e-2 | -5.364e+00 | 0.0127 | 713.0 |
| 371 | 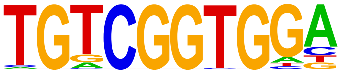 | At1g77640(AP2EREBP)/col-At1g77640-DAP-Seq(GSE60143)/Homer | 1e-2 | -5.357e+00 | 0.0128 | 451.0 |
| 372 | 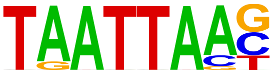 | ATHB24(ZFHD)/colamp-ATHB24-DAP-Seq(GSE60143)/Homer | 1e-2 | -5.347e+00 | 0.0129 | 503.0 |
| 373 | 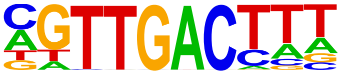 | WRKY29(WRKY)/colamp-WRKY29-DAP-Seq(GSE60143)/Homer | 1e-2 | -5.324e+00 | 0.0131 | 504.0 |
| 374 | 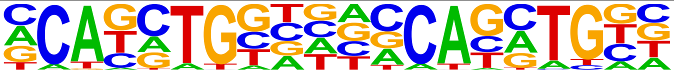 | Twist(bHLH)/HMLE-TWIST1-ChIP-Seq(Chang_et_al)/Homer | 1e-2 | -5.255e+00 | 0.0141 | 158.0 |
| 375 | 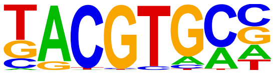 | HIF-1a(bHLH)/MCF7-HIF1a-ChIP-Seq(GSE28352)/Homer | 1e-2 | -5.238e+00 | 0.0142 | 584.0 |
| 376 | 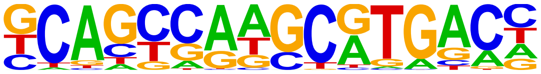 | PAX5(Paired,Homeobox)/GM12878-PAX5-ChIP-Seq(GSE32465)/Homer | 1e-2 | -5.124e+00 | 0.0159 | 690.0 |
| 377 | 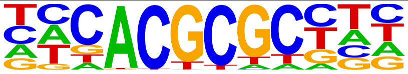 | FHY3(FAR1)/Arabidopsis-FHY3-ChIP-Seq(GSE30711)/Homer | 1e-2 | -5.104e+00 | 0.0162 | 1035.0 |
| 378 | 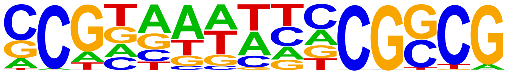 | AS2(LOBAS2)/col-AS2-DAP-Seq(GSE60143)/Homer | 1e-2 | -5.077e+00 | 0.0166 | 592.0 |
| 379 | 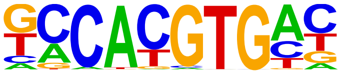 | NPAS2(bHLH)/Liver-NPAS2-ChIP-Seq(GSE39860)/Homer | 1e-2 | -5.043e+00 | 0.0171 | 1250.0 |
| 380 | 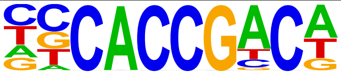 | AT1G44830(AP2EREBP)/col-AT1G44830-DAP-Seq(GSE60143)/Homer | 1e-2 | -5.040e+00 | 0.0171 | 924.0 |
| 381 | 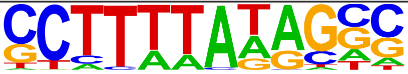 | TATA-Box(TBP)/Promoter/Homer | 1e-2 | -4.952e+00 | 0.0187 | 966.0 |
| 382 | 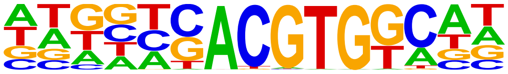 | bZIP3(bZIP)/col-bZIP3-DAP-Seq(GSE60143)/Homer | 1e-2 | -4.949e+00 | 0.0187 | 514.0 |
| 383 | 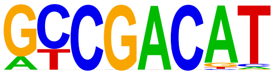 | DDF1(AP2EREBP)/col-DDF1-DAP-Seq(GSE60143)/Homer | 1e-2 | -4.931e+00 | 0.0190 | 633.0 |
| 384 | 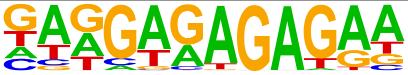 | BPC1(BBRBPC)/colamp-BPC1-DAP-Seq(GSE60143)/Homer | 1e-2 | -4.887e+00 | 0.0198 | 503.0 |
| 385 | 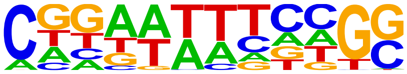 | LBD18(LOBAS2)/colamp-LBD18-DAP-Seq(GSE60143)/Homer | 1e-2 | -4.843e+00 | 0.0206 | 2302.0 |
| 386 | 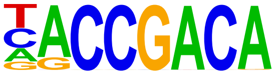 | bHLH10(bHLH)/colamp-bHLH10-DAP-Seq(GSE60143)/Homer | 1e-2 | -4.822e+00 | 0.0210 | 579.0 |
| 387 | 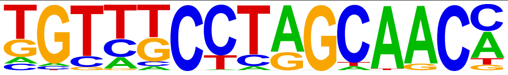 | Rfx6(HTH)/Min6b1-Rfx6.HA-ChIP-Seq(GSE62844)/Homer | 1e-2 | -4.820e+00 | 0.0210 | 1517.0 |
| 388 | 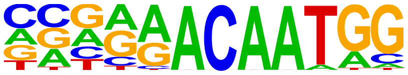 | Sox7(HMG)/ESC-Sox7-ChIP-Seq(GSE133899)/Homer | 1e-2 | -4.733e+00 | 0.0228 | 275.0 |
| 389 | 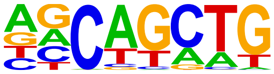 | SCL(bHLH)/HPC7-Scl-ChIP-Seq(GSE13511)/Homer | 1e-2 | -4.643e+00 | 0.0249 | 6486.0 |

S4 Table. Full list of de novo motif discovery from ChIP-seq data targeting PBX1.

### Homer *de novo* Motif Results (/wynton/group/marcucio/2022CHM/Data2022Mar/Motif/HomerPBX1IDR/)

Known Motif Enrichment Results
Gene Ontology Enrichment Results
If Homer is having trouble matching a motif to a known motif, try copy/pasting the matrix file into STAMP
More information on motif finding results: HOMER | Description of Results | Tips
Total target sequences = 13431
Total background sequences = 36373
* - possible false positive

| Rank | Motif | P-value | log P-pvalue | % of Targets | % of Background | STD(Bg STD) | Best Match/Details | Motif File |
|---|---|---|---|---|---|---|---|---|
| 1 | 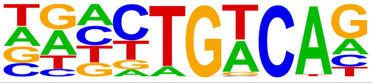 | 1e-823 | -1.896e+03 | 29.90% | 10.54% | 49.0bp (62.6bp) | Tgif1(Homeobox)/mES-Tgif1-ChIP-Seq(GSE55404)/Homer(0.955) More Information \| Similar Motifs Found | motif file (matrix) |
| 2 | 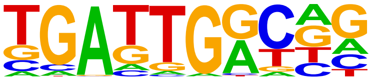 | 1e-480 | -1.107e+03 | 18.00% | 6.18% | 53.0bp (64.4bp) | Pknox1(Homeobox)/ES-Prep1-ChIP-Seq(GSE63282)/Homer(0.903) More Information \| Similar Motifs Found | motif file (matrix) |
| 3 | 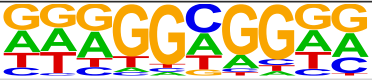 | 1e-158 | -3.646e+02 | 44.58% | 33.40% | 54.8bp (62.0bp) | POL003.1_GC-box/Jaspar(0.867) More Information \| Similar Motifs Found | motif file (matrix) |
| 4 | 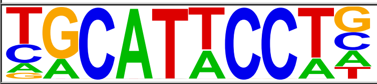 | 1e-121 | -2.793e+02 | 4.85% | 1.67% | 55.2bp (59.7bp) | TEAD3/MA0808.1/Jaspar(0.930) More Information \| Similar Motifs Found | motif file (matrix) |
| 5 | 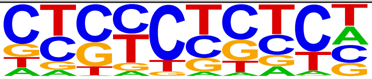 | 1e-105 | -2.425e+02 | 42.83% | 33.75% | 55.9bp (62.4bp) | SeqBias: GA-repeat(0.797) More Information \| Similar Motifs Found | motif file (matrix) |
| 6 | 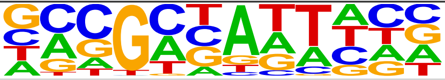 | 1e-86 | -1.998e+02 | 42.19% | 33.96% | 55.5bp (63.5bp) | pho/dmmpmm(Bergman)/fly(0.716) More Information \| Similar Motifs Found | motif file (matrix) |
| 7 | 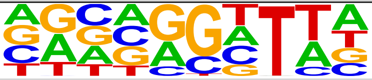 | 1e-82 | -1.905e+02 | 18.13% | 12.32% | 56.8bp (61.4bp) | bcd/dmmpmm(Bigfoot)/fly(0.794) More Information \| Similar Motifs Found | motif file (matrix) |
| 8 | 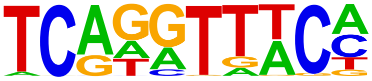 | 1e-75 | -1.742e+02 | 4.10% | 1.68% | 55.3bp (61.2bp) | Six1(Homeobox)/Myoblast-Six1-ChIP-Chip(GSE20150)/Homer(0.935) More Information \| Similar Motifs Found | motif file (matrix) |
| 9 | 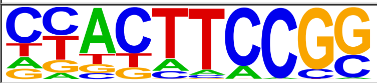 | 1e-65 | -1.512e+02 | 8.83% | 5.22% | 53.0bp (59.3bp) | ELK4/MA0076.2/Jaspar(0.949) More Information \| Similar Motifs Found | motif file (matrix) |
| 10 | 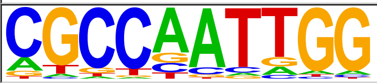 | 1e-65 | -1.497e+02 | 5.36% | 2.67% | 54.0bp (64.5bp) | NFY(CCAAT)/Promoter/Homer(0.766) More Information \| Similar Motifs Found | motif file (matrix) |
| 11 | 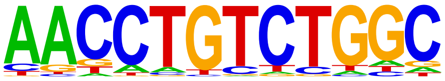 | 1e-50 | -1.158e+02 | 1.33% | 0.34% | 53.3bp (53.7bp) | Smad2(MAD)/ES-SMAD2-ChIP-Seq(GSE29422)/Homer(0.772) More Information \| Similar Motifs Found | motif file (matrix) |
| 12 | 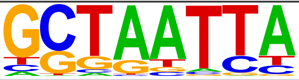 | 1e-48 | -1.106e+02 | 21.64% | 16.75% | 57.5bp (62.1bp) | Ro/dmmpmm(Noyes_hd)/fly(0.910) More Information \| Similar Motifs Found | motif file (matrix) |
| 13 | 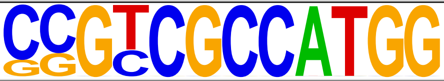 | 1e-45 | -1.039e+02 | 10.92% | 7.50% | 52.0bp (60.4bp) | YY2/MA0748.2/Jaspar(0.799) More Information \| Similar Motifs Found | motif file (matrix) |
| 14 | 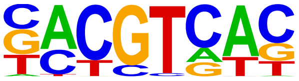 | 1e-43 | -9.922e+01 | 10.28% | 7.03% | 55.0bp (64.2bp) | TGA9(bZIP)/colamp-TGA9-DAP-Seq(GSE60143)/Homer(0.908) More Information \| Similar Motifs Found | motif file (matrix) |
| 15 | 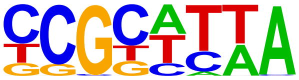 | 1e-41 | -9.549e+01 | 21.90% | 17.33% | 55.9bp (65.0bp) | Unknown3/Arabidopsis-Promoters/Homer(0.702) More Information \| Similar Motifs Found | motif file (matrix) |
| 16 | 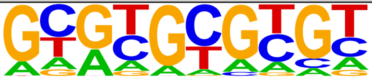 | 1e-30 | -7.091e+01 | 13.58% | 10.39% | 55.7bp (67.4bp) | TCFL5/MA0632.2/Jaspar(0.775) More Information \| Similar Motifs Found | motif file (matrix) |
| 17 | 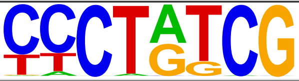 | 1e-29 | -6.742e+01 | 8.82% | 6.31% | 57.3bp (64.6bp) | Dref/dmmpmm(Bigfoot)/fly(0.701) More Information \| Similar Motifs Found | motif file (matrix) |
| 18 | 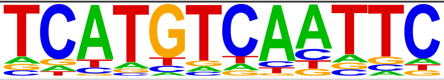 | 1e-28 | -6.666e+01 | 0.16% | 0.00% | 46.3bp (1.7bp) | CG11617/MA0173.1/Jaspar(0.697) More Information \| Similar Motifs Found | motif file (matrix) |
| | | | | | | | VRN1(ABI3VP1)/col-VRN1-DAP- | |

| | | | | | | | | |
|---|---|---|---|---|---|---|---|---|
| 19 | 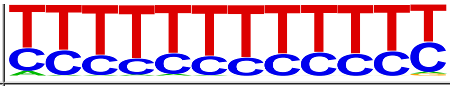 | 1e-25 | -5.766e+01 | 2.61% | 1.42% | 57.2bp (72.3bp) | Seq(GSE60143)/Homer(0.853) More Information \| Similar Motifs Found | motif file (matrix) |
| 20 | 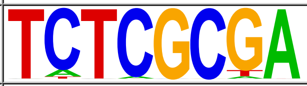 | 1e-23 | -5.340e+01 | 1.46% | 0.65% | 54.7bp (64.8bp) | ZBTB33/MA0527.1/Jaspar(0.901) More Information \| Similar Motifs Found | motif file (matrix) |
| 21 | 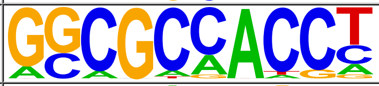 | 1e-14 | -3.438e+01 | 3.61% | 2.48% | 53.9bp (59.4bp) | RPN4/MA0373.1/Jaspar(0.861) More Information \| Similar Motifs Found | motif file (matrix) |
| 22 * | 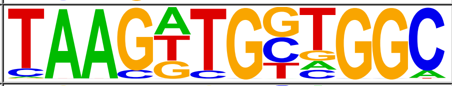 | 1e-10 | -2.458e+01 | 0.29% | 0.08% | 62.3bp (7.2bp) | YY1(Zf)/Promoter/Homer(0.751) More Information \| Similar Motifs Found | motif file (matrix) |
| 23 * | 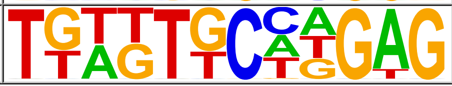 | 1e-4 | -1.076e+01 | 0.06% | 0.01% | 66.4bp (44.7bp) | slbo/dmmpmm(Down)/fly(0.645) More Information \| Similar Motifs Found | motif file (matrix) |

S5 Table. Full list of known motif discovery from ChIP-seq data targeting PBX3.

### Homer Known Motif Enrichment Results (/wynton/group/marcucio/2022CHM/Data2022Mar/Motif/HomerPBX3IDR)

Homer *de novo* Motif Results
Gene Ontology Enrichment Results
Known Motif Enrichment Results (txt file)
Total Target Sequences = 36119, Total Background Sequences = 35379

| Rank | Motif | Name | P-value | log P-pvalue | q-value (Benjamini) | # Target Sequences with Motif | % o Targ Sequ with Mot |
|---|---|---|---|---|---|---|---|
| 1 | 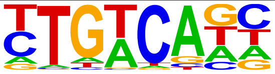 | Tgif1(Homeobox)/mES-Tgif1-ChIP-Seq(GSE55404)/Homer | 1e-2802 | -6.454e+03 | 0.0000 | 23203.0 | 64.2 |
| 2 | 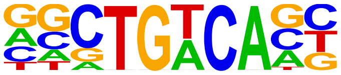 | Meis1(Homeobox)/MastCells-Meis1-ChIP-Seq(GSE48085)/Homer | 1e-2776 | -6.394e+03 | 0.0000 | 16533.0 | 45.7 |
| 3 | 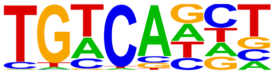 | Tgif2(Homeobox)/mES-Tgif2-ChIP-Seq(GSE55404)/Homer | 1e-2403 | -5.535e+03 | 0.0000 | 23110.0 | 63.9 |
| 4 | 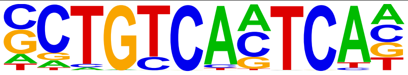 | Pbx3(Homeobox)/GM12878-PBX3-ChIP-Seq(GSE32465)/Homer | 1e-1661 | -3.825e+03 | 0.0000 | 4067.0 | 11.2 |
| 5 | 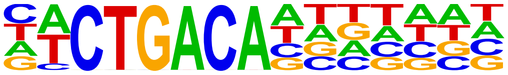 | GRF9(GRF)/colamp-GRF9-DAP-Seq(GSE60143)/Homer | 1e-1358 | -3.129e+03 | 0.0000 | 9651.0 | 26.7 |
| 6 | 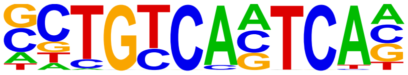 | Pknox1(Homeobox)/ES-Prep1-ChIP-Seq(GSE63282)/Homer | 1e-1346 | -3.100e+03 | 0.0000 | 3558.0 | 9.85 |
| 7 | 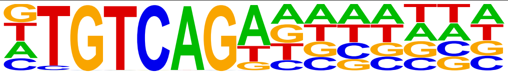 | AtGRF6(GRF)/col-AtGRF6-DAP-Seq(GSE60143)/Homer | 1e-1335 | -3.076e+03 | 0.0000 | 10601.0 | 29.3 |
| 8 | 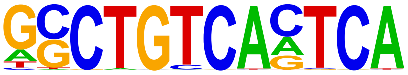 | PBX1(Homeobox)/MCF7-PBX1-ChIP-Seq(GSE28007)/Homer | 1e-807 | -1.860e+03 | 0.0000 | 1628.0 | 4.51 |
| 9 | 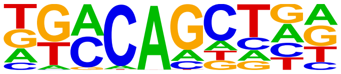 | bZIP18(bZIP)/colamp-bZIP18-DAP-Seq(GSE60143)/Homer | 1e-490 | -1.129e+03 | 0.0000 | 25460.0 | 70.4 |
| 10 | 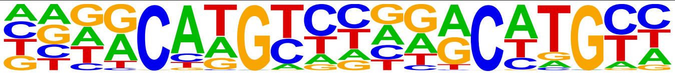 | p63(p53)/Keratinocyte-p63-ChIP-Seq(GSE17611)/Homer | 1e-481 | -1.108e+03 | 0.0000 | 2822.0 | 7.81 |
| 11 | 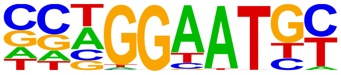 | TEAD4(TEA)/Tropoblast-Tead4-ChIP-Seq(GSE37350)/Homer | 1e-452 | -1.043e+03 | 0.0000 | 4926.0 | 13.6 |
| 12 | 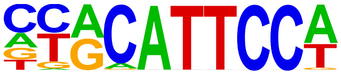 | TEAD1(TEAD)/HepG2-TEAD1-ChIP-Seq(Encode)/Homer | 1e-433 | -9.985e+02 | 0.0000 | 5416.0 | 14.9 |
| 13 | 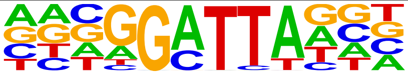 | bcd(Homeobox)/Embryo-Bcd-ChIP-Seq(GSE86966)/Homer | 1e-413 | -9.524e+02 | 0.0000 | 6120.0 | 16.9 |
| 14 | 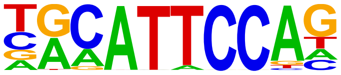 | TEAD3(TEA)/HepG2-TEAD3-ChIP-Seq(Encode)/Homer | 1e-400 | -9.219e+02 | 0.0000 | 6105.0 | 16.9 |
| 15 | 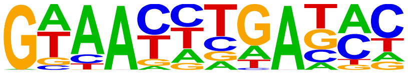 | Six2(Homeobox)/NephronProgenitor-Six2-ChIP-Seq(GSE39837)/Homer | 1e-393 | -9.052e+02 | 0.0000 | 5459.0 | 15.1 |
| 16 | 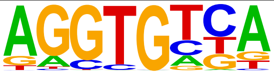 | Tbx5(T-box)/HL1-Tbx5.biotin-ChIP-Seq(GSE21529)/Homer | 1e-388 | -8.941e+02 | 0.0000 | 15503.0 | 42.9 |
| 17 | 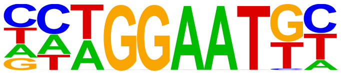 | TEAD(TEA)/Fibroblast-PU.1-ChIP-Seq(Unpublished)/Homer | 1e-365 | -8.418e+02 | 0.0000 | 4086.0 | 11.3 |
| 18 | 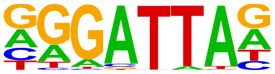 | GSC(Homeobox)/FrogEmbryos-GSC-ChIP-Seq(DRA000576)/Homer | 1e-362 | -8.337e+02 | 0.0000 | 6170.0 | 17.0 |
| 19 | 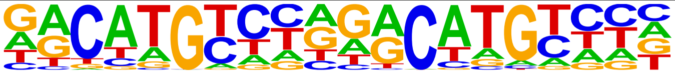 | p53(p53)/Saos-p53-ChIP-Seq(GSE15780)/Homer | 1e-339 | -7.817e+02 | 0.0000 | 1122.0 | 3.11 |
| 20 | 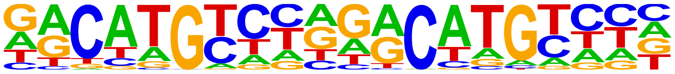 | p53(p53)/Saos-p53-ChIP-Seq/Homer | 1e-339 | -7.817e+02 | 0.0000 | 1122.0 | 3.11 |

| | | | | | | | |
|---|---|---|---|---|---|---|---|
| 21 | 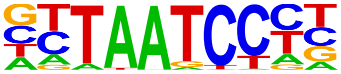 | Otx2(Homeobox)/EpiLC-Otx2-ChIP-Seq(GSE56098)/Homer | 1e-337 | -7.764e+02 | 0.0000 | 4493.0 | 12.4 |
| 22 | 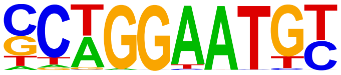 | TEAD2(TEA)/Py2T-Tead2-ChIP-Seq(GSE55709)/Homer | 1e-331 | -7.637e+02 | 0.0000 | 3213.0 | 8.89 |
| 23 | 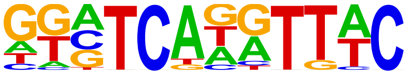 | Six1(Homeobox)/Myoblast-Six1-ChIP-Chip(GSE20150)/Homer | 1e-323 | -7.441e+02 | 0.0000 | 1903.0 | 5.27 |
| 24 | 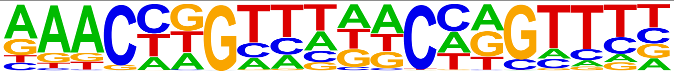 | GRHL2(CP2)/HBE-GRHL2-ChIP-Seq(GSE46194)/Homer | 1e-319 | -7.359e+02 | 0.0000 | 2512.0 | 6.95 |
| 25 | 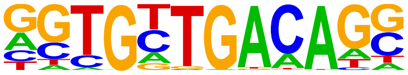 | Tbx20(T-box)/Heart-Tbx20-ChIP-Seq(GSE29636)/Homer | 1e-298 | -6.867e+02 | 0.0000 | 2425.0 | 6.71 |
| 26 | 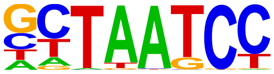 | CRX(Homeobox)/Retina-Crx-ChIP-Seq(GSE20012)/Homer | 1e-295 | -6.797e+02 | 0.0000 | 10523.0 | 29.1 |
| 27 | 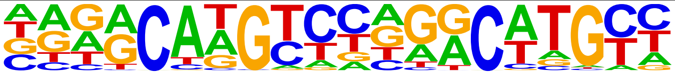 | p73(p53)/Trachea-p73-ChIP-Seq(PRJNA310161)/Homer | 1e-293 | -6.747e+02 | 0.0000 | 705.0 | 1.95 |
| 28 | 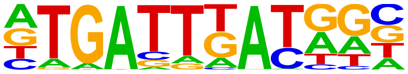 | PBX2(Homeobox)/K562-PBX2-ChIP-Seq(Encode)/Homer | 1e-281 | -6.481e+02 | 0.0000 | 4956.0 | 13.7 |
| 29 | 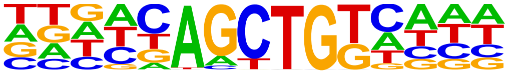 | bZIP52(bZIP)/colamp-bZIP52-DAP-Seq(GSE60143)/Homer | 1e-216 | -4.992e+02 | 0.0000 | 8842.0 | 24.4 |
| 30 | 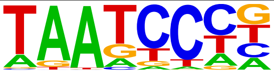 | Pitx1(Homeobox)/Chicken-Pitx1-ChIP-Seq(GSE38910)/Homer | 1e-197 | -4.536e+02 | 0.0000 | 17625.0 | 48.7 |
| 31 | 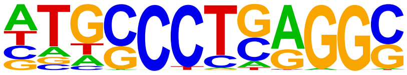 | AP-2alpha(AP2)/Hela-AP2alpha-ChIP-Seq(GSE31477)/Homer | 1e-189 | -4.367e+02 | 0.0000 | 4148.0 | 11.4 |
| 32 | 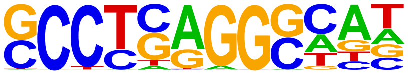 | AP-2gamma(AP2)/MCF7-TFAP2C-ChIP-Seq(GSE21234)/Homer | 1e-188 | -4.345e+02 | 0.0000 | 5259.0 | 14.5 |
| 33 | 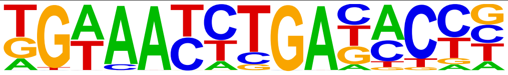 | Six4(Homeobox)/MCF7-SIX4-ChIP-Seq(Encode)/Homer | 1e-173 | -3.992e+02 | 0.0000 | 503.0 | 1.39 |
| 34 | 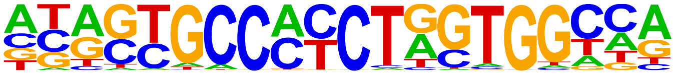 | CTCF(Zf)/CD4+-CTCF-ChIP-Seq(Barski_et_al.)/Homer | 1e-171 | -3.944e+02 | 0.0000 | 970.0 | 2.69 |
| 35 | 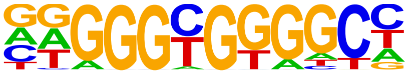 | KLF1(Zf)/HUDEP2-KLF1-CutnRun(GSE136251)/Homer | 1e-154 | -3.569e+02 | 0.0000 | 3241.0 | 8.97 |
| 36 | 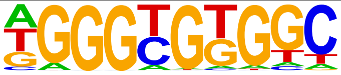 | KLF5(Zf)/LoVo-KLF5-ChIP-Seq(GSE49402)/Homer | 1e-150 | -3.470e+02 | 0.0000 | 4594.0 | 12.7 |
| 37 | 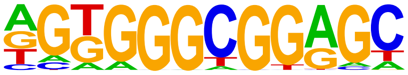 | Sp5(Zf)/mES-Sp5.Flag-ChIP-Seq(GSE72989)/Homer | 1e-150 | -3.466e+02 | 0.0000 | 4064.0 | 11.2 |
| 38 | 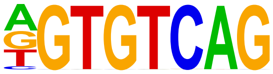 | CELF2(RRM)/JSL1-CELF2-CLIP-Seq(GSE71264)/Homer | 1e-150 | -3.457e+02 | 0.0000 | 2478.0 | 6.86 |
| 39 | 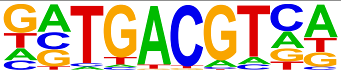 | bZIP50(bZIP)/colamp-bZIP50-DAP-Seq(GSE60143)/Homer | 1e-149 | -3.448e+02 | 0.0000 | 4584.0 | 12.6 |
| 40 | 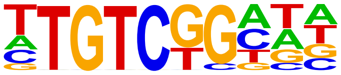 | ARF2(ARF)/col-ARF2-DAP-Seq(GSE60143)/Homer | 1e-139 | -3.217e+02 | 0.0000 | 12355.0 | 34.2 |
| 41 | 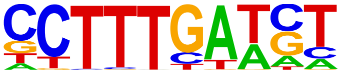 | LEF1(HMG)/H1-LEF1-ChIP-Seq(GSE64758)/Homer | 1e-126 | -2.909e+02 | 0.0000 | 3720.0 | 10.3 |
| 42 | 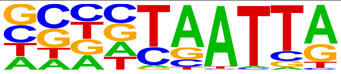 | DLX1(Homeobox)/BasalGanglia-Dlx1-ChIP-seq(GSE124936)/Homer | 1e-124 | -2.874e+02 | 0.0000 | 6866.0 | 19.0 |
| 43 | 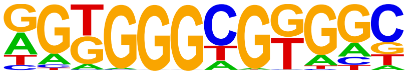 | KLF14(Zf)/HEK293-KLF14.GFP-ChIP-Seq(GSE58341)/Homer | 1e-124 | -2.864e+02 | 0.0000 | 6592.0 | 18.2 |
| 44 | 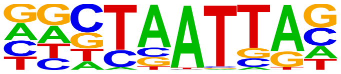 | En1(Homeobox)/SUM149-EN1-ChIP-Seq(GSE120957)/Homer | 1e-123 | -2.842e+02 | 0.0000 | 8838.0 | 24.4 |
| 45 | 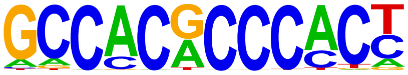 | Klf9(Zf)/GBM-Klf9-ChIP-Seq(GSE62211)/Homer | 1e-123 | -2.833e+02 | 0.0000 | 1336.0 | 3.70 |

| | | | | | | | |
|---|---|---|---|---|---|---|---|
| 46 | 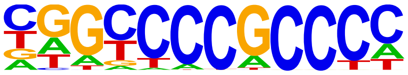 | Sp2(Zf)/HEK293-Sp2.eGFP-ChIP-Seq(Encode)/Homer | 1e-119 | -2.753e+02 | 0.0000 | 5633.0 | 15.5 |
| 47 | 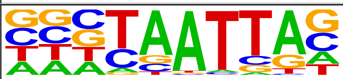 | DLX2(Homeobox)/BasalGanglia-Dlx2-ChIP-seq(GSE124936)/Homer | 1e-115 | -2.671e+02 | 0.0000 | 7436.0 | 20.5 |
| 48 | 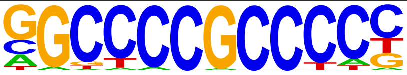 | Sp1(Zf)/Promoter/Homer | 1e-113 | -2.606e+02 | 0.0000 | 1766.0 | 4.89 |
| 49 | 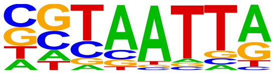 | DLX5(Homeobox)/BasalGanglia-Dlx5-ChIP-seq(GSE124936)/Homer | 1e-112 | -2.590e+02 | 0.0000 | 4401.0 | 12.1 |
| 50 | 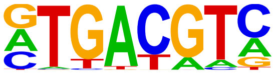 | TGA9(bZIP)/colamp-TGA9-DAP-Seq(GSE60143)/Homer | 1e-110 | -2.536e+02 | 0.0000 | 5677.0 | 15.7 |
| 51 | 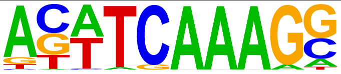 | Tcf3(HMG)/mES-Tcf3-ChIP-Seq(GSE11724)/Homer | 1e-109 | -2.514e+02 | 0.0000 | 1627.0 | 4.50 |
| 52 | 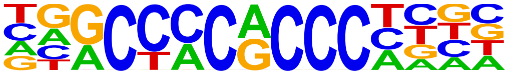 | KLF3(Zf)/MEF-Klf3-ChIP-Seq(GSE44748)/Homer | 1e-106 | -2.444e+02 | 0.0000 | 1766.0 | 4.89 |
| 53 | 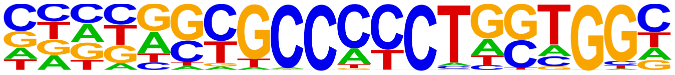 | BORIS(Zf)/K562-CTCFL-ChIP-Seq(GSE32465)/Homer | 1e-103 | -2.380e+02 | 0.0000 | 1291.0 | 3.57 |
| 54 | 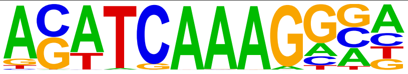 | Tcf4(HMG)/Hct116-Tcf4-ChIP-Seq(SRA012054)/Homer | 1e-99 | -2.294e+02 | 0.0000 | 2596.0 | 7.19 |
| 55 | 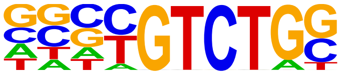 | Smad4(MAD)/ESC-SMAD4-ChIP-Seq(GSE29422)/Homer | 1e-98 | -2.276e+02 | 0.0000 | 7191.0 | 19.9 |
| 56 | 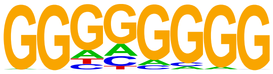 | Maz(Zf)/HepG2-Maz-ChIP-Seq(GSE31477)/Homer | 1e-96 | -2.231e+02 | 0.0000 | 5625.0 | 15.5 |
| 57 | 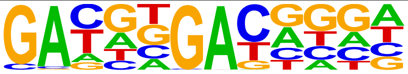 | Knotted(Homeobox)/Corn-KN1-ChIP-Seq(GSE39161)/Homer | 1e-89 | -2.070e+02 | 0.0000 | 6418.0 | 17.7 |
| 58 | 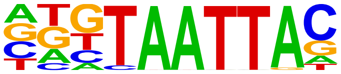 | Dlx3(Homeobox)/Kerainocytes-Dlx3-ChIP-Seq(GSE89884)/Homer | 1e-86 | -2.002e+02 | 0.0000 | 3769.0 | 10.4 |
| 59 | 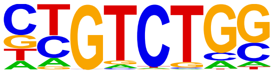 | Smad2(MAD)/ES-SMAD2-ChIP-Seq(GSE29422)/Homer | 1e-85 | -1.980e+02 | 0.0000 | 7060.0 | 19.5 |
| 60 | 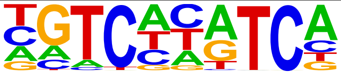 | Replumless(BLH)/Arabidopsis-RPL.GFP-ChIP-Seq(GSE78727)/Homer | 1e-84 | -1.954e+02 | 0.0000 | 5015.0 | 13.8 |
| 61 | 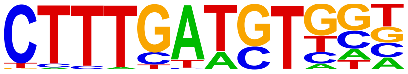 | Tcf7(HMG)/GM12878-TCF7-ChIP-Seq(Encode)/Homer | 1e-83 | -1.933e+02 | 0.0000 | 1896.0 | 5.25 |
| 62 | 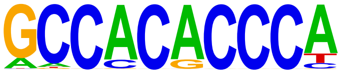 | Klf4(Zf)/mES-Klf4-ChIP-Seq(GSE11431)/Homer | 1e-83 | -1.924e+02 | 0.0000 | 1008.0 | 2.79 |
| 63 | 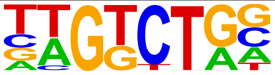 | Smad3(MAD)/NPC-Smad3-ChIP-Seq(GSE36673)/Homer | 1e-81 | -1.868e+02 | 0.0000 | 11352.0 | 31.4 |
| 64 | 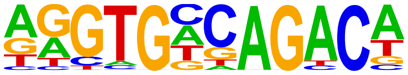 | Tbox:Smad(T-box,MAD)/ESCd5-Smad2_3-ChIP-Seq(GSE29422)/Homer | 1e-79 | -1.825e+02 | 0.0000 | 1045.0 | 2.89 |
| 65 | 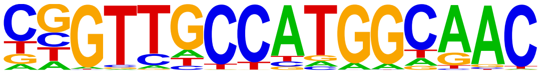 | RFX(HTH)/K562-RFX3-ChIP-Seq(SRA012198)/Homer | 1e-78 | -1.811e+02 | 0.0000 | 533.0 | 1.48 |
| 66 | 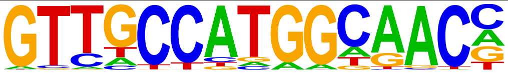 | Rfx2(HTH)/LoVo-RFX2-ChIP-Seq(GSE49402)/Homer | 1e-78 | -1.810e+02 | 0.0000 | 553.0 | 1.53 |
| 67 | 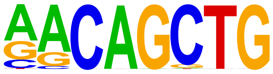 | MyoG(bHLH)/C2C12-MyoG-ChIP-Seq(GSE36024)/Homer | 1e-76 | -1.760e+02 | 0.0000 | 6493.0 | 17.9 |
| 68 | 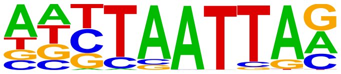 | Lhx3(Homeobox)/Neuron-Lhx3-ChIP-Seq(GSE31456)/Homer | 1e-76 | -1.753e+02 | 0.0000 | 7897.0 | 21.8 |
| 69 | 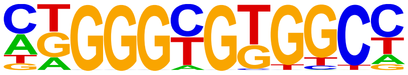 | KLF6(Zf)/PDAC-KLF6-ChIP-Seq(GSE64557)/Homer | 1e-75 | -1.740e+02 | 0.0000 | 3988.0 | 11.0 |
| 70 | 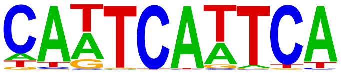 | WUS1(Homeobox)/colamp-WUS1-DAP-Seq(GSE60143)/Homer | 1e-74 | -1.713e+02 | 0.0000 | 2141.0 | 5.93 |

| | | | | | | | |
|---|---|---|---|---|---|---|---|
| 71 | 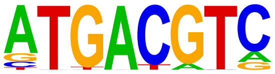 | TGA10(bZIP)/colamp-TGA10-DAP-Seq(GSE60143)/Homer | 1e-73 | -1.684e+02 | 0.0000 | 2607.0 | 7.22 |
| 72 | 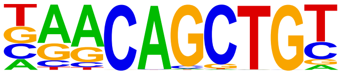 | Myf5(bHLH)/GM-Myf5-ChIP-Seq(GSE24852)/Homer | 1e-70 | -1.628e+02 | 0.0000 | 4902.0 | 13.5 |
| 73 | 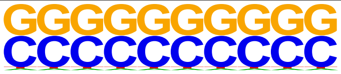 | SeqBias: CG bias | 1e-67 | -1.556e+02 | 0.0000 | 25456.0 | 70.4 |
| 74 | 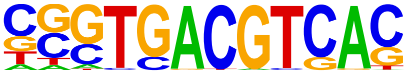 | CRE(bZIP)/Promoter/Homer | 1e-67 | -1.551e+02 | 0.0000 | 1303.0 | 3.61 |
| 75 | 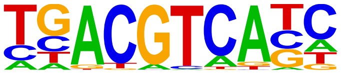 | TGA6(bZIP)/colamp-TGA6-DAP-Seq(GSE60143)/Homer | 1e-66 | -1.537e+02 | 0.0000 | 2548.0 | 7.05 |
| 76 | 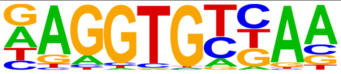 | Tbx6(T-box)/ESC-Tbx6-ChIP-Seq(GSE93524)/Homer | 1e-65 | -1.498e+02 | 0.0000 | 4551.0 | 12.6 |
| 77 | 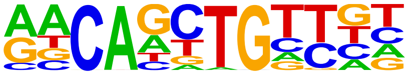 | HLH-1(bHLH)/cElegans-Embryo-HLH1-ChIP-Seq(modEncode)/Homer | 1e-64 | -1.489e+02 | 0.0000 | 4296.0 | 11.8 |
| 78 | 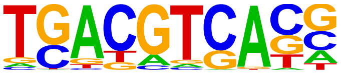 | FEA4(bZIP)/Corn-FEA4-ChIP-Seq(GSE61954)/Homer | 1e-64 | -1.486e+02 | 0.0000 | 4827.0 | 13.3 |
| 79 | 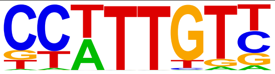 | Sox3(HMG)/NPC-Sox3-ChIP-Seq(GSE33059)/Homer | 1e-62 | -1.434e+02 | 0.0000 | 7509.0 | 20.7 |
| 80 | 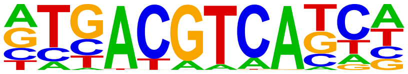 | TGA4(bZIP)/colamp-TGA4-DAP-Seq(GSE60143)/Homer | 1e-61 | -1.426e+02 | 0.0000 | 1372.0 | 3.80 |
| 81 | 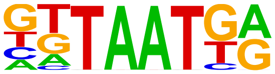 | Nkx6.1(Homeobox)/Islet-Nkx6.1-ChIP-Seq(GSE40975)/Homer | 1e-61 | -1.421e+02 | 0.0000 | 11474.0 | 31.7 |
| 82 | 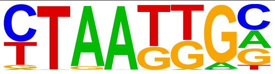 | Isl1(Homeobox)/Neuron-Isl1-ChIP-Seq(GSE31456)/Homer | 1e-61 | -1.416e+02 | 0.0000 | 8315.0 | 23.0 |
| 83 | 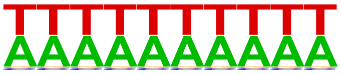 | SeqBias: A/T bias | 1e-60 | -1.382e+02 | 0.0000 | 29583.0 | 81.9 |
| 84 | 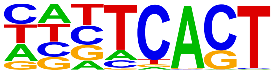 | STZ(C2H2)/colamp-STZ-DAP-Seq(GSE60143)/Homer | 1e-57 | -1.323e+02 | 0.0000 | 23216.0 | 64.2 |
| 85 | 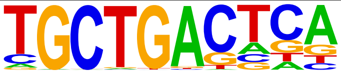 | MafA(bZIP)/Islet-MafA-ChIP-Seq(GSE30298)/Homer | 1e-55 | -1.288e+02 | 0.0000 | 4885.0 | 13.5 |
| 86 | 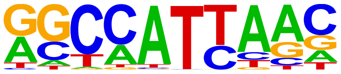 | Nanog(Homeobox)/mES-Nanog-ChIP-Seq(GSE11724)/Homer | 1e-55 | -1.286e+02 | 0.0000 | 17174.0 | 47.5 |
| 87 | 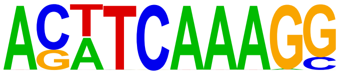 | TCFL2(HMG)/K562-TCF7L2-ChIP-Seq(GSE29196)/Homer | 1e-55 | -1.284e+02 | 0.0000 | 595.0 | 1.65 |
| 88 | 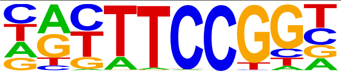 | Elk4(ETS)/Hela-Elk4-ChIP-Seq(GSE31477)/Homer | 1e-55 | -1.283e+02 | 0.0000 | 1869.0 | 5.17 |
| 89 | 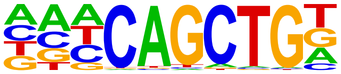 | Ap4(bHLH)/AML-Tfap4-ChIP-Seq(GSE45738)/Homer | 1e-54 | -1.257e+02 | 0.0000 | 7498.0 | 20.7 |
| 90 | 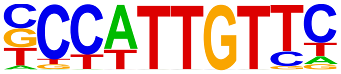 | Sox2(HMG)/mES-Sox2-ChIP-Seq(GSE11431)/Homer | 1e-53 | -1.243e+02 | 0.0000 | 3733.0 | 10.3 |
| 91 | 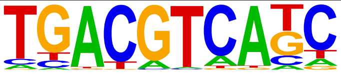 | TGA1(bZIP)/colamp-TGA1-DAP-Seq(GSE60143)/Homer | 1e-53 | -1.243e+02 | 0.0000 | 1822.0 | 5.04 |
| 92 | 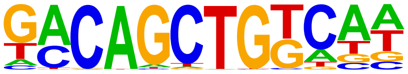 | bZIP69(bZIP)/col-bZIP69-DAP-Seq(GSE60143)/Homer | 1e-52 | -1.220e+02 | 0.0000 | 952.0 | 2.64 |
| 93 | 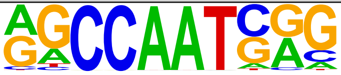 | NFY(CCAAT)/Promoter/Homer | 1e-52 | -1.210e+02 | 0.0000 | 2758.0 | 7.64 |
| 94 | 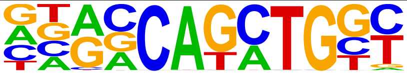 | Atoh1(bHLH)/Cerebellum-Atoh1-ChIP-Seq(GSE22111)/Homer | 1e-50 | -1.160e+02 | 0.0000 | 5583.0 | 15.4 |
| 95 | 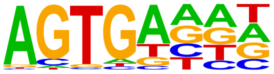 | At5g04390(C2H2)/col200-At5g04390-DAP-Seq(GSE60143)/Homer | 1e-49 | -1.139e+02 | 0.0000 | 21818.0 | 60.4 |

| | | | | | | | |
|---|---|---|---|---|---|---|---|
| 96 | 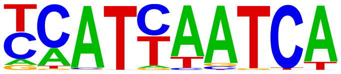 | Pdx1(Homeobox)/Islet-Pdx1-ChIP-Seq(SRA008281)/Homer | 1e-48 | -1.119e+02 | 0.0000 | 4407.0 | 12.2 |
| 97 | 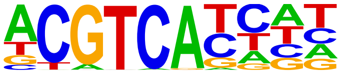 | TGA2(bZIP)/colamp-TGA2-DAP-Seq(GSE60143)/Homer | 1e-47 | -1.095e+02 | 0.0000 | 2316.0 | 6.41 |
| 98 | 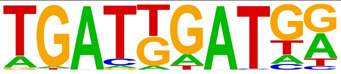 | HOXA1(Homeobox)/mES-Hoxa1-ChIP-Seq(SRP084292)/Homer | 1e-47 | -1.089e+02 | 0.0000 | 1368.0 | 3.79 |
| 99 | 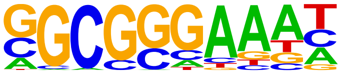 | E2F4(E2F)/K562-E2F4-ChIP-Seq(GSE31477)/Homer | 1e-45 | -1.047e+02 | 0.0000 | 1858.0 | 5.14 |
| 100 | 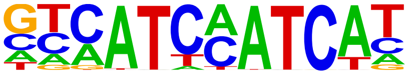 | HOXA2(Homeobox)/mES-Hoxa2-ChIP-Seq(Donaldson_et_al.)/Homer | 1e-44 | -1.033e+02 | 0.0000 | 605.0 | 1.67 |
| 101 | 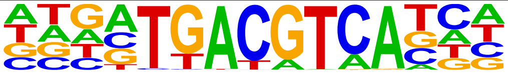 | TGA5(bZIP)/col-TGA5-DAP-Seq(GSE60143)/Homer | 1e-44 | -1.021e+02 | 0.0000 | 450.0 | 1.25 |
| 102 | 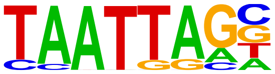 | Lhx2(Homeobox)/HFSC-Lhx2-ChIP-Seq(GSE48068)/Homer | 1e-44 | -1.019e+02 | 0.0000 | 4926.0 | 13.6 |
| 103 | 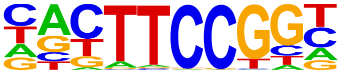 | Elk1(ETS)/Hela-Elk1-ChIP-Seq(GSE31477)/Homer | 1e-43 | -1.004e+02 | 0.0000 | 1876.0 | 5.19 |
| 104 | 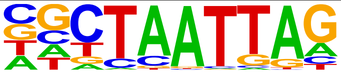 | LHX9(Homeobox)/Hct116-LHX9.V5-ChIP-Seq(GSE116822)/Homer | 1e-42 | -9.748e+01 | 0.0000 | 6169.0 | 17.0 |
| 105 | 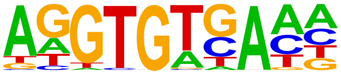 | Tbet(T-box)/CD8-Tbet-ChIP-Seq(GSE33802)/Homer | 1e-42 | -9.699e+01 | 0.0000 | 4427.0 | 12.2 |
| 106 | 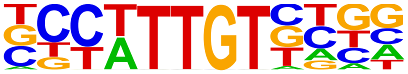 | Sox21(HMG)/ESC-SOX21-ChIP-Seq(GSE110505)/Homer | 1e-41 | -9.658e+01 | 0.0000 | 7662.0 | 21.2 |
| 107 | 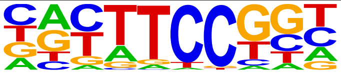 | Fli1(ETS)/CD8-FLI-ChIP-Seq(GSE20898)/Homer | 1e-41 | -9.551e+01 | 0.0000 | 3925.0 | 10.8 |
| 108 | 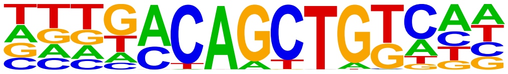 | VIP1(bZIP)/col-VIP1-DAP-Seq(GSE60143)/Homer | 1e-40 | -9.430e+01 | 0.0000 | 1318.0 | 3.65 |
| 109 | 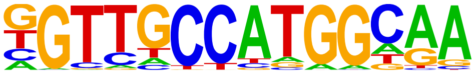 | Rfx1(HTH)/NPC-H3K4me1-ChIP-Seq(GSE16256)/Homer | 1e-40 | -9.318e+01 | 0.0000 | 864.0 | 2.39 |
| 110 | 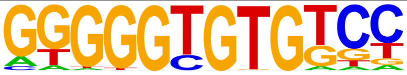 | KLF10(Zf)/HEK293-KLF10.GFP-ChIP-Seq(GSE58341)/Homer | 1e-40 | -9.228e+01 | 0.0000 | 1596.0 | 4.42 |
| 111 | 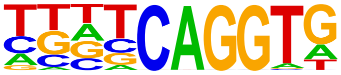 | NGA4(ABI3VP1)/col-NGA4-DAP-Seq(GSE60143)/Homer | 1e-39 | -9.172e+01 | 0.0000 | 10590.0 | 29.3 |
| 112 | 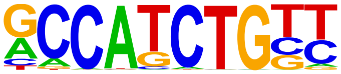 | NeuroD1(bHLH)/Islet-NeuroD1-ChIP-Seq(GSE30298)/Homer | 1e-38 | -8.883e+01 | 0.0000 | 3783.0 | 10.4 |
| 113 | 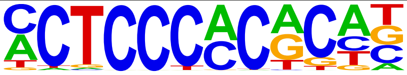 | WT1(Zf)/Kidney-WT1-ChIP-Seq(GSE90016)/Homer | 1e-38 | -8.872e+01 | 0.0000 | 2142.0 | 5.93 |
| 114 | 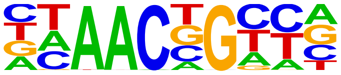 | BMYB(HTH)/Hela-BMYB-ChIP-Seq(GSE27030)/Homer | 1e-37 | -8.690e+01 | 0.0000 | 7346.0 | 20.3 |
| 115 | 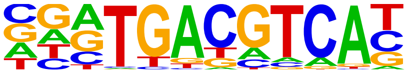 | Atf7(bZIP)/3T3L1-Atf7-ChIP-Seq(GSE56872)/Homer | 1e-35 | -8.240e+01 | 0.0000 | 1860.0 | 5.15 |
| 116 | 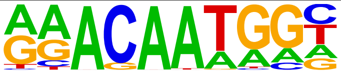 | Sox15(HMG)/CPA-Sox15-ChIP-Seq(GSE62909)/Homer | 1e-35 | -8.131e+01 | 0.0000 | 4115.0 | 11.3 |
| 117 | 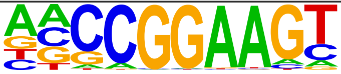 | ELF1(ETS)/Jurkat-ELF1-ChIP-Seq(SRA014231)/Homer | 1e-34 | -8.050e+01 | 0.0000 | 1590.0 | 4.40 |
| 118 | 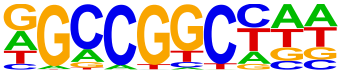 | RAP211(AP2EREBP)/colamp-RAP211-DAP-Seq(GSE60143)/Homer | 1e-34 | -8.016e+01 | 0.0000 | 8739.0 | 24.1 |
| 119 | 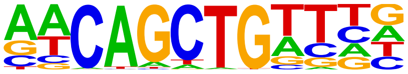 | E-box/Drosophila-Promoters/Homer | 1e-34 | -7.967e+01 | 0.0000 | 1683.0 | 4.66 |
| 120 | | AMYB(HTH)/Testes-AMYB-ChIP-Seq(GSE44588)/Homer | 1e-34 | -7.962e+01 | 0.0000 | 7289.0 | 20.1 |

| | | | | | | | |
|---|---|---|---|---|---|---|---|
| | 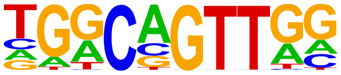 | | | | | | |
| 121 | 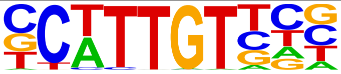 | Sox10(HMG)/SciaticNerve-Sox3-ChIP-Seq(GSE35132)/Homer | 1e-34 | -7.944e+01 | 0.0000 | 7121.0 | 19.7 |
| 122 | 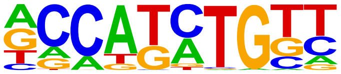 | Olig2(bHLH)/Neuron-Olig2-ChIP-Seq(GSE30882)/Homer | 1e-34 | -7.925e+01 | 0.0000 | 8862.0 | 24.5 |
| 123 | 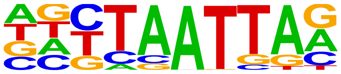 | Lhx1(Homeobox)/EmbryoCarcinoma-Lhx1-ChIP-Seq(GSE70957)/Homer | 1e-33 | -7.802e+01 | 0.0000 | 5189.0 | 14.3 |
| 124 | 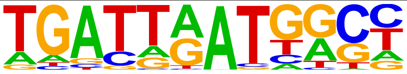 | Hoxb4(Homeobox)/ES-Hoxb4-ChIP-Seq(GSE34014)/Homer | 1e-33 | -7.784e+01 | 0.0000 | 1021.0 | 2.83 |
| 125 | 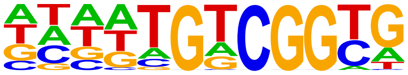 | AT1G71450(AP2EREBP)/col-AT1G71450-DAP-Seq(GSE60143)/Homer | 1e-33 | -7.783e+01 | 0.0000 | 5587.0 | 15.4 |
| 126 | 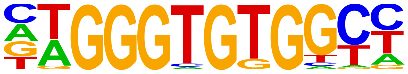 | EKLF(Zf)/Erythrocyte-Klf1-ChIP-Seq(GSE20478)/Homer | 1e-33 | -7.759e+01 | 0.0000 | 417.0 | 1.15 |
| 127 | 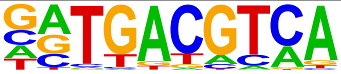 | Atf1(bZIP)/K562-ATF1-ChIP-Seq(GSE31477)/Homer | 1e-33 | -7.754e+01 | 0.0000 | 2731.0 | 7.56 |
| 128 | 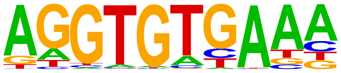 | Tbx21(T-box)/GM12878-TBX21-ChIP-Seq(Encode)/Homer | 1e-33 | -7.681e+01 | 0.0000 | 4057.0 | 11.2 |
| 129 | 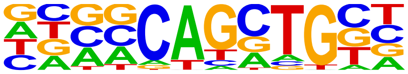 | Ascl1(bHLH)/NeuralTubes-Ascl1-ChIP-Seq(GSE55840)/Homer | 1e-33 | -7.628e+01 | 0.0000 | 7581.0 | 20.9 |
| 130 | 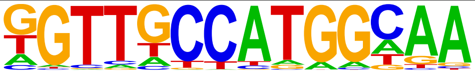 | X-box(HTH)/NPC-H3K4me1-ChIP-Seq(GSE16256)/Homer | 1e-32 | -7.578e+01 | 0.0000 | 521.0 | 1.44 |
| 131 | 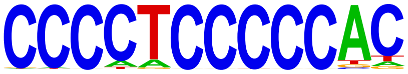 | Zfp281(Zf)/ES-Zfp281-ChIP-Seq(GSE81042)/Homer | 1e-32 | -7.567e+01 | 0.0000 | 605.0 | 1.67 |
| 132 | 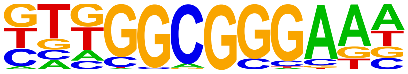 | E2F3(E2F)/MEF-E2F3-ChIP-Seq(GSE71376)/Homer | 1e-32 | -7.513e+01 | 0.0000 | 2605.0 | 7.21 |
| 133 | 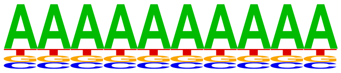 | SeqBias: polyA-repeat | 1e-31 | -7.356e+01 | 0.0000 | 34471.0 | 95.4 |
| 134 | 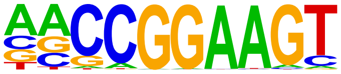 | ETS(ETS)/Promoter/Homer | 1e-31 | -7.270e+01 | 0.0000 | 1037.0 | 2.87 |
| 135 | 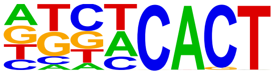 | AZF1(C2H2)/colamp-AZF1-DAP-Seq(GSE60143)/Homer | 1e-31 | -7.245e+01 | 0.0000 | 19774.0 | 54.7 |
| 136 | 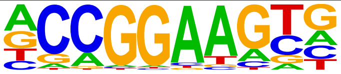 | ETV4(ETS)/HepG2-ETV4-ChIP-Seq(ENCODE)/Homer | 1e-31 | -7.152e+01 | 0.0000 | 3826.0 | 10.5 |
| 137 | 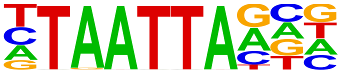 | ATHB23(ZFHD)/col-ATHB23-DAP-Seq(GSE60143)/Homer | 1e-31 | -7.141e+01 | 0.0000 | 3993.0 | 11.0 |
| 138 | 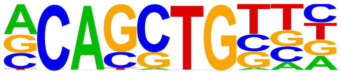 | Ptf1a(bHLH)/Panc1-Ptf1a-ChIP-Seq(GSE47459)/Homer | 1e-29 | -6.870e+01 | 0.0000 | 12657.0 | 35.0 |
| 139 | 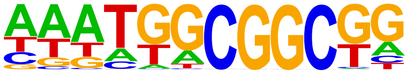 | ABR1(AP2EREBP)/colamp-ABR1-DAP-Seq(GSE60143)/Homer | 1e-29 | -6.864e+01 | 0.0000 | 4538.0 | 12.5 |
| 140 | 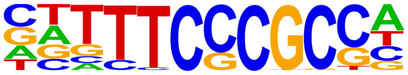 | E2F7(E2F)/Hela-E2F7-ChIP-Seq(GSE32673)/Homer | 1e-29 | -6.832e+01 | 0.0000 | 493.0 | 1.36 |
| 141 | 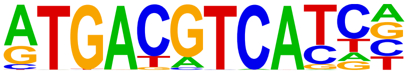 | JunD(bZIP)/K562-JunD-ChIP-Seq/Homer | 1e-29 | -6.824e+01 | 0.0000 | 397.0 | 1.10 |
| 142 | 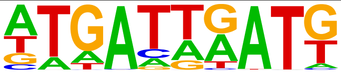 | LIN-39(Homeobox)/cElegans.L3-LIN39-ChIP-Seq(modEncode)/Homer | 1e-28 | -6.465e+01 | 0.0000 | 4886.0 | 13.5 |
| 143 | 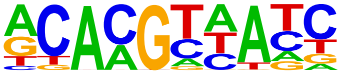 | ANAC038(NAC)/col-ANAC038-DAP-Seq(GSE60143)/Homer | 1e-27 | -6.403e+01 | 0.0000 | 8625.0 | 23.8 |
| 144 | 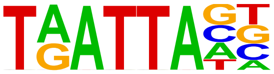 | ATHB25(ZFHD)/colamp-ATHB25-DAP-Seq(GSE60143)/Homer | 1e-27 | -6.275e+01 | 0.0000 | 4790.0 | 13.2 |

| | | | | | | | |
|---|---|---|---|---|---|---|---|
| 145 | 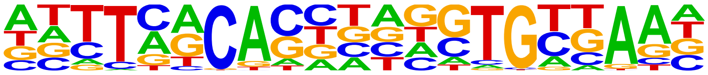 | Brachyury(T-box)/Mesoendoderm-Brachyury-ChIP-exo(GSE54963)/Homer | 1e-27 | -6.261e+01 | 0.0000 | 1469.0 | 4.07 |
| 146 | 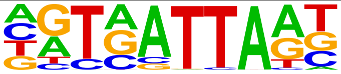 | ATHB33(ZFHD)/col-ATHB33-DAP-Seq(GSE60143)/Homer | 1e-27 | -6.219e+01 | 0.0000 | 5753.0 | 15.9 |
| 147 | 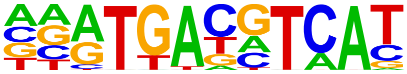 | CREB5(bZIP)/LNCaP-CREB5.V5-ChIP-Seq(GSE137775)/Homer | 1e-26 | -6.196e+01 | 0.0000 | 1448.0 | 4.01 |
| 148 | 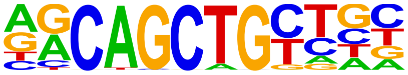 | MyoD(bHLH)/Myotube-MyoD-ChIP-Seq(GSE21614)/Homer | 1e-26 | -6.194e+01 | 0.0000 | 5225.0 | 14.4 |
| 149 | 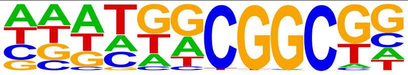 | RAP26(AP2EREBP)/colamp-RAP26-DAP-Seq(GSE60143)/Homer | 1e-26 | -6.126e+01 | 0.0000 | 5585.0 | 15.4 |
| 150 | 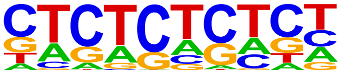 | GAGA-repeat/Arabidopsis-Promoters/Homer | 1e-26 | -6.121e+01 | 0.0000 | 4064.0 | 11.2 |
| 151 | 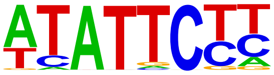 | KAN2(G2like)/colamp-KAN2-DAP-Seq(GSE60143)/Homer | 1e-26 | -6.038e+01 | 0.0000 | 5007.0 | 13.8 |
| 152 | 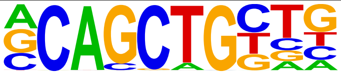 | Tcf12(bHLH)/GM12878-Tcf12-ChIP-Seq(GSE32465)/Homer | 1e-26 | -6.010e+01 | 0.0000 | 5755.0 | 15.9 |
| 153 | 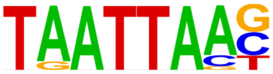 | ATHB24(ZFHD)/colamp-ATHB24-DAP-Seq(GSE60143)/Homer | 1e-26 | -5.991e+01 | 0.0000 | 3220.0 | 8.91 |
| 154 | 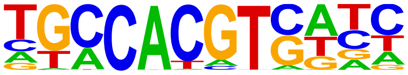 | GBF3(bZIP)/Arabidopsis-GBF3-ChIP-Seq(GSE80564)/Homer | 1e-25 | -5.926e+01 | 0.0000 | 1609.0 | 4.45 |
| 155 | 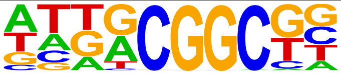 | ERF115(AP2EREBP)/colamp-ERF115-DAP-Seq(GSE60143)/Homer | 1e-25 | -5.911e+01 | 0.0000 | 6863.0 | 19.0 |
| 156 | 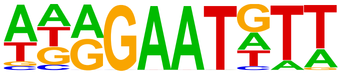 | AT2G38300(G2like)/col-AT2G38300-DAP-Seq(GSE60143)/Homer | 1e-25 | -5.878e+01 | 0.0000 | 4942.0 | 13.6 |
| 157 | 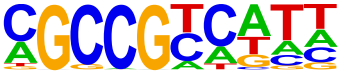 | ERF8(AP2EREBP)/colamp-ERF8-DAP-Seq(GSE60143)/Homer | 1e-25 | -5.771e+01 | 0.0000 | 4929.0 | 13.6 |
| 158 | 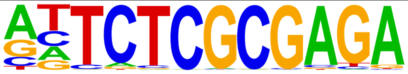 | GFX(?)/Promoter/Homer | 1e-24 | -5.700e+01 | 0.0000 | 94.0 | 0.26 |
| 159 | 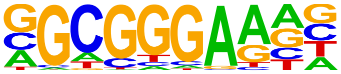 | E2F6(E2F)/Hela-E2F6-ChIP-Seq(GSE31477)/Homer | 1e-24 | -5.629e+01 | 0.0000 | 2378.0 | 6.58 |
| 160 | 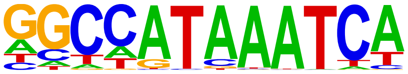 | Hoxc9(Homeobox)/Ainv15-Hoxc9-ChIP-Seq(GSE21812)/Homer | 1e-24 | -5.588e+01 | 0.0000 | 2280.0 | 6.31 |
| 161 | 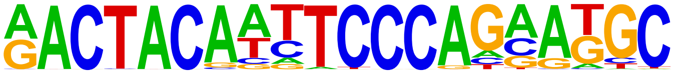 | GFY-Staf(?,Zf)/Promoter/Homer | 1e-24 | -5.585e+01 | 0.0000 | 251.0 | 0.69 |
| 162 | 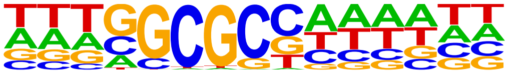 | E2FA(E2FDP)/colamp-E2FA-DAP-Seq(GSE60143)/Homer | 1e-24 | -5.566e+01 | 0.0000 | 1035.0 | 2.87 |
| 163 | 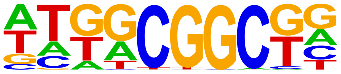 | At2g33710(AP2EREBP)/colamp-At2g33710-DAP-Seq(GSE60143)/Homer | 1e-24 | -5.563e+01 | 0.0000 | 7871.0 | 21.7 |
| 164 | 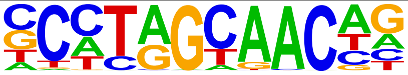 | Rfx5(HTH)/GM12878-Rfx5-ChIP-Seq(GSE31477)/Homer | 1e-24 | -5.549e+01 | 0.0000 | 1164.0 | 3.22 |
| 165 | 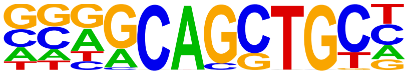 | Ascl2(bHLH)/ESC-Ascl2-ChIP-Seq(GSE97712)/Homer | 1e-24 | -5.541e+01 | 0.0000 | 5971.0 | 16.5 |
| 166 | 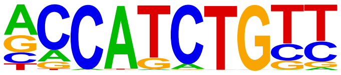 | NeuroG2(bHLH)/Fibroblast-NeuroG2-ChIP-Seq(GSE75910)/Homer | 1e-23 | -5.511e+01 | 0.0000 | 7200.0 | 19.9 |
| 167 | 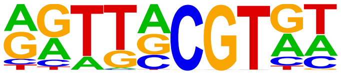 | NAM(NAC)/col-NAM-DAP-Seq(GSE60143)/Homer | 1e-23 | -5.511e+01 | 0.0000 | 4261.0 | 11.8 |
| 168 | 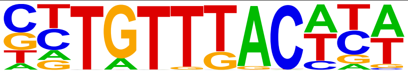 | Foxa2(Forkhead)/Liver-Foxa2-ChIP-Seq(GSE25694)/Homer | 1e-23 | -5.465e+01 | 0.0000 | 3249.0 | 8.99 |
| 169 | 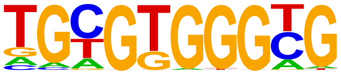 | Egr1(Zf)/K562-Egr1-ChIP-Seq(GSE32465)/Homer | 1e-23 | -5.444e+01 | 0.0000 | 2233.0 | 6.18 |

| | | | | | | | |
|---|---|---|---|---|---|---|---|
| 170 | 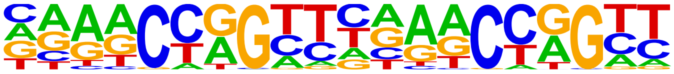 | Tcfcp2l1(CP2)/mES-Tcfcp2l1-ChIP-Seq(GSE11431)/Homer | 1e-23 | -5.444e+01 | 0.0000 | 520.0 | 1.44 |
| 171 | 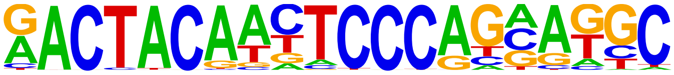 | Ronin(THAP)/ES-Thap11-ChIP-Seq(GSE51522)/Homer | 1e-23 | -5.405e+01 | 0.0000 | 179.0 | 0.50 |
| 172 | 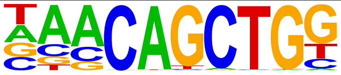 | Tcf21(bHLH)/ArterySmoothMuscle-Tcf21-ChIP-Seq(GSE61369)/Homer | 1e-23 | -5.346e+01 | 0.0000 | 5371.0 | 14.8 |
| 173 | 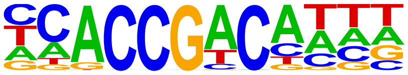 | DEAR2(AP2EREBP)/colamp-DEAR2-DAP-Seq(GSE60143)/Homer | 1e-23 | -5.335e+01 | 0.0000 | 3046.0 | 8.43 |
| 174 | 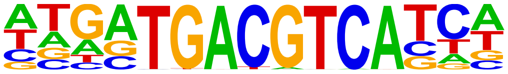 | TGA3(bZIP)/colamp-TGA3-DAP-Seq(GSE60143)/Homer | 1e-22 | -5.274e+01 | 0.0000 | 276.0 | 0.76 |
| 175 | 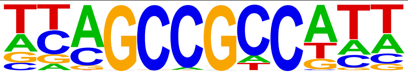 | ERF13(AP2EREBP)/colamp-ERF13-DAP-Seq(GSE60143)/Homer | 1e-22 | -5.249e+01 | 0.0000 | 5467.0 | 15.1 |
| 176 | 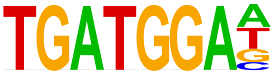 | HAP3(CCAATHAP3)/col-HAP3-DAP-Seq(GSE60143)/Homer | 1e-22 | -5.243e+01 | 0.0000 | 1575.0 | 4.36 |
| 177 | 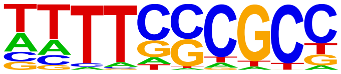 | DEL2(E2FDP)/col-DEL2-DAP-Seq(GSE60143)/Homer | 1e-22 | -5.197e+01 | 0.0000 | 1297.0 | 3.59 |
| 178 | 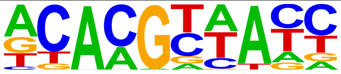 | ANAC046(NAC)/colamp-ANAC046-DAP-Seq(GSE60143)/Homer | 1e-22 | -5.112e+01 | 0.0000 | 7723.0 | 21.3 |
| 179 | 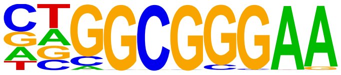 | E2F1(E2F)/Hela-E2F1-ChIP-Seq(GSE22478)/Homer | 1e-21 | -5.052e+01 | 0.0000 | 1046.0 | 2.90 |
| 180 | 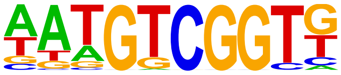 | DREB19(AP2EREBP)/colamp-DREB19-DAP-Seq(GSE60143)/Homer | 1e-21 | -5.043e+01 | 0.0000 | 1624.0 | 4.50 |
| 181 | 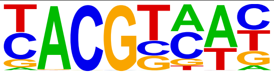 | ATAF1(NAC)/col-ATAF1-DAP-Seq(GSE60143)/Homer | 1e-21 | -4.993e+01 | 0.0000 | 9614.0 | 26.6 |
| 182 | 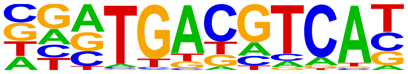 | Atf2(bZIP)/3T3L1-Atf2-ChIP-Seq(GSE56872)/Homer | 1e-21 | -4.972e+01 | 0.0000 | 1240.0 | 3.43 |
| 183 | 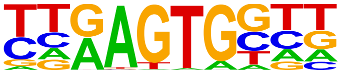 | Bapx1(Homeobox)/VertebralCol-Bapx1-ChIP-Seq(GSE36672)/Homer | 1e-21 | -4.891e+01 | 0.0000 | 7819.0 | 21.6 |
| 184 | 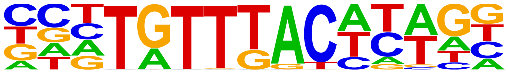 | Foxa3(Forkhead)/Liver-Foxa3-ChIP-Seq(GSE77670)/Homer | 1e-21 | -4.889e+01 | 0.0000 | 1270.0 | 3.52 |
| 185 | 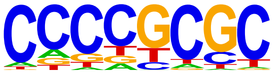 | SUT1?/SacCer-Promoters/Homer | 1e-21 | -4.855e+01 | 0.0000 | 20912.0 | 57.8 |
| 186 | 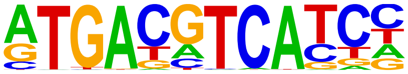 | c-Jun-CRE(bZIP)/K562-cJun-ChIP-Seq(GSE31477)/Homer | 1e-20 | -4.819e+01 | 0.0000 | 1143.0 | 3.16 |
| 187 | 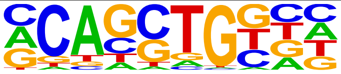 | HEB(bHLH)/mES-Heb-ChIP-Seq(GSE53233)/Homer | 1e-20 | -4.776e+01 | 0.0000 | 8747.0 | 24.2 |
| 188 | 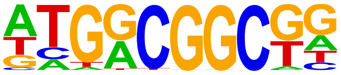 | ERF7(AP2EREBP)/col-ERF7-DAP-Seq(GSE60143)/Homer | 1e-20 | -4.746e+01 | 0.0000 | 5686.0 | 15.7 |
| 189 | 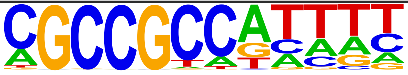 | RAP212(AP2EREBP)/col-RAP212-DAP-Seq(GSE60143)/Homer | 1e-20 | -4.727e+01 | 0.0000 | 3735.0 | 10.3 |
| 190 | 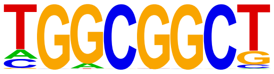 | ERF105(AP2EREBP)/colamp-ERF105-DAP-Seq(GSE60143)/Homer | 1e-20 | -4.685e+01 | 0.0000 | 6552.0 | 18.1 |
| 191 | 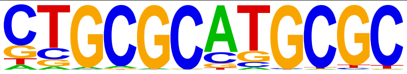 | NRF1(NRF)/MCF7-NRF1-ChIP-Seq(Unpublished)/Homer | 1e-19 | -4.603e+01 | 0.0000 | 642.0 | 1.78 |
| 192 | 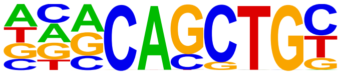 | E2A(bHLH)/proBcell-E2A-ChIP-Seq(GSE21978)/Homer | 1e-19 | -4.570e+01 | 0.0000 | 7091.0 | 19.6 |
| 193 | 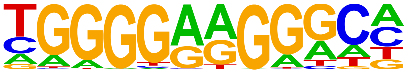 | ZNF467(Zf)/HEK293-ZNF467.GFP-ChIP-Seq(GSE58341)/Homer | 1e-19 | -4.378e+01 | 0.0000 | 3227.0 | 8.93 |
| 194 | 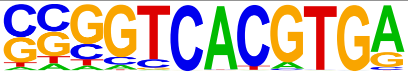 | E-box(bHLH)/Promoter/Homer | 1e-18 | -4.375e+01 | 0.0000 | 335.0 | 0.93 |

| | | | | | | | |
|---|---|---|---|---|---|---|---|
| 195 | 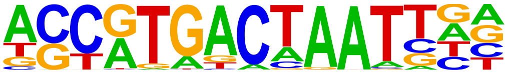 | PAX3:FKHR-fusion(Paired,Homeobox)/Rh4-PAX3:FKHR-ChIP-Seq(GSE19063)/Homer | 1e-18 | -4.372e+01 | 0.0000 | 878.0 | 2.43 |
| 196 | 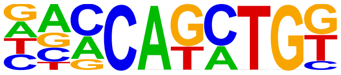 | BHLHA15(bHLH)/NIH3T3-BHLHB8.HA-ChIP-Seq(GSE119782)/Homer | 1e-18 | -4.331e+01 | 0.0000 | 6718.0 | 18.6 |
| 197 | 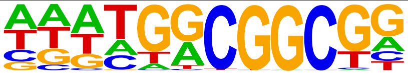 | ERF4(AP2EREBP)/colamp-ERF4-DAP-Seq(GSE60143)/Homer | 1e-18 | -4.329e+01 | 0.0000 | 4931.0 | 13.6 |
| 198 | 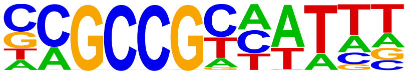 | RRTF1(AP2EREBP)/colamp-RRTF1-DAP-Seq(GSE60143)/Homer | 1e-18 | -4.316e+01 | 0.0000 | 1161.0 | 3.21 |
| 199 | 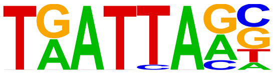 | ATHB34(ZFHD)/colamp-ATHB34-DAP-Seq(GSE60143)/Homer | 1e-18 | -4.249e+01 | 0.0000 | 3375.0 | 9.34 |
| 200 | 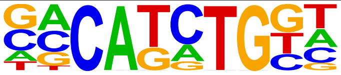 | TCF4(bHLH)/SHSY5Y-TCF4-ChIP-Seq(GSE96915)/Homer | 1e-18 | -4.244e+01 | 0.0000 | 6900.0 | 19.1 |
| 201 | 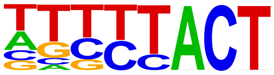 | At3g60580(C2H2)/col-At3g60580-DAP-Seq(GSE60143)/Homer | 1e-18 | -4.225e+01 | 0.0000 | 19050.0 | 52.7 |
| 202 | 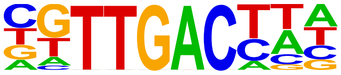 | WRKY28(WRKY)/col-WRKY28-DAP-Seq(GSE60143)/Homer | 1e-18 | -4.205e+01 | 0.0000 | 3630.0 | 10.0 |
| 203 | 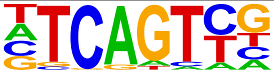 | Initiator/Drosophila-Promoters/Homer | 1e-18 | -4.185e+01 | 0.0000 | 9692.0 | 26.8 |
| 204 | 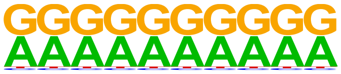 | SeqBias: G/A bias | 1e-18 | -4.169e+01 | 0.0000 | 36117.0 | 99.9 |
| 205 | 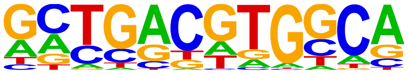 | O2(bZIP)/Corn-O2-ChIP-Seq(GSE63991)/Homer | 1e-17 | -4.141e+01 | 0.0000 | 684.0 | 1.89 |
| 206 | 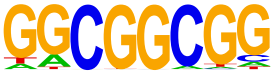 | AT1G28160(AP2EREBP)/colamp-AT1G28160-DAP-Seq(GSE60143)/Homer | 1e-17 | -4.133e+01 | 0.0000 | 7686.0 | 21.2 |
| 207 | 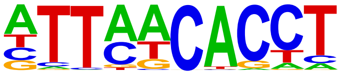 | Eomes(T-box)/H9-Eomes-ChIP-Seq(GSE26097)/Homer | 1e-17 | -4.110e+01 | 0.0000 | 8535.0 | 23.6 |
| 208 | 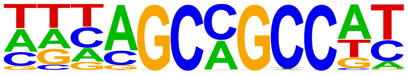 | ERF15(AP2EREBP)/colamp-ERF15-DAP-Seq(GSE60143)/Homer | 1e-17 | -4.086e+01 | 0.0000 | 10531.0 | 29.1 |
| 209 | 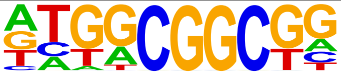 | ERF11(AP2EREBP)/col-ERF11-DAP-Seq(GSE60143)/Homer | 1e-17 | -4.049e+01 | 0.0000 | 4449.0 | 12.3 |
| 210 | 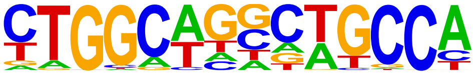 | Tlx?(NR)/NPC-H3K4me1-ChIP-Seq(GSE16256)/Homer | 1e-17 | -4.030e+01 | 0.0000 | 1535.0 | 4.25 |
| 211 | 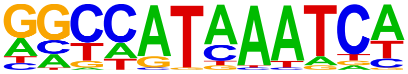 | HOXA9(Homeobox)/HSC-Hoxa9-ChIP-Seq(GSE33509)/Homer | 1e-17 | -4.025e+01 | 0.0000 | 2888.0 | 8.00 |
| 212 | 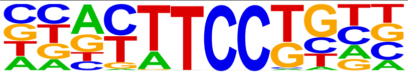 | Etv2(ETS)/ES-ER71-ChIP-Seq(GSE59402)/Homer | 1e-17 | -3.985e+01 | 0.0000 | 3219.0 | 8.91 |
| 213 | 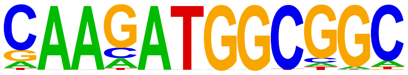 | YY1(Zf)/Promoter/Homer | 1e-17 | -3.957e+01 | 0.0000 | 425.0 | 1.18 |
| 214 | 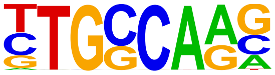 | NF1-halfsite(CTF)/LNCaP-NF1-ChIP-Seq(Unpublished)/Homer | 1e-17 | -3.931e+01 | 0.0000 | 7495.0 | 20.7 |
| 215 | 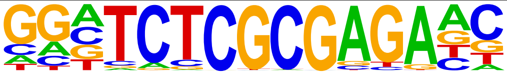 | ZBTB33(Zf)/GM12878-ZBTB33-ChIP-Seq(GSE32465)/Homer | 1e-16 | -3.844e+01 | 0.0000 | 194.0 | 0.54 |
| 216 | 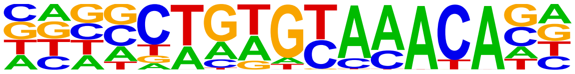 | Fox:Ebox(Forkhead,bHLH)/Panc1-Foxa2-ChIP-Seq(GSE47459)/Homer | 1e-16 | -3.765e+01 | 0.0000 | 4143.0 | 11.4 |
| 217 | 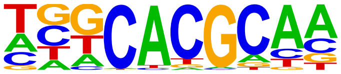 | Arnt:Ahr(bHLH)/MCF7-Arnt-ChIP-Seq(Lo_et_al.)/Homer | 1e-16 | -3.753e+01 | 0.0000 | 3293.0 | 9.12 |
| 218 | 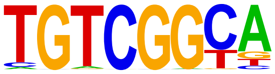 | AT1G12630(AP2EREBP)/colamp-AT1G12630-DAP-Seq(GSE60143)/Homer | 1e-16 | -3.737e+01 | 0.0000 | 1453.0 | 4.02 |
| 219 | 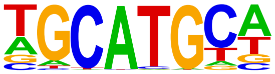 | RBFox2(?)/Heart-RBFox2-CLIP-Seq(GSE57926)/Homer | 1e-16 | -3.698e+01 | 0.0000 | 9181.0 | 25.4 |

| | | | | | | | |
|---|---|---|---|---|---|---|---|
| 220 | 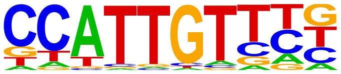 | Sox17(HMG)/Endoderm-Sox17-ChIP-Seq(GSE61475)/Homer | 1e-16 | -3.698e+01 | 0.0000 | 2594.0 | 7.18 |
| 221 | 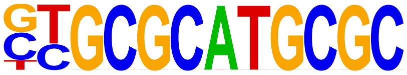 | NRF(NRF)/Promoter/Homer | 1e-15 | -3.664e+01 | 0.0000 | 662.0 | 1.83 |
| 222 | 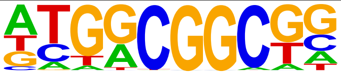 | ERF3(AP2EREBP)/colamp-ERF3-DAP-Seq(GSE60143)/Homer | 1e-15 | -3.660e+01 | 0.0000 | 3806.0 | 10.5 |
| 223 | 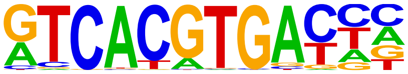 | TFE3(bHLH)/MEF-TFE3-ChIP-Seq(GSE75757)/Homer | 1e-15 | -3.653e+01 | 0.0000 | 284.0 | 0.79 |
| 224 | 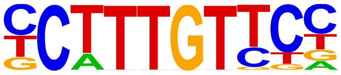 | Sox4(HMG)/proB-Sox4-ChIP-Seq(GSE50066)/Homer | 1e-15 | -3.644e+01 | 0.0000 | 3599.0 | 9.96 |
| 225 | 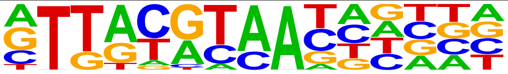 | NFIL3(bZIP)/HepG2-NFIL3-ChIP-Seq(Encode)/Homer | 1e-15 | -3.620e+01 | 0.0000 | 2465.0 | 6.82 |
| 226 | 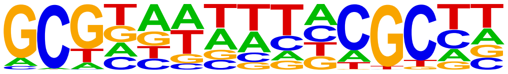 | LBD23(LOBAS2)/colamp-LBD23-DAP-Seq(GSE60143)/Homer | 1e-15 | -3.542e+01 | 0.0000 | 4224.0 | 11.6 |
| 227 | 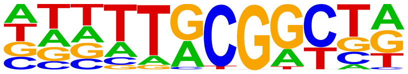 | At5g08750(C3H)/col-At5g08750-DAP-Seq(GSE60143)/Homer | 1e-15 | -3.516e+01 | 0.0000 | 1534.0 | 4.25 |
| 228 | 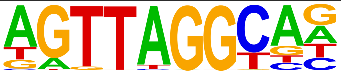 | MYB116(MYB)/colamp-MYB116-DAP-Seq(GSE60143)/Homer | 1e-15 | -3.508e+01 | 0.0000 | 2088.0 | 5.78 |
| 229 | 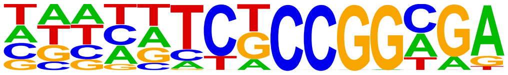 | AT5G05550(Trihelix)/col-AT5G05550-DAP-Seq(GSE60143)/Homer | 1e-15 | -3.504e+01 | 0.0000 | 5123.0 | 14.1 |
| 230 | 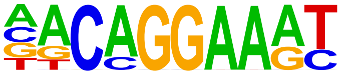 | EWS:FLI1-fusion(ETS)/SK_N_MC-EWS:FLI1-ChIP-Seq(SRA014231)/Homer | 1e-14 | -3.415e+01 | 0.0000 | 1973.0 | 5.46 |
| 231 | 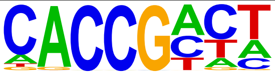 | At1g75490(AP2EREBP)/colamp-At1g75490-DAP-Seq(GSE60143)/Homer | 1e-14 | -3.325e+01 | 0.0000 | 6547.0 | 18.1 |
| 232 | 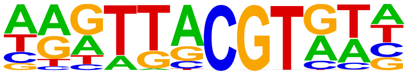 | NAP(NAC)/col-NAP-DAP-Seq(GSE60143)/Homer | 1e-14 | -3.298e+01 | 0.0000 | 3224.0 | 8.93 |
| 233 | 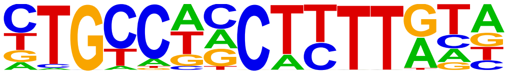 | ZNF7(Zf)/HepG2-ZNF7.Flag-ChIP-Seq(Encode)/Homer | 1e-13 | -3.180e+01 | 0.0000 | 2195.0 | 6.08 |
| 234 | 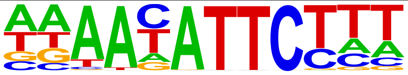 | AT2G40260(G2like)/colamp-AT2G40260-DAP-Seq(GSE60143)/Homer | 1e-13 | -3.166e+01 | 0.0000 | 5847.0 | 16.1 |
| 235 | 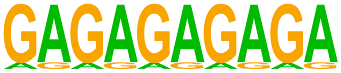 | SeqBias: GA-repeat | 1e-13 | -3.159e+01 | 0.0000 | 23284.0 | 64.4 |
| 236 | 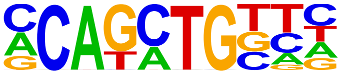 | Twist2(bHLH)/Myoblast-Twist2.Ty1-ChIP-Seq(GSE127998)/Homer | 1e-13 | -3.157e+01 | 0.0000 | 8336.0 | 23.0 |
| 237 | 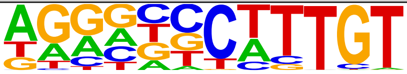 | Sox9(HMG)/Limb-SOX9-ChIP-Seq(GSE73225)/Homer | 1e-13 | -3.149e+01 | 0.0000 | 3306.0 | 9.15 |
| 238 | 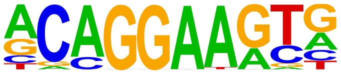 | ERG(ETS)/VCaP-ERG-ChIP-Seq(GSE14097)/Homer | 1e-13 | -3.122e+01 | 0.0000 | 5532.0 | 15.3 |
| 239 | 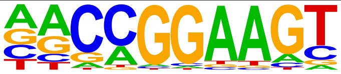 | GABPA(ETS)/Jurkat-GABPa-ChIP-Seq(GSE17954)/Homer | 1e-13 | -3.110e+01 | 0.0000 | 2882.0 | 7.98 |
| 240 | 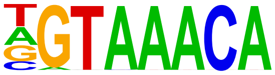 | Foxo3(Forkhead)/U2OS-Foxo3-ChIP-Seq(E-MTAB-2701)/Homer | 1e-13 | -3.084e+01 | 0.0000 | 2780.0 | 7.70 |
| 241 | 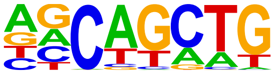 | SCL(bHLH)/HPC7-Scl-ChIP-Seq(GSE13511)/Homer | 1e-13 | -3.049e+01 | 0.0000 | 22088.0 | 61.1 |
| 242 | 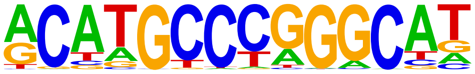 | p53(p53)/mES-cMyc-ChIP-Seq(GSE11431)/Homer | 1e-12 | -2.982e+01 | 0.0000 | 78.0 | 0.22 |
| 243 | 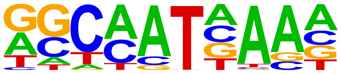 | Hoxa9(Homeobox)/ChickenMSG-Hoxa9.Flag-ChIP-Seq(GSE86088)/Homer | 1e-12 | -2.979e+01 | 0.0000 | 11107.0 | 30.7 |
| 244 | | ERF10(AP2EREBP)/col-ERF10-DAP-Seq(GSE60143)/Homer | 1e-12 | -2.979e+01 | 0.0000 | 3705.0 | 10.2 |

| | | | | | | | |
|---|---|---|---|---|---|---|---|
| | 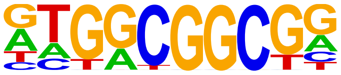 | | | | | | |
| 245 | 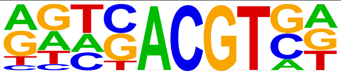 | HY5(bZIP)/colamp-HY5-DAP-Seq(GSE60143)/Homer | 1e-12 | -2.966e+01 | 0.0000 | 2931.0 | 8.11 |
| 246 | 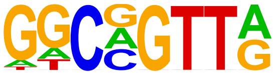 | MYB(HTH)/ERMYB-Myb-ChIPSeq(GSE22095)/Homer | 1e-12 | -2.965e+01 | 0.0000 | 8060.0 | 22.3 |
| 247 | 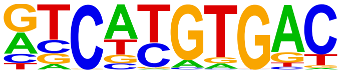 | MITF(bHLH)/MastCells-MITF-ChIP-Seq(GSE48085)/Homer | 1e-12 | -2.932e+01 | 0.0000 | 2983.0 | 8.26 |
| 248 | 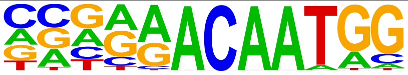 | Sox7(HMG)/ESC-Sox7-ChIP-Seq(GSE133899)/Homer | 1e-12 | -2.924e+01 | 0.0000 | 1019.0 | 2.82 |
| 249 | 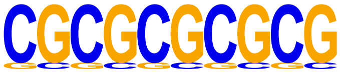 | SeqBias: CG-repeat | 1e-12 | -2.889e+01 | 0.0000 | 7138.0 | 19.7 |
| 250 | 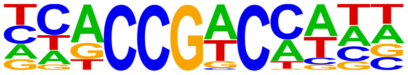 | At1g19210(AP2EREBP)/colamp-At1g19210-DAP-Seq(GSE60143)/Homer | 1e-12 | -2.784e+01 | 0.0000 | 3520.0 | 9.74 |
| 251 | 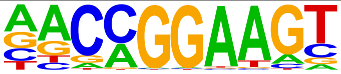 | ETV1(ETS)/GIST48-ETV1-ChIP-Seq(GSE22441)/Homer | 1e-12 | -2.774e+01 | 0.0000 | 4490.0 | 12.4 |
| 252 | 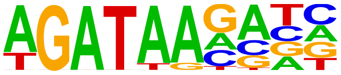 | TRPS1(Zf)/MCF7-TRPS1-ChIP-Seq(GSE107013)/Homer | 1e-11 | -2.739e+01 | 0.0000 | 7506.0 | 20.7 |
| 253 | 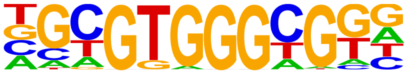 | Egr2(Zf)/Thymocytes-Egr2-ChIP-Seq(GSE34254)/Homer | 1e-11 | -2.685e+01 | 0.0000 | 635.0 | 1.76 |
| 254 | 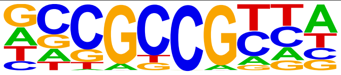 | CRF10(AP2EREBP)/col100-CRF10-DAP-Seq(GSE60143)/Homer | 1e-11 | -2.615e+01 | 0.0000 | 6131.0 | 16.9 |
| 255 | 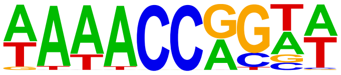 | AT1G76870(Trihelix)/col-AT1G76870-DAP-Seq(GSE60143)/Homer | 1e-11 | -2.603e+01 | 0.0000 | 1362.0 | 3.77 |
| 256 | 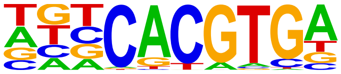 | SPCH(bHLH)/Seedling-SPCH-ChIP-Seq(GSE57497)/Homer | 1e-11 | -2.587e+01 | 0.0000 | 4113.0 | 11.3 |
| 257 | 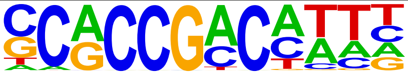 | At5g65130(AP2EREBP)/colamp-At5g65130-DAP-Seq(GSE60143)/Homer | 1e-11 | -2.577e+01 | 0.0000 | 1111.0 | 3.08 |
| 258 | 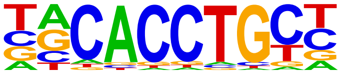 | Snail1(Zf)/LS174T-SNAIL1.HA-ChIP-Seq(GSE127183)/Homer | 1e-11 | -2.569e+01 | 0.0000 | 3344.0 | 9.26 |
| 259 | 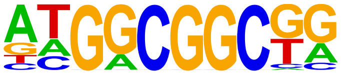 | AT4G18450(AP2EREBP)/col-AT4G18450-DAP-Seq(GSE60143)/Homer | 1e-11 | -2.548e+01 | 0.0000 | 2817.0 | 7.80 |
| 260 | 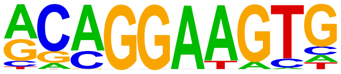 | ETS1(ETS)/Jurkat-ETS1-ChIP-Seq(GSE17954)/Homer | 1e-10 | -2.492e+01 | 0.0000 | 3558.0 | 9.85 |
| 261 | 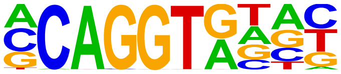 | ZEB1(Zf)/PDAC-ZEB1-ChIP-Seq(GSE64557)/Homer | 1e-10 | -2.471e+01 | 0.0000 | 6627.0 | 18.3 |
| 262 | 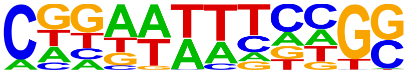 | LBD18(LOBAS2)/colamp-LBD18-DAP-Seq(GSE60143)/Homer | 1e-10 | -2.446e+01 | 0.0000 | 9876.0 | 27.3 |
| 263 | 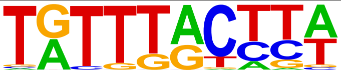 | FOXM1(Forkhead)/MCF7-FOXM1-ChIP-Seq(GSE72977)/Homer | 1e-10 | -2.440e+01 | 0.0000 | 3986.0 | 11.0 |
| 264 | 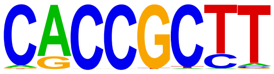 | At5g18450(AP2EREBP)/col-At5g18450-DAP-Seq(GSE60143)/Homer | 1e-10 | -2.415e+01 | 0.0000 | 7100.0 | 19.6 |
| 265 | 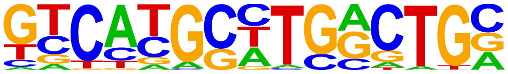 | Pax8(Paired,Homeobox)/Thyroid-Pax8-ChIP-Seq(GSE26938)/Homer | 1e-10 | -2.409e+01 | 0.0000 | 1192.0 | 3.30 |
| 266 | 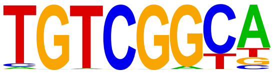 | Rap210(AP2EREBP)/col-Rap210-DAP-Seq(GSE60143)/Homer | 1e-10 | -2.403e+01 | 0.0000 | 1534.0 | 4.25 |
| 267 | 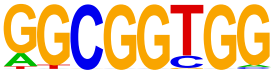 | AT3G57600(AP2EREBP)/col-AT3G57600-DAP-Seq(GSE60143)/Homer | 1e-10 | -2.389e+01 | 0.0000 | 4413.0 | 12.2 |
| 268 | 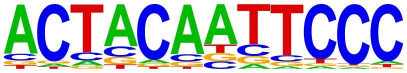 | GFY(?)/Promoter/Homer | 1e-10 | -2.387e+01 | 0.0000 | 271.0 | 0.75 |

| | | | | | | | |
|---|---|---|---|---|---|---|---|
| 269 | 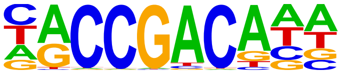 | At4g16750(AP2EREBP)/col-At4g16750-DAP-Seq(GSE60143)/Homer | 1e-10 | -2.378e+01 | 0.0000 | 1804.0 | 4.99 |
| 270 | 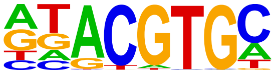 | HIF-1b(HLH)/T47D-HIF1b-ChIP-Seq(GSE59937)/Homer | 1e-10 | -2.367e+01 | 0.0000 | 5676.0 | 15.7 |
| 271 | 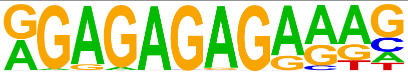 | FRS9(ND)/col-FRS9-DAP-Seq(GSE60143)/Homer | 1e-10 | -2.343e+01 | 0.0000 | 799.0 | 2.21 |
| 272 | 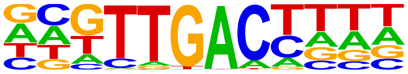 | WRKY27(WRKY)/colamp-WRKY27-DAP-Seq(GSE60143)/Homer | 1e-10 | -2.326e+01 | 0.0000 | 2515.0 | 6.96 |
| 273 | 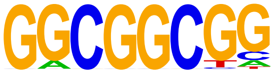 | ERF104(AP2EREBP)/col-ERF104-DAP-Seq(GSE60143)/Homer | 1e-9 | -2.284e+01 | 0.0000 | 5303.0 | 14.6 |
| 274 | 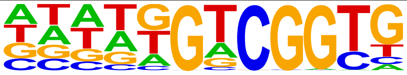 | At1g36060(AP2EREBP)/colamp-At1g36060-DAP-Seq(GSE60143)/Homer | 1e-9 | -2.278e+01 | 0.0000 | 1811.0 | 5.01 |
| 275 | 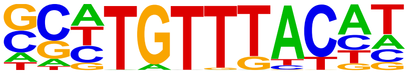 | FOXK2(Forkhead)/U2OS-FOXK2-ChIP-Seq(E-MTAB-2204)/Homer | 1e-9 | -2.205e+01 | 0.0000 | 2374.0 | 6.57 |
| 276 | 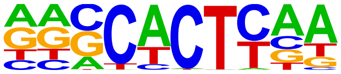 | Nkx2.5(Homeobox)/HL1-Nkx2.5.biotin-ChIP-Seq(GSE21529)/Homer | 1e-9 | -2.178e+01 | 0.0000 | 7547.0 | 20.8 |
| 277 | 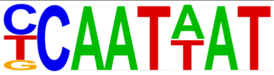 | ATHB21(HB)/colamp-ATHB21-DAP-Seq(GSE60143)/Homer | 1e-9 | -2.173e+01 | 0.0000 | 1621.0 | 4.49 |
| 278 | 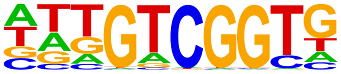 | CEJ1(AP2EREBP)/col-CEJ1-DAP-Seq(GSE60143)/Homer | 1e-9 | -2.171e+01 | 0.0000 | 2536.0 | 7.02 |
| 279 | 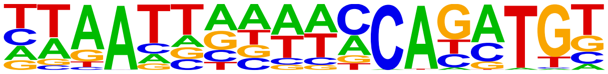 | Pitx1:Ebox(Homeobox,bHLH)/Hindlimb-Pitx1-ChIP-Seq(GSE41591)/Homer | 1e-9 | -2.153e+01 | 0.0000 | 815.0 | 2.26 |
| 280 | 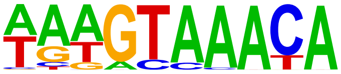 | FOXA1(Forkhead)/LNCAP-FOXA1-ChIP-Seq(GSE27824)/Homer | 1e-9 | -2.137e+01 | 0.0000 | 4633.0 | 12.8 |
| 281 | 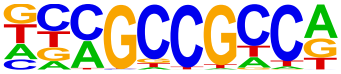 | ERF5(AP2EREBP)/colamp-ERF5-DAP-Seq(GSE60143)/Homer | 1e-9 | -2.133e+01 | 0.0000 | 2994.0 | 8.29 |
| 282 | 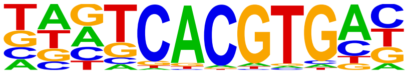 | BIM3(bHLH)/col-BIM3-DAP-Seq(GSE60143)/Homer | 1e-9 | -2.119e+01 | 0.0000 | 466.0 | 1.29 |
| 283 | 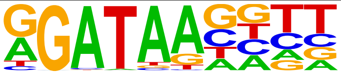 | At1g49010(MYBrelated)/col-At1g49010-DAP-Seq(GSE60143)/Homer | 1e-9 | -2.103e+01 | 0.0000 | 6381.0 | 17.6 |
| 284 | 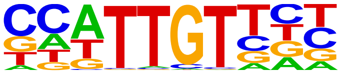 | Sox6(HMG)/Myotubes-Sox6-ChIP-Seq(GSE32627)/Homer | 1e-9 | -2.092e+01 | 0.0000 | 6345.0 | 17.5 |
| 285 | 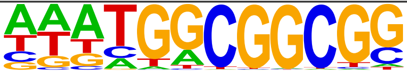 | ERF9(AP2EREBP)/colamp-ERF9-DAP-Seq(GSE60143)/Homer | 1e-9 | -2.092e+01 | 0.0000 | 2014.0 | 5.58 |
| 286 | 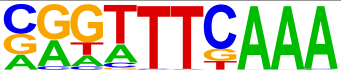 | CHR(?)/Hela-CellCycle-Expression/Homer | 1e-8 | -2.033e+01 | 0.0000 | 2240.0 | 6.20 |
| 287 | 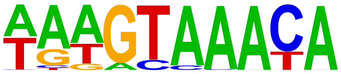 | FOXA1(Forkhead)/MCF7-FOXA1-ChIP-Seq(GSE26831)/Homer | 1e-8 | -1.985e+01 | 0.0000 | 3776.0 | 10.4 |
| 288 | 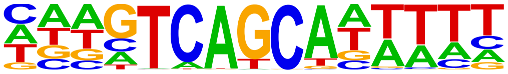 | MafF(bZIP)/HepG2-MafF-ChIP-Seq(GSE31477)/Homer | 1e-8 | -1.956e+01 | 0.0000 | 1359.0 | 3.76 |
| 289 | 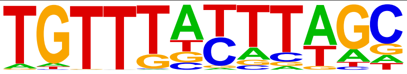 | FoxD3(forkhead)/ZebrafishEmbryo-Foxd3.biotin-ChIP-seq(GSE106676)/Homer | 1e-8 | -1.954e+01 | 0.0000 | 3445.0 | 9.54 |
| 290 | 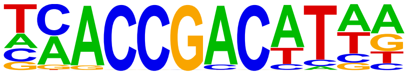 | DREB2(AP2EREBP)/col-DREB2-DAP-Seq(GSE60143)/Homer | 1e-8 | -1.953e+01 | 0.0000 | 1165.0 | 3.23 |
| 291 | 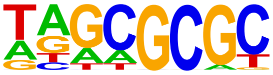 | DPL-1(E2F)/cElegans-Adult-ChIP-Seq(modEncode)/Homer | 1e-8 | -1.909e+01 | 0.0000 | 2930.0 | 8.11 |
| 292 | 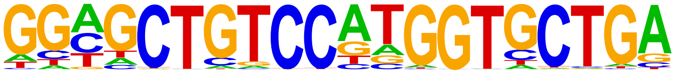 | REST-NRSF(Zf)/Jurkat-NRSF-ChIP-Seq/Homer | 1e-8 | -1.891e+01 | 0.0000 | 58.0 | 0.16 |
| 293 | 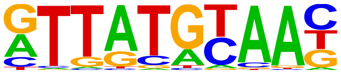 | HLF(bZIP)/HSC-HLF.Flag-ChIP-Seq(GSE69817)/Homer | 1e-8 | -1.886e+01 | 0.0000 | 3142.0 | 8.70 |

| | | | | | | | |
|---|---|---|---|---|---|---|---|
| 294 | 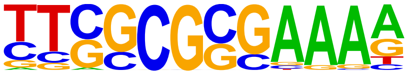 | E2F(E2F)/Hela-CellCycle-Expression/Homer | 1e-8 | -1.877e+01 | 0.0000 | 145.0 | 0.40 |
| 295 | 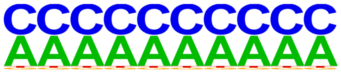 | SeqBias: C/A-bias | 1e-8 | -1.867e+01 | 0.0000 | 36108.0 | 99.9 |
| 296 | 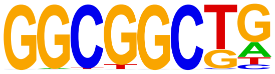 | AT5G23930(mTERF)/col-AT5G23930-DAP-Seq(GSE60143)/Homer | 1e-8 | -1.848e+01 | 0.0000 | 8482.0 | 23.4 |
| 297 | 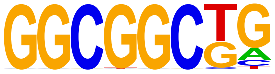 | ERF2(AP2EREBP)/colamp-ERF2-DAP-Seq(GSE60143)/Homer | 1e-7 | -1.840e+01 | 0.0000 | 4254.0 | 11.7 |
| 298 | 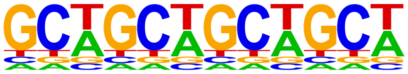 | SeqBias: GCW-triplet | 1e-7 | -1.839e+01 | 0.0000 | 36122.0 | 100. |
| 299 | 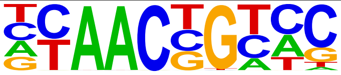 | MYB77(MYB)/col-MYB77-DAP-Seq(GSE60143)/Homer | 1e-7 | -1.836e+01 | 0.0000 | 5691.0 | 15.7 |
| 300 | 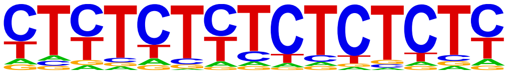 | GAGA-repeat/SacCer-Promoters/Homer | 1e-7 | -1.821e+01 | 0.0000 | 14418.0 | 39.9 |
| 301 | 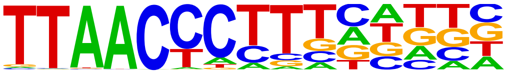 | ZNF652/HepG2-ZNF652.Flag-ChIP-Seq(Encode)/Homer | 1e-7 | -1.820e+01 | 0.0000 | 890.0 | 2.46 |
| 302 | 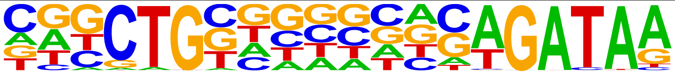 | GATA:SCL(Zf,bHLH)/Ter119-SCL-ChIP-Seq(GSE18720)/Homer | 1e-7 | -1.807e+01 | 0.0000 | 509.0 | 1.41 |
| 303 | 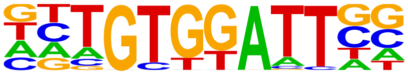 | Foxh1(Forkhead)/hESC-FOXH1-ChIP-Seq(GSE29422)/Homer | 1e-7 | -1.799e+01 | 0.0000 | 2098.0 | 5.81 |
| 304 | 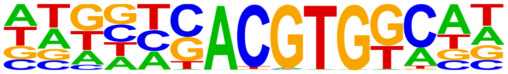 | bZIP3(bZIP)/col-bZIP3-DAP-Seq(GSE60143)/Homer | 1e-7 | -1.780e+01 | 0.0000 | 1693.0 | 4.69 |
| 305 | 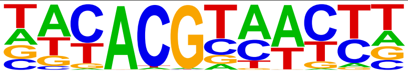 | ANAC047(NAC)/colamp-ANAC047-DAP-Seq(GSE60143)/Homer | 1e-7 | -1.773e+01 | 0.0000 | 2428.0 | 6.72 |
| 306 | 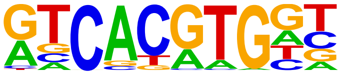 | Usf2(bHLH)/C2C12-Usf2-ChIP-Seq(GSE36030)/Homer | 1e-7 | -1.723e+01 | 0.0000 | 1019.0 | 2.82 |
| 307 | 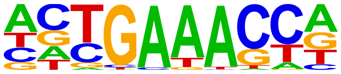 | IRF4(IRF)/GM12878-IRF4-ChIP-Seq(GSE32465)/Homer | 1e-7 | -1.698e+01 | 0.0000 | 1811.0 | 5.01 |
| 308 | 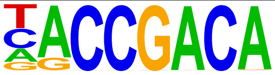 | bHLH10(bHLH)/colamp-bHLH10-DAP-Seq(GSE60143)/Homer | 1e-7 | -1.691e+01 | 0.0000 | 787.0 | 2.18 |
| 309 | 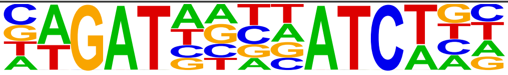 | GATA(Zf),IR4/iTreg-Gata3-ChIP-Seq(GSE20898)/Homer | 1e-7 | -1.685e+01 | 0.0000 | 370.0 | 1.02 |
| 310 | 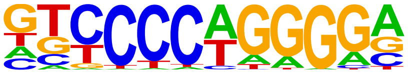 | EBF1(EBF)/Near-E2A-ChIP-Seq(GSE21512)/Homer | 1e-7 | -1.656e+01 | 0.0000 | 2998.0 | 8.30 |
| 311 | 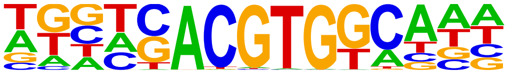 | GBF5(bZIP)/colamp-GBF5-DAP-Seq(GSE60143)/Homer | 1e-7 | -1.639e+01 | 0.0000 | 1073.0 | 2.97 |
| 312 | 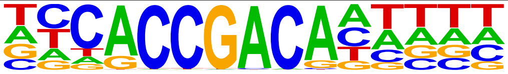 | At4g32800(AP2EREBP)/colamp-At4g32800-DAP-Seq(GSE60143)/Homer | 1e-7 | -1.631e+01 | 0.0000 | 304.0 | 0.84 |
| 313 | 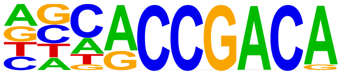 | TINY(AP2EREBP)/col-TINY-DAP-Seq(GSE60143)/Homer | 1e-7 | -1.625e+01 | 0.0000 | 732.0 | 2.03 |
| 314 | 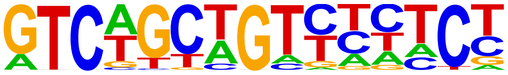 | ZNF317(Zf)/HEK293-ZNF317.GFP-ChIP-Seq(GSE58341)/Homer | 1e-7 | -1.617e+01 | 0.0000 | 410.0 | 1.14 |
| 315 | 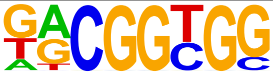 | ESE3(AP2EREBP)/col-ESE3-DAP-Seq(GSE60143)/Homer | 1e-6 | -1.596e+01 | 0.0000 | 6182.0 | 17.1 |
| 316 | 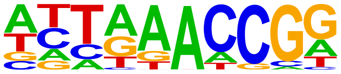 | Unknown3/Arabidopsis-Promoters/Homer | 1e-6 | -1.587e+01 | 0.0000 | 467.0 | 1.29 |
| 317 | 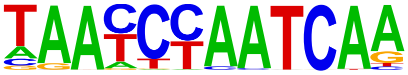 | Duxbl(Homeobox)/NIH3T3-Duxbl.HA-ChIP-Seq(GSE119782)/Homer | 1e-6 | -1.584e+01 | 0.0000 | 225.0 | 0.62 |
| 318 | 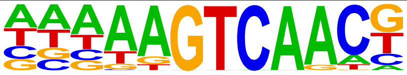 | WRKY22(WRKY)/colamp-WRKY22-DAP-Seq(GSE60143)/Homer | 1e-6 | -1.575e+01 | 0.0000 | 1674.0 | 4.63 |

| | | | | | | | |
|---|---|---|---|---|---|---|---|
| 319 | 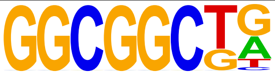 | ERF1(AP2EREBP)/colamp-ERF1-DAP-Seq(GSE60143)/Homer | 1e-6 | -1.554e+01 | 0.0000 | 3835.0 | 10.6 |
| 320 | 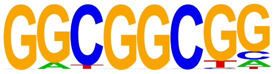 | ESE1(AP2EREBP)/col-ESE1-DAP-Seq(GSE60143)/Homer | 1e-6 | -1.552e+01 | 0.0000 | 4504.0 | 12.4 |
| 321 | 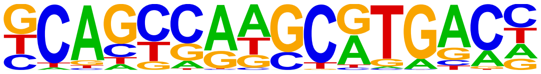 | PAX5(Paired,Homeobox)/GM12878-PAX5-ChIP-Seq(GSE32465)/Homer | 1e-6 | -1.495e+01 | 0.0000 | 1565.0 | 4.33 |
| 322 | 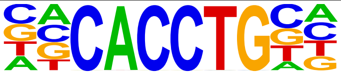 | E2A(bHLH),near_PU.1/Bcell-PU.1-ChIP-Seq(GSE21512)/Homer | 1e-6 | -1.488e+01 | 0.0000 | 5271.0 | 14.5 |
| 323 | 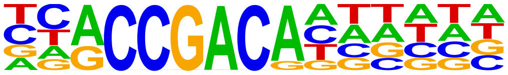 | At2g44940(AP2EREBP)/colamp-At2g44940-DAP-Seq(GSE60143)/Homer | 1e-6 | -1.465e+01 | 0.0000 | 548.0 | 1.52 |
| 324 | 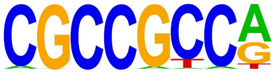 | CRF4(AP2EREBP)/colamp-CRF4-DAP-Seq(GSE60143)/Homer | 1e-6 | -1.420e+01 | 0.0000 | 3399.0 | 9.41 |
| 325 | 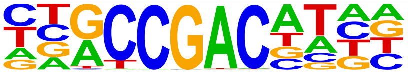 | CBF3(AP2EREBP)/colamp-CBF3-DAP-Seq(GSE60143)/Homer | 1e-6 | -1.405e+01 | 0.0000 | 994.0 | 2.75 |
| 326 | 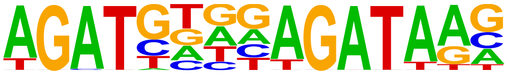 | GATA3(Zf),DR4/iTreg-Gata3-ChIP-Seq(GSE20898)/Homer | 1e-6 | -1.400e+01 | 0.0000 | 325.0 | 0.90 |
| 327 | 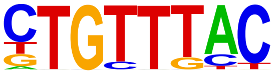 | Foxo1(Forkhead)/RAW-Foxo1-ChIP-Seq(Fan_et_al.)/Homer | 1e-6 | -1.400e+01 | 0.0000 | 7248.0 | 20.0 |
| 328 | 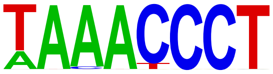 | TRP2(MYBrelated)/colamp-TRP2-DAP-Seq(GSE60143)/Homer | 1e-6 | -1.398e+01 | 0.0000 | 553.0 | 1.53 |
| 329 | 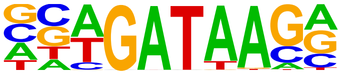 | Gata4(Zf)/Heart-Gata4-ChIP-Seq(GSE35151)/Homer | 1e-5 | -1.351e+01 | 0.0000 | 3758.0 | 10.4 |
| 330 | 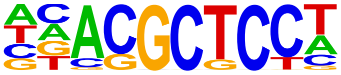 | MYB88(MYB)/col-MYB88-DAP-Seq(GSE60143)/Homer | 1e-5 | -1.345e+01 | 0.0000 | 7849.0 | 21.7 |
| 331 | 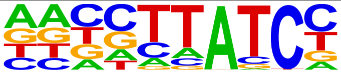 | At5g08520(MYBrelated)/colamp-At5g08520-DAP-Seq(GSE60143)/Homer | 1e-5 | -1.336e+01 | 0.0000 | 3506.0 | 9.71 |
| 332 | 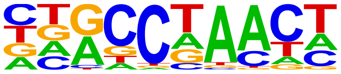 | BOS1(MYB)/col-BOS1-DAP-Seq(GSE60143)/Homer | 1e-5 | -1.316e+01 | 0.0000 | 2710.0 | 7.50 |
| 333 | 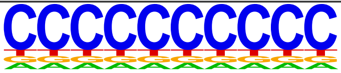 | SeqBias: polyC-repeat | 1e-5 | -1.308e+01 | 0.0000 | 35785.0 | 99.0 |
| 334 | 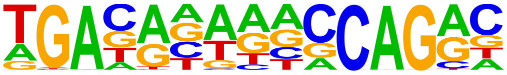 | Hand2(bHLH)/Mesoderm-Hand2-ChIP-Seq(GSE61475)/Homer | 1e-5 | -1.299e+01 | 0.0000 | 1509.0 | 4.18 |
| 335 | 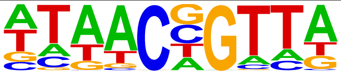 | MYB70(MYB)/col-MYB70-DAP-Seq(GSE60143)/Homer | 1e-5 | -1.261e+01 | 0.0000 | 6407.0 | 17.7 |
| 336 | 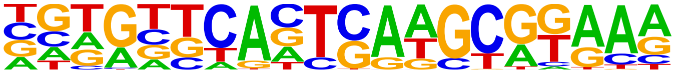 | PAX6(Paired,Homeobox)/Forebrain-Pax6-ChIP-Seq(GSE66961)/Homer | 1e-5 | -1.260e+01 | 0.0000 | 378.0 | 1.05 |
| 337 | 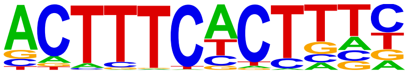 | PRDM1(Zf)/Hela-PRDM1-ChIP-Seq(GSE31477)/Homer | 1e-5 | -1.258e+01 | 0.0000 | 1817.0 | 5.03 |
| 338 | 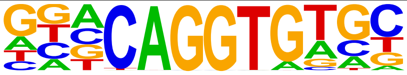 | ZEB2(Zf)/SNU398-ZEB2-ChIP-Seq(GSE103048)/Homer | 1e-5 | -1.242e+01 | 0.0000 | 3343.0 | 9.25 |
| 339 | 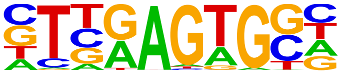 | Nkx2.2(Homeobox)/NPC-Nkx2.2-ChIP-Seq(GSE61673)/Homer | 1e-5 | -1.230e+01 | 0.0000 | 6672.0 | 18.4 |
| 340 | 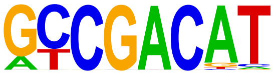 | DDF1(AP2EREBP)/col-DDF1-DAP-Seq(GSE60143)/Homer | 1e-5 | -1.229e+01 | 0.0000 | 905.0 | 2.51 |
| 341 | 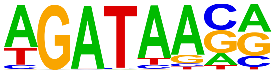 | GATA3(Zf)/iTreg-Gata3-ChIP-Seq(GSE20898)/Homer | 1e-5 | -1.216e+01 | 0.0000 | 5440.0 | 15.0 |
| 342 | 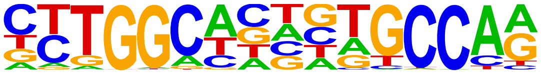 | NF1(CTF)/LNCAP-NF1-ChIP-Seq(Unpublished)/Homer | 1e-5 | -1.211e+01 | 0.0000 | 1114.0 | 3.08 |
| 343 | | AT5G61620(MYBrelated)/colamp-AT5G61620-DAP- | 1e-5 | -1.200e+01 | 0.0000 | 4594.0 | 12.7 |

| | | | | | | | |
|---|---|---|---|---|---|---|---|
| | 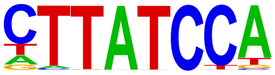 | Seq(GSE60143)/Homer | | | | | |
| 344 | 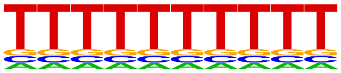 | VRN1(ABI3VP1)/col-VRN1-DAP-Seq(GSE60143)/Homer | 1e-5 | -1.199e+01 | 0.0000 | 993.0 | 2.75 |
| 345 | 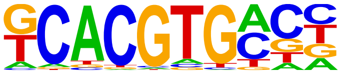 | bHLHE41(bHLH)/proB-Bhlhe41-ChIP-Seq(GSE93764)/Homer | 1e-5 | -1.192e+01 | 0.0000 | 4640.0 | 12.8 |
| 346 | 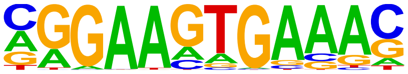 | PU.1-IRF(ETS:IRF)/Bcell-PU.1-ChIP-Seq(GSE21512)/Homer | 1e-5 | -1.181e+01 | 0.0000 | 4780.0 | 13.2 |
| 347 | 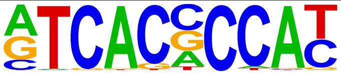 | Srebp1a(bHLH)/HepG2-Srebp1a-ChIP-Seq(GSE31477)/Homer | 1e-5 | -1.171e+01 | 0.0000 | 684.0 | 1.89 |
| 348 | 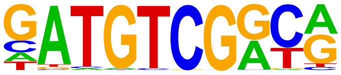 | DDF2(AP2EREBP)/col-DDF2-DAP-Seq(GSE60143)/Homer | 1e-5 | -1.159e+01 | 0.0000 | 116.0 | 0.32 |
| 349 | 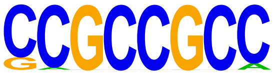 | ERF73(AP2EREBP)/col-ERF73-DAP-Seq(GSE60143)/Homer | 1e-4 | -1.151e+01 | 0.0000 | 4260.0 | 11.7 |
| 350 | 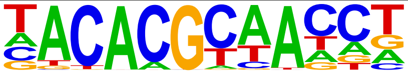 | ANAC079(NAC)/colamp-ANAC079-DAP-Seq(GSE60143)/Homer | 1e-4 | -1.139e+01 | 0.0000 | 1481.0 | 4.10 |
| 351 | 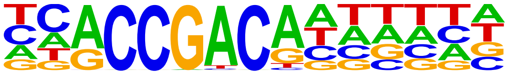 | AT3G16280(AP2EREBP)/colamp-AT3G16280-DAP-Seq(GSE60143)/Homer | 1e-4 | -1.109e+01 | 0.0000 | 848.0 | 2.35 |
| 352 | 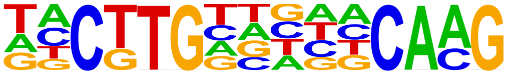 | ANAC017(NAC)/colamp-ANAC017-DAP-Seq(GSE60143)/Homer | 1e-4 | -1.108e+01 | 0.0000 | 297.0 | 0.82 |
| 353 | 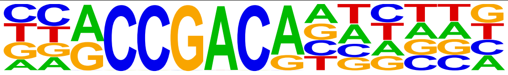 | AT3G60490(AP2EREBP)/colamp-AT3G60490-DAP-Seq(GSE60143)/Homer | 1e-4 | -1.100e+01 | 0.0000 | 754.0 | 2.09 |
| 354 | 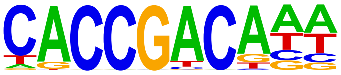 | At4g31060(AP2EREBP)/colamp-At4g31060-DAP-Seq(GSE60143)/Homer | 1e-4 | -1.083e+01 | 0.0001 | 1020.0 | 2.82 |
| 355 | 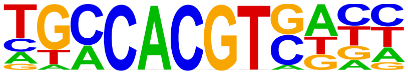 | bZIP28(bZIP)/col-bZIP28-DAP-Seq(GSE60143)/Homer | 1e-4 | -1.079e+01 | 0.0001 | 927.0 | 2.57 |
| 356 | 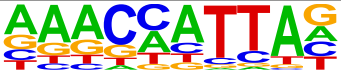 | Barx1(Homeobox)/Stomach-Barx1.3xFlag-ChIP-Seq(GSE69483)/Homer | 1e-4 | -1.077e+01 | 0.0001 | 1917.0 | 5.31 |
| 357 | 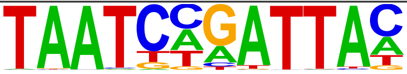 | Pax7(Paired,Homeobox),long/Myoblast-Pax7-ChIP-Seq(GSE25064)/Homer | 1e-4 | -1.071e+01 | 0.0001 | 114.0 | 0.32 |
| 358 | 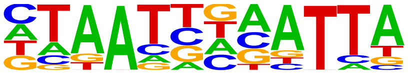 | Prop1(Homeobox)/GHFT1-PROP1.biotin-ChIP-Seq(GSE77302)/Homer | 1e-4 | -1.035e+01 | 0.0001 | 2230.0 | 6.17 |
| 359 | 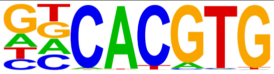 | BMAL1(bHLH)/Liver-Bmal1-ChIP-Seq(GSE39860)/Homer | 1e-4 | -1.032e+01 | 0.0001 | 6295.0 | 17.4 |
| 360 | 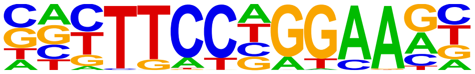 | Stat3+il21(Stat)/CD4-Stat3-ChIP-Seq(GSE19198)/Homer | 1e-4 | -1.029e+01 | 0.0001 | 2274.0 | 6.30 |
| 361 | 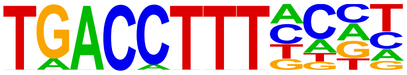 | Nur77(NR)/K562-NR4A1-ChIP-Seq(GSE31363)/Homer | 1e-4 | -1.021e+01 | 0.0001 | 640.0 | 1.77 |
| 362 | 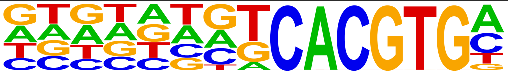 | BIM1(bHLH)/colamp-BIM1-DAP-Seq(GSE60143)/Homer | 1e-4 | -1.017e+01 | 0.0001 | 429.0 | 1.19 |
| 363 | 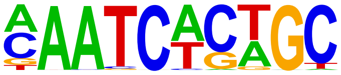 | Gfi1b(Zf)/HPC7-Gfi1b-ChIP-Seq(GSE22178)/Homer | 1e-4 | -1.012e+01 | 0.0001 | 3180.0 | 8.80 |
| 364 | 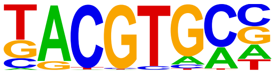 | HIF-1a(bHLH)/MCF7-HIF1a-ChIP-Seq(GSE28352)/Homer | 1e-4 | -1.007e+01 | 0.0001 | 1358.0 | 3.76 |
| 365 | 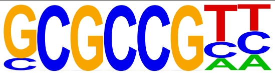 | PUCHI(AP2EREBP)/colamp-PUCHI-DAP-Seq(GSE60143)/Homer | 1e-4 | -1.001e+01 | 0.0001 | 4467.0 | 12.3 |
| 366 | 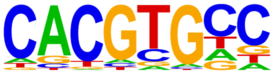 | IBL1(bHLH)/Seedling-IBL1-ChIP-Seq(GSE51120)/Homer | 1e-4 | -9.901e+00 | 0.0001 | 6743.0 | 18.6 |
| 367 | | WRKY7(WRKY)/colamp-WRKY7- | 1e-4 | -9.805e+00 | 0.0002 | 10.0 | 0.03 |

| | | | | | | | |
|---|---|---|---|---|---|---|---|
| | 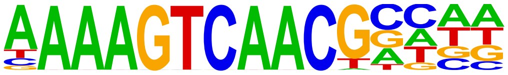 | DAP-Seq(GSE60143)/Homer | | | | | |
| 368 | 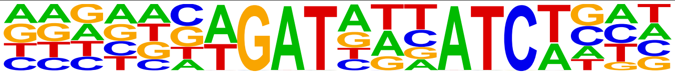 | GATA(Zf),IR3/iTreg-Gata3-ChIP-Seq(GSE20898)/Homer | 1e-4 | -9.801e+00 | 0.0002 | 569.0 | 1.58 |
| 369 | 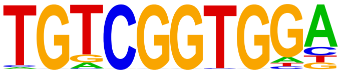 | At1g77640(AP2EREBP)/col-At1g77640-DAP-Seq(GSE60143)/Homer | 1e-4 | -9.636e+00 | 0.0002 | 537.0 | 1.49 |
| 370 | 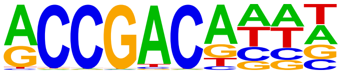 | AT1G77200(AP2EREBP)/colamp-AT1G77200-DAP-Seq(GSE60143)/Homer | 1e-4 | -9.283e+00 | 0.0003 | 1753.0 | 4.85 |
| 371 | 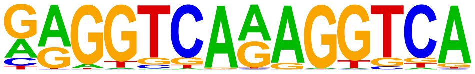 | TR4(NR),DR1/Hela-TR4-ChIP-Seq(GSE24685)/Homer | 1e-4 | -9.259e+00 | 0.0003 | 238.0 | 0.66 |
| 372 | 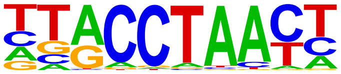 | MYB57(MYB)/col-MYB57-DAP-Seq(GSE60143)/Homer | 1e-3 | -9.201e+00 | 0.0003 | 1229.0 | 3.40 |
| 373 | 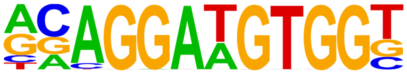 | ETS:RUNX(ETS,Runt)/Jurkat-RUNX1-ChIP-Seq(GSE17954)/Homer | 1e-3 | -9.111e+00 | 0.0003 | 227.0 | 0.63 |
| 374 | 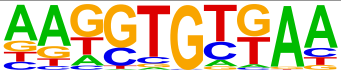 | Tbr1(T-box)/Cortex-Tbr1-ChIP-Seq(GSE71384)/Homer | 1e-3 | -9.008e+00 | 0.0003 | 5362.0 | 14.8 |
| 375 | 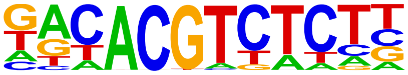 | bHLH157(bHLH)/col-bHLH157-DAP-Seq(GSE60143)/Homer | 1e-3 | -8.974e+00 | 0.0003 | 875.0 | 2.42 |
| 376 | 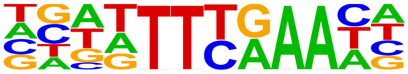 | TCX2(CPP)/colamp-TCX2-DAP-Seq(GSE60143)/Homer | 1e-3 | -8.768e+00 | 0.0004 | 5110.0 | 14.1 |
| 377 | 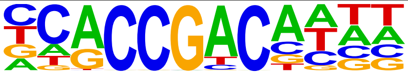 | RAP21(AP2EREBP)/colamp-RAP21-DAP-Seq(GSE60143)/Homer | 1e-3 | -8.722e+00 | 0.0004 | 414.0 | 1.15 |
| 378 | 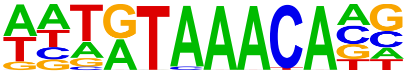 | FoxL2(Forkhead)/Ovary-FoxL2-ChIP-Seq(GSE60858)/Homer | 1e-3 | -8.701e+00 | 0.0004 | 2893.0 | 8.01 |
| 379 | 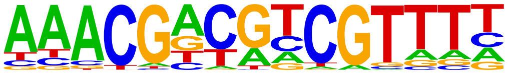 | Unknown2/Arabidopsis-Promoters/Homer | 1e-3 | -8.691e+00 | 0.0004 | 19.0 | 0.05 |
| 380 | 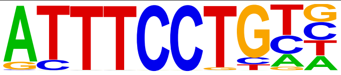 | EWS:ERG-fusion(ETS)/CADO_ES1-EWS:ERG-ChIP-Seq(SRA014231)/Homer | 1e-3 | -8.638e+00 | 0.0005 | 2325.0 | 6.44 |
| 381 | 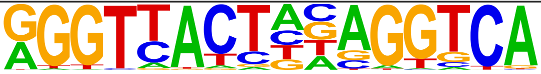 | LXRE(NR),DR4/RAW-LXRb.biotin-ChIP-Seq(GSE21512)/Homer | 1e-3 | -8.600e+00 | 0.0005 | 132.0 | 0.37 |
| 382 | 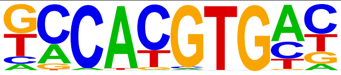 | NPAS2(bHLH)/Liver-NPAS2-ChIP-Seq(GSE39860)/Homer | 1e-3 | -8.586e+00 | 0.0005 | 3756.0 | 10.4 |
| 383 | 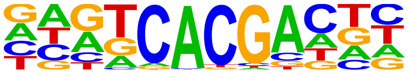 | Npas4(bHLH)/Neuron-Npas4-ChIP-Seq(GSE127793)/Homer | 1e-3 | -8.301e+00 | 0.0007 | 3558.0 | 9.85 |
| 384 | 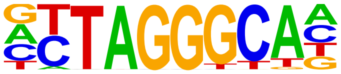 | TBP3(MYBrelated)/col-TBP3-DAP-Seq(GSE60143)/Homer | 1e-3 | -8.247e+00 | 0.0007 | 1808.0 | 5.01 |
| 385 | 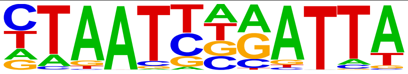 | Phox2a(Homeobox)/Neuron-Phox2a-ChIP-Seq(GSE31456)/Homer | 1e-3 | -8.028e+00 | 0.0009 | 1348.0 | 3.73 |
| 386 | 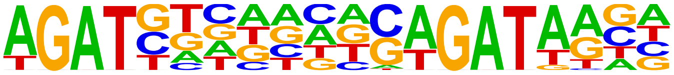 | GATA3(Zf),DR8/iTreg-Gata3-ChIP-Seq(GSE20898)/Homer | 1e-3 | -8.002e+00 | 0.0009 | 306.0 | 0.85 |
| 387 | 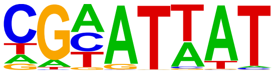 | AtHB32(ZFHD)/col200-AtHB32-DAP-Seq(GSE60143)/Homer | 1e-3 | -7.793e+00 | 0.0011 | 4320.0 | 11.9 |
| 388 | 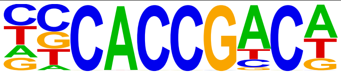 | AT1G44830(AP2EREBP)/col-AT1G44830-DAP-Seq(GSE60143)/Homer | 1e-3 | -7.662e+00 | 0.0012 | 1019.0 | 2.82 |
| 389 | 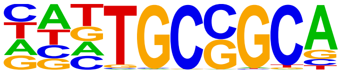 | Zfp57(Zf)/H1-ZFP57.HA-ChIP-Seq(GSE115387)/Homer | 1e-3 | -7.645e+00 | 0.0012 | 2185.0 | 6.05 |
| 390 | 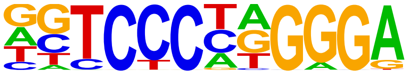 | EBF(EBF)/proBcell-EBF-ChIP-Seq(GSE21978)/Homer | 1e-3 | -7.576e+00 | 0.0013 | 537.0 | 1.49 |
| 391 | | CBF2(AP2EREBP)/colamp-CBF2-DAP- | 1e-3 | -7.509e+00 | 0.0014 | 965.0 | 2.67 |

| | | | | | | | |
|---|---|---|---|---|---|---|---|
| | 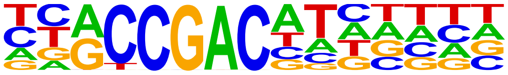 | Seq(GSE60143)/Homer | | | | | |
| 392 | 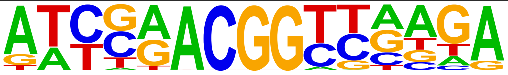 | ZIM(C2C2gata)/col-ZIM-DAP-Seq(GSE60143)/Homer | 1e-3 | -7.398e+00 | 0.0016 | 27.0 | 0.07 |
| 393 | 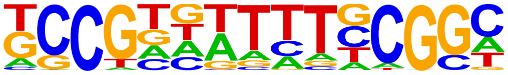 | LBD13(LOBAS2)/colamp-LBD13-DAP-Seq(GSE60143)/Homer | 1e-3 | -7.375e+00 | 0.0016 | 3036.0 | 8.40 |
| 394 | 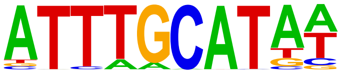 | Oct4(POU,Homeobox)/mES-Oct4-ChIP-Seq(GSE11431)/Homer | 1e-3 | -7.335e+00 | 0.0017 | 1786.0 | 4.94 |
| 395 | 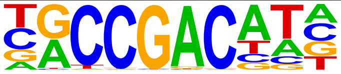 | CBF1(AP2EREBP)/colamp-CBF1-DAP-Seq(GSE60143)/Homer | 1e-3 | -7.320e+00 | 0.0017 | 1491.0 | 4.13 |
| 396 | 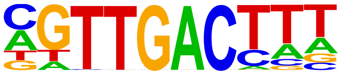 | WRKY29(WRKY)/colamp-WRKY29-DAP-Seq(GSE60143)/Homer | 1e-3 | -7.267e+00 | 0.0018 | 2620.0 | 7.25 |
| 397 | 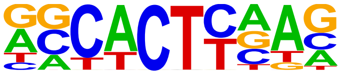 | Nkx2.1(Homeobox)/LungAC-Nkx2.1-ChIP-Seq(GSE43252)/Homer | 1e-3 | -7.216e+00 | 0.0019 | 9936.0 | 27.5 |
| 398 | 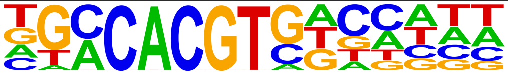 | ABF2(bZIP)/col-ABF2-DAP-Seq(GSE60143)/Homer | 1e-3 | -7.132e+00 | 0.0020 | 717.0 | 1.98 |
| 399 | 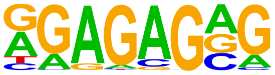 | Trl(Zf)/S2-GAGAfactor-ChIP-Seq(GSE40646)/Homer | 1e-3 | -7.094e+00 | 0.0021 | 10036.0 | 27.7 |
| 400 | 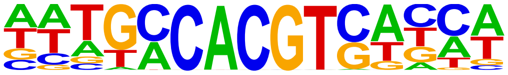 | GBF6(bZIP)/colamp-GBF6-DAP-Seq(GSE60143)/Homer | 1e-3 | -7.058e+00 | 0.0022 | 737.0 | 2.04 |
| 401 | 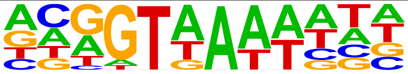 | GT2(Trihelix)/colamp-GT2-DAP-Seq(GSE60143)/Homer | 1e-3 | -7.041e+00 | 0.0022 | 3891.0 | 10.7 |
| 402 | 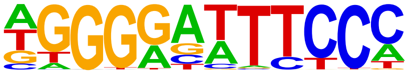 | NFkB-p65(RHD)/GM12787-p65-ChIP-Seq(GSE19485)/Homer | 1e-3 | -6.943e+00 | 0.0024 | 1420.0 | 3.93 |
| 403 | 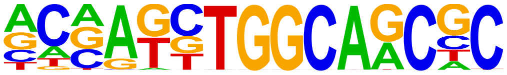 | Unknown3/Drosophila-Promoters/Homer | 1e-2 | -6.900e+00 | 0.0025 | 446.0 | 1.23 |
| 404 | 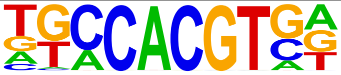 | bZIP16(bZIP)/colamp-bZIP16-DAP-Seq(GSE60143)/Homer | 1e-2 | -6.894e+00 | 0.0025 | 921.0 | 2.55 |
| 405 | 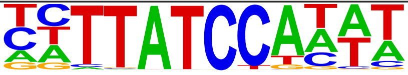 | At1g74840(MYBrelated)/col100-At1g74840-DAP-Seq(GSE60143)/Homer | 1e-2 | -6.881e+00 | 0.0026 | 2273.0 | 6.29 |
| 406 | 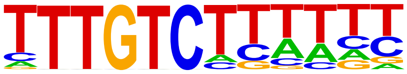 | SGR5(C2H2)/colamp-SGR5-DAP-Seq(GSE60143)/Homer | 1e-2 | -6.813e+00 | 0.0027 | 2732.0 | 7.56 |
| 407 | 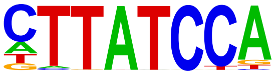 | At5g47390(MYBrelated)/col-At5g47390-DAP-Seq(GSE60143)/Homer | 1e-2 | -6.772e+00 | 0.0028 | 3952.0 | 10.9 |
| 408 | 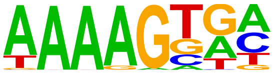 | CDF3(C2C2dof)/colamp-CDF3-DAP-Seq(GSE60143)/Homer | 1e-2 | -6.710e+00 | 0.0030 | 6895.0 | 19.0 |
| 409 | 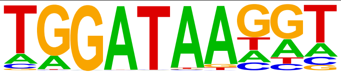 | AT5G56840(MYBrelated)/colamp-AT5G56840-DAP-Seq(GSE60143)/Homer | 1e-2 | -6.652e+00 | 0.0032 | 4003.0 | 11.0 |
| 410 | 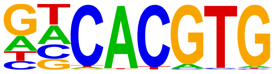 | CLOCK(bHLH)/Liver-Clock-ChIP-Seq(GSE39860)/Homer | 1e-2 | -6.631e+00 | 0.0032 | 1939.0 | 5.37 |
| 411 | 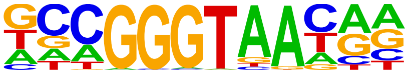 | REB1/SacCer-Promoters/Homer | 1e-2 | -6.606e+00 | 0.0033 | 523.0 | 1.45 |
| 412 | 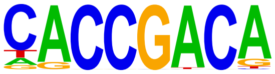 | ERF38(AP2EREBP)/col-ERF38-DAP-Seq(GSE60143)/Homer | 1e-2 | -6.531e+00 | 0.0036 | 1237.0 | 3.42 |
| 413 | 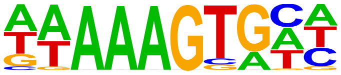 | AT1G69570(C2C2dof)/col-AT1G69570-DAP-Seq(GSE60143)/Homer | 1e-2 | -6.512e+00 | 0.0036 | 4685.0 | 12.9 |
| 414 | 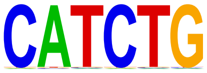 | Tal1 | 1e-2 | -6.472e+00 | 0.0038 | 5895.0 | 16.3 |
| 415 | 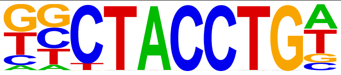 | Zelda(Zf)/Embryo-zld-ChIP-Seq(GSE65441)/Homer | 1e-2 | -6.418e+00 | 0.0040 | 1664.0 | 4.61 |

| | | | | | | | |
|---|---|---|---|---|---|---|---|
| 416 | 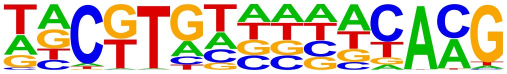 | CUC2(NAC)/colamp-CUC2-DAP-Seq(GSE60143)/Homer | 1e-2 | -6.300e+00 | 0.0044 | 936.0 | 2.59 |
| 417 | 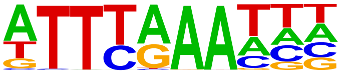 | SOL1(CPP)/colamp-SOL1-DAP-Seq(GSE60143)/Homer | 1e-2 | -6.206e+00 | 0.0049 | 4924.0 | 13.6 |
| 418 | 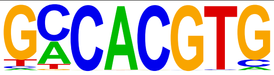 | E-box/Arabidopsis-Promoters/Homer | 1e-2 | -6.130e+00 | 0.0052 | 1767.0 | 4.89 |
| 419 | 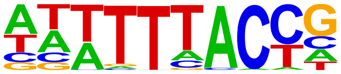 | AT5G47660(Trihelix)/colamp-AT5G47660-DAP-Seq(GSE60143)/Homer | 1e-2 | -6.038e+00 | 0.0057 | 5184.0 | 14.3 |
| 420 | 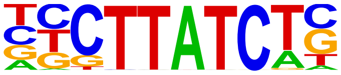 | Gata2(Zf)/K562-GATA2-ChIP-Seq(GSE18829)/Homer | 1e-2 | -6.028e+00 | 0.0058 | 2347.0 | 6.50 |
| 421 | 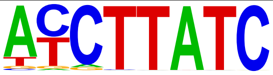 | At5g05790(MYBrelated)/col-At5g05790-DAP-Seq(GSE60143)/Homer | 1e-2 | -5.994e+00 | 0.0060 | 2964.0 | 8.21 |
| 422 | 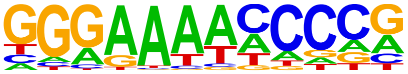 | Dorsal(RHD)/Embryo-dl-ChIP-Seq(GSE65441)/Homer | 1e-2 | -5.993e+00 | 0.0060 | 626.0 | 1.73 |
| 423 | 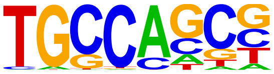 | HIC1(Zf)/Treg-ZBTB29-ChIP-Seq(GSE99889)/Homer | 1e-2 | -5.925e+00 | 0.0064 | 8804.0 | 24.3 |
| 424 | 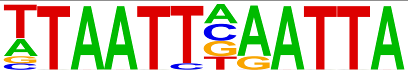 | Phox2b(Homeobox)/CLBGA-PHOX2B-ChIP-Seq(GSE90683)/Homer | 1e-2 | -5.915e+00 | 0.0064 | 675.0 | 1.87 |
| 425 | 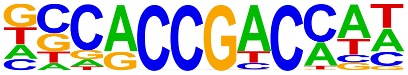 | At4g28140(AP2EREBP)/colamp-At4g28140-DAP-Seq(GSE60143)/Homer | 1e-2 | -5.819e+00 | 0.0070 | 909.0 | 2.52 |
| 426 | 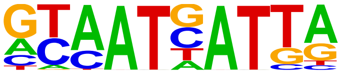 | ATHB15(HB)/col-ATHB15-DAP-Seq(GSE60143)/Homer | 1e-2 | -5.817e+00 | 0.0070 | 1005.0 | 2.78 |
| 427 | 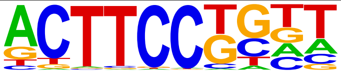 | Elf4(ETS)/BMDM-Elf4-ChIP-Seq(GSE88699)/Homer | 1e-2 | -5.809e+00 | 0.0071 | 2987.0 | 8.27 |
| 428 | 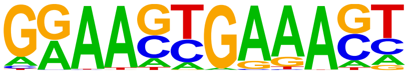 | IRF8(IRF)/BMDM-IRF8-ChIP-Seq(GSE77884)/Homer | 1e-2 | -5.770e+00 | 0.0073 | 996.0 | 2.76 |
| 429 | 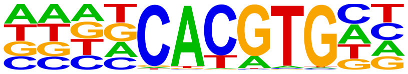 | BIM2(bHLH)/col-BIM2-DAP-Seq(GSE60143)/Homer | 1e-2 | -5.717e+00 | 0.0077 | 3119.0 | 8.63 |
| 430 | 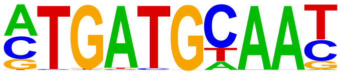 | Atf4(bZIP)/MEF-Atf4-ChIP-Seq(GSE35681)/Homer | 1e-2 | -5.702e+00 | 0.0078 | 1299.0 | 3.60 |
| 431 | 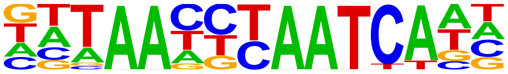 | DUX4(Homeobox)/Myoblasts-DUX4.V5-ChIP-Seq(GSE75791)/Homer | 1e-2 | -5.689e+00 | 0.0079 | 110.0 | 0.30 |
| 432 | 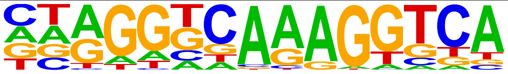 | PPARa(NR),DR1/Liver-Ppara-ChIP-Seq(GSE47954)/Homer | 1e-2 | -5.685e+00 | 0.0079 | 2776.0 | 7.69 |
| 433 | 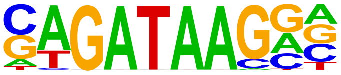 | Gata1(Zf)/K562-GATA1-ChIP-Seq(GSE18829)/Homer | 1e-2 | -5.666e+00 | 0.0080 | 2114.0 | 5.85 |
| 434 | 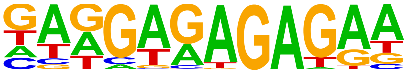 | BPC1(BBRBPC)/colamp-BPC1-DAP-Seq(GSE60143)/Homer | 1e-2 | -5.602e+00 | 0.0086 | 1902.0 | 5.27 |
| 435 | 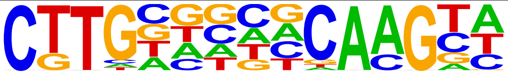 | AT1G19040(NAC)/col-AT1G19040-DAP-Seq(GSE60143)/Homer | 1e-2 | -5.394e+00 | 0.0105 | 285.0 | 0.79 |
| 436 | 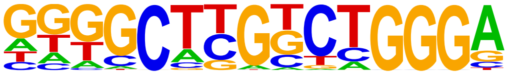 | Zfp809(Zf)/ES-Zfp809-ChIP-Seq(GSE70799)/Homer | 1e-2 | -5.390e+00 | 0.0105 | 736.0 | 2.04 |
| 437 | 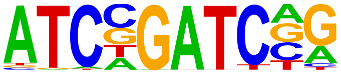 | GATA19(C2C2gata)/colamp-GATA19-DAP-Seq(GSE60143)/Homer | 1e-2 | -5.277e+00 | 0.0118 | 260.0 | 0.72 |
| 438 | 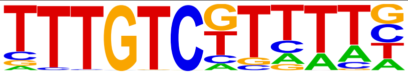 | IDD7(C2H2)/col-IDD7-DAP-Seq(GSE60143)/Homer | 1e-2 | -5.238e+00 | 0.0122 | 1455.0 | 4.03 |
| 439 | 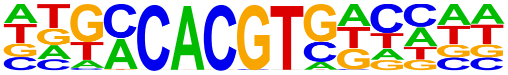 | AREB3(bZIP)/col-AREB3-DAP-Seq(GSE60143)/Homer | 1e-2 | -5.176e+00 | 0.0130 | 993.0 | 2.75 |
| 440 | 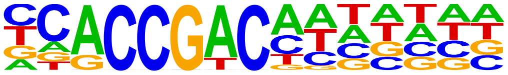 | DEAR3(AP2EREBP)/colamp-DEAR3-DAP-Seq(GSE60143)/Homer | 1e-2 | -5.104e+00 | 0.0139 | 697.0 | 1.93 |

| | | | | | | | |
|---|---|---|---|---|---|---|---|
| 441 | 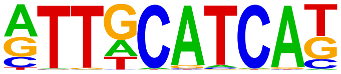 | Chop(bZIP)/MEF-Chop-ChIP-Seq(GSE35681)/Homer | 1e-2 | -5.065e+00 | 0.0144 | 978.0 | 2.71 |
| 442 | 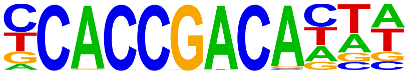 | AT1G01250(AP2EREBP)/col-AT1G01250-DAP-Seq(GSE60143)/Homer | 1e-2 | -5.064e+00 | 0.0144 | 230.0 | 0.64 |
| 443 | 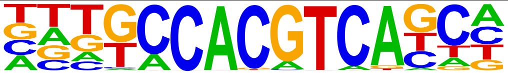 | bZIP44(bZIP)/colamp-bZIP44-DAP-Seq(GSE60143)/Homer | 1e-2 | -5.007e+00 | 0.0152 | 89.0 | 0.25 |
| 444 | 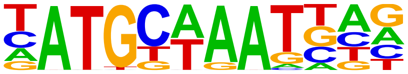 | Brn1(POU,Homeobox)/NPC-Brn1-ChIP-Seq(GSE35496)/Homer | 1e-2 | -4.991e+00 | 0.0154 | 1097.0 | 3.04 |
| 445 | 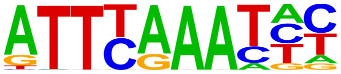 | AT2G20110(CPP)/colamp-AT2G20110-DAP-Seq(GSE60143)/Homer | 1e-2 | -4.973e+00 | 0.0157 | 4864.0 | 13.4 |
| 446 | 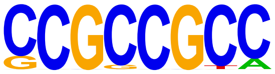 | SHN3(AP2EREBP)/col-SHN3-DAP-Seq(GSE60143)/Homer | 1e-2 | -4.919e+00 | 0.0165 | 2844.0 | 7.87 |
| 447 | 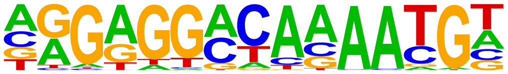 | ZNF675(Zf)/HEK293-ZNF675.GFP-ChIP-Seq(GSE58341)/Homer | 1e-2 | -4.737e+00 | 0.0197 | 425.0 | 1.18 |
| 448 | 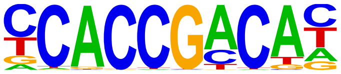 | DREB26(AP2EREBP)/col-DREB26-DAP-Seq(GSE60143)/Homer | 1e-2 | -4.713e+00 | 0.0202 | 784.0 | 2.17 |
| 449 | 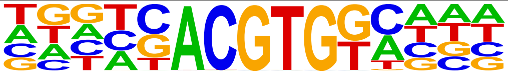 | bZIP53(bZIP)/colamp-bZIP53-DAP-Seq(GSE60143)/Homer | 1e-2 | -4.686e+00 | 0.0207 | 1031.0 | 2.85 |
| 450 | 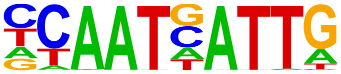 | ATHB18(Homeobox)/colamp-ATHB18-DAP-Seq(GSE60143)/Homer | 1e-2 | -4.681e+00 | 0.0207 | 541.0 | 1.50 |
| 451 | 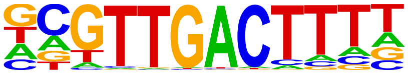 | WRKY21(WRKY)/colamp-WRKY21-DAP-Seq(GSE60143)/Homer | 1e-2 | -4.672e+00 | 0.0209 | 250.0 | 0.69 |

S6 Table. Full list of de novo motif discovery from ChIP-seq data targeting PBX3.

### Homer *de novo* Motif Results (/wynton/group/marcucio/2022CHM/Data2022Mar/Motif/HomerPBX3IDR/)

Known Motif Enrichment Results
Gene Ontology Enrichment Results
If Homer is having trouble matching a motif to a known motif, try copy/pasting the matrix file into STAMP
More information on motif finding results: HOMER | Description of Results | Tips
Total target sequences = 36123
Total background sequences = 35367
* - possible false positive

| Rank | Motif | P-value | log P-pvalue | % of Targets | % of Background | STD(Bg STD) | Best Match/Details | Motif File |
|---|---|---|---|---|---|---|---|---|
| 1 | 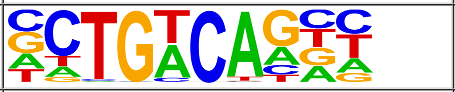 | 1e-3291 | -7.579e+03 | 49.34% | 20.25% | 49.3bp (64.6bp) | Meis1(Homeobox)/MastCells-Meis1-ChIP-Seq(GSE48085)/Homer(0.962) More Information \| Similar Motifs Found | motif file (matrix) |
| 2 | 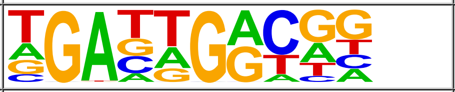 | 1e-1664 | -3.832e+03 | 18.73% | 5.54% | 51.6bp (62.7bp) | Pknox1(Homeobox)/ES-Prep1-ChIP-Seq(GSE63282)/Homer(0.860) More Information \| Similar Motifs Found | motif file (matrix) |
| 3 | 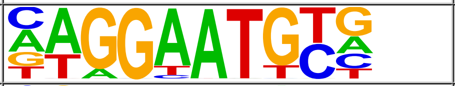 | 1e-618 | -1.425e+03 | 12.68% | 5.33% | 53.2bp (59.8bp) | TEAD3/MA0808.1/Jaspar(0.964) More Information \| Similar Motifs Found | motif file (matrix) |
| 4 | 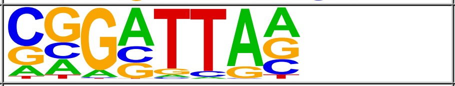 | 1e-457 | -1.054e+03 | 31.30% | 20.98% | 55.8bp (61.0bp) | OTX1/MA0711.1/Jaspar(0.959) More Information \| Similar Motifs Found | motif file (matrix) |
| 5 | 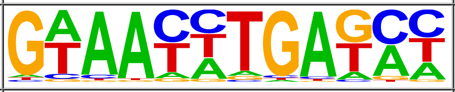 | 1e-262 | -6.044e+02 | 6.41% | 2.88% | 53.9bp (58.5bp) | Six1(Homeobox)/Myoblast-Six1-ChIP-Chip(GSE20150)/Homer(0.975) More Information \| Similar Motifs Found | motif file (matrix) |
| 6 | 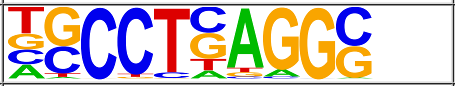 | 1e-252 | -5.808e+02 | 7.94% | 3.99% | 53.5bp (60.7bp) | TFAP2A/MA0003.4/Jaspar(0.948) More Information \| Similar Motifs Found | motif file (matrix) |
| 7 | 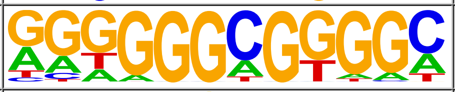 | 1e-239 | -5.520e+02 | 11.50% | 6.73% | 53.5bp (60.9bp) | KLF14(Zf)/HEK293-KLF14.GFP-ChIP-Seq(GSE58341)/Homer(0.955) More Information \| Similar Motifs Found | motif file (matrix) |
| 8 | 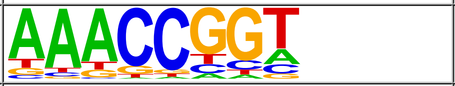 | 1e-233 | -5.387e+02 | 28.96% | 21.62% | 56.4bp (60.8bp) | TFCP2/MA0145.3/Jaspar(0.908) More Information \| Similar Motifs Found | motif file (matrix) |
| 9 | 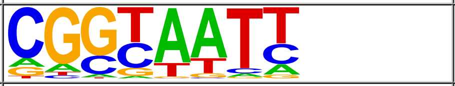 | 1e-196 | -4.526e+02 | 23.52% | 17.31% | 55.6bp (62.2bp) | PH0024.1_Dlx5/Jaspar(0.779) More Information \| Similar Motifs Found | motif file (matrix) |
| 10 | 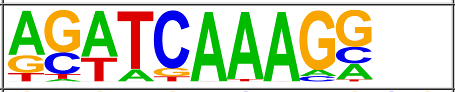 | 1e-166 | -3.844e+02 | 5.85% | 3.05% | 55.1bp (60.2bp) | LEF1(HMG)/H1-LEF1-ChIP-Seq(GSE64758)/Homer(0.988) More Information \| Similar Motifs Found | motif file (matrix) |
| 11 | 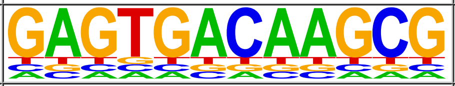 | 1e-132 | -3.054e+02 | 2.19% | 0.80% | 53.0bp (66.2bp) | PBX3/MA1114.1/Jaspar(0.771) More Information \| Similar Motifs Found | motif file (matrix) |
| 12 | 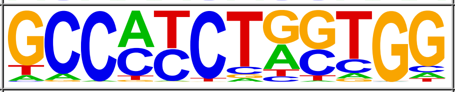 | 1e-125 | -2.901e+02 | 2.07% | 0.75% | 50.7bp (59.8bp) | BORIS(Zf)/K562-CTCFL-ChIP-Seq(GSE32465)/Homer(0.923) More Information \| Similar Motifs Found | motif file (matrix) |
| 13 | 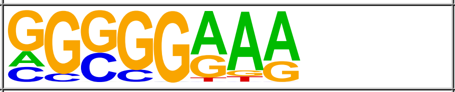 | 1e-94 | -2.168e+02 | 28.68% | 23.95% | 56.9bp (63.1bp) | RO3G_00049(RRM)/Rhizopus_oryzae-RNCMPT00205-PBM/HughesRNA(0.817) More Information \| Similar Motifs Found | motif file (matrix) |
| 14 | 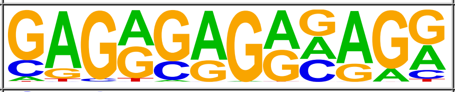 | 1e-80 | -1.842e+02 | 6.44% | 4.28% | 54.7bp (55.3bp) | FRS9(ND)/col-FRS9-DAP-Seq(GSE60143)/Homer(0.848) More Information \| Similar Motifs Found | motif file (matrix) |
| 15 | 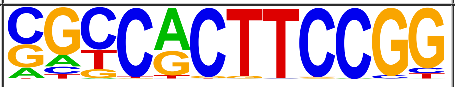 | 1e-74 | -1.710e+02 | 1.20% | 0.43% | 53.4bp (56.7bp) | ZBTB7A/MA0750.2/Jaspar(0.945) More Information \| Similar Motifs Found | motif file (matrix) |
| 16 | 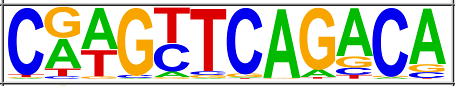 | 1e-65 | -1.512e+02 | 0.17% | 0.01% | 55.4bp (42.3bp) | PB0060.1_Smad3_1/Jaspar(0.707) More Information \| Similar Motifs Found | motif file (matrix) |
| 17 | 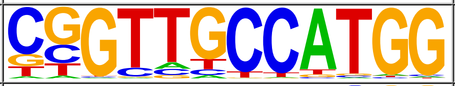 | 1e-54 | -1.255e+02 | 0.73% | 0.23% | 52.8bp (55.6bp) | RFX3/MA0798.2/Jaspar(0.929) More Information \| Similar Motifs Found | motif file (matrix) |
| 18 | 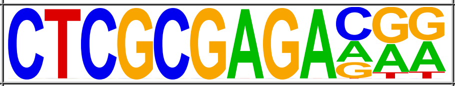 | 1e-51 | -1.180e+02 | 0.14% | 0.01% | 55.7bp (0.0bp) | GFX(?)/Promoter/Homer(0.907) More Information \| Similar Motifs Found | motif file (matrix) |
| | | | | | | | FEA4(bZIP)/Corn-FEA4-ChIP- | |

| | | | | | | | | |
|---|---|---|---|---|---|---|---|---|
| 19 | 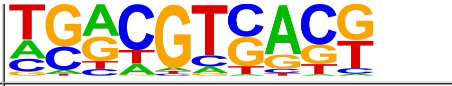 | 1e-45 | -1.036e+02 | 0.71% | 0.25% | 52.5bp (65.1bp) | Seq(GSE61954)/Homer(0.872) More Information \| Similar Motifs Found | motif file (matrix) |
| 20 | 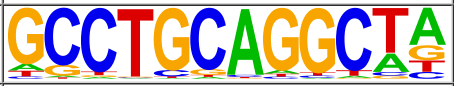 | 1e-39 | -9.116e+01 | 1.45% | 0.77% | 53.8bp (62.9bp) | SOK2/MA0385.1/Jaspar(0.768) More Information \| Similar Motifs Found | motif file (matrix) |
| 21 | 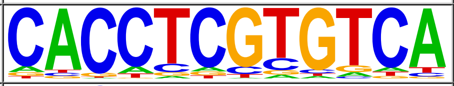 | 1e-38 | -8.976e+01 | 0.21% | 0.03% | 51.8bp (28.5bp) | HLHm5/dmmpmm(Pollard)/fly(0.631) More Information \| Similar Motifs Found | motif file (matrix) |
| 22 | 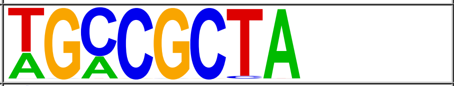 | 1e-34 | -7.916e+01 | 3.45% | 2.39% | 57.1bp (67.1bp) | STP3/MA0396.1/Jaspar(0.780) More Information \| Similar Motifs Found | motif file (matrix) |
| 23 | 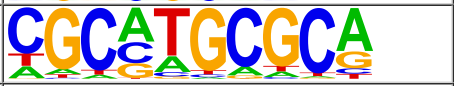 | 1e-33 | -7.647e+01 | 1.05% | 0.53% | 53.6bp (69.3bp) | NRF1/MA0506.1/Jaspar(0.932) More Information \| Similar Motifs Found | motif file (matrix) |
| 24 | 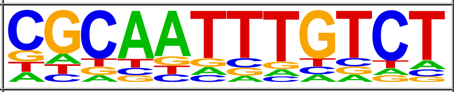 | 1e-32 | -7.376e+01 | 0.12% | 0.01% | 47.9bp (49.5bp) | SGR5(C2H2)/colamp-SGR5-DAP-Seq(GSE60143)/Homer(0.730) More Information \| Similar Motifs Found | motif file (matrix) |
| 25 | 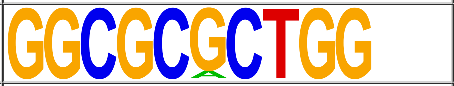 | 1e-23 | -5.348e+01 | 0.11% | 0.01% | 58.4bp (57.7bp) | POL006.1_BREu/Jaspar(0.797) More Information \| Similar Motifs Found | motif file (matrix) |
| 26 | 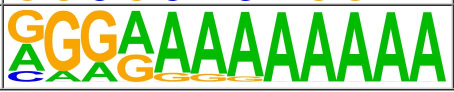 | 1e-15 | -3.503e+01 | 2.15% | 1.60% | 57.0bp (57.0bp) | REM19(REM)/colamp-REM19-DAP-Seq(GSE60143)/Homer(0.859) More Information \| Similar Motifs Found | motif file (matrix) |
